# A chromosome-level genome assembly of the European Beech (*Fagus sylvatica*) reveals anomalies for organelle DNA integration, repeat content and distribution of SNPs

**DOI:** 10.1101/2021.03.22.436437

**Authors:** Bagdevi Mishra, Bartosz Ulaszewski, Joanna Meger, Jean-Marc Aury, Catherine Bodénès, Isabelle Lesur-Kupin, Markus Pfenninger, Corinne Da Silva, Deepak K Gupta, Erwan Guichoux, Katrin Heer, Céline Lalanne, Karine Labadie, Lars Opgenoorth, Sebastian Ploch, Grégoire Le Provost, Jérôme Salse, Ivan Scotti, Stefan Wötzel, Christophe Plomion, Jaroslaw Burczyk, Marco Thines

**Affiliations:** Senckenberg Biodiversity and Climate Research Centre (BiK-F), Senckenberg Gesellschaft für Naturforschung, Senckenberganlage 25, D-60325 Frankfurt am Main, Germany; Goethe University, Department for Biological Sciences, Institute of Ecology, Evolution and Diversity, Max-von-Laue-Str. 9, D-60438 Frankfurt am Main, Germany; Kazimierz Wielki University, Department of Genetics, ul. Chodkiewicza 30, 85-064 Bydgoszcz, Poland; Génomique Métabolique, Genoscope, Institut François Jacob, CEA, CNRS, Univ Evry, Université Paris-Saclay, F-91057, Evry, France; INRAE, Univ. Bordeaux, BIOGECO, F-33610 Cestas, France; HelixVenture, F-33700, Mérignac, France; Philipps University Marburg, Faculty of Biology, Plant Ecology and Geobotany, 35043, Marburg, Germany; LOEWE Centre for Translational Biodiversity Genomics (TBG), Georg-Voigt-Str. 14-16, D-60325 Frankfurt am Main (Germany); INRAE, UCA, GDEC, F-63100 Clermont-Ferrand, France; INRAE, URFM, F-84914, Avignon, France

**Keywords:** Chromosomes, *Fagaceae*, genome architecture, genomics, Hi-C, repeat elements, SNPs

## Abstract

The European Beech is the dominant climax tree in most regions of Central Europe and valued for its ecological versatility and hardwood timber. Even though a draft genome has been published recently, higher resolution is required for studying aspects of genome architecture and recombination. Here we present a chromosome-level assembly of the more than 300 year-old reference individual, Bhaga, from the Kellerwald-Edersee National Park (Germany). Its nuclear genome of 541 Mb was resolved into 12 chromosomes varying in length between 28 Mb and 73 Mb. Multiple nuclear insertions of parts of the chloroplast genome were observed, with one region on chromosome 11 spanning more than 2 Mb of the genome in which fragments up to 54,784 bp long and covering the whole chloroplast genome were inserted randomly. Unlike in *Arabidopsis thaliana*, ribosomal cistrons are present in *Fagus sylvatica* only in four major regions, in line with FISH studies. On most assembled chromosomes, telomeric repeats were found at both ends, while centromeric repeats were found to be scattered throughout the genome apart from their main occurrence per chromosome. The genome-wide distribution of SNPs was evaluated using a second individual from Jamy Nature Reserve (Poland). SNPs, repeat elements and duplicated genes were unevenly distributed in the genomes, with one major anomaly on chromosome 4. The genome presented here adds to the available highly resolved plant genomes and we hope it will serve as a valuable basis for future research on genome architecture and for understanding the past and future of European Beech populations in a changing climate.

## 1. Introduction

Many lowland and mountainous forests in Central Europe are dominated by the European Beech (*Fagus sylvatica*) (Durrant et al., 2016). This tree is a shade-tolerant hardwood tree that can survive as a sapling in the understorey for decades until enough light becomes available for rapid growth and maturation (Wagner et al., 2010; Ligot et al., 2013). Beech trees reach ages of 200-300 years, but older individuals are known e.g. from suboptimal habitats, especially close to the tree line (Di Filippo et al., 2012). Under optimal water availability, European Beech is able to outcompete most other tree species, forming monospecific stands (Leuschner et al., 2006), but both stagnant soil water and drought restrict its presence in natural habitats (Jump at al., 2006; Geßler at al., 2007). Particularly, dry summers, which have recently been observed in Central Europe and that are predicted to increase as a result of climate change (Coumou and Rahmstorf, 2012; Spinoni at al., 2015), will intensify climatic stress as already now severe damage has been observed (Geßler at al., 2007; Reif at al., 2017). In order to cope with this, human intervention in facilitating regeneration of beech forests with more drought-resistant genotypes might be a useful strategy (Rose et al., 2009; Bolte and Degen, 2010). However, for the selection of drought-resistant genotypes, whole genome sequences of trees that thrive in comparatively dry conditions and the comparison with trees that are declining in drier conditions are necessary to identify genes associated with tolerating these adverse conditions (Pfenninger et al., 2020). Such genome-wide association studies rely on well-assembled reference genomes onto which genome data from large-scale resequencing projects can be mapped (e.g. (Atwell et al., 2010)).

Due to advances in library construction and sequencing, chromosome-level assemblies have been achieved for a variety of genomes from various kingdoms of live, including animals (Michael and VanBuren, 2020; Priest at al., 2020; Rhie at al., 2020). While the combination of short- and long-read sequencing has brought about a significant improvement in the assembly of the gene space and regions with moderate repeat-element presence, chromosome conformation information libraries, such as Hi-C (Lieberman-Aiden et al., 2009), have enabled associating scaffolds across highly repetitive regions, enabling the construction of super-scaffolds of chromosomal scale (e.g. (Yin et al., 2020)). Recently, the first chromosome-level assemblies have been published for tree and shrub species, e.g. the tea tree (*Camellia sinensis* (Chen et al., 2020)), loquat (*Eriobotrya japonica* (Jiang et al., 2020)), walnut (*Juglans regia* (Marrano et al., 2020)), Chinese tupelo (*Nyssa sinensis* (Yang et al., 2019)), fragrant rosewood (*Dalbergia odorifera* (Hong et al., 2020)), wheel tree (*Trochodendron aralioides* (Strijk at. Al., 2019)), azalea (*Rhododendron simsii* (Yang et al., 2020)), agrarwood tree (*Aquilaria sinensis* (Nong et al., 2020)), and tea olive (*Osmanthus fragrans* (Yang et al., 2018)). However, such resources are currently lacking for species of the *Fagaceae*, which includes the economically and ecologically important genera *Castanea, Fagus*, and *Quercus* (Kermer at al., 2012). For this family, various draft assemblies have been published (Sork et al., 2016; Martínez-García et al., 2016; Plomion et al., 2016), including European Beech (Mishra et al., 2018), but none is so far resolved on a chromosome scale. To achieve this, we have sequenced the genome of the more than 300 year-old beech individual, Bhaga, from the Kellerwald-Edersee National Park (Germany), and compared it to an individual from the Jamy Nature Reserve (Poland), to get first insights into the genome architecture and variability of *Fagus sylvatica*.

## 2. Materials and Methods

### 2.1. Sampling and processing

#### 2.1.1. Reference genome

The more than 300 year-old beech individual Bhaga (Fig. 1) lives on a rocky outcrop on the edge of a cliff in the Kellerwald-Edersee National Park in Hesse, Germany (51°10’09”N 8°57’47”E). Dormant buds were previously collected for the extraction of high molecular weight DNA and obtaining the sequence data described in Mishra et al. (2018). The same tree was sampled again in February 2018 for obtaining bud samples for constructing Hi-C libraries. Hi-C libraries construction and sequencing was done by a commercial sequencing provider (BGI, Hong Kong, China). For an initial assessment of genome variability and to obtain its genome sequence, Illumina reads derived from the Polish individual, Jamy, reported in Mishra et al. (Mishra et al., 2021a), were used.

**Fig. 1.**
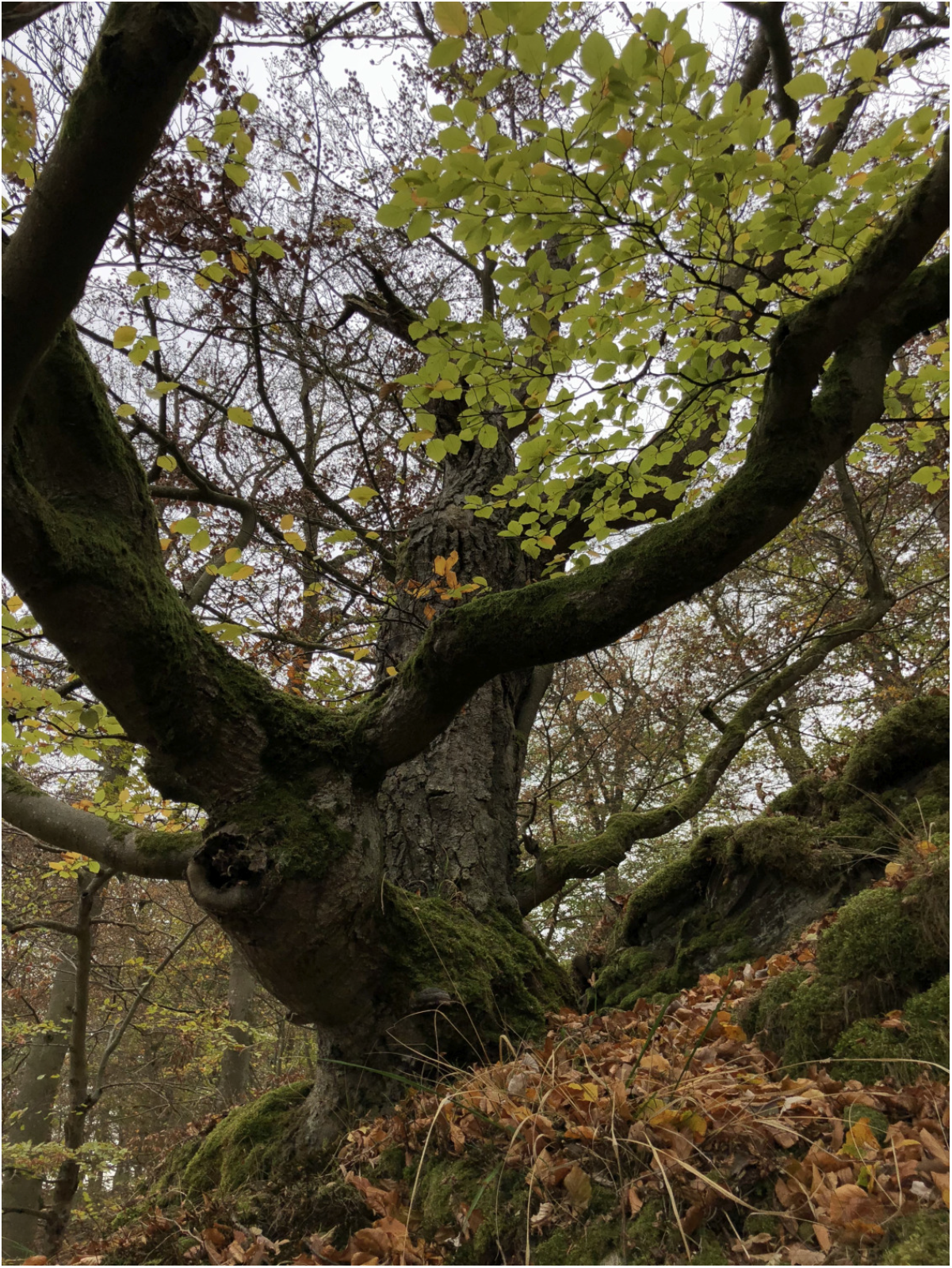
The more than 300 year-old *Fagus sylvatica* reference individual Bhaga on a cliff over the Edersee in the Kellerwald Edersee National Park (Germany)

#### 2.1.2. Progeny trial and linkage map construction

For a progeny trial establishment seeds were sampled from a single mother tree (accession MSSB). About 1,000 beechnuts were collected during two successive campaigns in the fall 2013 and 2016 using a net under the mother tree located in the southern range of the species in the south-west of France (Saint- Symphorien 44° 25’ 41.138’’ N 0° 29’ 23.125” W). Seeds were germinated and raised the following springs at the National Forest Office nursery in Guémené-Penfao (47° 37’ 59.99’’ N -1° 49’ 59.99” W) and then planted at the Nouzilly (47° 32′ 36″ N 0° 45′ 0″ E) experimental unit PAO of INRAE in February 2017 (537 saplings corresponding to the 1^st^ campaign, used for the paternity reconstruction) and at the National Forest Office nursery in Guémené-Penfao in January 2019 (429 saplings corresponding to the 2^nd^ campaign, used for linkage mapping). for relatedness assessment among the half-sib progeny of MSSB, young leaves after bud burst were sampled from saplings in the nursery in spring 2014 (1^st^ campaign) and 2017 (2^nd^ campaign), immediately frozen in dry ice and then stored at -80°C before subsequent genetic analyses. Likewise, leaves were sampled on the mother tree and 19 surrounding adult trees (expected fathers). Nuclear DNA was extracted individually from 10 mg of tissue using the DNeasy Plant Mini Kit (QIAGEN, DE) following the manufacturer’s instructions. DNA concentration was measured on a ND-8000 NanoDrop spectrophotometer (Thermo Scientific, Wilmington, USA). For additional transcriptome construction a total of six different organs were sampled on the MSSB accession, including: two types of buds (quiescent buds and swelling buds just before bud break) during dormancy release the 15th of March 2017, male flowers and female flowers collected the 3rd of May 2017, leaves and xylem collected the 28th of June 2017. Each organ was immediately flash-frozen in liquid nitrogen and stored at -80°C before RNA extraction. For short read sequencing (Illumina), total RNA was extracted from these six samples following the procedure described in Le Provost et al. (2007). Residual genomic DNA was removed before purification using DNase RQ1 (Promega, Madisson, WI, USA) according to the manufacturer’s instructions. The quantity and the quality of each extract was determined using an Agilent 2100 Bioanalyser (Agilent Technologies, Inc., Santa Clara, CA, USA). For long read sequencing (Oxford Nanopore Technologies) total RNA was extracted as described above and depleted using the Ribo-Zero rRNA Removal Kit Plant Leaves (Illumina, San Diego, CA, USA). RNA was then purified and concentrated on a RNA Clean Concentrator^™^-5 column (Zymo Research, Irvine, CA, USA).

For the linkage mapping, vegetative buds from the individuals from the first and second campaign were sampled on the 28^th^ of February 2018 in Nouzilly at the ecodormancy stage from 200 genotypes (i.e. 200 half-sibs that constitute the mapping population) and were frozen on dry ice and then stored at -80°C. RNA was extracted from bud scale-free leaves following the procedure described above. These 200 genotypes included two relatively large full-sib families comprising 49 full-sibs (family MSSBxSSP12) and 36 full-sibs (family MSSBxMSSH) (see results section).

### 2.2. Chromosomal pseudo-molecules and their annotation

#### 2.2.1 Building of chromosomal pseudo-molecules using Hi-C reads

The previous scaffold-level assembly was constructed with Illumina shotgun short reads and PacBio long reads (Mishra et al., 2018). For a chromosome-level assembly, intermediate results from the previous assembly were used as the starting material. Sequence homology of the 6699 scaffolds generated from the DBG2OLC hybrid assembler (Ye et al., 2016), to the separately assembled chloroplast and mitochondria of beech, were inferred using blast v2.10.1 (Altschul et al., 1990). All scaffolds that match in full length to any of the organelle with identity > 99 % and gaps and/or mismatches ≤ 3 were discarded. The remaining 6657 scaffolds along with Hi-C data (116 Mb) were used in ALLHiC (Zhang et al., 2019) for building the initial chromosome-level assembly. The cleaned Illumina reads were aligned to the initial assembly using Bowtie2 software (Langmead and Salzberg, 2012) and then, sorted and indexed bam files of the concordantly aligned read pairs for all the sequences were used in Pilon (Walker et al., 2014) to improve the correctness of the assembly. The final assemblies for Bhaga and Jamy were deposited under the accession numbers PRJEB43845 and PRJNA450822, respectively.

The completeness of the assembly was evaluated with plant-specific (viridiplantae_odb10.2019-11- 20) and eudicot-specific (eudicots_odb10.2019-11-20) Benchmarking Universal Single-Copy Orthologs (BUSCO v4.1.4) (Seppey et al., 2019).

#### 2.2.2. Gene prediction

Cleaned transcriptomic Illumina reads (minimum read length: 70; average read quality: 25 and read pairs containing no N) were aligned to the assembly using Hisat (Kim et al., 2015) in order to generate splice-aware alignments. The sorted and indexed bam file (samtools, v1.9 (Li et al., 2009)) of the splice alignments was used in “Eukaryotic gene finding” pipeline of OmicsBox (Accessed March 3, 2020) which uses Augustus (Stanke and Morgenstern, 2005) for gene prediction. For prediction, few parameters were changed from the default values. Minimum intron length was set to 20 and minimum exon length was set to 200 and complete genes (with start and stop codon) of a minimum of 180 bp length were predicted, by choosing *Arabidopsis thaliana* as the closest organism.

#### 2.2.3. Assessment of the gene space

The protein sequences of the PLAZA genes for *A. thaliana, Vitis vinifera*, and *Eucalyptus grandis* were downloaded from plaza v4.5 dicots (Accessed October 21, 2020) dataset and were used along with the predicted proteins from our assembly to make protein clusters using cd-hit v.4.8.1 (Li and Godzik, 2006; Fu et al., 2012). The number of exons per genes was assessed and compared to the complete coding genes from *A. thaliana, Populus trichocarpa*, and *Castanea mollissima*, in line with the comparison made in the scaffold level assembly (Mishra et al., 2018).

#### 2.2.4. Functional annotation of genes

The predicted genes were translated into proteins using transeq (EMBOSS:6.6.0.0 (Rice et al., 2000)) and were queried against the non-redundant database from NCBI (downloaded on 2020-06-24) using diamond (v0.9.30) software (Buchfink and Xie, 2015) to find homology of the predicted proteins to sequences of known functions. For prediction of protein family membership and the presence of functional domains and sites in the predicted proteins, Interproscan v5.39.77 (Jones et al., 2014) was used. Result files from both diamond and Interproscan (in Xml format) were used in the blast2go (Götz et al., 2008) module of OmicsBox and taking both homology and functional domains into consideration, the final functional annotations were assigned to the genes. The density of coding space for each 100 kb region stretch was calculated for all the chromosomes.

#### 2.2.5. Repeat prediction and analysis

A repeat element database was generated using RepeatScout (v1.0.5) (Price et al., 2005), which was used in RepeatMasker (v4.0.5) (Smit and Hubley, 2007) to predict repeat elements. The predicted repeat elements were further filtered on the basis of their copy numbers. Those repeats represented with at least 10 copies in the genome were retained as the final set of repeat elements of the genome. Repeat fractions per 100 kb region for each of the chromosomes were calculated for accessing patterns of repeat distribution over the genome.

In a separate analysis, repeat elements present in *Fagus sylvatica* were identified by a combination of homology-based and de novo approaches using RepeatModeler 2.0 (Flynn et al., 2020) and RepeatMasker v. 4.1.1 (Tarailo-Graovac and Chen, 2009). First, we identified and classified repetitive elements de novo and generated a library of consensus sequences using RepeatModeler 2.0 (Flynn et al., 2020). We then annotated repeats in the assembly with RepeatMasker 4.1.1 (Tarailo-Graovac and Chen, 2009) using the custom repeat library generated in the previous step.

#### 2.2.6. Telomeric and Centromeric repeat identification

Tandem repeat finder (TRF version 4.0.9) (Benson, 1999) was used with parameters 2, 7, 7, 80, 10, 50 and 500 for Match, Mismatch, Delta, PM, PI, Minscore and MaxPeriod, respectively (Marrano et al., 2020), and all tandem repeats with monomer length up to 500 bp were predicted. Repeat frequencies of all the monomers were plotted against the length of the monomers to identify all high-frequency repeats. As the repeats were fetched by TRF program with different start and end positions and the identical repeats were falsely identified as different ones, the program MARS (Ayad and Pissis, 2017) was used to align the monomers of the different predicted repeats, and the repeat frequencies were adjusted accordingly. The chromosomal locations of telomeric and centromeric repeats were identified by blasting the repeats to the chromosomes. For confirmation of centromeric locations, pericentromeres of *A. thaliana* were blasted against the chromosomes of Bhaga.

#### 2.2.7. Organelle integration

Separately assembled chloroplast (Mishra et al. 2021a) and mitochondrial (Mishra et al. 2021b) genomes were aligned to the genomic assembly using blastn with an e-value cut-off of 10e-10. Information for different match lengths and different identity cut-offs were tabulated and analysed. Locations of integration into the nuclear genome were inferred at different length cut-offs for sequence homology (identity) equal to or more than 95%. The number of insertions per non-overlapping window of 100 kb was calculated separately for both organelles.

#### 2.2.8. SNP identification and assessment

The DNA isolated from the Polish individual Jamy was shipped to Macrogen Inc. (Seoul, Rep. of Korea) for library preparation with 350 bp targeted insert size using TruSeq DNA PCR Free preparation kit (Illumina, USA) and sequencing on HiSeq X device (Illumina, USA) using PE-150 mode. The generated 366,127,860 raw read pairs (55.3 Gb) were processed with AfterQC v 0.9.1 (Chen et al., 2017) for quality control, filtering, trimming and error removal with default parameters resulting in 54.12 Gbp of high-quality data. Illumina shotgun genomic data from Jamy was mapped to the chromosome-level assembly using stringent parameters (--very-sensitive mode of mapping) in bowtie2 (Li, 2011). The sam formatted output of Bowtie2 was converted to binary format and sorted according to the coordinates using samtools version 1.9 (Li et al., 2009). SNPs were called from the sorted mapped data using bcftools (version: 1.10.2) (Li, 2011) call function. SNPs were called for only those genomic locations with sequencing depth ≥ 10 bases. All locations 3 bp upstream and downstream of gaps were excluded. For determining heterozygous and homozygous states in Bhaga, sites with more than one base called and a ratio between the alternate and the reference allele of ≥ 0.25 and < 0.75 in were considered as heterozygous SNP. Where the ratio was ≥ 0.75, the position was considered homozygous. In addition, homozygous SNPs were called by comparison to Jamy, where the consensus base in Jamy was different than in Bhaga and Bhaga was homozygous at that position. SNP density was calculated for each chromosome in 100 kb intervals.

#### 2.2.9. Genome browser

A genome browser was set up using JBrowse v.1.16.10 (Buels et al., 2016). Tracks for the predicted gene model, annotated repeat elements were added using the gff files. Separate tracks for the SNP locations and the locations of telomere and centromere were added as bed files. A track depicting the GC content was also added. The genome browser can be accessed from http://beechgenome.net.

### 2.3. Pedigree reconstruction

#### 2.3.1. SNP assay design and genotyping for relatedness assessment among half-sibs

We used a multiplexed assay using the MassARRAY® MALDI-TOF platform (iPLEX MassArray, Agena BioScience, USA) to genotype the mother tree (MSSB), its half-sib progeny from the 1^st^ campaign and 19 putative fathers. PCR and extension primers were designed from flanking sequences (60pb of either side) of 40 loci (Supplementary file 5) available from Lalagüe et al. (2014) and Ouayjan and Hampe (2018). Data analysis was performed with Typer Analyzer 4.0.26.75 (Agena BioScience). We filtered out all monomorphic SNPs, as well as loci with a weak or ambiguous signal (i.e., displaying more than three clusters of genotypes or unclear cluster delimitation). Thirty-six SNPs were finally retained for the paternity analysis.

#### 2.3.2. Sibship assignment

Paternity analysis was carried out using Cervus 3.0 software (Kalinowski et al., 2007, Marshall et al. 1998) to check the identity of the maternal parent and identify the paternal parent among 19 candidate fathers growing in the neighbourhood of mother tree MSSB. Cervus was run assuming a 0.1% genotyping error rate. The pollen donor of each offspring was assigned by likelihood ratios assuming the strict confidence criterion (95%). We performed simulations with the following parameters: number of offspring genotypes = 100 000, number of candidate fathers = 19, mistyping rate = 0.01 and proportion of loci typed = 0.9755. Zero mismatch was allowed for each offspring and the supposed father. The Cervus selfing option was used because self-pollination may occur.

### 2.4. Unigene set construction

#### 2.4.1. Library construction and sequencing

Six Illumina RNA-Seq libraries (one for each organ) were constructed from 500ng total RNA using the TruSeq Stranded mRNA kit (Illumina, San Diego, CA, USA), which allows for mRNA strand orientation (the orientation of sequences relative to the antisense strand is recorded). Each library was sequenced using 151 bp paired end reads chemistry on a HS4000 Illumina sequencer.

One Nanopore cDNA library was also prepared from entire female flowers RNA. The cDNA library was obtained from 50 ng RNA according to the Oxford Nanopore Technologies (Oxford Nanopore Technologies Ltd, Oxford, UK) protocol “cDNA-PCR Sequencing (SQK-PCS108)” with a 14 cycles PCR (6 minutes for elongation time). ONT adapters were ligated to 190 ng of cDNA. The Nanopore library was sequenced using a MinION Mk1b with R9.4.1 flowcells.

#### 2.4.2. Bioinformatic analysis

Short-read RNA-Seq data (Illumina) from the six tissues were assembled using Velvet (Zerbino et al., 2010) 1.2.07 and Oases (Schulz et al., 2012) 0.2.08, using a k-mer size of 63 bp. Reads were mapped back to the contigs with BWA-mem (Li et al., 2009) and the consistent paired-end reads were selected. Chimeric contigs were identified and splitted (uncovered regions) based on coverage information from consistent paired-end reads. Moreover, open reading frames (ORF) and domains were searched using respectively TransDecoder (Haas et al., 2013) and CDDsearch (Marchler-Bauer et al., 2011). We only allowed breaks outside ORF and domains. Finally, the read strand information was used to correctly orient the RNA-seq contigs.

Long-read RNA-Seq data (Oxford Nanopore Technologies) from female flowers were corrected using NaS (Madoui et al., 2015) with default parameters.

Contigs obtained from short reads as well as corrected long reads were then aligned on a draft version of MSSB genome assembly (unpublished) using BLAT (Kent, 2002). The best matches (based on BLAT score) for each contig were selected. Then, Est2genome (Mott, 1997) was used to refine the alignments and we kept alignments with an identity percent and a coverage at least of 95% and 80%, respectively. Finally, for each genomic cluster, the sequence with the best match against *Quercus robur* or *Castanea mollissima* proteins was kept. This procedure yielded 34,987 unigenes (below referred to as the 35K unigene set).

### 2.5. Genotyping-by-sequencing of the mapping population

#### 2.5.1. RNAseq libraries construction

The 200 RNA samples were prepared as described above (Unigene set construction section), using the TruSeq Stranded mRNA kit (Illumina, San Diego, CA, USA), from 500 ng total RNA. Libraries were multiplexed onto Illumina Novaseq 6000 using S4 chemistry (2×150 read length), targeting approximately 30 million reads per sample.

#### 2.5.2. RNAseq reads processing for the MSSB accession

We first identified SNPs in the MSSB reference unigene. To this end, a trimming procedure was applied to the MSSB sequences to remove adapters, primers, ribosomal reads and nucleotides with quality value lower than 20 from both ends of the reads and reads shorter than 30 nucleotides as described previously (Alberti et al., 2017). Trimmed reads were aligned onto the 35K unigene set using bwa mem 0.7.17. Biallelic SNPs were identified using two methods: samtools 1.8 / bcftools 1.9 (Danecek et al. 2021) and GATK 3.8 (van der Auwera et al. 2020) with java 1.8.0_72. We kept SNPs identified by both methods.

#### 2.5.3. Identification of SNPs from RNAseq data and offspring genotype inference

We called SNPs and bioinformatically genotyped the mapping population at each MSSB polymorphic site, based on the paired-end Illumina sequencing of 200 RNAseq libraries. The 200 raw-read datasets were trimmed following the same procedure used for MSSB. Reads were aligned to the 35K unigene set using bwa mem 0.7.17. Genotypes were recovered from the 200 libraries at the 507,905 polymorphic positions, identified in MSSB, using GATK 3.8.

We then applied the following four-step filtering procedure: i/ for each SNP of a given half-sib, polymorphic genotypes were set to monomorphic if the sequencing depth for this individual at this position was lower than 20X; ii/ we kept SNPs only if at least 50% of the mapping population (i.e. 100 half-sibs) were heterozygous at this site; iii/ we kept only polymorphic sites consistent with a 1:1 heterozygote:homozygote genotype ratio, according to a Chi-square test with a 90% confidence interval (Chi-square < 6.635, 1 d.f.), corresponding to heterozygous loci in the mother tree and monomorph in all possible fathers; iv/ finally, for each contig, we retained only the SNP with fewest missing data in the mapping population.

### 2.6. Linkage map construction

Half-sibs presenting too many missing data were discarded. As a result, 182 individuals (out of 200 selected from the first and second campaign) with valid genotypes for at least 4,127 loci were kept for further analyses. A preliminary analysis was then performed using R-qtl package to group linked SNP markers into robust linkage groups (LG) (LOD = 8) (Supplementary file 6). Given the large number of markers per LG, marker ordering was performed within each LG using JoinMap 4.1 (Kyazma, Wageningen, NL). To this end, linkage groups of the maternal parent (MSSB) were constructed using a four-step procedure: i) The maximum likelihood (ML) algorithm of JoinMap was first used with a minimum linkage LOD score of 5 to calculate the number of crossing-overs (CO) for each individual and to estimate the position of all mapped SNPs, ii/ then, the regression algorithm (with a minimum LOD of 5 and default parameters: recombination frequency of 0.4 and maximum threshold value of 1 for the jump) was used for a subset of evenly spaced SNPs (referred to below as set #1 SNPs) along each LG, iii) the maternal linkage maps of the two full-sib families, identified from the paternity test, were constructed using this subset of markers and individuals, providing two genetic maps (referred to below as set #2 and set #3 SNPs) with higher confidence in genetic distance estimates and marker ordering, both parents being known; iv) finally, from these two SNP datasets, we created a final dataset (set #4) combining sets #2 and #3. For these 3 marker sets (#2, #3 and #4), a first map was constructed using the ML algorithm to calculate the number of CO and a second map was established using the regression algorithm excluding SNPs with high conflict of positions and reducing the number of CO.

### 2.7. Genomic scaffold anchoring

Sequences of the unigenes encompassing SNP markers included in the linkage map, were aligned on the genome assembly using BLAT with default parameters, except “-minScore=80”. Unigenes presenting more than one alignment were filtered out. In other words, when a second best match having a score equal to or greater than 90% of the best score the marker was tagged as ambiguous. For all the remaining alignments we kept only the alignment with the best score.

## 3. Results

### 3.1. General genome features

#### 3.1.1. Genomic composition and completeness

The final assembly of the Bhaga genome was based on hybrid assembly of PacBio and Illumina reads as well as scaffolding using a Hi-C library. It was resolved into 12 chromosomes, spanning 535.4 Mb of the genome and 155 unassigned contigs of 4.9 Mb, which to 79% consisted of unplaced repeat regions that precluded their unequivocal placement. It revealed a high level of BUSCO gene detection (97.4%), surpassing that of the previous assembly and other genome assemblies available for members of the *Fagaceae* (Table 1). Of the complete assembly, 57.12% were annotated as interspersed repeat regions and 1.97% consisted of simple sequence repeats (see Supplementary File 1 for details regarding the repeat types and abundances).

**Table 1.** Comparison of BUSCO completeness in Fagaceae genomes available and in the present study (*Fagus sylvatica* V2).

| Species | Complete<br>genes | Single<br>genes | Duplicated<br>genes | Fragmented<br>genes | Missing<br>genes |
| --- | --- | --- | --- | --- | --- |
| *Fagus sylvatica* V2 | 97.4% | 90.3% | 7.1% | 1.3% | 1.3% |
| *Fagus sylvatica* V1 (Mishra et al., 2018) | 96.6% | 85.6% | 11% | 1.8% | 1.6% |
| *Castanea mollissima* (Wang et al., 2020) | 92.4% | 88.8% | 3.7% | 1.5% | 6.1% |
| *Quercus lobata* v3 (Sork et al., 2016) | 93.5% | 87.6% | 5.9% | 1.0% | 5.5% |

The gene prediction pipeline yielded 63,736 complete genes with start and stop codons and a minimum length of 180 bp. Out of these, 2,472 genes had alternate splice variants. For 86.8% of all genes, a functional annotation could be assigned. Gene density varied widely in the genome, ranging from zero per 100 kb window to 49.7%, with an average and median of 18.2% and 17.6%, respectively. Gene lengths ranged from 180 to 54,183 bp, with an average and median gene length of 3,919 and 3,082 bp, respectively. In *Fagus sylvatica* 4.9 exons per gene were found on average, corresponding well to other high-quality plant genome drafts. The distribution of exons and introns in comparison to *J. regia* and *A. thaliana* are presented in Table 2. An analysis of PLAZA genes identified 28,326 such genes in *F. sylvatica*, out of which 1,776 genes were present in three other species used for comparison (Supplementary File 2).

**Table 2.**
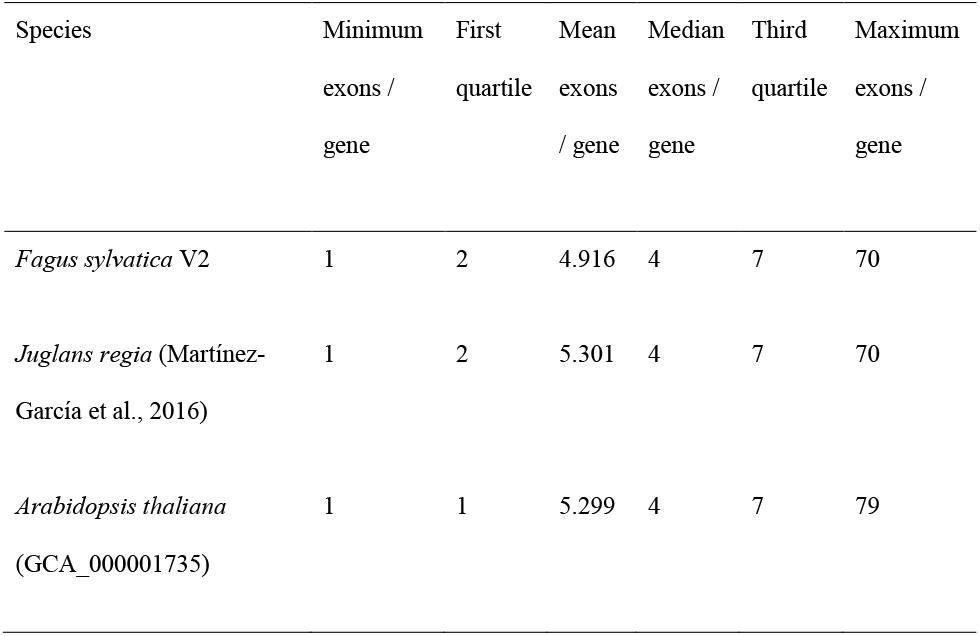
Distribution of exons in *Fagus sylvatica* in comparison to *Juglans regia* and *Arabidopsis thaliana*.

#### 3.1.2. Telomere and centromere predictions

The results given above indicate a high quality of the genome assembly and the gene annotations. To ascertain that the chromosomes were fully resolved, telomeric and centromeric regions were predicted in the genome. The tandem repeat element TTTAGGG was the most abundant repeat in the genome and was the building block of the telomeric repeats. Out of 12 chromosomes, 8 have stretches of telomeric repeats towards both ends of the chromosomes and the other 4 chromosomes have telomeric repeats towards only one end of chromosomes (Fig. 2). One unplaced scaffold of 110,653 bp which is composed of 12,051 bp of telomeric repeats at one end, probably represents one of the missing chromosome-ends.

**Fig. 2.**
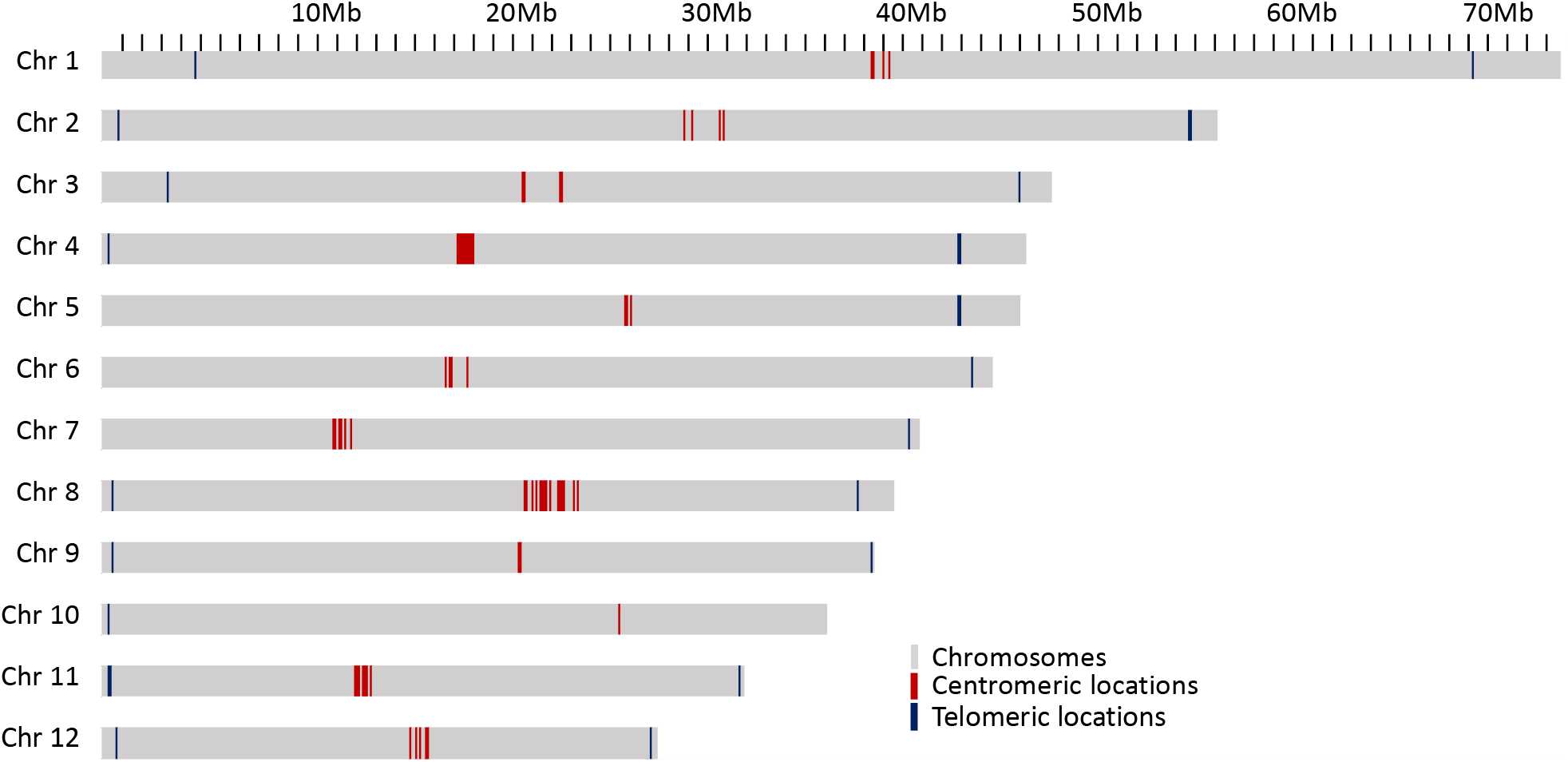
Locations of probable centromeric repeats on the chromosomes presented as red lines and telomeric locations as blue line on the chromosomes.

Two different types of potential centromeric repeats were observed, consisting of 79 bp and 80 bp monomer units (Supplementary File 3). Centromeric repeats were also observed in higher numbers outside the main centromeric region on several chromosomes (Supplementary File 3). However, except for chromosome 10, there was a clear clustering of centromeric repeats within each of the chromosomes, likely corresponding to the actual centromere of the respective chromosomes, and supported also by complementary evidence, such as similarities to centromeric regions of *A. thaliana*, high gypsy element content and low GC content (Supplementary File 3).

#### 3.1.3. Integration of organelle DNA in the nuclear genome

As it has previously been shown that organelle DNA insertions can be uneven across the genome and associated with chromatin structure (Wang and Timmis 2013), their distribution in the genome of Bhaga was analysed. For both chloroplast (Mishra et al. 2021a) and mitochondria (Mishra et al. 2021b), multiple integrations of fragments of variable length of their genomic DNA were observed in all chromosomes (Figs. 3, 4). These fragments varied in length from the minimum size threshold (100 bp) to 54,784 bp for the chloroplast and 26,510 bp for the mitochondrial DNA. The identity of the integrated organelle DNA with the corresponding stretches in the organelle genome ranged from the minimum threshold tested of 95% to 100%. Nuclear-integrated fragments of organelle DNA exceeding 10 kbp were found on six chromosomes for the chloroplast, but only on one chromosome for the mitochondrial genome (Figs. 3, 4).

**Fig. 3.**
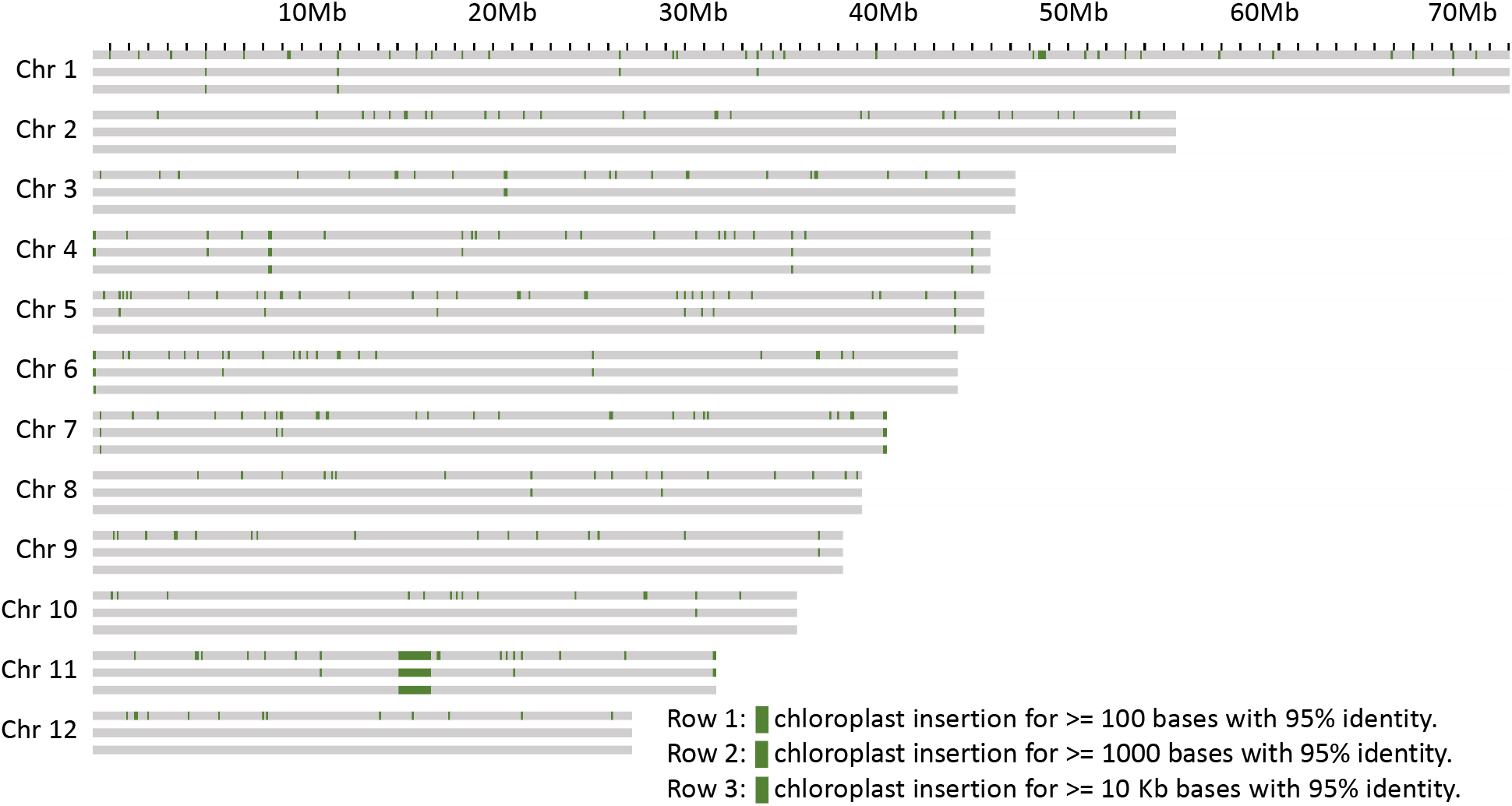
Chloroplast genome insertions within 100 kb windows on the chromosomes. Each chromosome is represented as three rows, the first with insertions more than 100 bp long, the second row with more than 1 kb and the third with more than 10 kb.

**Fig. 4.**
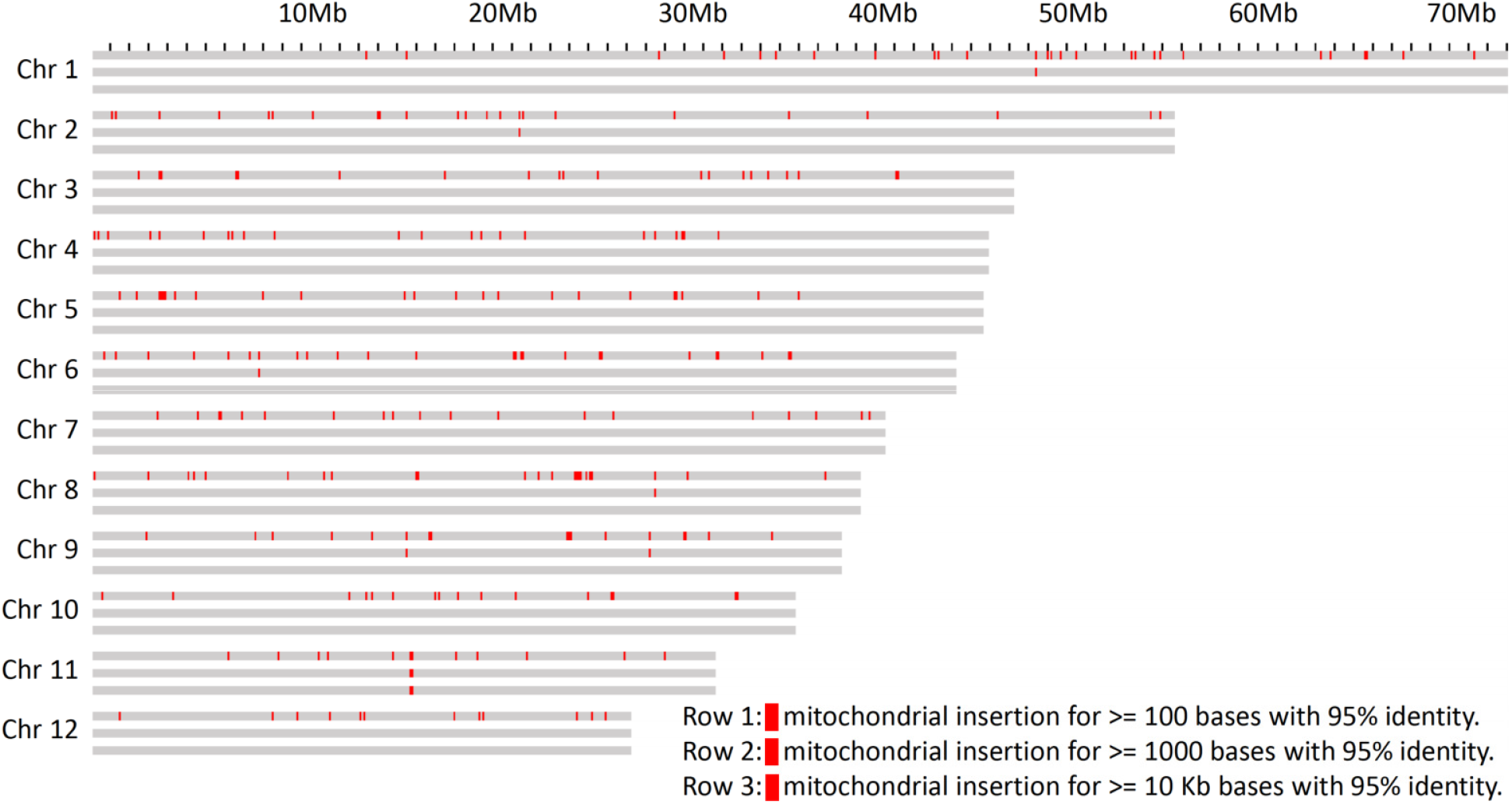
Mitochondrion genome insertions within 100 kb windows on the chromosomes. Each chromosome is represented as three rows, the first with insertions more than 100 bp long, the second row with more than 1 kb and the third with more than 10 kb.

Nuclear insertions with sequence identity > 99% were about ten times more frequent for chloroplast than for mitochondrial DNA with 173 vs. 16 for fragments > 1 kb and 115 vs. 11 for fragments > 5 kb, respectively. Eight of these matches of mitochondria were located on unplaced contigs. Overall, mitochondrial insertions tended to be smaller and show a slightly higher sequence similarity (Supplementary File 4), suggesting that they might be purged from the nuclear genome quicker than the chloroplast genome insertions.

The integration of organelle DNA into the nuclear genome was mostly even, but tandem-like integrations of chloroplast DNA on chromosome 2 were observed (Fig. 3). In addition, insertions of both organelles were found close to the ends in 4 of the 24 chromosome ends (4, 6, 7, and 8). For the insertions further than 500 kb away from the chromosome ends the integration sites of mitochondrion DNA were sometimes found within the same 100 kb windows where the chloroplast DNA insertion was found. If some regions of the genome are more amenable for the integration of organelle DNA than others needs to be clarified in future studies. A major anomaly was found on chromosome 11, where in a stretch of about 2 Mb (from about Mb 16-18 on that chromosome) consisting mainly of multiple insertions of both chloroplast and mitochondrial DNA was observed. In this region, an insertion of more than 20 kb of mitochondrial DNA was flanked by multiple very long integrations of parts of the chloroplast genome on both sides (Figs. 3, 4). Thus, these integrations appeared almost repeat-like at this particular location.

#### 3.1.4. Repeat elements and gene space

The most abundant repeat elements were LTR elements and LINEs, covering 11.49% and 3.66% of the genome, respectively. A detailed list of the element types found, their abundance and proportional coverage of the genome is given in Supplementary File 1. Repeat elements presence was variable across the chromosomes (Fig. 5). While the repeat content per 100 kb window exceeded 50 % over more than 88% of chromosome 1, this was the case for only 37.5% of chromosome 9. Chromosomes showed an accumulation of repeat elements towards their ends, except for chromosome 10, where only a moderate increase was observed on one of the ends, and chromosome 1, where repeat elements were more evenly distributed. Repeat content was unevenly distributed, with a patchy distribution of repeat-rich and repeat-poor regions of variable length.

**Fig. 5.**
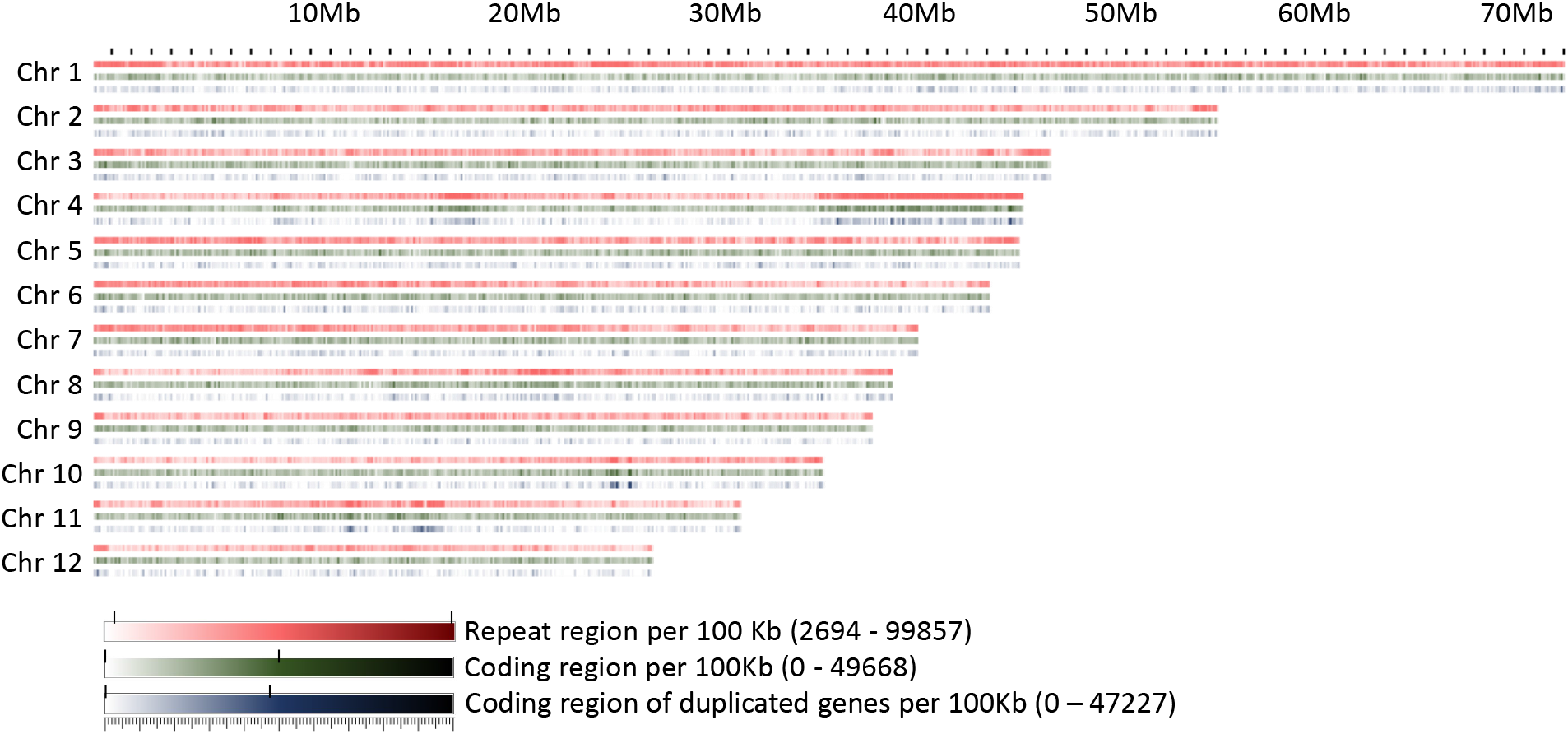
Repeat regions, coding regions, and regions coding for genes present within 100 kb windows on the chromosomes.

A conspicuous anomaly was noticed in chromosome 4, where at one end a large region of about 10 Mb was found in which 97% of the 100 kb windows had a repeat content greater than 70%. This region also contained a high proportion of duplicated or multiplicated genes (Fig. 5). Additional regions containing more than 20% of duplicated genes within a window of at least 1 Mb were identified on chromosomes 4, 10, and 11. On chromosome 11, two clusters were detected, one of which corresponded to the site of organelle DNA insertions described above.

The ribosomal cistrons were reported to be located at the telomeres of four different chromosomes in *F. sylvatica* (Ribeiro et al., 2011). Due to the highly repetitive nature of the ribosomal repeats and their placement near the telomeres, they could not be assigned with certainty to specific chromosomes and thus remained in four unplaced contigs. However, the 5S unit, which is separate from the other ribosomal units in F. sylvatica, could be placed near the centromeric locations of chromosomes 1 and 2, in line with the locations inferred by fluorescence microscopy (Ribeiro et al., 2011).

Coding space was more evenly distributed over the chromosomes, with the exception of the regions with high levels of duplicated or multiplied genes. Apart from this, a randomly fluctuating proportion of coding space was observed, with only few regions that seemed to be slightly enriched or depleted in terms of coding space, e.g. in the central part of chromosome 8.

#### 3.1.5. Distribution of single nucleotide polymorphisms

To study, if the distribution of single nucleotide polymorphisms (SNPs) correlates with the feature reported above, they were identified on the basis of the comparison of the two individuals investigated in this study, Bhaga and Jamy. A total of 2,787,807 SNPs were identified out of which 1,271,410 SNPs were homozygous (i.e. an alternating base on both chromosomes between Bhaga and Jamy) and 1,582,804 were heterozygous (representing two alleles within Bhaga). A total of 269,756 SNPs fell inside coding regions out of which 119,946 were homozygous.

Heterozygous SNPs were very unequally distributed over the chromosomes (Fig. 6). Several regions, the longest of which comprised more than 30 Mb on chromosome 6, contained only very low amounts of heterozygous SNPs. Apart from the chromosome ends, where generally few heterozygous positions were observed, all chromosomes contained at least one window of 1 Mb where only very few heterozygous SNPs were present. On chromosomes 2, 3, 4, 6, and 9 such areas extended beyond 5 Mb. On chromosome 4 this region corresponded to the repeat region anomaly reported in the previous paragraph, but for the region poor in heterozygous SNPs on chromosome 9, no association with a repeat-rich region could be observed.

**Fig. 6.**
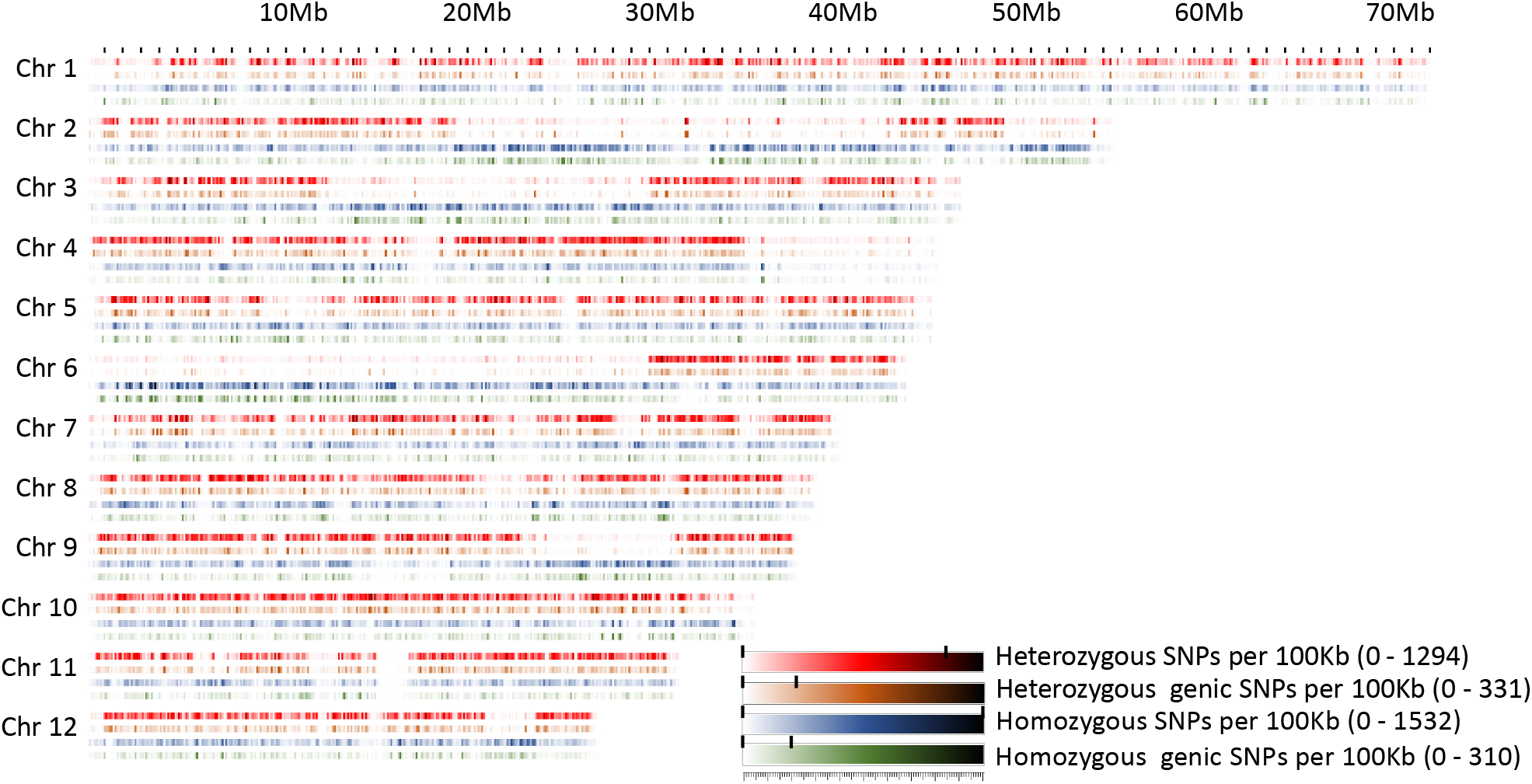
Homozygous and heterozygous SNPs in *Fagus sylvatica* present within 100 kb windows on the chromosomes.

Homozygous SNPs differentiating Bhaga and Jamy, often followed a different pattern. All regions with low heterozygous SNP frequency longer than 5 Mb had an above-average homozygous SNP frequency, with the exception of the anomalous repeat-rich region on chromosome 4, which had very low frequencies for both homozygous and heterozygous SNPs. However, there were also two regions of more than 1 Mb length on chromosome 11 that also showed low frequencies of both SNP categories (Fig. 6).

Generally, the frequency of overall and intergenic SNPs per 100 kb window corresponded well for both heterozygous and homozygous SNPs, suggesting neutral evolution. However, there were some regions in which genic and intergenic SNP frequencies were uncoupled. For example, on chromosome 1 a high overall heterozygous SNP frequency was observed at 37.7, 48.2 and 56 Mb, but genic heterozygous SNP frequency was low despite normal gene density, suggesting the presence of highly conserved genes. In line with this, also the frequency of homozygous genic SNPs was equally low in the corresponding areas. Similary, homozygous SNP frequencies were also decoupled on chromosome 1, where a low frequency was observed at 4.2, 7.1, 38.2, 62.1, and 64.8 Mb, but a high genic SNP frequency was observed. This suggests the presence of diversifying genes in the corresponding 100 kb windows, such as genes involved in coping with biotic or abiotic stress.

In line with the different distribution over the chromosomes, with large areas poor in heterozygous SNPs, there were much more windows with low numbers of heterozygous SNPs than windows with homozygous SNPs (Fig. 7). Notably, at intermediate SNP frequencies, homozygous SNPs were found in more 100 kb windows, while at very high SNP frequencies, heterozygous SNPs were more commonly found. This pattern is consistent with predominant local pollination, but occasional introgression of highly distinct genotypes.

**Fig. 7.**
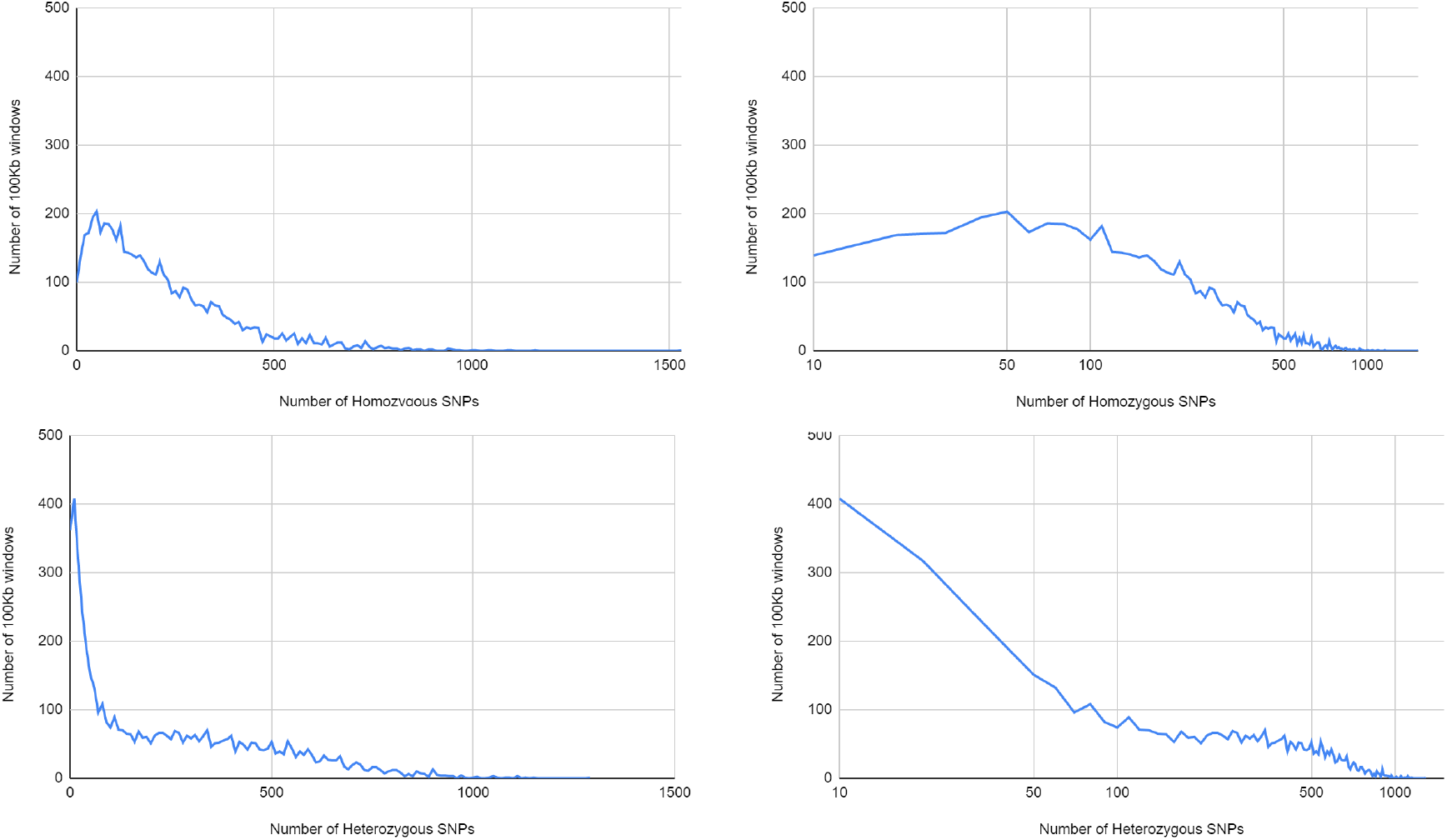
Distribution of homozygous and heterozygous SNPS in non-overlapping 100 kb windows.

#### 3.1.6. Genome browser

A genome browser for the genome of Bhaga, with the various genomic features outlined above annotated, is available at beechgenome.net. Predicted genes, annotated repeat elements and homozygous and heterozygous SNPs are available in “B. Annotations”. The telomeric and centromeric locations, as well as the GC content details are available in “C. Other Details”.

### 3.2. Validation of chromosomal-scale pseudomolecules

#### 3.2.1. Pedigree reconstruction

The analysis of the 36 SNPs using Cervus allowed the identification of candidate fathers and reconstruct full-sib families. For 317 of the 537 offspring a likely father was identified. The 19 candidate fathers were represented in the progeny, although their contributions were variable (0.8 to 21%). For the other offpring, no father could be assigned, i.e. the pollen donor is not present among the surrounding trees (corresponding to 210 genotypes, i.e. 39.1% of the samples when 0 mismatch is allowed, and 22 % when 1 mismatch is allowed). The two largest families comprised 68 (MSSBxMSSH) and 86 (MSSBxSSP12) full-sibs. Few years after plantations, 36 genotypes for the former and 49 for the latter survived (Table 3).

**Table 3.** Size of the full-sib families identified from pedigree reconstruction.

| candidate<br>father | size of the<br>full-sib<br>family |
| --- | --- |
| MSSB | 47 |
| MSSH | 68 |
| SSP01 | 24 |
| SSP02 | 27 |
| SSP03 | 4 |
| SSP04 | 10 |
| SSP05 | 16 |
| SSP06 | 13 |
| SSP07 | 9 |
| SSP08 | 17 |
| SSP09 | 12 |
| SSP10 | 9 |
| SSP11 | 17 |
| SSP12 | 86 |
| SSP13 | 15 |
| SSP14 | 10 |
| SSP15 | 2 |
| SSP16 | 13 |
| SSP17 | 3 |
| SSP18 | 8 |
| sum | 410 |

#### 3.2.2. A new unigene set for European beech

Our study provides a new reference unigene set for *Fagus sylvatica* based on short and long NGS reads obtained from cDNA libraries constructed from six different tissues. The first unigene set for this species was established back in 2015 using a combination of Sanger and Roche-454 reads (Lesur et al. 2015). The sequences were assembled into 21000 contigs. A second step was achieved by Müller et al. (2017) using NGS data (Illumina) resulting in 44000 contigs. Tis third transcript catalog contains a total of 34,987 items. When compared to the oak proteome (to date the best annotated among Fagaceae species), this new reference provides the most complete transcript catalog (Table 4).

**Table 4.** Summary statistics for three Fagus sylvatica unigene sets. The last column gives the number of homologous proteins (blastX E10-5) against the most complete fagaceae proteome (25,808 proteins) to date, that of Quercus robur (Plomion et al. 2018).

|  | Technologies | assembler | # contigs in the unigene | Identified oak proteins | # contigs with identified proteins |
| --- | --- | --- | --- | --- | --- |
| Lesur et al. 2015 | Sanger | MIRA | 21,057 | 22,684 | 16,512 |
|  | 454 Roche |  |  |  |  |
| Muller et al. 2017 | Illumina | CLCBio | 44,335 | 24,804 | 24,480 |
| This study* | Illumina | Velvet | 34,987 | 24,826 | 22,347 |
|  | ONT | Oases | 33,013** | 24,811 | 21,886 |
|  |  |  | (≥200bp) |  |  |

\*In addition to Illumina and ONT RNAseq, contigs obtained from Lesur et al. 2015 were also included in the analysis. This first unigene provided a total of 609 transcripts to the new reference unigene. \* \*\*Transcripts longer than 200bp are available online (ENA accession HBVZ01000000). Smaller contigs are available upon request.

#### 3.2.3. Identification of RNAseq-based SNP markers for linkage mapping

Sequencing of the six tissues (collected on the MSSB accession) using an RNA-Seq approach, led to 408,111,505 Illumina paired-end reads. A total of 383,149,091 trimmed sequences were used to identify putative segregating SNPs in MSSB.

On average, 82.67% of the reads were properly aligned on the reference unigene, ranging from 72.94% for the male flowers to 86.46% for leaves. We identified 613,885 and 507,905 SNPs using Samtools/bcftools and GATK, respectively. A total of 507,905 SNPs in MSSB were finally identified by both methods.

Sequencing of the 200 siblings, followed by trimming of the raw data, led to a total of 9,155,925,565 reads. On average, 78.64% of the reads were properly aligned on the reference unigene (min. 72.6% - max. 83.04%). We found 267,361 polymorphic sites in at least one out of the 200 half-sibs. Our four-step filtering process yielded a final set of 6,385 SNPs spread over 6,385 contigs, with at least 20X coverage.

#### 3.2.4. Linkage map construction

Beech is a diploid species with 2n=2x=24. The 12 expected linkage groups (LG) were retrieved using SNPs from set #1 using the R-qtl package. The number of SNP markers per LG ranged from 231 to 412. However, the detailed linkage analysis, carried out with JoinMap for each LG, revealed an unexpectedly high number of crossing-overs and oversized LGs compared to previous linkage mapping analyses performed in beech (Scalfi et al., 2004) or oak (Bodénès et al., 2016), probably owing to genotyping errors among the 182 hal-sibs. Because of this, we established genetic linkage maps based on the two largest full-sib families identified from the paternity analysis, and only used the corresponding two sets of mapped SNPs (sets #2 and #3) to create a combined genetic linkage map based on the analysis of 182 half-sibs. A total of 768 SNPs were available for the combined maternal linkage map, 368 of which were unambiguously mapped on the 13 longest LGs. The size of LGs varied from 64 to 279 cM and comprised 8 to 56 SNPs (Table 5). High colinearity was observed between the homologous linkage groups obtained from the three different maps (Fig. 8).

**Table 5.**
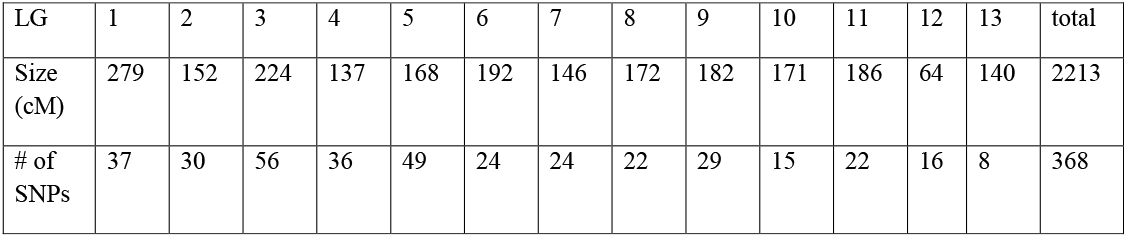
Characteristics of the combined maternal linkage map in terms of genetic size (cM) and number of SNP markers for each linkage group (LG).

| LG | 1 | 2 | 3 | 4 | 5 | 6 | 7 | 8 | 9 | 10 | 11 | 12 | 13 | total |
| --- | --- | --- | --- | --- | --- | --- | --- | --- | --- | --- | --- | --- | --- | --- |
| Size (cM) | 279 | 152 | 224 | 137 | 168 | 192 | 146 | 172 | 182 | 171 | 186 | 64 | 140 | 2213 |
| # of SNPs | 37 | 30 | 56 | 36 | 49 | 24 | 24 | 22 | 29 | 15 | 22 | 16 | 8 | 368 |

**Fig. 8.**
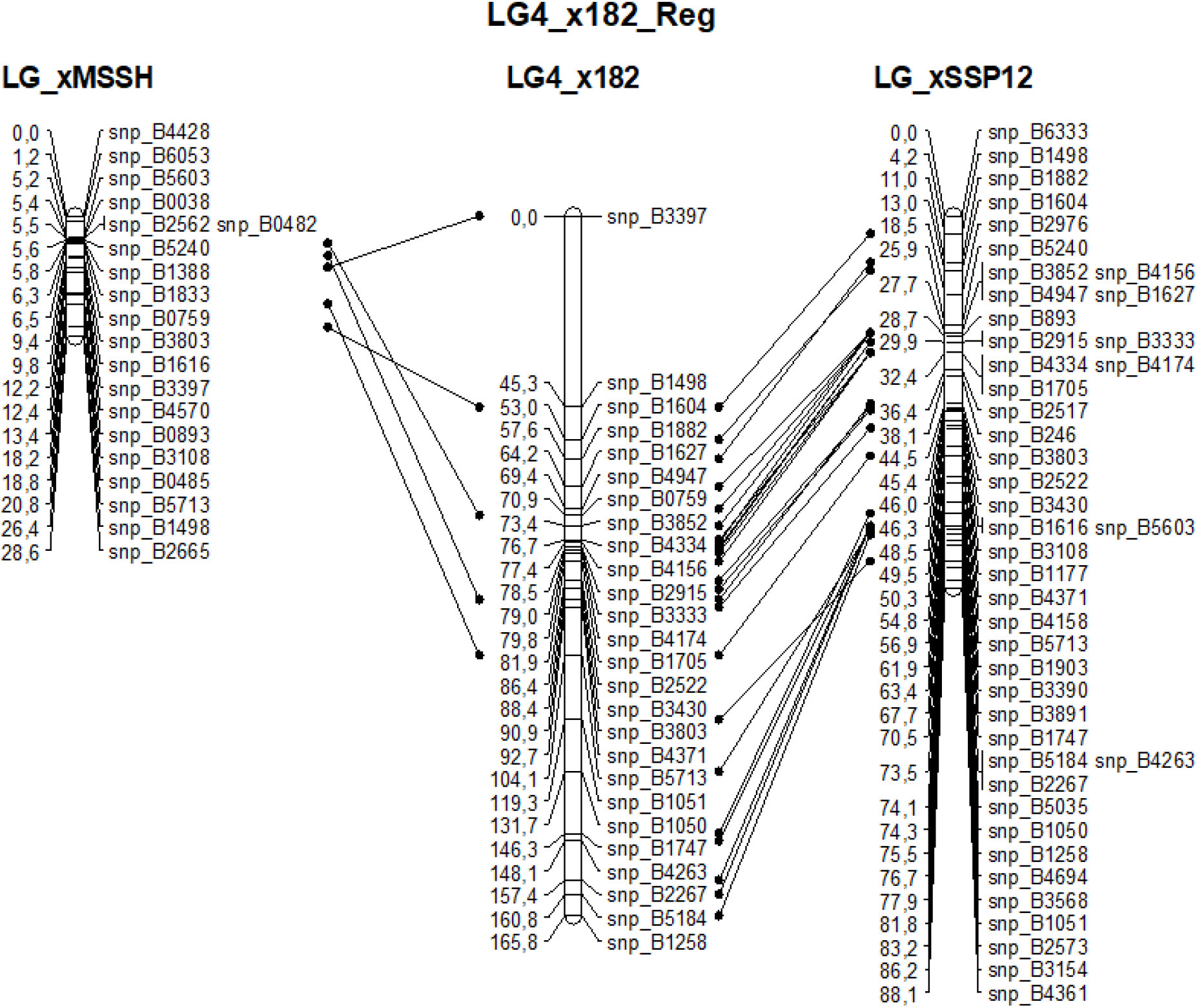
Example of the high collinearity between homologous maternal (MSSB) linkage group #4 obtained from the analysis of three sets of offspring: xMSSH and xSSP12 correspond to the two largest full-sib families and x182 correspond to the cosegregation analysis of their mapped markers in the 182 half-sibs.

#### 3.2.5. Alignment of Bhaga genomic scaffolds to the SNP-based linkage map of beech

The 368 mapped markers were aligned on the 12 genomic scaffolds (Bagha_1 to Bagha_12) of the Fagus sylvatica genome assembly. The alignments were filtered and congruence between scaffolds and linkage groups were checked. Most of the markers from a given LG mapped on a single scaffold (Table 6) providing a genetic validation of the physical assembly obtained for the Bagha genome sequence. Notable exceptions were: (i) LG11 and LG12, which corresponded to Bagha_#8; these two chromosomal arms could not be merged into a single LG, and (ii) LG13 and scaffold #11, which presented too few markers for unambiguous assignment to one or more scaffolds and LGs, respectively.

**Table 6.** Number of SNP markers of a given linkage group (LG) aligned to a specified scaffold (Bhaga_i) of the *Fagus sylvatica* assembly.

|  | Bhag a_1 | Bhag a_2 | Bhag a_3 | Bhag a_4 | Bhag a_5 | Bhag a_6 | Bhag a_7 | Bhag a_8 | Bhag a_9 | Bhag a_10 | Bhag a_11 | Bhag a_12 |
| --- | --- | --- | --- | --- | --- | --- | --- | --- | --- | --- | --- | --- |
| LG1 | 2 |  |  |  |  | 26 |  |  |  |  |  |  |
| LG2 | 1 |  |  |  | 1 |  | 23 |  |  |  |  |  |
| LG3 |  | 42 |  |  |  |  |  |  |  |  |  |  |
| LG4 |  | 1 | 1 |  | 22 | 1 | 1 |  |  |  | 1 |  |
| LG5 |  |  |  |  |  |  |  |  |  |  |  | 42 |
| LG6 | 1 |  | 16 | 1 |  |  |  |  |  |  |  |  |
| LG7 | 16 |  | 1 |  |  |  |  | 1 |  | 2 |  |  |
| LG8 |  |  |  |  | 1 |  |  |  |  | 15 |  |  |
| LG9 |  |  |  |  |  | 1 |  |  | 25 |  |  |  |
| LG10 | 1 |  |  | 12 |  |  |  |  |  | 1 |  |  |
| LG11 |  |  |  |  |  |  |  | 20 |  |  |  |  |
| LG12 | 1 |  |  |  | 1 |  | 1 | 10 |  |  |  |  |
| LG13 |  | 1 |  |  | 1 | 1 |  |  |  |  | 2 |  |

## 4. Discussion

### 4.1. General genome features

The genome assembled and analysed in this study compares well with previously published *Fagaceae* genomes, both in terms of size and gene space. We here confirm the base chromosome number of 12, as was previously reported based on chromosome counts (Ribeiro et al., 2011). The number of exons per gene is moderately higher than in the previously published genome of the same individual (Mishra et al., 2018), reflecting the higher contiguity of the presented chromosome-level assembly. Despite the lower chromosome number of the beech genome, it is structurally similar to the available genomes of genus *Juglans*, which is the most closely related genus for which chromosome-level assemblies are available, with continuous sequences from telomere to telomere (*J. regia* (Marrano et al., 2020); *J. sigillata* (Ning et al., 2020); *J. regia × J. microcarpa* (Zhu et al., 2019)).

### 4.2. Telomere and centromere predictions

Telomeres are inherently difficult to resolve because of long stretches of GC-rich repeats that can cause artefacts during library preparation (Aird et al., 2011) and can lead to biased mapping (Dohm et al., 2008). However, using long-read sequencing and Hi-C scaffolding, we could identify telomeric repeats on all chromosomes. It seems likely that several of the unplaced contigs of 4.9 Mb, which included telomeric sequences, were not correctly anchored in the assembly due to ambiguous Hi-C association data resulting from the high sequence similarity of telomeric repeats, because of which for four chromosomes we could identify telomeric repeats only on one of the ends. This might also be due to the presence of ribosomal cistrons on four chromosome ends, which might have interfered with the Hi-C linkage due to their length and very high sequence similarity. On the outermost regions of the chromosomes, no longer telomeric repeat stretches were present most likely due to their ambiguous placement in the assembly, because of very high sequence similarity.

Centromere repeats were identified by screening the genome for repeats of intermediate sizes, and were found to be present predominantly within a single location per chromosome. However, lower amounts of centromeric repeat units were also observed to be scattered throughout the genome. The function of the centromeric repeats outside of the centromere remains largely enigmatic but could be associated with chromosome structuring (Alves et al., 2012) or centromere repositioning (Mandáková et al., 2020; Klein and O’Neill, 2018). Interestingly, we could find two major groups of potential centromeric repeat units of different lengths, which did not always coincide. The location of the main occurrence of the centromere-defining repeat unit agreed well with the location previously inferred using chromosome preparations and fluorescence microscopy (Ribeiro et al., 2011).

### 4.3. Integration of organelle DNA in the nuclear genome

Organelle DNA integration has been frequently found in all kingdoms of life for which high-resolution genomes are available (Zhang et al., 2020; Guo et al., 2008; Stegemann et al., 2003). It can be assumed that this transfer of organelle DNA to the nucleus is the seed of transfer of chloroplast genes to the nuclear genome (Huang et al., 2003). However, apart from a few hints (Yang et al., 2017) it is unclear, which factors stabilise the chloroplast genome so that its content in non-parasitic plants stays relatively stable over long evolutionary timescales (Xiong et al., 2009; Wang et al., 2007). In the present study, it has been found that the insertion of organelle DNA insertions are located mainly in repeat-rich regions of the beech genome. However, their presence in regions without pronounced repeat density might suggest that repeats are not the only factor associated with the insertion of organelle DNA. Nevertheless, it appears that some regions are generally amenable to the integration of organelle DNA, as in several cases chloroplast and mitochondrion insertions were observed in close proximity. The reason for this is unclear, but is known that open chromatin is more likely to accumulate insertions (Wang and Timmis 2013). The potential presence of areas in the genome that are less protected from the insertion of foreign DNA could open up potential molecular biology applications for creating stable transformants.

An anomaly regarding organelle DNA insertion was observed on chromosome 11. Around a central insertion of mitochondrion DNA, multiple insertions of chloroplast DNA were found. The whole region spans more than 2 Mb, which is significantly longer than the organelle integration hotspots reported in other species (Zhang et al., 2020). The evolutionary origin of this large chromosome region is unclear, but given its repetitive nature it is conceivable that it resulted from a combination of an integration of long fragments and repeat element activity. The presence of multiple copies at the location implies an unusual genome structure in this area, but further analyses, ideally including multiple additional individuals, will be necessary to elucidate the basis for this.

### 4.4. Distribution of single nucleotide polymorphisms (SNPs)

SNP content was found to vary across all chromosomes leading to a mosaic pattern. While most of the areas of high or low SNP density were rather short and not correlated to any other patterns, there were several regions > 1 Mbp that exhibited a similar polymorphism type, suggesting non-neutral evolution.

The longest of those stretches poor in both heterozygous and homozygous positions was found on chromosome 4, and corresponded to a region rich in both genes and repeat elements. This is remarkable and probably due to a recent proliferation, as repeat-rich regions are usually less stable and more prone to accumulate mutations (Wang et al., 2020; Flynn et al., 2018; Ho et al., 2020).

Most regions with lower abundance of heterozygous SNPs than on average were found to be particularly high in homozygous SNPs. The longest of such stretches was found on chromosome 6, comprising about two thirds of the entire chromosome. Three more such regions longer than 5 Mbp were found on other chromosomes. The evolutionary significance of this is unclear, but it is conceivable that these areas contain locale specific variants for which no alternative alleles are shared within the same stand. For confirmation of this hypothesis, it would be important to evaluate genetic markers from additional individuals of the same stand. Locally adaptive alleles could be fixed relatively easy by local inbreeding (Ceballos et al., 2018), considering the low seed dispersion kernel of European Beech (Martínez and González-Taboada, 2009). The presence of genes involved in local adaptation could explain the rather high amount of homozygous SNPs in the same location, as the stands from which the two studied individuals came from differ in soil, water availability, continentality, and light availability. However, more individuals from geographically separated similar stands need to be investigated to disentangle the effects of inbreeding and local adaptation.

In summary, homozygous and heterozygous SNPs were rather uniformly distributed throughout the major part of the genome, suggesting neutral evolution or balancing selection.

## 5. Conclusions

The chromosome-level assembly of the ultra-centennial individual Bhaga from the Kellerwald- Edersee National Park in Germany and its comparison with the individual Jamy from the Jamy Nature Reserve in Poland has revealed several notable genomic features. The prediction of the telomeres and centromeres as well as ribosomal DNA corresponded well with data gained from chromosome imaging (Ribeiro et al., 2011), suggesting state-of-the-art accuracy of the assembly. Interestingly, several anomalies were observed in the genome, corresponding to regions with abundant integrations of organelle DNA, low frequency of both heterozygous and homozygous SNPs, and long chromosome stretches almost homozygous but with a high frequency of SNPs differentiating the individuals.

Taken together, the data presented here suggest a strongly partitioned genome architecture and potentially divergent selection regimes in the stands of the two individuals investigated here. Future comparisons of additional genomes to the reference will help understanding the significance of variant sites identified in this study and shed light on the fundamental processes involved in local adaptation of a long-lived tree species exposed to a changing climate.

## Supporting information

All supplementary files except for S6

## 6. Data availability

The data sets supporting the results of this article are available in the GenBank repository, under the accession number PRJEB24056 for the *Fagus sylvatica* reference individual Bhaga, PRJNA450822 for the individual Jamy, PRJEB46583 for sequencing of a new unigene set and PRJEB46593 for RNA-seq-based genetic mapping in European beech.

## 8. Author Contributions

M.T. conceived the study, wih contributions from C.P. and J.B. B.U., C.B., J.B., J.M., J.M.A, L.O., M.T., and S.P. provided materials. All authors conducted laboratory experiments or analysed the data. All authors were involved in data interpretation. B.M. and M.T. wrote the manuscript with contributions from the other authors. All authors read and approved the final manuscript.

## 9. Funding and Acknowledgment

This study was supported by grants of the German Science Foundation (Th1632-18-1), National Science Centre, Poland (2012/04/A/NZ9/00F500), the Polish Ministry of Science and Higher Education under the program “Regional Initiative of Excellence” in 2019–2022 (Grant No. 008/RID/2018/19), and the LOEWE initiative of the government of Hessen in the framework of the LOEWE Centre for Translational Biodiversity Genomics (TBG).

For the validation of the chromosome-scale assembly, we thank (i) INRAE (Ecodiv division) and the Genoscope: the Commissariat à l’Energie Atomique et aux Energies Alternatives (CEA), (ii) France Génomique (ANR-10-INBS-09-08, Beech genome project) for providing funding for sequencing, (iii) the Genotoul Bioinformatics Platform Toulouse Occitanie (Bioinfo Genotoul, https://doi.org/10.15454/1.5572369328961167E12) for providing computing resources, (iv) the Genome Transcriptome Facility of Bordeaux (grant from Investissements d’Avenir, Convention attributive d’aide EquipEx Xyloforest ANR-10-EQPX-16-01) which carried out the mass array assay. IL received funding from the European Union’s Horizon 2020 research and innovation program (GENTREE project, No. 676876). We thank many of our colleagues from INRAE (UMR Biogeco) including : Benjamin Dencausse (seed collection in Saint-Symphorien, plantation and tissue sampling in Nouzilly and Guémené-Penfao), Jean-Charles Leplé (bud collection in Nouzilly), Parick Reynet, Edith Reuzeau, Yannick Mellerin (seed collection in Saint-Symphorien), Martine Martin-Clotte (sample preparation for ONF nursery), Maxime Doeland, Jean Broué (DNA extraction and genotyping), Adib Ouayjan (selection of SNPs for pedigree reconstruction), Adline Delcamp and Emilie Chancerel (mass-array genotyping). We are grateful to the administrative nursery of Guémené-Penfao managed by the Office National des Forêts (ONF) especially Jean-Pierre Huvelin and Olivier Forestier (germination of seeds of the 1st and 2nd campaign, plantation of the 2nd campaign and monitoring of the plantation), as well as the experimental units PAO of INRAE Nouzilly (Benoit Luwez) and GBFOR of INRAE Orléans (Dominique Veisse) for the plantation of the 1st campaign and the monitoring of the plantation.

## Supplementary Files

**Supplementary file 1**. Details of annotated repeat elements in Fagus sylvatica.

**Supplementary file 2**. Venn diagram showing shared PLAZA proteins of *Arabidopsis thaliana* (27615), *Eucalyptus grandis* (36331), and *Vitis vinifera* (26346) with those of *Fagus sylvatica* (28326).

**Supplementary file 3**. Centromeric feature annotation.

**Supplementary file 4**. Details of the conservation of organelle DNA insertions in the nuclear genome.

**Supplementary file 5**. Multiplexed SNP assay. SNP_ID (a) following Ouayjan et al. (2018). SNPs discarded from the analyses are highlighted in yellow; Locus_Name_pos_SNP (b) corresponds to the locus name given by Lalagüe et al. (2014); seq SNP corresponds to sequences of the SNP flanking regions. The targeted SNP is indicated in brackets [ / ].

**Supplementary file 6**. Genotyping data. List of 4127 SNPs and their associated linkage groups based on R_qtl (second raw) and JoinMap (third raw) analyses.

