## Supplementary material for "A chromosome-level genome assembly of the European Beech (*Fagus sylvatica*) reveals anomalies for organelle DNA integration, repeat content and distribution of SNPs": All supplementary files except for S6

Supplementary File 1. Details of annotated repeat elements in *Fagus sylvatica*.

| Repeat elements | # of elements | size | percentage of genome |
|---|---|---|---|
| Retroelements | 87628 | 81859802 bp | 15.15 % |
| LINEs: | 36016 | 19755558 bp | 3.66 % |
| RTE/Bov-B | 5707 | 963701 bp | 0.18 % |
| L1/CIN4 | 30309 | 18791857 bp | 3.48 % |
| LTR elements: | 51612 | 62104244 bp | 11.49 % |
| Ty1/Copia | 26740 | 25500497 bp | 4.72 % |
| Gypsy/DIRS1 | 22010 | 34391985 bp | 6.36 % |
| DNA transposons | 27884 | 14339425 bp | 2.65 % |
| hobo-Activator | 14367 | 4870881 bp | 0.90 % |
| Tourist/Harbinger | 4205 | 1873197 bp | 0.35 % |
| Rolling-circles | 7484 | 3397980 bp | 0.63 % |

Supplementary File 2. Number of PLAZA genes shared between seleected highly resolved plant genomes.

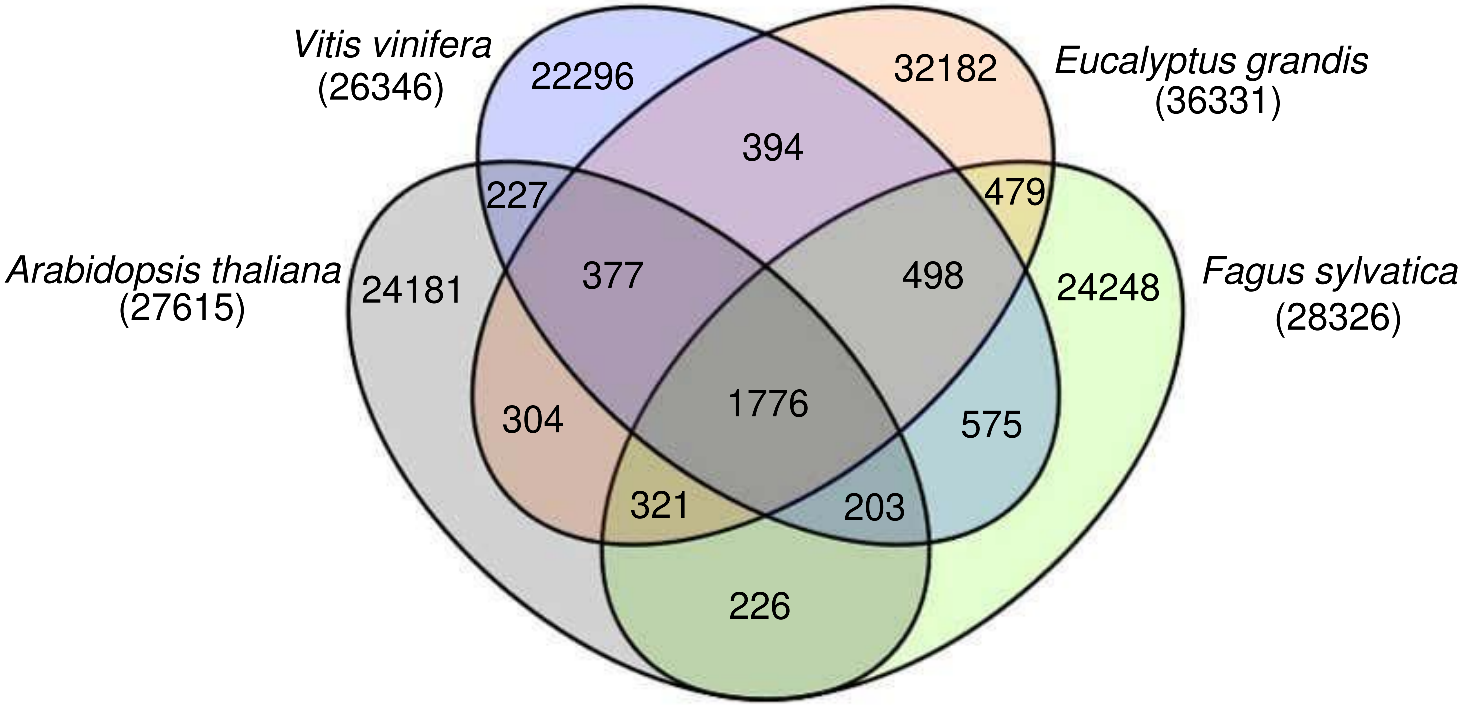

Supplementary File 3. Annotation of centromere-associated features along the 12 chromosomes.

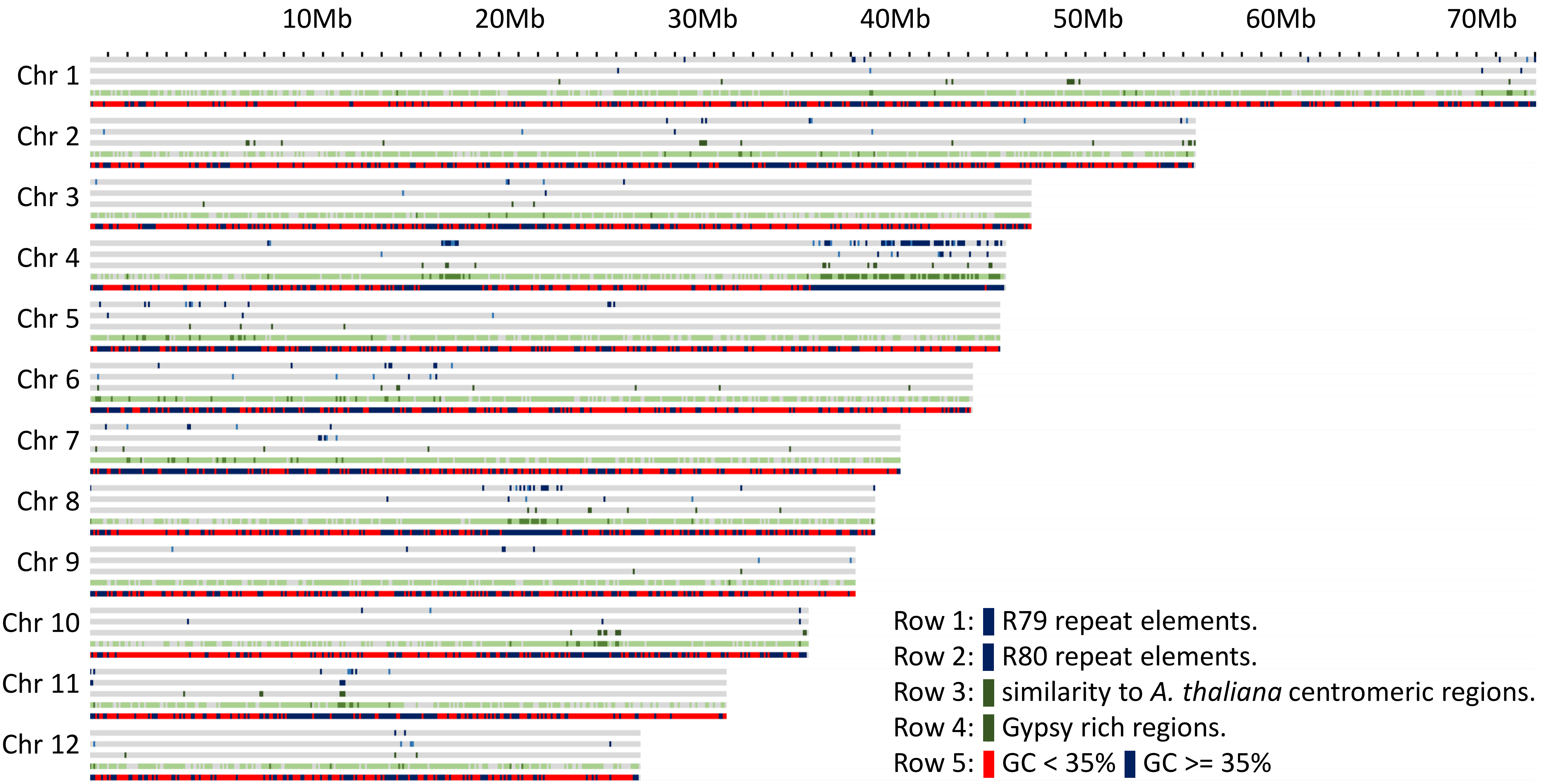

Supplementary File 4. Details of the insertion of organelle genome fragments into the nuclear genome.

Chloroplast Insertions

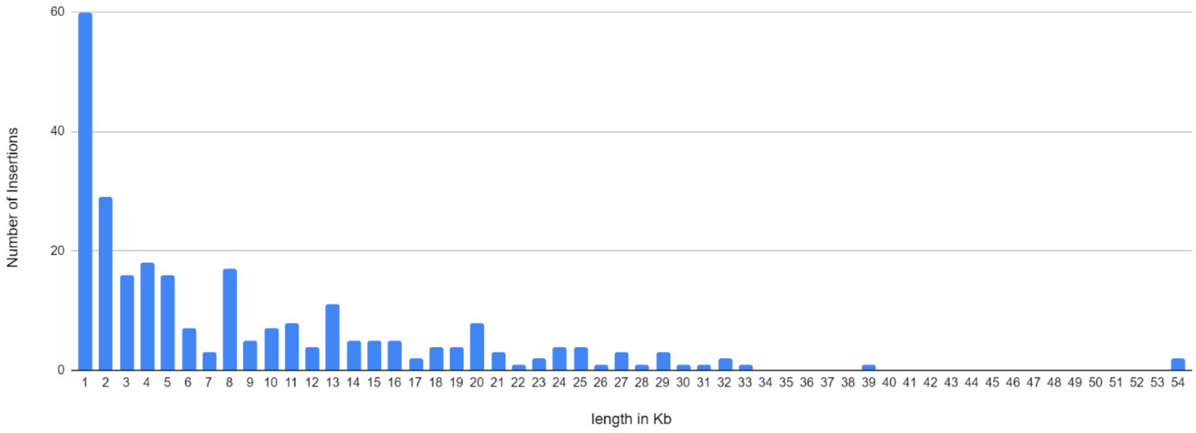


Column chart showing number of insertions for different fragment sizes of chloroplast DNA. The horizontal axis shows the sizes of the fragments (in kb). The vertical axis shows the numbers of insertions for the specified fragment sizes. Inserted fragments less than 1 kb in length are not included in this plot.

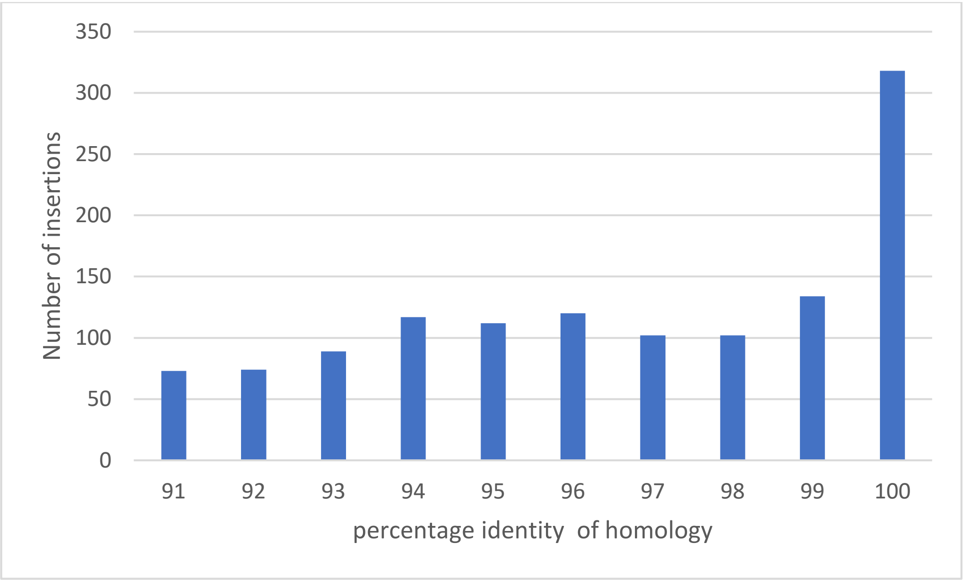


Column chart showing number of insertions at different percentage identity level of the chloroplast genome fragments from 90% to 100% identity. Inserted fragments less than 100 bp in length are excluded for this analysis.

Mitochondrial insertions

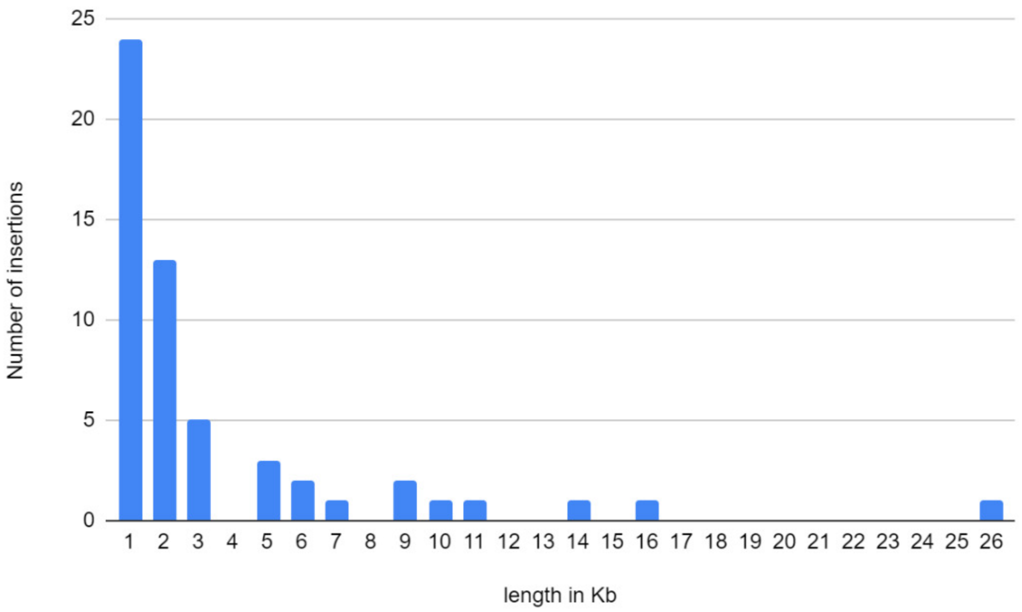


Column chart showing number of insertions for different fragment sizes of mitochondrion DNA. The horizontal axis shows the sizes of the fragments (in kb). The vertical axis shows the numbers of insertions for the specified fragment sizes. Inserted fragments less than 1 kb in length are not included in this plot.

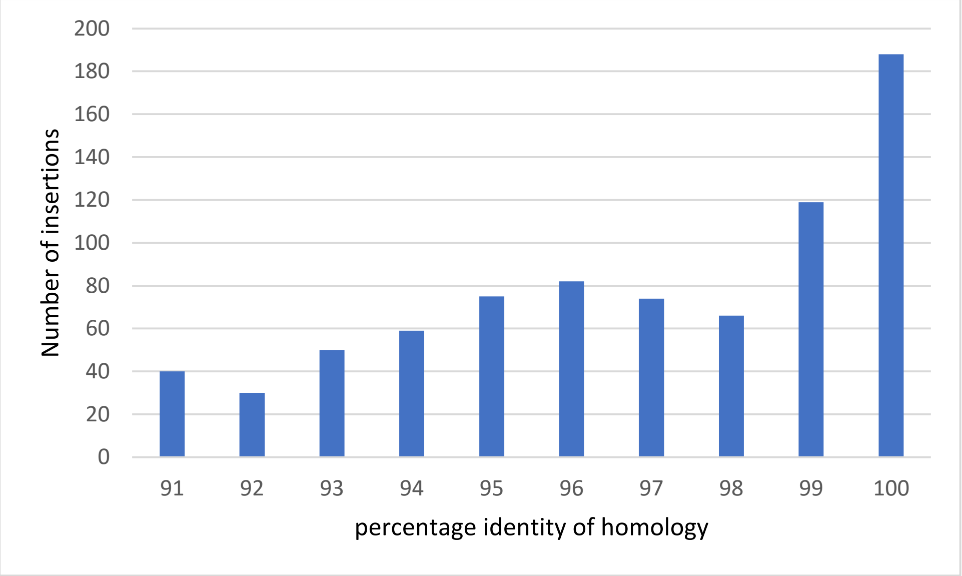


Column chart showing number of insertions at different percentage identity level of the mitochondrion genome fragments from 90% to 100% identity. Inserted fragments less than 100 bp in length are excluded for this analysis.

**Supplementary File 5** Multiplexed SNP assay

| SNP number | SNP_ID (a) | Locus_Name_pos_SNP (b) | Seq SNP |
|---|---|---|---|
| 1 | 10 | 00066_edited.merged.removebadseq.renamed-600 | TCTACAAAACTAATGAAAGTTCAGCAGCTCTAATGACTGAGGATTCTGGTATTAGGAAGATTCTTCCTAAATCYAAGATATGTGACTTTGAGTTTGACCC[C/T]TGCGGGTACTCCATGAATTCTATCGAAGGYGATGCAATTTCTACAATCCATGTTACACCAGAAGATGGTTTCAGTTATGCRAGCTTTGAAGCAGCTGGCT |
| 2 | 33 | 00133_edited_HIrecoded.merged.removebadseq.renamed-637 | TTTTCCACTTCAAATGAATTCATTTGGTAACTGCACCATTTGCTGGGAGGTGAAACAGACATCTGTTGTTGGACTTCCCTWACGTGTTAGTGAATTTTGG[A/T]TGAAGTTGAAATGTGTCTACTGCTAAAGGAGGATTCCTTCTAGAGCCACGGAGATGCCTGAGTAGAATCCTGTCTTCTATKGCCATACTTGATGTTCCTT |
| 3 | 46 | 00142_edited_HIrecoded.merged.removebadseq.renamed-375 | ATATTGTTAATMAGAAWTTTTTTTTTYYCCTCCATCCCYRGCATAGGTAAATGYGTATGAGATGTATTAATCATTCYCCCTCAGAATTGATTTTCTGTGG[C/T]GGCATTTTCAGATGCATGCCTTGTTTTAACTTCWCTGGCTWGYACAGAATGAAGAGTACTCTGAGAAAACCATTTCCCCTTTRAGATAATGCGAGTTTCT |
| 4 | 50 | 00142_edited_HIrecoded.merged.removebadseq.renamed-458 | GAATTGATTTTCTGTGGYGGCATTTTCAGATGCATGCCTTGTTTTAACTTCWCTGGCTWGYACAGAATGAAGAGTACTCTGAGAAAACCATTTCCCCTTT[A/G]AGATAATGCGAGTTTCTAGTTACTTTGGTTGCCAATTGCTCTT |
| 5 | 61 | 091_2_edited_HIori.merged.removebadseq.renamed-231 | ACATTTTGCCMCTGCAGCCAGTTGGCCGCYTGGTCTTGAATAAGAACATTGATAACTTCTTTGCAGAAAATGAACARCTTGCATTTTGCCCTGCCATTGT[G/T]GTTCCTGGTGTCTACTACTCAGATGATAAGTTGCTCCAGACTCGTATCTTTTCCTATTCTGATACTCAGAGGCACCGTCTCGGACCAAACTATCTGCAGC |
| 6 | 64 | 091_2_edited_HIori.merged.removebadseq.renamed-479 | GGTTTCATGAATTTCATGCACAGGGATGAGGAGGTACTTAAAGCTGGCTACTGCATCTTTTAGTTAGTCRGACTTGTATGCTACCTTAGTGCYTTAGGCC[C/T]ACAAATGGTGTAAGATGCTAAGGCRGTATTGCAATTTTATGCTTGTCTAATTYGTGTGAATTTATTCTGATAAGATATTATGATCATAATCTTTAGGTCA |
| 7 | 81 | 102_edited_HIrecoded.merged.removebadseq.renamed-745 | ATCAGAAGAGTACTGAGCTCCTGATCAGGAAGYTGCCCTTCCAGAGGCTTGTCCGTGAAATTGCCCAGGACTTCAAGGTRTTGTTTTTATTTATTTAGTT[G/T]GGTATTGTTTGGGTTTTGAGTTTGTTTTGTTTTGTTGCTAATGGGTTTGTTTTTATGTTTCAGACTGATCTGCGTTTCCAGAGTCACGCTGTGTTGGCTC |
| 8 | 87 | 10_edited_HIrecoded.merged.removebadseq.renamed-381 | AGTGTTCTGATGCAAATTGAAGTTAGATTGACTTTCAATGGCTTCTTCAGACTTCCAGTAGTCTGTACCTGTTTCAAAATACTGGTTTGGTCTGTGATTT[C/T]GGTAATTCTACAGGATTGCCATGTAMTTAAATATCGTTATCAAAATGTAAATAAACTGCGTTTTWTTTTATTAAAAATAAAAAAAATAARRGTTTTCCTC |
| 9 | 101 | 10_edited_HIrecoded.merged.removebadseq.renamed-978 | TTGACAAGAATGCCAAGATAGGAAGAAATGTGGTCATYGCAAATGGTGATGTAAGTTTGGCTCTCTGTGCTCTTTAAATTTTAAATTGCTCTGCACTTCA[A/C]GCTTATCCAAGTTATTGAAGAATRGTGTTTCACTGTTCATTGACACTGTRCTTAMGGAAATTCTTTGACTTCTCAGGGTGTTGAGGAAGCAGATAGGCCA |
| 10 | 109 | 110_3_edited.merged.removebadseq.renamed-173 | GGTGGCCGAAGAACCAACATCTTCGATCCATTTTCTCTCGACATGTGGGATCCCTTTGATGGTGTTTTCACCTCTGCACTTGCCAATGTACCTGCCTCTG[A/C]ACGTGAAACGTCTGCGTTTGCCAACACAAGGATTGATTGGAGGGAGACCCCAGAAGCTCATGTTTTCAAAGCTGACCTTCCGGGGCTGAAGAAAGAGGAG |
| 11 | 140 | 130_edited.merged.removebadseq.renamed-363 | GGCTGAAGCTGCTGGGTCTGGAGTTTCTCCATCCCTCAAGAACTTCTTGCTCAGCATTGTGGCTGGTGGTGTTGTGYTTACTRTRATTGTTGGTGCTGTG[A/G]TAGGAGTCTCCAACTTTGATCCAGTCAAGCGGACCTGAGAGACCAATTCTGAGAAGAGAATGCTGTAATTTTATAAATTCACTYTCATGTATCTGTGTCA |
| 12 | 154 | 134_2_edited_HIrecoded2.merged.removebadseq.renamed-50 | GAGGAAACTGTGGATGTGGTACTGGCTGCAAGTGCGGCAGCGGCTGTGG[A/C]GGGTAATCTTCTAACTTCTTCTGTATTTCATCTTKSTGTTTTCCTAACTATATGCTTAGCATATCCGGCGAGGCTTATTGGTATCACGCGTCGCGAGCGT |
| 13 | 180 | 145_2_edited_HIrecoded.merged.removebadseq.renamed-722 | AATGAAACGCAAACAGTGTTGTCTTTTYTAATGACAATTAAATGTAATTTAACTTTGTGTTTGAAAGTTTTACCCTATTTACATTATGCAAAAAGCTAAA[A/C]AACATCTTAGCTTTCTAGCCAYATTATCTTGTTCCTCTGCGAATGAGCTTTRGCTCAAATGGGTACTTCCTCCTYCCATAAGAATGKGTRGAGTATTAGG |
| 14 | 194 | 148_1_edited_HIrecoded.merged.removebadseq.renamed-378 | TGGGACGCCGAGTTYGTCAAAGTTGACCAGGCCACCCTCTTTGATCTCATTCTGGTATAATTACCTTTTTACTCTCTTATTATCGTTTTCTTTGGGCTAG[G/T]KCTTCCAGATTTGGATCTGGATGTTAGTTAGGGTTTCGCTAKYCTGAATTTACGATTCTAGGGTTYCGAAATKTTGAGGATTTTCGATCGAAATTTGCTT |
| 15 | 221 | 150_2_edited_HIrecoded.merged.removebadseq.renamed-381 | TGTTTATACCATGAATTGTCGTTCTTAATTATATTCCTTTTATTATAAATTTAACCTTTTATRGTTGCCTTCAGCATTCTTTCTTGTTTCTGATTTGACA[A/G]TTGTGAATTTTAACAGGTTTACAAGTCAGGGTCCTTCGGGAGGTCACTGGATATCACCAAATTTACTAGCTACCATGAGTTGCGTGGTGAGTTGGCTCGT |
| 16 | 226 | 150_2_edited_HIrecoded.merged.removebadseq.renamed-693 | GTTTCCATCATTGTTTGTTGTTTCCTTTTCTTTTATAATTATAACTGCRTGATAGGACYGTTAATTTTTTAAAATTRTCTTTTGGATYGGAAGGCATTTT[A/G]AGTATTCATTTGCATCTCAGAGCCTAAGACTCAATTTAAATCATTTGGACCATCTGATATCAGAATTTTTCTGGCATGTTTCATATGAAAAAAATTAATA |
| 17 | 231 | 150_2_edited_HIrecoded.merged.removebadseq.renamed-956 | CATTTGTATGATGCTGTAAMTGGTGGTCATGGAGGTCTATATTCTTTATACRAATRTCTTATTTGATTTGGAGATTCTATCCTGATGTCCACACTTAATT[A/G]GYCAAGCTTGATAAGATAGTGATTTTTCCACAACAGGGAATTTGTGAACAGTGTGTGGTGTATCAAAATACTATCACCACAAGAAGTGCAGCAAATGGGC |
| 18 | 236 | 154_1_edited.merged.removebadseq.renamed-34 | TCCTCATCACCCCTTTGAAAAAAGCCCCAATAT[C/T]ACCACAAATATTTCACACCTACACCTCCACAAACACATCCAAGTAGCCCACTCAACATGCCAAGGCACACTATACCCAGAACTATGCGTCTCAACACTCT |
| 19 | 260 | 155_2_edited.merged.removebadseq.renamed-444 | ATGCCCAAAAGGTCGTTTTTGCCCCGGGTGATTTTGTCGAAATTACCATTTTCGACCTTGCATCTCTCTGAKATGAAGGAAATTTTCCACGCGCAGGAGA[A/G]ATCTAGTATGGAGCRCGTATTGRTGAACGCGCTGGCRGAGTGTGAGAGGGCTCCGAGCCTGGRTGAGACCAAGCRGTGCGTTGGGTCGGTGGAGGACATG |
| 20 | 267 | 155_2_edited.merged.removebadseq.renamed-574 | CGCTGGCRGAGTGTGAGAGGGCTCCGAGCCTGGRTGAGACCAAGCRGTGCGTTGGGTCGGTGGAGGACATGATAGACTTTGCYGTCTCGGTCTTGGGCCA[C/T]AATGTRGTTGTGAGGAGCACTGAAAATGTGWGTGGGTCAAAGCAAAACGTGATGATTGGATCAGTCAAASGSATCAACGGTGGAGAAGTGACGAAATCGG |
| 21 | 283 | 155_3_edited.merged.removebadseq.renamed-169 | TGTCTAGTTATCTGGATCACATTTTGGTACTGGTGATGACTGCATCKCTGTTGGACAAGGAAATTCTCAGGTCACTATTTCAGGTGTCAACTGTGGACCT[C/G]GTCATGGCATCAGGTACTTTTGAAATCTCCAATGCTTGTAACCTCTATGTTTACTTGGGCACATACATCATTTATGTGAAGCTACTTAATGTAATATATT |
| 22 | 291 | 17_edited.merged.removebadseq.renamed-1081 | ACCATCCACCGGCTTCCATGGCTATGTATGGGGTGACTAAAACAGCCCTTCTTGGTCTTACCAAGGTATGACATGGTTCCCATATCTATGTTTTATGTTA[G/T]TTGTGTATTCAAACAAATCATGGGGTATTCTTCTTTCCTTGTTCTGATTCTTTTTCAATCATCATTTAGGCCCTTGCAGCTGAGATGGCTCCAGATACAC |
| 23 | 296 | 19_edited_HIrecoded.merged.removebadseq.renamed-215 | CCRACCACCGGTGGAGTCAAGAAGCCCCATCGTTACCGYCCTGGTACTGTTGCCCTCCGGTGAGTCAGCTTATTTATCTCTGTTTTGCYTTTTAATGAAA[C/T]TTGCTTTATTGGTGATTTGCCTTATACGTGTTTGTCAAATTGTTACAGTGAAATCCGTAAGTATCAGAAGAGTACTGAGCTCCTGATCAGGAAGYTGCCC |
| 24 | 334 | 23_1_edited_HIori.merged.removebadseq.renamed-741 | KATTTAAATAGCATATGTGCAYGAATATGCTTGRTTTTTSRTGATTATCCTTACATTCTTATGGAAACATTTTGGCMAAAACCGTCTTGTTTTTGGATGA[A/G]TAAGTAATTTATTCAAACCAGAAGAGACTTGCAAGCAGGTTACMAKAAGAAACWGCAAGCCTTRACACTCAATACAGCTCCAMAAACTTCAAACAAAAAT |
| 25 | 340 | 23_1_edited_HIori.merged.removebadseq.renamed-852 | ATTCAAACCAGAAGAGACTTGCAAGCAGGTTACMAKAAGAAACWGCAAGCCTTRACACTCAATACAGCTCCAMAAACTTCAAACAAAAATCACTAATATA[A/C]ATTAGACTCCACAAAGCAAGGACCATTTTGTTATGCTAATGATTACATATATGCTCATTTTGATGTGAACTCTTAGGGGATTGAGATTTTCATGATGTTT |
| 26 | 375 | 30_2_edited_HIrecoded.merged.removebadseq.renamed-435 | GAGTATTATTTGACTCCCAAACTAAACCATGTGGGCTACATAATAATATTGTTGTTCYTTTTTTATGTTTTTKGGTTAAATNATGATCTCATGGGCTAGA[A/C]ATGGGTGTTGGCTGCAGGTGTGGAGCATGTTGGGGGAGACATGTTTGTTAGTGTTCCTAAAGGAGATGCCATTTTCATGAAGGTGGGTTTGWACTTATCT |
| 27 | 384 | 30_2_edited_HIrecoded.merged.removebadseq.renamed-936 | TAYTTTTTGTGGAAAGACACTTTAACCTATTAACTAGCACATGAATCATAAGGTGAAAATGAAATTTAAAGAAAAATTGATTAGTCAAAGACATTTGATC[C/T]TTGWTTATGCTAYTGACATGTGTGTGCTCCTATGTACAGTGGATATGCCATGATTGGAGTGATGAGCACTGCTCAAAATTTTTGAAGAACTGCTATGATG |
| 28 | 387 | 30_2_edited_HIrecoded.merged.removebadseq.renamed-1173 | GTATTCTTCCTGTAGCCCCGGACAGTAGCCTTGCTGCTAAGGGAGTCATCCATATTGATGTTATCATGTTGGCTCATAATCCGGGTGGGAAAGAGAGGAC[C/T]GAGAAGGAGTTCGAGGMCTTGGCYAAAGGGGCAGGATTTCAAGGGTTCAAAGTAGTGTGTTCTGCTTTCAACACTTACATCATGGAATTCCTTAAAAAGM |
| 29 | 395 | 39_edited.merged.removebadseq.renamed-256 | GTCCTGCAACTCATCGGTCCTATAGGTGTATTGTCAGGTATGGATATCGCGATGTTCATCAGGATATCCATTCCTTTGAAACTGAGCTAGTTGAAAGACT[A/C]GCTGATTTCATTCGCTATGATTGGCRKAGAACAAATGGTACTAATTCTTGCAATGAGGATGATGCATCAAGATCTAATGAATCAACTAGTGAGTGTAGAT |
| 30 | 412 | 50_edited_HIrecoded.merged.removebadseq.renamed-320 | AGATGACGTGGARGCTGGTTCTGACGTGTCATTATGGAGTTATTCAATTTGATTGTTATTACTATTAATTATAACAATCATGTYGTAATAYATGTTAGAA[A/C]GTTTGGTTTAGCATTATAGCTGCTTGCATTTTAGACAGTCTGATCATTGTTACTTGTTAGGGAAAGAATTGTTACTTTCCGTGCTTATTGTTTTTGATGA |
| 31 | 435 | 52_1_edited.merged.removebadseq.renamed-235 | CAAACCGCYGTACTCATYGAAACCCTCACTGCTCTCGGCGCTGAGGTTCGTTGGTGTTCATGCAACATCTTCTCAACCCAAGACCACGCCGCCGCCGCCA[C/T]CGCTCGCGACWCYKCTGCTGTGTTCGCGTGGAAGGGAGAGACWCTBCAGGAGTACTGGTGGTGTACCGAGCGTGCACTTGATTGGGGCCCAGGTGGTGGA |
| 32 | 440 | 52_1_edited.merged.removebadseq.renamed-350 | CTGCTGTGTTCGCGTGGAAGGGAGAGACWCTBCAGGAGTACTGGTGGTGTACCGAGCGTGCACTTGATTGGGGCCCAGGTGGTGGACCCGATCTCATCGT[C/T]GATGAYGGTGGTGATGCYACTTTGTTGATCCATGAGGGTGTGAAAGCTGAGGAGATCTATGAGAAGACTGGTACCGTTCCAGACCCTTCATCYACAGACA |
| 33 | 444 | 52_2_edited.merged.removebadseq.renamed-65 | ATGTYATGATTGCCGGAAAGGTTGCCGTTGTCGCTGGATATGGTGATGTGGGAAAGGGTTGTGC[A/T]GCTGCCTTGAAACAAGCTGGAGCCCGTGTCATCGTSACTGAGATTGATCCAATTTGTGCTCTTCAGGCCCTTATGGAAGGCCTCCAAGTTCTAACCCTYG |
| 34 | 470 | 58_edited_HIori.merged.removebadseq.renamed-736 | AGAACTTTTTTCAATGACCTCAATCTTTGTATACTTTCATGCTGCTTAACTTTTTTTGGTGATGCAGGGGAGAGATTGGGAAATGGATTGCTTTYGTTGC[A/T]GTTATATTGCGCCTTTTTTTCCCACGTCATTTCCCAGGTATAATATAAGTTTCACTCACTATTGATTGATCGTGTTTATGCRTTAATGCTTCTAAGTAAT |
| 35 | 477 | 60_edited_HIrecoded.merged.removebadseq.renamed-249 | YTGCTAGTGCCCAGGGAAGCCTTGTTGATGAGATTAGGGAATACCTGAAGAAGGGTTCCTCTGCAAGTCAGCTGAAACCGTTTTTGGATTCACTCTCTGG[G/T]ACTTCCCAAGAAKTCATGGATGCCCTRTTYRAAGCTCTCTTTGGGGGTCTTGGAAAGGGGTTTGCTAAAGAGGTGACCAAGAAGAAGAACTTCCTTGCAG |
| 36 | 502 | 68_edited.merged.removebadseq.renamed-321 | GTTGATGTTTCAGTGGTTGACCTCACTGTGAGGCTGGAGAAGAAGGCTACCTATGAWCAAATTAAAGCTGCTATCAAGTAAGGGCAGTGAATMATTTTTT[G/T]TTATTTGCTTTTAGTGTCTGTTGTATGTATGGTAGGTTGTATTGCTATGTGGGTGATGAACGTTTAGTTGCTTTACCWTCTGACAATTGTGTGGTGAAAG |
| 37 | 504 | 70_edited.merged.removebadseq.renamed-119 | AGCTCCCAAAGGACACTTCACGCTCATTCCAAGCTCCAACTGGCTGGTCCGGGCGTTTCTGGGCCCGAACCGGGTGCAACTTCGACGGATCCGGAGCCGG[A/G]TCATGTCTCACTGGCGACTGTGGTTCAGGCCAAGTAGAATGCAACGGGTTCGGAGCCGCCCCACCCGCAACTCTCGCGGAGTTCACACTCGGYTCTGGCG |
| 38 | 511 | 73_edited_HIori.merged.removebadseq.renamed-166 | TGCAGAAGATGAGCTCATGAGAATCCTYSTTCTTGTCCGAAACTTCTTACCTGGATACCATCAAGCTATTAGTGGGGAATGGAATGTTGCAGCTATTGCT[C/T]ACAGGGCTTATGATCTTGAGGGAAAGACAGTTGGAACKRTTGGTGCTGGACGAATTGGCAAACTTTTACTTCAACGTTTGAAACCTTTTAACTGTAATCT |
| 39 | 515 | 73_edited_HIori.merged.removebadseq.renamed-615 | GATCTCAATTTTGTTAGATACTATGTTGTGATACAATTTGAATTTGTTTGCTGTAGAATGTCATATTTCTATGTTACTTATARGCTATCATTATGTTGGT[G/T]TATGCACTGAAATTCAAACTTATGAGAATTCTGATCTTGTGATTCANNGGGTTATTTGACAAANNNNNNNNNNNNCACATCTCTTGTTAGGCTATATTATT |
| 40 | 535 | 88-1_edited.merged.removebadseq.renamed-727 | TGGGACCATTCTCAGAGCAAACTCCCACATACCTGACTGGTGAATTCCCTGGTGACTATGGCTGGGACACTGCTGGTTTATCAGCAGACCCTGAGACCTT[C/T]GCAAAGAACCGTGAGCTCGAGGTGATCCACAGCCGATGGGCCATGCTTGGAGCTCTGGGCTGTGTCTTCCCTGAASTCCTTTCAAAGAATGGTGTCAAGT |

SNP_ID (a) following Ouayjan et al. (2018). SNPs discarded from the analyses are highlighted in yellow; Locus_Name_pos_SNP (b) corresponds to the locus name given by Lalagüe et al. (2014); seq SNP corresponds to sequences of the SNP flanking regions. The targeted SNP is indicated in brackets [ / ].
