## Supplementary material for "A chromosome-level genome assembly of the European Beech (*Fagus sylvatica*) reveals anomalies for organelle DNA integration, repeat content and distribution of SNPs": S6

**Supplementary file 6.** Genotyping data. List of 4127 SNPs and their associated linkage groups based on R_qtl (second raw) and JoinMap (third raw) analyses.

| SNP_ID | snp_B0029 | snp_B0043 | snp_B0050 | snp_B0051 | snp_B0067 | snp_B0087 | snp_B0099 | snp_B0106 | snp_B0160 | snp_B0193 | snp_B0203 | snp_B0205 | snp_B0249 | snp_B0265 | snp_B0302 | snp_B0330 | snp_B0349 | snp_B0376 | snp_B0381 | snp_B0392 | snp_B0417 | snp_B0418 |
|---|---|---|---|---|---|---|---|---|---|---|---|---|---|---|---|---|---|---|---|---|---|---|
| LG_Rqtl | 1 | 1 | 1 | 1 | 1 | 1 | 1 | 1 | 1 | 1 | 1 | 1 | 1 | 1 | 1 | 1 | 1 | 1 | 1 | 1 | 1 | 1 |
| LG_JoinMap | LG3 | -- | -- | -- | -- | -- | -- | -- | -- | -- | -- | -- | -- | -- | -- | -- | -- | -- | -- | -- | -- | -- |
| SSB002 | AA | AB | AA | AB | AA | AA | AA | AA | AB | AB | AB | AA | AB | AA | AB | AB | AA | AB | AA | AA | AB | AA |
| SSB003 | AB | AB | AB | AB | AB | AB | AA | AB | AB | AB | AB | AA | AB | AB | AB | AB | AB | AB | AB | AA | AB | AA |
| SSB007 | AA | AB | AB | AB | AB | AA | AA | AB | AA | AA | AB | AA | AA | AA | AB | AA | AB | AA | AB | AB | AA | AB |
| SSB010 | AA | AA | AA | AB | AA | AA | AB | AA | AA | AA | AA | AA | - | AA | AB | - | AA | - | - | AB | - | AB |
| SSB014 | AB | AB | AB | AA | AB | AB | AA | AA | AB | AA | AA | AB | AA | AB | AA | AB | AA | AB | AA | AA | AB | AB |
| SSB016 | AA | AA | - | - | - | AA | - | AB | AA | AB | AA | - | AA | AA | AA | - | - | AB | AA | AA | AA | - |
| SSB020 | AA | AB | AA | AA | AB | AB | AB | AB | AA | AA | AB | AA | AA | AB | AA | AA | AB | AB | AA | AB | AA | AA |
| SSB022 | AA | AB | AB | AB | AB | AB | AB | AA | AB | AB | AB | AB | AA | AB | AB | AB | AA | AA | AB | AA | AA | AA |
| SSB024 | AA | AA | AA | AA | AA | AB | AA | AA | AA | AB | AB | AA | AB | AB | AB | AA | AA | AB | AA | AA | AA | AB |
| SSB030 | AA | AA | AA | AA | AB | AB | AB | AA | AA | AB | AB | AA | AA | AA | AB | AA | AB | AB | AB | AB | AA | AA |
| SSB031 | AA | AA | AA | AB | AB | AB | AB | AB | AA | AB | AB | AA | AB | AB | AB | AB | AB | AB | AB | AB | AB | AA |
| SSB033 | AA | AB | AB | AB | AB | AB | AB | AA | AB | AB | AA | AA | AB | AB | AB | AB | AA | AB | AB | AB | AA | AA |
| SSB039 | AA | AA | AA | AA | AA | AB | AA | AA | AA | AB | AB | AA | AA | AA | AA | AA | AA | AB | AA | AA | AA | AB |
| SSB041 | AB | AB | AB | AA | AB | AB | AA | AA | AA | AB | AA | AA | AA | AA | AB | AA | AB | AB | AA | AB | AA | AA |
| SSB048 | AA | AA | AA | AB | AB | AA | AB | AA | AA | AA | AA | AA | AB | AA | AB | AA | AB | AB | AB | AB | AA | AB |
| SSB049 | AA | AB | AB | AB | AA | AA | AA | AA | AA | AB | AB | - | AA | AA | AB | AB | AA | AB | AB | AB | AA | AB |
| SSB052 | AB | AA | AB | AB | AA | AB | AA | AA | AB | AA | AB | AB | AB | AA | AB | AB | AB | AA | AB | AB | AA | AB |
| SSB070 | AB | AA | - | - | - | AA | - | AB | AB | AA | AA | - | - | AB | - | - | - | - | AB | AA | - | - |
| SSB075 | AB | AB | AB | AB | AB | AB | AB | AB | AB | AB | AB | AB | AB | AB | AB | AB | AB | AB | AB | AB | AB | AB |
| SSB077 | AA | AB | AB | AB | AB | AA | AB | AB | AB | AB | AB | AA | AA | AB | AB | AB | AA | AB | AB | AA | AA | AA |
| SSB080 | AA | AB | AB | AB | AB | AB | AB | AA | AB | AB | AB | AA | AA | AB | AB | AB | AB | AB | AB | AB | AB | AA |
| SSB084 | AB | AA | AA | AB | AB | AB | AB | AA | AA | AA | AB | AB | AB | AB | AB | AB | AB | AA | AB | AB | AA | AB |
| SSB086 | AA | AB | - | - | - | AA | AA | AB | AA | - | AB | AA | - | AA | - | - | - | - | - | AB | AA | - |
| SSB097 | AB | AB | AB | AB | AB | AB | AB | AA | AA | AA | AB | AB | AB | AB | AB | AB | AB | AA | AB | AB | AA | AB |
| SSB098 | AB | AA | AA | AB | AA | AB | AA | AA | AA | AA | AB | AB | AA | AA | AB | AA | AB | AA | AA | AA | AA | AA |
| SSB110 | AB | AB | AB | AB | AA | AB | AA | AB | AA | AB | AA | AB | AB | AA | AB | AB | AB | AA | AB | AA | AB | AA |
| SSB111 | AB | AB | AB | AB | AB | AA | AB | AB | AA | AB | AA | AA | AA | AA | AB | AA | AB | AB | AB | AA | AB | AA |
| SSB119 | AA | AB | AB | AB | AB | AA | AB | AA | AA | AB | AA | AA | AA | AA | AB | AB | AA | AA | AB | AA | AA | AA |
| SSB122 | AA | AA | AB | AB | - | AA | AB | AA | AA | AB | AB | - | - | AA | AB | - | AB | AB | AB | AB | - | AB |
| SSB123 | AB | AA | AB | - | - | AA | - | AB | AB | AA | AB | - | - | AB | - | - | - | - | - | AA | - | AB |
| SSB127 | AB | AB | AA | AB | AA | AA | AB | AA | AA | AA | AA | AA | AB | AB | AA | AA | AB | AA | AA | AA | AA | AB |
| SSB129 | AB | AB | AA | AB | AA | AA | AB | AA | AA | AA | AA | AB | AB | AB | AA | AB | AA | AA | AB | AA | AA | AB |
| SSB130 | AA | AB | - | AB | AB | AA | AB | AB | AB | AA | AA | - | - | AB | AA | AB | - | AA | AB | AA | AA | - |
| SSB132 | AB | AB | AB | AB | - | AB | AB | AB | AB | AA | AB | - | AB | AB | AB | - | - | AB | AB | AB | - | AB |
| SSB137 | AA | AA | AA | AB | - | AA | AA | AA | AA | AB | AB | AA | - | AB | AB | - | AA | - | AA | AA | - | - |
| SSB145 | AB | AA | AA | AA | AA | AB | AA | AB | AA | AB | AB | AA | AB | AB | AA | AA | AB | AB | AA | AB | - | AB |
| SSB148 | AB | AA | AA | AB | AB | AB | AA | AB | AA | AB | AB | AB | AB | AB | AA | AA | AB | AB | AB | AB | AB | AB |
| SSB149 | AA | AB | AA | AA | AB | AB | AB | AB | AA | AB | AB | AB | AA | AB | AA | AA | AB | AB | AA | AB | AB | AA |

| snp_B1351 | snp_B1391 | snp_B1426 | snp_B1427 | snp_B1434 | snp_B1449 | snp_B1454 | snp_B1467 | snp_B1499 | snp_B1520 | snp_B1571 | snp_B1592 | snp_B1597 | snp_B1602 | snp_B1610 | snp_B1615 | snp_B1621 |
|---|---|---|---|---|---|---|---|---|---|---|---|---|---|---|---|---|
| 1 | 1 | 1 | 1 | 1 | 1 | 1 | 1 | 1 | 1 | 1 | 1 | 1 | 1 | 1 | 1 | 1 |
| -- | -- | -- | -- | -- | LG3 | -- | -- | -- | LG3 | -- | -- | -- | -- | -- | LG3 | -- |
| AB | AA | AB | AA | AB | AB | AB | AB | AB | AB | AB | AB | AA | AA | AA | AB | AA |
| AB | AB | AA | AB | AB | AB | AB | AB | AB | AB | AB | AB | AB | AB | AB | AB | AB |
| AA | - | AA | AB | AA | AA | AA | AB | AB | AB | AB | AB | AB | AB | AB | AB | AB |
| AB | AB | - | - | AB | AA | AB | - | AB | AB | AA | - | AB | AB | - | AA | AA |
| AA | AB | AB | AA | AB | AB | AB | AB | AA | AA | AA | AB | AB | AA | AB | AA | AB |

| snp_B1655 | snp_B1675 | snp_B1682 | snp_B1693 | snp_B1710 | snp_B1717 | snp_B1723 | snp_B1736 | snp_B1755 | snp_B1758 | snp_B1769 | snp_B1770 | snp_B1771 | snp_B1773 | snp_B1790 | snp_B1794 | snp_B1818 | snp_B1828 | snp_B1835 | snp_B1853 |
|---|---|---|---|---|---|---|---|---|---|---|---|---|---|---|---|---|---|---|---|
| 1 | 1 | 1 | 1 | 1 | 1 | 1 | 1 | 1 | 1 | 1 | 1 | 1 | 1 | 1 | 1 | 1 | 1 | 1 | 1 |
| -- | -- | -- | -- | LG3 | -- | LG3 | LG3 | LG3 | -- | -- | -- | -- | LG3 | -- | -- | -- | -- | LG3 | LG3 |
| AB | AA | AA | AB | AA | AA | AA | AB | AB | AA | AA | AB | AA | AA | AB | AB | AA | AA | AA | AA |
| AB | AB | AB | AB | AB | AB | AB | AB | AB | AB | AB | AB | AB | AB | AB | AB | AB | AB | AB | AB |
| AB | AA | AB | AB | AA | AA | AB | AA | AA | AB | AB | AB | AB | AA | AA | AB | AA | AA | AA | AB |
| AB | AA | AB | AB | AA | AA | AB | AA | AA | - | AA | AA | AA | AB | AB | AB | AA | AA | AA | AA |
| AA | AB | AA | AA | AB | AB | AA | AB | AB | AB | AA | AB | AB | AA | AB | AB | AA | AA | AB | AB |

| snp_B1883 | snp_B1885 | snp_B1906 | snp_B1911 | snp_B1926 | snp_B1950 | snp_B1965 | snp_B1986 | snp_B1992 | snp_B2004 | snp_B2009 | snp_B2025 | snp_B2031 | snp_B2034 | snp_B2039 | snp_B2049 | snp_B2061 | snp_B2066 | snp_B2085 |
|---|---|---|---|---|---|---|---|---|---|---|---|---|---|---|---|---|---|---|
| 1 | 1 | 1 | 1 | 1 | 1 | 1 | 1 | 1 | 1 | 1 | 1 | 1 | 1 | 1 | 1 | 1 | 1 | 1 |
| -- | -- | -- | -- | -- | -- | -- | -- | -- | -- | -- | LG3 | -- | -- | -- | -- | -- | -- | -- |
| AA | AA | AB | AA | AB | AA | AB | AB | AA | AB | AB | AB | AA | AA | AB | AB | AA | AA | AA |
| AB | AA | AB | AB | AB | AB | AB | AB | AB | AB | AB | AB | AB | AB | AB | AB | AB | AB | AB |
| AB | AA | AB | AA | AB | AA | AB | AB | AB | AB | AB | AB | AB | AA | AA | AB | AB | AA | AA |
| AB | AA | AA | AA | AA | AA | AB | AB | AA | AB | AB | AB | AB | AB | AA | AB | AA | AB | AA |
| AA | AB | AA | AA | AA | AB | AB | AA | AA | AA | AA | AA | AA | AB | AA | AA | AA | AB | AB |

| snp_B2086 | snp_B2100 | snp_B2103 | snp_B2112 | snp_B2113 | snp_B2137 | snp_B2145 | snp_B2163 | snp_B2173 | snp_B2176 | snp_B2181 | snp_B2220 | snp_B2225 | snp_B2255 | snp_B2258 | snp_B2275 | snp_B2282 | snp_B2371 | snp_B2377 | snp_B2383 | snp_B2400 |
|---|---|---|---|---|---|---|---|---|---|---|---|---|---|---|---|---|---|---|---|---|
| 1 | 1 | 1 | 1 | 1 | 1 | 1 | 1 | 1 | 1 | 1 | 1 | 1 | 1 | 1 | 1 | 1 | 1 | 1 | 1 | 1 |
| -- | -- | -- | -- | -- | -- | -- | -- | -- | -- | -- | -- | LG3 | LG3 | -- | -- | LG3 | -- | -- | -- | -- |
| AB | AA | AA | AB | AA | AA | AA | AB | AA | AA | AA | AB | AA | AB | AA | AA | AA | AA | AB | AB | AB |
| AB | AB | AB | AB | AB | AB | AB | AB | AB | AB | AB | AB | AB | AB | AA | AB | AB | AB | AB | AB | AB |
| AA | AB | AA | AB | AB | AB | AB | AB | AA | AB | AB | AB | AA | AA | AB | AA | AB | AB | AB | AA | AB |
| AA | AA | AB | AA | AA | AB | AB | AA | AB | AB | AA | AB | AA | AA | AB | AA | - | AA | AA | AB | AB |
| AA | AA | AB | AA | AA | AA | AA | AA | AA | AA | AB | AA | AA | AB | AA | AB | AB | AB | AB | AB | AA |

| snp_B2407 | snp_B2445 | snp_B2510 | snp_B2516 | snp_B2531 | snp_B2533 | snp_B2541 | snp_B2547 | snp_B2549 | snp_B2558 | snp_B2559 | snp_B2572 | snp_B2601 | snp_B2602 | snp_B2613 | snp_B2621 | snp_B2622 | snp_B2623 | snp_B2625 | snp_B2654 | snp_B2670 | snp_B2718 | snp_B2772 | snp_B2776 | snp_B2793 | snp_B2807 | snp_B2808 | snp_B2829 | snp_B2838 | snp_B2840 | snp_B2844 | snp_B2849 | snp_B2854 | snp_B2876 | snp_B2890 | snp_B2907 | snp_B2914 | snp_B2918 | snp_B2923 | snp_B2927 | snp_B2932 | snp_B2950 | snp_B2984 | snp_B2992 | snp_B2999 | snp_B3017 | snp_B3044 | snp_B3058 | snp_B3071 | snp_B3085 | snp_B3086 | snp_B3092 | snp_B3098 | snp_B3111 | snp_B3159 | snp_B3162 | snp_B3178 | snp_B3179 | snp_B3181 | snp_B3202 | snp_B3206 | snp_B3244 | snp_B3306 | snp_B3319 | snp_B3330 | snp_B3380 | snp_B3389 | snp_B3393 | snp_B3447 | snp_B3470 | snp_B3476 | snp_B3499 | snp_B3503 | snp_B3522 | snp_B3528 | snp_B3536 | snp_B3542 |
|---|---|---|---|---|---|---|---|---|---|---|---|---|---|---|---|---|---|---|---|---|---|---|---|---|---|---|---|---|---|---|---|---|---|---|---|---|---|---|---|---|---|---|---|---|---|---|---|---|---|---|---|---|---|---|---|---|---|---|---|---|---|---|---|---|---|---|---|---|---|---|---|---|---|---|---|---|
| 1 | 1 | 1 | 1 | 1 | 1 | 1 | 1 | 1 | 1 | 1 | 1 | 1 | 1 | 1 | 1 | 1 | 1 | 1 | 1 | 1 | 1 | 1 | 1 | 1 | 1 | 1 | 1 | 1 | 1 | 1 | 1 | 1 | 1 | 1 | 1 | 1 | 1 | 1 | 1 | 1 | 1 | 1 | 1 | 1 | 1 | 1 | 1 | 1 | 1 | 1 | 1 | 1 | 1 | 1 | 1 | 1 | 1 | 1 | 1 | 1 | 1 | 1 | 1 | 1 | 1 | 1 | 1 | 1 | 1 | 1 | 1 | 1 | 1 | 1 | 1 | 1 |
| -- | -- | -- | -- | -- | -- | -- | -- | -- | -- | LG3 | -- | -- | -- | -- | -- | LG3 | -- | -- | LG3 | -- | -- | -- | -- | LG3 | -- | LG3 | -- | -- | -- | -- | -- | -- | -- | -- | LG3 | -- | -- | LG3 | -- | -- | -- | -- | -- | -- | -- | -- | -- | -- | -- | -- | -- | -- | LG3 | -- | -- | -- | -- | -- | -- | -- | -- | -- | -- | -- | -- | -- | -- | -- | LG3 | -- | -- | LG3 | -- | LG3 | -- | -- |

| snp_B3546 | snp_B3549 | snp_B3564 | snp_B3584 | snp_B3610 | snp_B3651 | snp_B3658 | snp_B3660 | snp_B3663 | snp_B3677 | snp_B3680 | snp_B3686 | snp_B3689 | snp_B3697 | snp_B3709 | snp_B3710 | snp_B3732 | snp_B3736 | snp_B3740 |
|---|---|---|---|---|---|---|---|---|---|---|---|---|---|---|---|---|---|---|
| 1 | 1 | 1 | 1 | 1 | 1 | 1 | 1 | 1 | 1 | 1 | 1 | 1 | 1 | 1 | 1 | 1 | 1 | 1 |
| -- | -- | -- | LG3 | LG3 | LG3 | -- | -- | -- | -- | -- | -- | -- | -- | -- | -- | -- | -- | -- |
| AB | AA | AA | AA | AA | AB | AB | AA | AB | AA | AB | AB | AA | AA | AB | AA | AB | AB | AB |
| AB | AA | AB | AB | AA | AB | AB | AA | AB | AB | AB | AB | AB | AB | AB | AB | AB | AB | AB |

| snp_B3747 | snp_B3748 | snp_B3758 | snp_B3784 | snp_B3789 | snp_B3808 | snp_B3811 | snp_B3814 | snp_B3825 | snp_B3829 | snp_B3838 | snp_B3846 | snp_B3858 | snp_B3864 | snp_B3876 | snp_B3888 | snp_B3904 | snp_B3926 | snp_B3927 | snp_B3937 | snp_B3940 | snp_B3992 |
|---|---|---|---|---|---|---|---|---|---|---|---|---|---|---|---|---|---|---|---|---|---|
| 1 | 1 | 1 | 1 | 1 | 1 | 1 | 1 | 1 | 1 | 1 | 1 | 1 | 1 | 1 | 1 | 1 | 1 | 1 | 1 | 1 | 1 |
| -- | -- | -- | -- | -- | -- | LG3 | -- | -- | -- | -- | -- | LG3 | LG3 | -- | -- | -- | -- | -- | -- | LG3 | -- |
| AB | AA | AA | AA | AB | AA | AB | AB | AB | AB | AA | AA | AB | AB | AB | AA | AB | AB | AB | AA | AA | AA |
| AB | AB | AA | AB | AB | AB | AB | AB | AB | AB | AB | AA | AB | AB | AB | AB | AB | AB | AB | AB | AB | AB |

| snp_B4007 | snp_B4011 | snp_B4017 | snp_B4025 | snp_B4026 | snp_B4036 | snp_B4055 | snp_B4060 | snp_B4066 | snp_B4073 | snp_B4080 | snp_B4105 | snp_B4108 | snp_B4111 | snp_B4128 | snp_B4133 | snp_B4152 | snp_B4162 | snp_B4172 | snp_B4183 | snp_B4215 |
|---|---|---|---|---|---|---|---|---|---|---|---|---|---|---|---|---|---|---|---|---|
| 1 | 1 | 1 | 1 | 1 | 1 | 1 | 1 | 1 | 1 | 1 | 1 | 1 | 1 | 1 | 1 | 1 | 1 | 1 | 1 | 1 |
| LG3 | -- | -- | -- | -- | -- | -- | -- | -- | -- | -- | -- | -- | -- | -- | -- | -- | -- | -- | -- | -- |
| AB | AA | AB | AB | AB | AA | AA | AA | AB | AA | AA | AB | AB | AB | AA | AB | AA | AA | AA | AA | AB |
| AB | AB | AB | AB | AB | AB | AB | AB | AB | AA | AB | AB | AB | AB | AB | AB | AB | AB | AB | AB | AB |

| snp_B4251 | snp_B4287 | snp_B4296 | snp_B4298 | snp_B4301 | snp_B4322 | snp_B4378 | snp_B4410 | snp_B4432 | snp_B4436 | snp_B4457 | snp_B4464 | snp_B4489 | snp_B4493 | snp_B4500 |
|---|---|---|---|---|---|---|---|---|---|---|---|---|---|---|
| 1 | 1 | 1 | 1 | 1 | 1 | 1 | 1 | 1 | 1 | 1 | 1 | 1 | 1 | 1 |
| -- | -- | -- | -- | -- | -- | LG3 | -- | -- | -- | -- | -- | -- | -- | -- |
| AB | AB | AB | AB | AB | AA | AB | AB | AA | AB | AA | AA | AA | AA | AB |
| AB | AB | AB | AB | AB | AB | AB | AB | AB | AB | AB | AB | AB | AB | AB |

| snp_B4512 | snp_B4520 | snp_B4530 | snp_B4538 | snp_B4548 | snp_B4563 | snp_B4568 | snp_B4577 | snp_B4583 | snp_B4590 | snp_B4602 | snp_B4609 | snp_B4624 | snp_B4646 | snp_B4653 | snp_B4671 | snp_B4683 | snp_B4697 | snp_B4706 | snp_B4732 |
|---|---|---|---|---|---|---|---|---|---|---|---|---|---|---|---|---|---|---|---|
| 1 | 1 | 1 | 1 | 1 | 1 | 1 | 1 | 1 | 1 | 1 | 1 | 1 | 1 | 1 | 1 | 1 | 1 | 1 | 1 |
| -- | -- | LG3 | LG3 | -- | -- | -- | -- | -- | -- | -- | -- | -- | -- | -- | -- | -- | -- | -- | -- |
| AA | AB | AB | AA | AA | AB | AB | AA | AB | AA | AA | AB | AA | AA | AA | AA | AB | AB | AA | AB |
| AB | AB | AB | AB | AA | AB | AB | AB | AB | AB | AB | AB | AB | AB | AB | AB | AB | AB | AB | AB |
| AB | AA | AB | AA | AA | AB | AB | AA | AB | AA | AA | AB | AB | AB | AB | AB | AB | AA | AB | AB |
| AA | AA | AA | AA | AA | AA | AB | AB | AA | AB | AB | - | AA | AB | AA | AA | AA | AA | AB | AA |
| AB | AA | AA | AB | AA | AB | AB | AA | AB | AA | AA | AA | AB | AA | AA | AA | AA | AB | AA | AB |
| AB | AB | AA | AA | - | - | AB | AA | AA | AA | AA | - | AA | AA | AB | AA | AA | AA | AA | AA |
| AB | AB | AA | AA | AA | AA | AB | AB | AA | AB | AA | AB | AB | AB | AB | AB | AB | AA | AA | AB |
| AA | AB | AB | AB | AB | AB | AA | AA | AB | AA | AA | AA | AA | AB | AA | AA | AB | AB | AA | AA |
| AA | AB | AB | AA | AA | AB | AB | AB | AB | AA | AA | AB | AA | AA | AA | AA | AB | AA | AB | AB |
| AA | AB | AB | AA | AA | AA | AB | AB | AA | AB | AB | AB | AA | AB | AA | AA | AB | AA | AA | AA |
| AA | AB | AB | AA | AA | AA | AB | AB | AA | AA | AB | AB | AB | AA | AA | AA | AB | AA | AA | AB |
| AA | AB | AB | AA | AB | AA | AB | AB | AB | AA | AA | AA | AA | AB | AA | AA | AB | AB | AA | AA |
| AA | AB | AB | AB | AA | AB | AB | AA | AA | AA | AB | AB | AA | AA | AA | AA | AA | AA | AB | AA |
| AB | AB | AA | AA | AB | AA | AB | AB | AB | AA | AB | AB | AB | AB | AA | AB | AB | AA | AA | AB |
| AA | AA | AA | AA | AA | AA | AB | AB | AA | AB | AA | AA | AB | AB | AA | AB | AB | AA | AB | AA |
| AB | AB | AB | AA | AB | AB | AB | AB | AA | AB | AA | AB | AA | AB | AB | AB | AB | AA | AA | AA |
| AA | AA | AB | AB | AB | AB | AA | AA | AB | AA | AB | AB | AB | AA | AA | AA | AB | AB | AA | AB |
| AB | AA | AA | AB | AB | - | AA | AA | - | - | AA | AA | AB | AA | - | AA | AB | AB | AA | - |
| AB | AB | AB | AB | AA | AB | AB | AB | AB | AB | AB | AB | AB | AB | AB | AB | AB | AB | AB | AB |
| AB | AB | AA | AA | AB | AA | AB | AA | AB | AB | AA | - | AA | AB | AB | AB | AA | AB | AB | AA |
| AA | AA | AB | AA | AA | AA | AB | AB | AB | AB | AB | AB | AB | AB | AA | AB | AB | AB | AB | AA |
| AA | AB | AB | AB | AA | AB | AA | AB | AA | AA | AB | AB | AB | AA | AA | AB | AB | AA | AA | AB |
| AA | AB | AB | AA | AB | - | AB | - | AA | - | - | - | AA | AB | - | AA | AB | AA | AA | - |
| AA | AA | AB | AB | AB | AB | AA | AB | AB | AB | AB | AB | AB | AA | AA | AB | AB | AB | AA | AB |
| AA | AA | AB | AA | AA | AA | AA | AB | AA | AA | AB | AA | AB | AA | AA | AB | AA | AA | AB | AB |
| AA | AB | AA | AB | AB | AA | AA | AB | AA | AA | AB | AA | AB | AA | AA | AB | AA | AA | AB | AB |
| AB | AB | AA | AA | AB | AA | AA | AA | AA | AB | AB | AB | AB | AB | AB | AB | AA | AB | AB | AB |
| AB | AB | AA | AA | AA | AA | AB | AA | AA | AB | AA | AB | AA | AB | AB | AB | AB | AB | AA | AB |
| AA | AB | AB | AB | AB | AA | AB | AB | AA | - | AB | AB | AB | AB | AA | AB | AB | AA | AA | AB |
| AB | AA | AA | AB | AB | - | AA | - | AB | AB | AB | - | AB | AA | - | AB | AA | AB | AB | - |
| AA | AA | AB | AB | AB | AB | AA | AA | AA | AB | AB | AA | AB | AB | AA | AA | AB | AA | AA | AB |
| AA | AA | AB | AB | AA | AB | AA | AA | AB | AB | AB | AA | AA | AB | AA | AA | AB | AA | AA | AA |
| AB | AA | AA | AA | AB | AA | AB | AA | AB | AB | - | - | AA | AB | AB | AA | AA | AB | AB | AA |
| AA | AA | AB | AB | AB | AB | AB | AB | AB | - | - | - | AB | AB | AB | AB | AB | AB | AB | AB |
| AA | AB | AA | AA | - | - | AA | AA | AA | AA | AA | AA | AB | AA | AA | AA | AA | AA | AB | - |
| AB | AB | AB | AB | AB | AB | AB | AB | AA | AA | - | AB | AB | AA | AB | AB | AB | AA | AB | - |
| AB | AB | AB | AB | AB | AB | AB | AB | AA | AA | AB | AB | AB | AA | AB | AB | AB | AA | AB | AB |
| AB | AB | AB | AA | AA | AB | AB | AB | AA | AA | AA | AB | AB | AA | AB | AB | AB | AA | AB | AB |
| AB | AB | AB | AB | AB | AB | AB | AB | AA | AA | AB | AB | AB | AA | AB | AB | AB | AA | AB | AB |
| AB | AB | AA | AA | AB | AB | AB | AA | AB | AA | AB | AB | AA | AA | AA | AB | AB | AB | AA | - |
| AA | AB | AA | AA | AB | AB | AA | AA | AB | AB | AB | AA | AB | AB | AA | AA | AA | AB | AA | AA |
| AA | AA | AB | AB | AB | AB | AA | AA | AB | AB | AB | AB | AB | AB | AA | AA | AB | AA | AB | AA |
| AB | AA | AA | AA | AB | AA | AB | AB | AA | AA | AA | AA | AB | AB | AB | AB | AA | AA | AA | AB |
| AA | AB | AB | AA | AA | - | AB | AB | AA | - | AB | - | AB | AB | AA | AB | AB | AA | AB | AB |
| AB | AB | AB | AB | AA | AB | AB | AA | AA | AB | AB | AB | AB | AB | AA | AB | AB | AA | AB | AA |
| AA | AA | AA | AB | AA | AA | AA | AA | AA | AA | AA | AB | AB | AB | AA | AB | AB | AA | AB | AB |
| AA | AA | AB | AB | AB | AB | AA | AB | AA | AB | AB | AB | AB | AB | AA | AB | AB | AA | AB | AB |
| AA | AA | AB | AA | AA | AA | AB | AB | AB | AA | AB | AA | AA | AB | AA | AB | AB | AA | AB | AA |
| AB | AB | AB | AB | AB | AB | AB | AB | AA | AB | AB | AB | AB | AB | AB | AB | AB | AA | AB | AB |
| AB | AB | AB | AA | AB | AB | AA | AB | AA | AB | AB | AB | AB | AB | AB | AB | AA | AB | AB | AB |
| AB | AA | AB | AB | AA | AB | AA | AA | AB | AA | AA | AA | AB | AA | AB | AA | AA | AA | AB | AB |
| AA | AA | AA | AB | - | AB | AA | AA | AB | AA | AA | AB | AA | AA | AB | AA | AA | AA | AB | AA |
| AA | AA | AA | AA | AB | AA | AA | AB | AA | AB | AB | AA | AA | AB | AA | AB | AA | AB | AA | AA |

[illegible]

| snp_B5746 | snp_B5755 | snp_B5785 | snp_B5787 | snp_B5803 | snp_B5833 | snp_B5842 | snp_B5844 | snp_B5847 | snp_B5867 | snp_B5876 | snp_B5880 | snp_B5881 | snp_B5882 | snp_B5941 | snp_B5943 | snp_B5945 | snp_B5962 | snp_B5981 | snp_B6015 | snp_B6027 |
|---|---|---|---|---|---|---|---|---|---|---|---|---|---|---|---|---|---|---|---|---|
| 1 | 1 | 1 | 1 | 1 | 1 | 1 | 1 | 1 | 1 | 1 | 1 | 1 | 1 | 1 | 1 | 1 | 1 | 1 | 1 | 1 |
| -- | -- | LG3 | -- | -- | -- | -- | -- | -- | -- | LG3 | -- | -- | -- | LG3 | -- | -- | -- | -- | -- | -- |
| AB | AB | AB | AA | AB | AB | AB | AB | AB | AB | AA | AA | AA | AB | AB | AA | AA | AB | AA | AA | AB |
| AB | AB | AB | AA | AB | AB | AB | AB | AB | AB | AB | AA | AB | AB | AB | AB | AA | AB | AA | AA | AB |
| AB | AA | AB | AA | AA | AB | AA | AA | AA | AB | AA | AB | AB | AA | AA | AA | AA | AB | AA | AA | AB |
| AA | AA | AB | AB | AB | - | AA | AA | AB | AB | AA | AB | AA | AA | AA | AB | AB | AB | AA | AA | - |
| AA | AB | AA | - | AB | AB | AB | AB | AA | AA | AB | AA | AA | AB | AB | AA | AB | AB | AB | AA | AA |
| - | AA | - | AA | AB | AA | AA | AA | AA | AA | AA | AA | - | - | AA | AA | AA | AB | AA | AB | - |
| AB | AB | AB | AB | AB | AA | AA | AA | AB | AB | AA | AA | AA | AA | AA | AB | AA | AB | AA | AA | AA |
| AA | AA | AA | AB | AA | AA | AB | AB | AA | AA | AA | AA | AA | AB | AB | AB | AA | AA | AB | AA | AB |
| AA | AB | AB | AA | AB | AB | AA | AA | AB | AB | AA | AB | AA | AA | AA | AA | AA | AB | AA | AA | AB |
| AA | AB | AB | AB | AB | AA | AB | AA | AB | AA | AA | AB | AA | AA | AA | AB | AA | AA | AB | AA | AA |
| AA | AB | AB | AB | AB | AA | AA | AA | AB | AB | AA | AB | AA | AB | AA | AB | AA | AA | AA | AA | AB |
| AB | AB | AB | AB | AB | AA | AA | AB | AB | AB | AA | AB | AA | AB | AA | AB | AA | AB | AA | AB | AB |
| AA | AB | AB | AA | AB | AA | AB | AA | AA | AA | AA | AA | AA | AA | AB | AA | AB | AB | AA | AA | AA |
| AA | AB | AB | AB | AB | AB | AA | AB | AB | AB | AA | AB | AA | AA | AA | AB | AA | AB | AB | AA | AA |
| AB | AA | AB | AB | AB | AA | AA | AA | AB | AB | AA | AA | AA | AA | AA | AB | AA | AB | AA | AB | AB |
| AB | AB | AB | AB | AA | AA | AA | AA | AB | AA | AA | AA | AA | AB | AA | AB | AA | AB | AA | AA | AB |
| AA | AA | AB | AA | AA | AB | AB | AB | AA | AA | AB | AB | AA | AB | AB | AA | AB | AA | AB | AA | AA |
| - | AA | - | - | AA | AA | AB | AA | AA | AA | AB | AA | AB | - | AB | - | AB | AB | - | AB | - |
| AB | AB | AB | AB | AB | AB | AB | AB | AA | AB | AB | AB | AB | AB | AB | AB | AB | AB | AA | AB | AB |
| AB | AB | AA | AB | AB | AA | AA | AB | AB | AA | AA | AA | AB | AB | AA | AB | AB | AB | AA | AB | AB |
| AA | AB | AB | AB | AA | AA | AB | AB | AB | AA | AA | AB | AA | AB | AA | AB | AA | AA | AA | AA | AB |
| AA | AB | AB | AA | AA | AB | AB | AB | AB | AB | AB | AB | AA | AB | AB | AA | AB | AA | AA | AA | AB |
| AB | - | - | AA | AB | - | - | AB | AB | AB | AA | AB | AA | - | AA | AB | AB | AB | AA | AB | - |
| AA | AB | AB | AA | AA | AB | AB | AB | AB | AB | AB | AB | AA | AB | AB | AB | AB | AA | AB | AA | AA |
| AA | AB | AB | AA | AA | AB | AA | AA | AB | AA | AB | AA | AA | AA | AA | AA | AB | AA | AA | AA | AB |
| AA | AA | AA | AB | AB | AB | AB | AA | AA | AB | AB | AB | AB | AB | AB | AB | AB | AA | AA | AB | AB |
| AB | AB | AB | AB | AB | AB | AA | AA | AB | AB | AA | AB | AB | AA | AA | AB | AA | AB | AA | AB | AB |
| AB | AA | AB | AB | AB | AB | AA | AA | AB | AB | AA | AB | AA | AA | AA | AB | AA | AB | AA | AB | AB |
| AB | AB | AB | - | AB | AB | AA | AA | AB | AB | AA | AB | AA | AA | AA | AA | AA | AB | AA | AB | - |
| AA | AB | AA | - | AA | AB | - | AA | AA | AB | AB | AA | AB | AB | AB | - | AB | - | AA | AA | - |
| AA | AA | AB | AB | AA | AB | AB | AA | AA | AB | AB | AB | AA | AA | AB | AB | AB | AA | AB | AA | AA |
| AA | AA | AA | AB | AA | AB | AB | AB | AA | AA | AB | AB | AA | AB | AB | AB | AB | AA | AB | AB | AA |
| AB | AA | AA | AA | AB | AA | AA | AB | AA | AA | AA | AA | AB | AB | AA | AB | AA | AB | AA | AA | - |
| AB | AA | - | - | AB | AB | AB | AB | AB | AB | AB | AB | AA | AB | AB | AB | AB | AB | AB | AB | - |
| - | AB | AA | AA | AA | AA | - | AA | AB | AA | AA | - | AA | - | AA | AA | AA | AA | AB | AB | - |
| AB | AB | AB | AA | AB | AB | AB | AA | AA | AB | AB | AB | AB | AA | AB | AA | AB | AB | AB | AB | AB |
| AB | AA | AB | AA | AB | AB | AA | AA | AB | AA | AB | AA | AB | AA | AB | AA | AB | AB | AB | AA | AB |
| AB | AA | AB | AA | AB | AA | AA | AA | AA | AB | AA | AA | AA | AA | AA | AB | AA | AB | AA | AA | AB |
| AB | AB | AB | AA | AB | AB | AA | AB | AA | AB | AB | AB | AB | AA | AA | AB | AA | AB | AB | AB | AB |
| AA | AB | AB | AA | AB | AA | AB | AB | AA | AA | AA | AB | AA | AB | AB | - | AA | AB | AB | AA | - |
| AA | AB | AA | AB | AA | AA | AB | AB | AA | AA | AA | AA | AA | AB | AA | AB | AA | AA | AA | AA | AA |
| AA | AA | AB | AA | AA | AB | AB | AB | AB | AB | AB | AB | AB | AB | AB | AA | AA | AB | AA | AB | AB |
| AB | AB | AA | AB | AB | AA | AA | AB | AB | AB | AA | AA | AA | AA | AB | AB | AB | AB | AB | AA | AA |
| AA | AB | AB | AB | AB | AB | AA | AA | AA | AB | AA | AB | AA | AB | AB | AB | AB | AA | AA | AB | AB |
| AA | AA | AA | AB | AA | AA | AB | AB | AA | AA | AA | AA | AA | AB | AA | AB | AA | AB | AA | AA | - |
| AA | AB | AB | AB | AA | AB | AA | AA | AB | AB | AB | AB | AA | AA | AB | AB | AB | AA | AB | AB | AA |
| AA | AA | AB | AB | AA | AA | AB | AA | AB | AB | AA | AA | AB | AA | AA | AB | AB | AA | AB | AA | AA |
| AA | AB | AB | AB | AB | AA | AB | AB | AB | AA | AA | AB | AA | AB | AA | AB | AB | AA | AA | AA | AB |
| AB | AA | AB | AB | AB | AA | AA | AA | AA | AB | AA | AA | AA | AA | AA | AB | AA | AB | AA | AA | - |
| AB | AB | AB | AB | AB | AB | AA | AA | AB | AB | AA | AA | AB | AB | AA | AA | AB | AB | AB | AB | AB |
| AB | AA | AA | AA | AA | AB | AB | AB | AB | AA | AA | AA | AB | AB | AB | AA | AA | AB | AB | AB | AB |
| AB | AA | AA | - | AA | AA | AB | AB | AB | AA | AB | AA | AB | AB | AB | - | AA | AB | AA | AA | - |
| AA | AA | AA | AB | AA | AB | AA | AA | AB | AB | AB | AA | AB | AA | AA | AB | AB | AA | AA | AB | AB |
| AA | AB | AA | AB | AA | AA | AA | AB | AB | AB | AB | AA | AA | AB | AA | AB | AB | AA | AA | AA | AB |
| AB | AB | AB | AB | AB | AA | AA | AA | - | AB | AA | AA | AA | AB | AA | AB | AA | AB | AB | AA | AA |
| AA | AB | AA | AB | AA | AA | AB | AA | AB | AB | AB | AB | AA | AB | AA | AB | AA | AA | AA | AB | AB |
| AA | AB | AA | AB | AA | AA | AA | AA | AB | AB | AA | AA | AA | AB | AA | AB | AA | AA | AA | AA | AB |
| AA | AB | AB | AB | AA | AB | AA | AA | AA | AA | AB | AB | AA | AA | AA | AB | AB | AA | AA | AA | AA |
| AB | - | AB | AA | AA | AA | AB | AB | AB | AA | AB | AA | AB | AB | AA | AA | AA | AB | AB | AA | AB |
| AB | AA | AB | AA | AA | AB | AA | AA | AB | AA | AA | AB | AA | AA | AA | AB | AB | AA | AB | AB | AA |
| AB | AB | AB | - | AB | AB | AB | AA | AB | AB | AB | AB | AB | AA | AB | - | AB | AB | AA | AB | AB |
| AA | AA | AA | AA | AB | AB | AB | AB | AA | AB | AB | AB | AB | AA | AB | AA | AB | AA | AA | AB | AB |
| AA | AB | AA | AB | AB | AB | AB | AA | AB | AA | AB | AB | AB | AA | AB | AA | AB | AB | AA | AA | AB |
| AB | AB | - | AB | AB | AB | AB | AA | AB | AB | AB | AB | AB | AA | AB | - | AB | AB | AB | AB | - |
| AA | AB | AA | AA | AB | AA | AA | AB | AB | AA | AA | AA | AB | AB | AB | AA | AA | AA | AB | AB | AB |
| AB | AB | AA | AB | AB | AA | AA | AA | AB | AA | AA | AA | AB | AB | AA | AB | AA | AB | AA | AB | AB |
| AB | AA | AA | AB | AB | AA | AA | AB | AA | AA | AB | AB | AB | AB | AA | AA | AB | AB | AA | AB | AB |
| AB | AA | AB | AB | AA | AA | AA | AB | AB | AA | AA | AA | AB | AB | AB | AB | AA | AA | AA | AB | AB |
| AA | AA | AA | AA | AB | AA | AB | AB | AA | AA | AA | AB | AB | AA | AB | AA | AA | AA | AB | AA | AA |
| AB | AB | AA | AB | AA | AB | AA | AB | AB | AA | AA | AB | AB | AB | AA | AB | AA | AA | AB | AB | AB |
| AA | AA | AB | AA | AA | AB | AB | AA | AA | AB | AA | AA | AA | AB | AB | AA | AB | AA | AB | AB | AA |
| AB | AB | AA | AA | AB | AA | AA | AA | AA | AB | AA | AA | AB | AA | AB | AB | AA | AA | AA | AA | AA |
| AB | AB | AB | AB | AA | AB | AB | AB | AB | AB | AA | AB | AA | AB | AB | AB | AA | AB | AB | AB | AB |
| AB | AA | AA | AB | AB | AB | AB | AA | AB | AA | AB | AA | AB | AA | AB | AB | AB | AB | AA | AB | AB |
| AB | AA | AB | AB | AB | AA | AB | AB | AB | AB | AA | AA | AA | AA | AA | AA | AA | AA | AA | AA | AA |
| AA | AB | AA | AB | AB | AA | AA | AB | AA | AA | AB | AB | AB | AA | AB | AA | AB | AB | AB | AA | AA |
| AB | AA | AA | AB | AB | AA | AA | AA | AA | AA | AA | AA | AB | AA | AA | AB | AB | AB | AA | AB | AA |
| AA | AB | AA | AA | AA | AA | AB | AB | AA | AA | AB | AB | AA | AB | AB | AA | AB | AA | AB | AB | AB |
| AB | AB | AB | AB | AB | AA | AA | AA | AA | AA | AB | AA | AB | AA | AB | AB | AB | AB | AA | AB | AB |
| AA | AA | AA | AB | AA | AA | AA | AB | AB | AA | AA | AA | AB | AB | AB | AA | AA | AA | AA | AA | AB |
| [illegible] | [illegible] | [illegible] | [illegible] | [illegible] | [illegible] | [illegible] | [illegible] | [illegible] | [illegible] | [illegible] | [illegible] | [illegible] | [illegible] | [illegible] | [illegible] | [illegible] | [illegible] | [illegible] | [illegible] | [illegible] |

| snp_B6151 | snp_B6153 | snp_B6166 | snp_B6208 | snp_B6243 | snp_B6360 | snp_B0026 | snp_B0042 | snp_B0055 | snp_B0066 | snp_B0071 | snp_B0074 | snp_B0104 | snp_B0135 | snp_B0157 | snp_B0183 | snp_B0207 | snp_B0208 | snp_B0258 |
|---|---|---|---|---|---|---|---|---|---|---|---|---|---|---|---|---|---|---|
| 1 | 1 | 1 | 1 | 1 | 1 | 2 | 2 | 2 | 2 | 2 | 2 | 2 | 2 | 2 | 2 | 2 | 2 | 2 |
| -- | -- | -- | -- | -- | -- | -- | -- | -- | -- | -- | -- | LG1 | -- | -- | LG1 | -- | -- | -- |
| [illegible] | [illegible] | [illegible] | [illegible] | [illegible] | [illegible] | [illegible] | [illegible] | [illegible] | [illegible] | [illegible] | [illegible] | [illegible] | [illegible] | [illegible] | [illegible] | [illegible] | [illegible] | [illegible] |

| snp_B0270 | snp_B0272 | snp_B0274 | snp_B0280 | snp_B0309 | snp_B0315 | snp_B0332 | snp_B0341 | snp_B0366 | snp_B0375 | snp_B0379 | snp_B0395 | snp_B0400 | snp_B0421 | snp_B0451 | snp_B0460 | snp_B0472 | snp_B0479 | snp_B0496 |
|---|---|---|---|---|---|---|---|---|---|---|---|---|---|---|---|---|---|---|
| 2 | 2 | 2 | 2 | 2 | 2 | 2 | 2 | 2 | 2 | 2 | 2 | 2 | 2 | 2 | 2 | 2 | 2 | 2 |
| -- | -- | -- | -- | -- | -- | -- | -- | -- | -- | -- | -- | -- | -- | -- | -- | -- | -- | -- |
| [illegible] | [illegible] | [illegible] | [illegible] | [illegible] | [illegible] | [illegible] | [illegible] | [illegible] | [illegible] | [illegible] | [illegible] | [illegible] | [illegible] | [illegible] | [illegible] | [illegible] | [illegible] | [illegible] |

| snp_B0507 | snp_B0534 | snp_B0549 | snp_B0564 | snp_B0594 | snp_B0608 | snp_B0639 | snp_B0642 | snp_B0652 | snp_B0660 | snp_B0681 | snp_B0703 | snp_B0716 | snp_B0775 | snp_B0780 | snp_B0795 | snp_B0807 | snp_B0825 |
|---|---|---|---|---|---|---|---|---|---|---|---|---|---|---|---|---|---|
| 2 | 2 | 2 | 2 | 2 | 2 | 2 | 2 | 2 | 2 | 2 | 2 | 2 | 2 | 2 | 2 | 2 | 2 |
| -- | -- | -- | -- | -- | -- | -- | -- | -- | -- | -- | -- | LG1 | -- | -- | -- | LG1 | -- |
| [illegible] | [illegible] | [illegible] | [illegible] | [illegible] | [illegible] | [illegible] | [illegible] | [illegible] | [illegible] | [illegible] | [illegible] | [illegible] | [illegible] | [illegible] | [illegible] | [illegible] | [illegible] |

| snp_B0826 | snp_B0859 | snp_B0881 | snp_B0891 | snp_B0897 | snp_B0904 | snp_B0925 | snp_B0929 | snp_B0953 | snp_B0965 | snp_B0972 | snp_B0973 | snp_B0974 | snp_B0983 | snp_B0988 | snp_B0998 | snp_B1001 | snp_B1010 | snp_B1021 | snp_B1022 | snp_B1023 | snp_B1027 | snp_B1028 | snp_B1030 | snp_B1067 | snp_B1071 | snp_B1079 | snp_B1086 | snp_B1135 | snp_B1166 | snp_B1168 | snp_B1173 | snp_B1178 | snp_B1201 | snp_B1207 | snp_B1210 | snp_B1285 | snp_B1290 | snp_B1305 | snp_B1311 | snp_B1321 | snp_B1327 | snp_B1337 | snp_B1341 | snp_B1355 | snp_B1364 | snp_B1370 | snp_B1393 | snp_B1412 | snp_B1418 | snp_B1422 | snp_B1441 | snp_B1459 | snp_B1466 | snp_B1471 | snp_B1472 | snp_B1489 | snp_B1516 | snp_B1552 | snp_B1559 | snp_B1562 | snp_B1563 | snp_B1622 | snp_B1624 | snp_B1626 | snp_B1636 | snp_B1640 | snp_B1659 | snp_B1687 | snp_B1709 | snp_B1713 | snp_B1724 | snp_B1742 | snp_B1746 | snp_B1756 | snp_B1777 | snp_B1801 |
|---|---|---|---|---|---|---|---|---|---|---|---|---|---|---|---|---|---|---|---|---|---|---|---|---|---|---|---|---|---|---|---|---|---|---|---|---|---|---|---|---|---|---|---|---|---|---|---|---|---|---|---|---|---|---|---|---|---|---|---|---|---|---|---|---|---|---|---|---|---|---|---|---|---|---|---|---|
| 2 | 2 | 2 | 2 | 2 | 2 | 2 | 2 | 2 | 2 | 2 | 2 | 2 | 2 | 2 | 2 | 2 | 2 | 2 | 2 | 2 | 2 | 2 | 2 | 2 | 2 | 2 | 2 | 2 | 2 | 2 | 2 | 2 | 2 | 2 | 2 | 2 | 2 | 2 | 2 | 2 | 2 | 2 | 2 | 2 | 2 | 2 | 2 | 2 | 2 | 2 | 2 | 2 | 2 | 2 | 2 | 2 | 2 | 2 | 2 | 2 | 2 | 2 | 2 | 2 | 2 | 2 | 2 | 2 | 2 | 2 | 2 | 2 | 2 | 2 | 2 | 2 |
| -- | -- | -- | -- | -- | -- | -- | -- | -- | -- | -- | -- | -- | -- | -- | -- | LG1 | -- | -- | -- | -- | LG1 | -- | LG1 | -- | -- | -- | -- | LG1 | -- | -- | -- | -- | LG1 | LG1 | -- | -- | -- | -- | -- | -- | -- | -- | -- | -- | -- | -- | -- | -- | -- | -- | -- | -- | -- | -- | -- | -- | -- | -- | LG1 | LG1 | -- | -- | LG1 | -- | -- | -- | -- | -- | -- | -- | -- | -- | -- | -- | -- | -- |
| [illegible] | [illegible] | [illegible] | [illegible] | [illegible] | [illegible] | [illegible] | [illegible] | [illegible] | [illegible] | [illegible] | [illegible] | [illegible] | [illegible] | [illegible] | [illegible] | [illegible] | [illegible] | [illegible] | [illegible] | [illegible] | [illegible] | [illegible] | [illegible] | [illegible] | [illegible] | [illegible] | [illegible] | [illegible] | [illegible] | [illegible] | [illegible] | [illegible] | [illegible] | [illegible] | [illegible] | [illegible] | [illegible] | [illegible] | [illegible] | [illegible] | [illegible] | [illegible] | [illegible] | [illegible] | [illegible] | [illegible] | [illegible] | [illegible] | [illegible] | [illegible] | [illegible] | [illegible] | [illegible] | [illegible] | [illegible] | [illegible] | [illegible] | [illegible] | [illegible] | [illegible] | [illegible] | [illegible] | [illegible] | [illegible] | [illegible] | [illegible] | [illegible] | [illegible] | [illegible] | [illegible] | [illegible] | [illegible] | [illegible] | [illegible] | [illegible] | [illegible] |

| snp_B1803 | snp_B1809 | snp_B1816 | snp_B1862 | snp_B1865 | snp_B1867 | snp_B1887 | snp_B1897 | snp_B1917 | snp_B1919 | snp_B1924 | snp_B1931 | snp_B1935 | snp_B1948 | snp_B1952 | snp_B1955 | snp_B1967 | snp_B1984 | snp_B1990 | snp_B2005 | snp_B2006 | snp_B2035 | snp_B2036 | snp_B2048 | snp_B2056 | snp_B2065 | snp_B2077 |
|---|---|---|---|---|---|---|---|---|---|---|---|---|---|---|---|---|---|---|---|---|---|---|---|---|---|---|
| 2 | 2 | 2 | 2 | 2 | 2 | 2 | 2 | 2 | 2 | 2 | 2 | 2 | 2 | 2 | 2 | 2 | 2 | 2 | 2 | 2 | 2 | 2 | 2 | 2 | 2 | 2 |
| -- | -- | -- | -- | LG1 | -- | -- | -- | -- | -- | -- | -- | -- | LG1 | -- | -- | -- | -- | -- | LG1 | -- | -- | -- | -- | -- | -- | -- |
| AB | AB | AB | AB | AA | AB | AB | AB | AB | AB | AB | AB | AB | AA | AB | AA | AB | AA | AB | AA | AB | AA | AB | AB | AA | AB | AA |
| AA | AA | AA | AA | AA | AB | AA | AA | AB | AA | AA | AA | AA | AB | AA | AA | AB | AB | AA | AA | AA | AB | AA | AA | AA | AA | AA |
| AA | AA | AA | AB | AA | AB | AA | AA | AA | AB | AA | AB | AB | AB | AA | AB | AA | AB | AA | AA | AA | AA | AB | AB | AA | AA | AB |
| AA | AB | AA | AB | - | AA | AA | AA | AA | AA | AA | AA | AB | AA | AA | AA | AA | - | AA | AB | AA | AA | AB | AB | AA | AB | AB |
| AB | AB | AB | AB | AB | AB | AA | AB | AB | AA | AA | AB | AB | AB | AA | AB | AB | AB | AB | AA | AB | AB | AA | AB | AA | AA | AA |
| AA | AA | AA | AA | - | - | AA | AA | AA | AA | AA | AA | - | AA | AA | AA | AA | - | - | AA | AA | - | AA | AA | AA | AA | AA |

| snp_B2123 | snp_B2130 | snp_B2132 | snp_B2144 | snp_B2166 | snp_B2180 | snp_B2193 | snp_B2203 | snp_B2205 | snp_B2211 | snp_B2242 | snp_B2247 | snp_B2261 | snp_B2276 | snp_B2311 | snp_B2348 | snp_B2359 | snp_B2367 | snp_B2375 | snp_B2424 | snp_B2428 | snp_B2434 | snp_B2471 | snp_B2515 | snp_B2537 | snp_B2542 |
|---|---|---|---|---|---|---|---|---|---|---|---|---|---|---|---|---|---|---|---|---|---|---|---|---|---|
| 2 | 2 | 2 | 2 | 2 | 2 | 2 | 2 | 2 | 2 | 2 | 2 | 2 | 2 | 2 | 2 | 2 | 2 | 2 | 2 | 2 | 2 | 2 | 2 | 2 | 2 |
| -- | -- | -- | -- | -- | -- | -- | -- | -- | -- | -- | -- | -- | -- | -- | -- | -- | -- | -- | -- | -- | -- | -- | LG1 | -- | -- |
| AA | AA | AB | AA | AB | AA | AB | AB | AA | AA | AA | AA | AB | AA | AB | AB | AB | AB | AB | AB | AB | AA | AB | AA | AA | AB |
| AA | AA | AA | AB | AB | AA | AB | AA | AA | AA | AA | AB | AA | AA | AA | AB | AB | AA | AA | AA | AA | AA | AA | AB | AB | AB |
| AA | AA | AA | AA | AA | AB | AA | AB | AA | AA | AA | AB | AB | AB | AB | AB | AA | AA | AA | AA | AA | AB | AB | AB | AA | AB |
| AA | AB | - | AA | AA | AA | AB | AA | AB | - | AB | AA | - | AA | AA | AA | AA | AB | - | AA | AA | AB | AA | AA | AA | - |
| AB | AA | AB | AB | AB | AA | AB | AB | AB | AA | AA | AB | AB | AB | AB | AB | AB | AA | AA | AB | AB | AA | AA | AB | AB | AB |
| AA | - | - | AA | AA | AA | AA | - | - | - | AA | AA | - | AA | AA | AA | - | - | - | AA | AA | AA | AA | AA | AA | - |

| snp_B2593 | snp_B2638 | snp_B2649 | snp_B2677 | snp_B2680 | snp_B2683 | snp_B2688 | snp_B2690 | snp_B2692 | snp_B2712 | snp_B2735 | snp_B2765 | snp_B2797 | snp_B2800 | snp_B2802 | snp_B2825 | snp_B2845 | snp_B2846 | snp_B2857 | snp_B2877 | snp_B2882 | snp_B2899 | snp_B2904 | snp_B2906 |
|---|---|---|---|---|---|---|---|---|---|---|---|---|---|---|---|---|---|---|---|---|---|---|---|
| 2 | 2 | 2 | 2 | 2 | 2 | 2 | 2 | 2 | 2 | 2 | 2 | 2 | 2 | 2 | 2 | 2 | 2 | 2 | 2 | 2 | 2 | 2 | 2 |
| -- | -- | -- | -- | LG1 | -- | -- | -- | -- | -- | -- | -- | -- | -- | -- | -- | -- | -- | -- | LG1 | -- | -- | -- | -- |
| AA | AB | AB | AB | AA | AB | AB | AB | AA | AA | AA | AB | AA | AB | AB | AA | AB | AB | AB | AB | AB | AB | AA | AA |
| AB | AA | AB | AA | AB | AA | AB | AB | AA | AA | AB | AA | AA | AA | AA | AB | AA | AA | AA | AA | AA | AA | AA | AA |
| AA | AA | AB | AA | AB | AA | AA | AA | AB | AA | AB | AB | AA | AB | AA | AB | AA | AB | AA | AB | AA | AA | AA | AA |
| - | AB | AB | AB | AA | AB | AB | AA | AA | AA | AA | AB | AA | AA | AB | AA | AA | AA | AB | AA | - | AB | AA | AB |
| AB | AA | AB | AA | AB | AB | AB | AB | AB | AB | AB | AB | AB | AB | AB | AB | AA | AA | AA | AB | AB | AB | AB | AA |
| - | AA | AA | AA | AA | - | AA | - | AA | AA | AA | AA | AA | AA | AA | - | AA | AA | AA | - | - | AA | AA | AA |

| snp_B2922 | snp_B2939 | snp_B2957 | snp_B3000 | snp_B3010 | snp_B3016 | snp_B3025 | snp_B3027 | snp_B3049 | snp_B3096 | snp_B3101 | snp_B3115 | snp_B3135 | snp_B3140 | snp_B3144 | snp_B3209 | snp_B3233 | snp_B3241 | snp_B3288 | snp_B3289 | snp_B3290 | snp_B3305 |
|---|---|---|---|---|---|---|---|---|---|---|---|---|---|---|---|---|---|---|---|---|---|
| 2 | 2 | 2 | 2 | 2 | 2 | 2 | 2 | 2 | 2 | 2 | 2 | 2 | 2 | 2 | 2 | 2 | 2 | 2 | 2 | 2 | 2 |
| -- | -- | -- | -- | -- | -- | -- | -- | -- | -- | -- | -- | -- | -- | -- | -- | -- | -- | -- | -- | -- | -- |
| AB | AA | AA | AB | AA | AA | AA | AB | AA | AB | AA | AA | AA | AA | AA | AB | AB | AB | AB | AB | AA | AB |
| AB | AA | AA | AA | AA | AA | AA | AB | AA | AA | AB | AA | AB | AA | AB | AA | AB | AA | AA | AA | AA | AA |
| AA | AA | AB | AB | AB | AA | AB | AA | AA | AB | AB | AA | AB | AA | AA | AB | AA | AB | AA | AB | AA | AB |
| AA | AA | AA | AA | - | AA | AB | AA | AA | AB | - | - | AA | AA | AA | AA | AA | AB | AB | - | AA | AB |
| AB | AB | AB | AA | AA | AB | AB | AB | AB | AB | AB | AA | AB | AA | AB | AB | AB | AB | AB | AB | AB | AB |
| - | AA | - | AA | - | AA | - | AA | AA | AA | - | - | AA | AA | AA | AA | AA | - | AA | - | - | AA |
| AB | AA | AA | AA | AB | AB | AB | AA | AB | AB | AA | AB | AB | AB | AB | AA | AB | AB | AB | AA | AA | AB |
| AB | AA | AB | AB | AB | AA | AA | AA | AB | AA | AA | AA | AA | AB | AA | AB | AA | AA | AA | AB | AB | AA |
| AA | AA | AB | AA | AA | AB | AB | AB | AB | AA | AB | AA | AB | AA | AB | AA | AA | AA | AA | AA | AB | AA |
| AA | AA | AB | AB | AB | AB | AB | AB | AB | AA | AB | AA | AB | AB | AB | AB | AB | AA | AA | AA | AA | AA |
| AA | AA | AB | AB | AA | AA | AB | AB | AB | AA | AB | AA | AB | AB | AA | AB | AA | AA | AA | AB | AA | AA |
| AA | AB | AB | AB | AB | AA | AB | AA | AA | AA | AA | AA | AA | AA | AA | AB | AA | AA | AA | AB | AB | AA |
| AB | AA | AA | AA | AA | AB | AA | AA | AB | AB | AA | AA | AA | AA | AA | AA | AB | AB | AB | AA | AA | AB |
| AB | AA | AA | AB | AA | AB | AB | AA | AB | AB | AA | AA | AA | AA | AA | AA | AB | AB | AB | AA | AA | AB |
| AA | AA | AB | AB | AA | AB | AA | AB | AB | AA | AB | AA | AA | AA | AA | AB | AA | AA | AB | AB | AA | AB |
| AA | AB | AA | AB | AB | AA | AB | AB | AA | AB | AB | AA | AB | AA | AB | AB | AA | AA | AA | AB | AA | AA |
| AB | AB | AB | AB | AB | AB | AB | AB | AA | AB | AA | AB | AB | AA | AB | AB | AB | AB | AB | AB | AB | AB |
| AB | AA | AA | AA | - | - | AA | AB | AB | AB | AB | - | AA | AB | AB | AA | AB | - | AA | - | - | - |
| AB | AB | AB | AB | AB | AB | AB | AB | AB | AB | AB | AB | AB | AB | AB | AB | AB | AB | AB | AB | AB | AB |
| AB | AB | AB | AB | - | AB | AB | AB | AB | AB | AB | AB | AB | AB | AB | AB | AB | AB | AB | AB | AB | AB |
| AA | AB | AB | AB | AB | AB | AB | AB | AB | AA | AB | AA | AB | AB | AB | AB | AB | AA | AB | AB | AA | AA |
| AA | AB | AA | AA | AA | AA | AB | AA | AA | AA | AB | AB | AA | AA | AA | AB | AA | AB | AA | AA | AA | AA |
| AB | AA | - | AB | - | - | AA | - | AA | AB | - | - | AA | AA | AA | AA | AA | - | AB | - | - | - |
| AA | AA | AA | AA | AA | AA | AA | AA | AA | AA | AB | AA | AA | AA | AA | AA | AA | AA | AA | AA | AA | AA |
| AB | AB | AA | AA | AA | AA | AA | AA | AA | AB | AA | AB | AA | AA | AA | AA | AB | AA | AB | AA | AA | AB |
| AA | AA | AA | AA | AA | AA | AA | AA | AA | AB | AB | AA | AA | AA | AA | AA | AA | AA | AA | AA | AA | AA |
| AA | AA | AA | AA | AB | AA | AA | AA | AA | AB | AA | AA | AA | AA | AA | AA | AA | AA | AB | AA | AA | AB |
| AB | AB | AB | AB | AB | AA | AB | AB | AA | AA | AB | AA | AB | AA | AB | AB | AB | AB | AA | AB | AB | AA |
| AA | AA | - | AB | AB | AB | - | AB | AA | AB | - | AA | AA | AA | AA | AB | AA | AB | AA | - | AA | AB |
| - | AB | AB | AB | - | - | AB | AB | AA | AB | - | - | AB | AA | AB | AB | AB | - | AB | - | AB | AB |
| AB | AA | AA | AA | AB | AB | AB | AB | AA | AB | AB | AA | AB | AB | AB | AA | AB | AB | AB | AA | AB | AB |
| AA | AA | AA | AA | AA | AA | AB | AB | AA | AA | AB | AA | AA | AA | AA | AA | AA | AA | AB | AA | AA | AB |

| snp_B3310 | snp_B3371 | snp_B3372 | snp_B3381 | snp_B3391 | snp_B3404 | snp_B3407 | snp_B3416 | snp_B3421 | snp_B3445 | snp_B3452 | snp_B3480 | snp_B3519 | snp_B3529 | snp_B3533 | snp_B3539 | snp_B3547 | snp_B3555 | snp_B3566 | snp_B3570 | snp_B3572 | snp_B3580 | snp_B3593 | snp_B3603 | snp_B3625 |
|---|---|---|---|---|---|---|---|---|---|---|---|---|---|---|---|---|---|---|---|---|---|---|---|---|
| 2 | 2 | 2 | 2 | 2 | 2 | 2 | 2 | 2 | 2 | 2 | 2 | 2 | 2 | 2 | 2 | 2 | 2 | 2 | 2 | 2 | 2 | 2 | 2 | 2 |
| -- | -- | -- | -- | -- | LG1 | -- | -- | -- | -- | -- | -- | -- | -- | -- | -- | -- | -- | LG1 | -- | -- | -- | -- | -- | -- |
| AB | AB | AB | AA | AA | AA | AB | AA | AB | AB | AA | AB | AB | AA | AA | AB | AB | AB | AA | AB | AA | AB | AB | AB | AB |
| AB | AB | AA | AA | AA | AA | AA | AA | AA | AB | AB | AB | AB | AA | AB | AA | AA | AA | AB | AA | AA | AA | AA | AA | AA |
| AB | AB | AA | AB | AB | AB | AA | AA | AA | AA | AB | AA | AB | AA | AB | AB | AA | AB | AB | AA | AB | AB | AB | AB | AA |
| AA | - | AA | AA | AB | AA | AA | AA | AB | AA | AA | AA | AA | AB | AB | AA | AA | AA | AA | AB | - | AA | AA | AA | AA |
| AB | AB | AB | AB | AA | AB | AA | AB | AA | AB | AB | AB | AB | AA | AB | AA | AB | AB | AB | AB | AB | AB | AB | AB | AB |
| AA | AA | - | - | - | - | AA | AA | AA | AA | AA | - | AA | AA | AA | AA | AA | AA | AA | - | AA | AA | AA | AA | AA |
| AB | AA | AA | AA | AB | AA | AB | AA | AA | AB | AA | AA | AB | AA | AB | AA | AB | AA | AA | AB | AB | AA | AB | AA | AB |
| AB | AA | AA | AB | AA | AB | AB | AA | AA | AA | AB | AB | AB | AA | AA | AB | AB | AB | AA | AB | AA | AB | AB | AB | AB |
| AA | AB | AB | AA | AB | AB | AB | AB | AA | AA | AA | AA | AB | AA | AB | AB | AB | AB | AB | AA | AB | AB | AB | AB | AB |
| AB | AB | AA | AB | AA | AB | AA | AB | AB | AA | AA | AA | AB | AA | AA | AB | AA | AB | AB | AA | AB | AB | AB | AB | AA |
| AB | AA | AB | AB | AA | AB | AA | AA | AB | AA | AB | AB | AB | AA | AB | AB | AB | AB | AB | AA | AB | AB | AB | AB | AB |
| AA | AA | AB | AB | AA | AA | AA | AB | AA | AA | AA | AA | AA | AA | AB | AA | AA | AB | AA | AA | AA | AB | AA | AB | AB |
| AA | AA | AA | AA | AA | AA | AA | AA | AA | AA | AA | AB | AA | AA | AA | AA | AA | AA | AA | AB | AB | AA | AB | AB | AA |
| AA | AA | AA | AA | AA | AB | AA | AA | AB | AB | AA | AA | AA | AA | AA | AA | AB | AA | AA | AB | AB | AA | AB | AB | AB |
| AA | AA | AB | AB | AA | AB | AB | AB | AA | AA | AB | AA | AB | AA | AB | AA | AB | AB | AA | AA | AB | AB | AA | AB | AB |
| AB | AB | AB | AB | AA | AA | AA | AA | AA | AA | AA | AB | AA | AA | AA | AB | AA | AA | AB | AB | AB | AB | AA | AA | AB |
| AB | AB | AB | AB | AA | AB | AB | AA | AA | AB | AB | AB | AA | AA | AA | AB | AB | AA | AB | AB | AB | AA | AA | AB | AB |
| AB | AA | AA | AA | - | - | AA | AA | AB | AB | AA | - | AA | AB | AB | AA | AA | AA | AB | - | - | AA | - | AA | AA |
| AB | AB | AB | AB | AB | AB | AB | AB | AB | AB | AB | AB | AB | AB | AB | AB | AB | AB | AB | AB | AB | AB | AB | AB | AB |
| AB | AB | AB | AB | AB | AB | AA | AB | AB | AB | AB | AB | AB | AB | AB | AB | AB | AB | AB | AB | AB | AB | AB | AB | AB |
| AB | AB | AA | AB | AA | AB | AB | AB | AB | AA | AA | AA | AB | AA | AB | AB | AB | AB | AB | AA | AB | AA | AB | AB | AB |
| AA | AA | AA | AB | AB | AA | AB | AA | AB | AA | AB | AA | AB | AA | AB | AB | AA | AB | AA | AA | AA | AB | AB | AB | AB |
| AB | AB | - | AA | - | - | AA | AA | AA | AB | AB | - | AB | AB | AA | AB | AA | AA | AA | AB | AB | AA | AA | AA | AA |
| AA | AA | AA | AA | AA | AA | AA | AA | AA | AA | AA | AA | AA | AA | AA | AA | AA | AA | AA | AA | AA | AA | AA | AA | AA |
| AA | AA | AA | AA | AB | AA | AB | AA | AB | AB | AA | AB | AA | AA | AA | AB | AA | AA | AA | AB | AB | AA | AA | AB | AA |
| AA | AA | AA | AA | AA | AA | AA | AA | AA | AA | AA | AA | AA | AA | AA | AA | AA | AA | AA | AA | AA | AA | AA | AA | AA |
| AA | AA | AA | AB | AA | AB | AB | AA | AB | AA | AA | AA | AA | AA | AB | AA | AB | AA | AA | AA | AA | AB | AA | AB | AA |
| AB | AB | AB | AB | AA | AB | AB | AA | AB | AB | AB | AB | AB | AB | AB | AB | AB | AB | AA | AB | AA | AB | AA | AB | AB |
| AB | AB | AB | AB | - | - | AB | AB | AA | AA | AA | - | AB | AA | AB | AB | AB | AB | AA | - | AA | AB | AB | AB | AB |
| AB | AB | AB | - | - | - | AB | AB | AB | - | AB | - | AB | AB | AB | AB | AB | AB | AB | - | - | AA | AB | AA | AB |
| AB | AA | AA | AA | AB | AA | AA | AB | AA | AB | AA | AA | AB | AB | AB | AB | AB | AA | AA | AB | - | AA | AB | AA | AA |
| AA | AB | AA | AA | AA | AA | AB | AA | AA | AA | AB | AB | AB | AA | AA | AA | AA | AA | AB | AA | AA | AA | AA | AA | AA |

| snp_B3629 | snp_B3630 | snp_B3637 | snp_B3652 | snp_B3685 | snp_B3726 | snp_B3727 | snp_B3755 | snp_B3764 | snp_B3766 | snp_B3769 | snp_B3783 | snp_B3785 | snp_B3787 | snp_B3818 | snp_B3834 | snp_B3844 | snp_B3863 | snp_B3910 | snp_B3915 | snp_B3920 | snp_B3929 | snp_B3955 | snp_B3968 | snp_B3980 | snp_B3984 | snp_B3997 | snp_B4048 | snp_B4059 | snp_B4082 |
|---|---|---|---|---|---|---|---|---|---|---|---|---|---|---|---|---|---|---|---|---|---|---|---|---|---|---|---|---|---|
| 2 | 2 | 2 | 2 | 2 | 2 | 2 | 2 | 2 | 2 | 2 | 2 | 2 | 2 | 2 | 2 | 2 | 2 | 2 | 2 | 2 | 2 | 2 | 2 | 2 | 2 | 2 | 2 | 2 | 2 |
| LG1 | -- | -- | -- | -- | -- | -- | -- | -- | -- | -- | LG1 | -- | -- | -- | -- | -- | -- | LG1 | -- | -- | -- | LG1 | -- | -- | -- | -- | -- | -- | -- |
| AB | AB | AB | AB | AA | AB | AA | AB | AB | AA | AA | AB | AB | AA | AB | AA | AB | AA | AA | AA | AB | AB | AA | AB | AA | AA | AB | AB | AB | AB |
| AA | AB | AB | AB | AA | AA | AA | AA | AA | AB | AB | AA | AB | AA | AB | AA | AB | AA | AA | AA | AB | AB | AA | AB | AA | AB | AA | AB | AB | AB |
| AA | AB | AB | AB | AB | AA | AA | AA | AA | AB | AB | AA | AB | AA | AB | AB | AA | AB | AA | AA | AB | AA | AA | AB | AB | AB | AB | AB | AB | AB |
| - | AA | AA | AA | - | AA | AA | AB | AB | AA | AA | AB | AB | AB | AA | AB | AA | - | AB | AB | AA | AA | AB | AA | AB | AB | AA | AA | AA | AB |
| AB | AA | AA | AB | AA | AA | AA | AA | AB | AB | AB | AA | AB | AB | AB | AA | AA | AA | AA | AA | AA | AB | AA | AB | AB | AB | AB | AB | AB | AB |
| - | AA | AA | AA | - | AA | AA | AA | - | - | - | AA | - | AA | AA | - | AA | AA | - | AA | AB | AA | AA | AA | AA | AA | - | - | - | - |
| AA | AA | AB | AB | AB | AB | AB | AB | AB | AB | AB | AB | AA | AA | AA | AB | AB | - | AB | AB | AA | AB | AB | AA | AB | AA | AA | AA | AA | AB |
| AB | AB | AB | AB | AA | AB | AB | AA | AA | AB | AB | AA | AA | AA | AA | AA | AB | AA | AA | AA | AA | AB | AB | AA | AB | AA | AA | AA | AB | AA |
| AB | AA | AB | AB | AB | AA | AA | AA | AB | AA | AB | AA | AA | AA | AB | AB | AA | AB | AA | AA | AB | AA | AA | AB | AA | AB | AA | AB | AB | AB |
| AA | AA | AB | AB | AB | AB | AA | AA | AB | AA | AB | AA | AA | AA | AB | AB | AA | AB | AA | AA | AA | AA | AA | AB | AA | AB | AB | AA | AB | AA |
| AA | AA | AB | AB | AA | AB | AB | AA | AA | AA | AB | AA | AA | AA | AB | AB | AB | AB | AA | AA | AA | AA | AA | AB | AA | AB | AB | AB | AB | AB |
| AB | AB | AB | AA | AB | AB | AB | AA | AA | AA | AA | AA | AA | AB | AA | AB | AA | - | AA | AA | AA | AA | AA | AA | AA | AA | AA | AA | AB | AA |
| AA | AA | AA | AA | AA | AA | AA | AA | AA | AA | AA | AA | AB | AA | AA | AA | AB | AB | AA | AA | AA | AB | AA | AA | AB | AA | AA | AA | AB | AA |
| AA | AA | AA | AB | AA | AB | AB | AB | AB | AB | AB | AB | AB | AA | AA | AA | AB | AA | AA | AA | AA | AB | AB | AA | AA | AB | AB | AA | AB | AB |
| AA | AA | AB | AA | AA | AB | AA | AA | AB | AA | AB | AA | AA | AB | AB | AB | AA | AA | AA | AA | AB | AB | AA | AB | AB | AA | AB | AB | AB | AB |
| AB | AA | AB | AA | AA | AB | AB | AB | AA | AB | AA | AA | AA | AB | AB | AA | AB | AA | AA | AB | AB | AA | AA | AB | AA | AB | AB | AB | AB | AA |
| AB | AB | AB | AA | AB | AB | AB | AB | AB | AB | AB | AB | AB | AB | AB | AB | AA | AB | AB | AB | AB | AB | AB | AB | AB | AA | AB | AB | AB | AB |
| - | AA | AB | AA | - | AA | AA | AB | - | AA | AA | AB | AA | AA | AB | - | AB | AB | - | AB | AA | AB | AB | AA | AB | AA | - | - | - | - |
| AB | AB | AB | AB | AB | AB | AB | AB | AB | AB | AB | AB | AB | AB | AB | AB | AB | AB | AB | AB | AB | AB | AB | AB | AB | AB | AB | AB | AB | AB |
| AB | AB | AB | AB | AB | AB | AB | AB | AB | AB | AB | AB | AB | AB | AB | AB | AB | AB | AB | AB | AA | AB | AB | AB | AB | AB | AB | AB | AB | AB |
| AA | AB | AB | AB | AB | AB | AA | AA | AB | AA | AB | AB | AA | AB | AB | AB | AA | AB | AA | AA | AB | AA | AB | AB | AA | AB | AB | AB | AB | AA |
| AB | AB | AA | AB | AB | AB | AB | AA | AA | AA | AB | AB | AA | AB | AA | AB | AB | AB | AB | AB | AB | AA | AB | AA | AB | AA | AB | AA | AB | AA |
| - | AB | AB | AA | - | AA | - | AB | - | - | AB | AB | - | AA | AA | AB | AA | - | - | AA | - | AB | AB | AA | AB | - | - | - | - | - |
| AA | AA | AA | AA | AA | AA | AA | AA | AA | AA | AA | AA | AA | AA | AA | AA | AA | AB | AA | AA | AB | AA | AA | AA | AA | AA | AA | AA | AA | AA |
| AB | AB | AA | AA | AB | AA | AB | AA | AA | AB | AA | AB | AB | AB | AA | AA | AB | - | AB | AB | AA | AB | AB | AA | AA | AB | AA | AA | AB | AB |
| AA | AA | AA | AA | AA | AA | AA | AA | AA | AA | AA | AA | AA | AA | AA | AA | AA | AB | AA | AA | AB | AA | AA | AA | AA | AA | AA | AA | AA | AA |
| AA | AA | AA | AB | AA | AA | AA | AB | AA | AA | AA | AB | AB | AA | AA | AA | AB | AA | AA | AA | AA | AB | AA | AA | AB | AA | AA | AA | AB | AA |
| AB | AA | AB | AB | AA | AA | AA | AA | AA | AB | AB | AA | AA | AB | AB | AB | AA | AA | AA | AA | AB | AB | AA | AB | AA | AA | AB | AB | AB | AB |
| - | AB | AB | AB | AA | AA | AA | AA | AA | AB | AB | AB | AB | AA | AA | AB | AA | AA | AA | AB | AA | AA | AB | AA | AB | AB | AB | AA | AB | AA |
| AB | AB | AB | AB | - | AB | AB | AB | - | AB | AB | AB | AB | AB | AB | - | AB | - | AA | AA | AB | AB | AB | AB | AB | AB | - | - | AA | - |
| AA | AA | AB | AB | AB | AB | AB | AB | AB | AB | AB | AB | AA | AA | AA | AB | AA | - | AB | AB | AA | AB | AB | AA | AB | AA | AA | AA | AA | AB |
| AA | AA | AA | AA | AA | AB | AA | AA | AB | AB | AB | AB | AB | AA | AB | AA | AB | AB | AA | AA | AB | AA | AB | AA | AB | AA | AB | AA | AA | AA |

| snp_B4100 | snp_B4101 | snp_B4169 | snp_B4185 | snp_B4192 | snp_B4194 | snp_B4197 | snp_B4199 | snp_B4202 | snp_B4203 | snp_B4214 | snp_B4221 | snp_B4272 | snp_B4281 | snp_B4303 | snp_B4314 | snp_B4323 |
|---|---|---|---|---|---|---|---|---|---|---|---|---|---|---|---|---|
| 2 | 2 | 2 | 2 | 2 | 2 | 2 | 2 | 2 | 2 | 2 | 2 | 2 | 2 | 2 | 2 | 2 |
| -- | LG1 | -- | LG1 | -- | -- | -- | -- | -- | -- | -- | -- | -- | -- | -- | -- | -- |
| AB | AA | AA | AA | AA | AB | AA | AA | AA | AB | AA | AA | AB | AB | AB | AB | AA |
| AA | AA | AA | AB | AB | AA | AB | AA | AA | AA | AA | AA | AA | AA | AA | AA | AA |
| AA | AA | AA | AA | AB | AB | AB | AB | AA | AA | AA | AB | AB | AB | AB | AA | AA |
| AA | AB | AB | AB | AB | AA | AA | - | AB | AB | AB | AB | - | AA | AB | AB | AA |
| AA | AB | AB | AA | AB | AB | AB | AA | AB | AB | AB | AA | AB | AA | AB | AA | AB |
| AA | AA | AA | AA | - | AA | AA | - | AA | AA | - | AA | - | AA | - | AA | AA |
| AB | AB | AB | AB | AB | AA | AA | AB | AA | AB | AB | AA | AB | AA | AA | AA | AB |
| AB | AA | AB | AA | AA | AB | AA | AB | AA | AA | AB | AA | AA | AB | AA | AA | AA |
| AB | AA | AB | AA | AB | AB | AB | AB | AB | AB | AB | AB | AB | AB | AA | AA | AB |
| AA | AA | AA | AA | AB | AB | AB | AB | AB | AB | AB | AB | AA | AA | AA | AB | AB |
| AB | AA | AA | AA | AB | AB | AA | AB | AB | AA | AB | AA | AA | AB | AB | AA | AA |
| AA | AA | AA | AA | AB | AB | AA | AB | AB | AB | AA | AB | AB | AB | AA | AA | AA |
| AA | AA | AB | AB | AA | AA | AB | AA | AA | AA | AB | AA | AA | AA | AB | AA | AB |
| AA | AB | AA | AB | AB | AB | AA | AB | AB | AB | AB | AB | AB | AB | AA | AB | AA |
| AA | AA | AA | AA | AB | AB | AB | AA | AA | AA | AA | AB | AA | AB | AB | AA | AA |
| AA | AA | AA | AA | AA | AA | AB | AB | AA | AA | AB | AA | AA | AA | AA | AA | AB |
| AB | AB | AB | AB | AB | AA | AB | AB | AB | AB | AB | AA | AB | AA | AB | AB | AB |
| AA | AB | - | AB | - | AA | AB | - | - | - | AA | - | - | AA | - | - | AA |
| AB | AB | AB | AB | AB | AB | AB | AB | AB | AB | AB | AB | AB | AB | AB | AB | AB |
| AA | AB | AB | AB | AB | AB | AB | AB | AB | AB | AB | AB | AB | AB | AB | AB | AB |
| AB | AA | AA | AA | AB | AB | AB | AB | AB | AB | AB | AB | AA | AB | AA | AB | AB |
| AB | AB | AA | AB | AB | AA | AA | AA | AB | AB | AA | AB | AA | AB | AB | AB | AA |
| AB | AB | - | AB | AA | AA | AB | - | - | AB | AB | - | - | AB | AA | - | AB |
| AA | AA | AA | AA | AA | AA | AA | AA | AA | AA | AA | AA | AA | AA | AA | AA | AA |
| AA | AB | AA | AB | AA | AA | AB | AA | AA | AB | AB | AA | AB | AA | AA | AB | AA |
| AA | AA | AA | AA | AA | AA | AA | AA | AA | AA | AA | AA | AA | AA | AA | AA | AA |
| AB | AB | AB | AA | AB | AB | AA | AA | AB | AA | AB | AA | AB | AB | AB | AB | AA |
| AB | AA | AA | AA | AA | AB | AA | AB | AA | AA | AA | AA | AA | AB | AA | AA | AA |
| AA | AA | AA | AA | AA | AB | AB | - | AA | AB | AA | AB | AA | AB | AB | AA | AA |
| AB | AB | - | AB | - | AB | AB | - | - | AB | AB | - | - | AB | AB | AB | AB |

| snp_B4356 | snp_B4364 | snp_B4415 | snp_B4435 | snp_B4463 | snp_B4480 | snp_B4509 | snp_B4516 | snp_B4527 | snp_B4529 | snp_B4531 | snp_B4594 |
|---|---|---|---|---|---|---|---|---|---|---|---|
| 2 | 2 | 2 | 2 | 2 | 2 | 2 | 2 | 2 | 2 | 2 | 2 |
| -- | -- | -- | -- | -- | -- | -- | -- | -- | -- | -- | -- |
| AB | AB | AA | AB | AB | AA | AB | AA | AB | AB | AB | AB |
| AA | AA | AB | AA | AB | AA | AA | AA | AA | AB | AA | AA |
| AA | AA | AB | AA | AA | AB | AA | AA | AA | AB | AA | AA |
| AB | AB | AA | AB | AB | AB | - | AA | AA | AA | - | AA |
| AB | AB | AB | AB | AB | AB | AB | AA | AB | AB | AA | AB |
| AA | AA | AB | AA | AA | AA | - | AA | AA | AA | - | AA |
| AA | AB | AB | AB | AA | AA | AB | AB | AB | AA | AB | AA |
| AB | AA | AA | AA | AA | AA | AA | AB | AA | AB | AB | AA |
| AB | AA | AB | AB | AB | AA | AA | AB | AA | AB | AA | AA |
| AA | AA | AB | AA | AB | AB | AA | AB | AA | AB | AB | AB |
| AA | AA | AB | AA | AB | AA | AA | AB | AA | AB | AB | AA |
| AB | AA | AB | AB | AB | AB | AA | AB | AA | AB | AB | AA |
| AB | AB | AA | AA | AA | AA | AB | AA | AB | AA | AA | AB |
| AB | AA | AA | AB | AB | AA | AB | AA | AA | AA | AA | AA |
| AA | AA | AA | AA | AA | AB | AA | AA | AB | AB | AB | AB |
| AA | AA | AB | AA | AA | AA | AA | AA | AB | AB | AA | AB |
| AA | AB | AB | AB | AB | AB | AB | AB | AB | AB | AB | AB |
| AB | AB | AA | AB | AB | AB | - | AB | - | - | - | AB |
| AB | AB | AB | AB | AB | AB | AB | AB | AB | AB | AB | AB |
| AB | AB | AB | AB | AB | AB | AB | AB | AB | AB | AB | AB |
| AA | AA | AB | AA | AB | AB | AA | AB | AA | AB | AB | AB |
| AA | AA | AA | AA | AB | AB | AA | AB | AA | AA | AB | AA |
| AA | AB | AA | AA | AA | AA | AB | AB | AB | AA | - | AB |
| AA | AA | AA | AA | AA | AA | AA | AA | AA | AA | AA | AA |
| AA | AB | AA | AB | AA | AB | AB | AB | AB | AA | AB | AB |
| AA | AA | AA | AA | AA | AA | AA | AA | AA | AA | AA | AA |
| AB | AB | AA | AB | AB | AA | AB | AB | AA | AB | AA | AA |
| AA | AA | AB | AA | AA | AB | AA | AB | AB | AB | AB | AB |
| AA | AA | AB | AA | AA | AB | AA | AA | AA | AB | AB | AA |
| AA | AB | AB | AB | AB | AB | - | AB | AB | AB | - | AB |

| snp_B4601 | snp_B4614 | snp_B4625 | snp_B4631 | snp_B4640 | snp_B4658 | snp_B4688 | snp_B4727 | snp_B4729 | snp_B4747 | snp_B4765 |
|---|---|---|---|---|---|---|---|---|---|---|
| 2 | 2 | 2 | 2 | 2 | 2 | 2 | 2 | 2 | 2 | 2 |
| -- | -- | -- | -- | -- | -- | LG1 | -- | -- | LG1 | -- |
| AB | AB | AB | AB | AA | AB | AA | AA | AB | AA | AB |
| AA | AA | AB | AB | AA | AA | AB | AA | AB | AB | AA |
| AB | AB | AB | AB | AB | AA | AB | AA | AA | AA | AA |
| AA | AA | AA | AA | AB | AB | AA | AB | AB | AB | AA |
| AB | AB | AB | AB | AA | AB | AB | AB | AA | AA | AB |
| AA | - | AA | AA | AA | AA | AA | AA | AA | AA | AA |
| AB | AB | AB | AA | AA | AB | AA | AA | AB | - | AB |
| AA | AA | AA | AB | AA | AA | AA | AA | AA | AA | AB |
| AA | AB | AB | AB | AB | AA | AB | AB | AB | AA | AA |
| AA | AA | AA | AB | AA | AA | AB | AB | AA | AA | AA |
| AB | AA | AB | AB | AA | AA | AB | AB | AA | AA | AB |
| AA | AA | AA | AB | AB | AB | AA | AB | AA | AA | AA |
| AB | AB | AA | AA | AA | AA | AA | AA | AA | AB | AB |
| AA | AB | AB | AB | AA | AB | AB | AB | AA | AA | AB |
| AA | AB | AB | AB | AA | AA | AA | AA | AA | AA | AA |
| AB | AA | AA | AA | AB | AA | AB | AA | AA | AA | AB |
| AB | AB | AB | AB | AB | AB | AB | AB | AB | AA | AB |
| AB | AB | - | AA | AB | AA | AA | AB | - | - | AB |
| AB | AB | AB | AB | AB | AB | AB | AB | AB | AB | AB |
| AB | AB | AB | AB | AB | AB | AB | AB | AA | AB | AB |
| AA | AA | AA | AB | AA | AA | AB | AB | AB | AA | AA |
| AA | AA | AA | AA | AA | AB | AA | AB | AB | AA | AA |
| - | AA | - | AA | AB | AB | AA | AA | - | - | AB |
| AA | AA | AA | AA | AA | AA | AA | AA | AA | AA | AA |
| AB | AA | AB | AA | AA | AA | AA | AA | AA | AB | AB |
| AA | AA | AA | AA | AA | AA | AA | AA | AA | AA | AA |
| AA | AA | AA | AB | AA | AB | AB | AB | AA | AA | AA |
| AA | AA | AB | AB | AB | AA | AB | AA | AA | AA | AA |
| AA | AA | AA | AB | AB | AA | AB | AA | AB | AA | AB |
| AB | AB | - | AB | AB | AB | AB | AB | AB | AA | AB |

| snp_B4771 | snp_B4777 | snp_B4778 | snp_B4783 | snp_B4787 | snp_B4799 | snp_B4805 | snp_B4825 | snp_B4827 | snp_B4835 | snp_B4836 |
|---|---|---|---|---|---|---|---|---|---|---|
| 2 | 2 | 2 | 2 | 2 | 2 | 2 | 2 | 2 | 2 | 2 |
| LG1 | -- | -- | -- | -- | -- | -- | -- | -- | -- | -- |
| AB | AA | AA | AB | AA | AB | AA | AA | AA | AB | AB |
| AB | AA | AA | AB | AB | AA | AA | AA | AA | AB | AA |
| AA | AA | AA | AB | AB | AA | AB | AB | AA | AB | AB |
| AB | AA | AB | AA | - | AB | AB | AA | AB | AA | AA |
| AA | AA | AB | AB | AB | AA | AA | AA | AB | AB | AA |
| AA | AA | AA | AA | AA | AA | AA | AA | AA | AA | - |
| AB | AA | AB | AA | - | AB | AB | AB | AB | AA | AA |
| AA | AA | AB | AA | AA | AA | AA | AB | AA | AB | AB |
| AB | AA | AB | AA | AA | AB | AB | AB | AA | AB | AB |
| AB | AA | AA | AB | AB | AA | AA | AB | AA | AB | AB |
| AA | AB | AB | AB | AA | AB | AA | AA | AA | AB | AB |
| AA | AA | AA | AA | AA | AA | AA | AB | AA | AA | AB |
| AB | AA | AB | AA | AA | AA | AA | AA | AA | AA | AA |
| AA | AA | AA | AA | AA | AA | AA | AA | AA | AA | AA |
| AA | AA | AA | AA | AA | AB | AA | AB | AA | AB | AB |
| AA | AB | AA | AB | AA | AA | AA | AB | AA | AA | - |
| AB | AB | AB | AB | AB | AA | AA | AB | AB | AB | AB |
| AB | AA | AB | AB | - | AA | AA | AB | AB | AA | - |
| AB | AB | AB | AB | AB | AB | AB | AB | AB | AB | AB |
| AA | AB | AB | AB | AB | AA | AB | AB | AB | AB | AB |
| AB | AA | AA | AB | AA | AB | AA | AB | AA | AB | AB |
| AB | AA | AA | AA | AA | AB | AB | AA | AB | AB | AA |
| AB | AB | AA | AA | AA | AA | AB | AB | AB | AA | - |
| AA | AA | AA | AA | AA | AA | AA | AA | AA | AA | AA |
| AA | AA | AA | AA | AA | AB | AB | AA | AB | AA | AA |
| AA | AA | AA | AA | AA | AA | AA | AA | AA | AA | AA |
| AA | AA | AB | AA | AA | AB | AA | AB | AB | AA | AA |
| AA | AB | AA | AB | AB | AB | AA | AB | AB | AB | AB |
| AA | AA | AA | AB | - | AB | AB | AB | AA | AB | - |
| AB | AB | AB | - | AB | AB | AA | AB | AA | AB | - |

| snp_B4855 | snp_B4867 | snp_B4869 | snp_B4885 | snp_B4898 | snp_B4903 | snp_B4908 | snp_B4942 | snp_B4957 | snp_B4967 | snp_B4996 | snp_B4999 |
|---|---|---|---|---|---|---|---|---|---|---|---|
| 2 | 2 | 2 | 2 | 2 | 2 | 2 | 2 | 2 | 2 | 2 | 2 |
| -- | LG1 | -- | -- | -- | -- | -- | -- | -- | -- | -- | -- |
| AB | AA | AB | AB | AB | AA | AB | AA | AB | AB | AA | AB |
| AA | AB | AA | AA | AA | AA | AA | AA | AA | AB | AA | AA |
| AA | AB | AA | AA | AA | AA | AA | AB | AB | AA | AA | AA |
| AA | AA | AA | AA | AB | AB | AB | AB | AB | AA | AA | AA |
| AB | AB | AB | AB | AA | AB | AB | AB | AB | AB | AA | AB |
| AA | AA | AA | AA | - | AA | AA | AA | AA | - | AA | AA |
| AB | AA | AA | AB | AB | AA | AA | AA | AB | AA | AB | AB |
| AA | AA | AB | AA | AA | AA | AA | AA | AB | AB | AB | AA |
| AA | AB | AA | AB | AB | AB | AB | AA | AB | AA | AA | AA |
| AA | AB | AA | AB | AA | AA | AB | AA | AB | AA | AB | AA |
| AA | AB | AA | AA | AB | AA | AB | AB | AA | AB | AB | AA |
| AA | AA | AB | AA | AA | AB | AB | AA | AA | AA | AB | AA |
| AA | AA | AA | AB | AA | AA | AA | AB | AB | AB | AA | AB |
| AB | AA | AB | AB | AA | AA | AB | AB | AA | AA | AB | AB |
| AB | AB | AA | AB | AA | AB | AA | AB | AA | AB | AA | AA |
| AA | AB | AA | AA | AA | AA | AA | AB | AA | AB | AA | AA |
| AB | AB | AB | AB | AB | AB | AB | AB | AB | AB | AB | AB |
| AB | AA | AA | AB | - | AA | - | AA | AB | - | AB | AA |
| AB | AB | AB | AB | AB | AB | AB | AB | AB | AB | AB | AB |
| AB | AB | AB | AB | AA | AB | AB | AB | AB | AB | AB | AB |
| AA | AB | AA | AB | AB | AA | AB | AA | AA | AA | AB | AA |
| AA | AA | AB | AA | AB | AB | AB | AA | AA | AA | AB | AA |
| AB | AA | AB | AB | AB | AA | - | AB | AA | - | AB | AB |
| AA | AA | AA | AA | AA | AA | AA | AA | AA | AA | AA | AA |
| AA | AA | AB | AA | AB | AB | AA | AB | AA | AB | AB | AB |
| AA | AA | AA | AA | AA | AA | AA | AA | AA | AA | AA | AA |
| AA | AA | AA | AA | AB | AB | AB | AB | AA | AA | AB | AA |
| AB | AB | AB | AA | AB | AB | AA | AA | AA | AB | AA | AA |
| AA | AB | AA | AA | AB | AB | AA | AB | AA | AA | AA | AA |
| AB | - | AB | AB | AB | AB | - | - | AB | AB | AB | AB |

| snp_B5012 | snp_B5027 | snp_B5116 | snp_B5139 | snp_B5140 | snp_B5159 | snp_B5168 | snp_B5185 | snp_B5206 | snp_B5223 | snp_B5233 | snp_B5267 | snp_B5272 | snp_B5286 |
|---|---|---|---|---|---|---|---|---|---|---|---|---|---|
| 2 | 2 | 2 | 2 | 2 | 2 | 2 | 2 | 2 | 2 | 2 | 2 | 2 | 2 |
| -- | -- | -- | -- | -- | -- | -- | -- | -- | -- | -- | -- | -- | -- |
| AA | AB | AA | AA | AB | AB | AA | AB | AA | AB | AB | AB | AB | AA |
| AA | AA | AA | AA | AA | AB | AB | AA | AA | AB | AA | AB | AB | AA |
| AB | AA | AB | AA | AA | AA | AB | AB | AA | AB | AA | AA | AB | AB |
| AA | AB | AB | AB | AA | AA | AA | AA | AA | AB | AB | AA | AA | - |
| AB | AA | AA | AB | AA | AB | AB | AA | AB | AB | AA | AB | AB | AA |
| - | AA | AA | AA | AA | AA | - | AA | AA | AA | - | AA | AA | AA |
| AA | AB | AB | AB | AA | AB | AA | AA | AB | AB | - | AB | AA | AB |
| AB | AA | AA | AB | AB | AA | AA | AB | AB | AA | AA | AB | AA | AA |
| AB | AB | AA | AB | AB | AA | AB | AB | AA | AB | AB | AA | AA | AB |
| AA | AA | AB | AB | AA | AA | AB | AA | AA | AA | AA | AA | AA | AA |
| AB | AB | AB | AB | AA | AA | AB | AB | AB | AA | AA | AA | AB | AA |
| AB | AA | AA | AA | AB | AA | AA | AA | AA | AB | AA | AA | AA | AA |
| AA | AA | AA | AB | AA | AA | AA | AA | AA | AA | AA | AB | AA | AA |
| AA | AA | AB | AA | AA | AA | AA | AA | AA | AB | AA | AB | AB | AA |
| AB | AB | AA | AA | AB | AA | AB | AA | AB | AB | AA | AA | AA | AA |
| AB | AA | AA | AA | AA | AA | AB | AA | AA | AA | AA | AA | AB | AA |
| AB | AA | AB | AB | AB | AB | AB | AA | AB | AB | AB | AB | AB | AA |
| - | AA | AB | - | AA | - | - | AA | AB | AB | - | AB | - | - |
| AB | AB | AB | AB | AB | AB | AB | AB | AB | AB | AB | AB | AB | AB |
| AB | AA | AB | AB | AB | AB | AB | AB | AB | AB | AA | AB | AB | - |
| AA | AB | AB | AB | AA | AA | AB | AB | AA | AA | AB | AA | AA | AA |
| AB | AB | AA | AA | AB | AA | AA | AB | AB | AB | AB | AA | AA | AB |
| - | AA | AB | AA | - | AB | - | AA | AA | - | AB | AB | AA | AA |
| AA | AA | AA | AA | AA | AA | AA | AA | AA | AA | AA | AA | AA | AA |
| AA | AA | AA | AA | AB | AB | AB | AB | AB | AA | AA | AB | AA | AB |
| AA | AA | AA | AA | AA | AA | AA | AA | AA | AA | AA | AA | AA | AA |
| AA | AB | AB | AB | AA | AA | AA | AB | AA | AB | AA | AA | AA | AA |
| AB | AB | AA | AA | AB | AA | AB | AB | AA | AA | AA | AB | AB | AA |
| AB | AA | AA | AA | AA | AA | AA | AA | AB | AA | AA | AA | AB | - |
| AB | AB | AB | AB | - | AB | AB | AB | AA | - | - | AB | AB | - |

| snp_B5289 | snp_B5299 | snp_B5318 | snp_B5325 | snp_B5415 | snp_B5430 | snp_B5451 | snp_B5460 | snp_B5475 | snp_B5502 | snp_B5522 | snp_B5536 | snp_B5545 | snp_B5569 | snp_B5590 | snp_B5604 | snp_B5614 | snp_B5650 | snp_B5655 | snp_B5658 | snp_B5661 | snp_B5673 | snp_B5678 | snp_B5679 | snp_B5681 | snp_B5690 | snp_B5695 | snp_B5702 | snp_B5707 | snp_B5731 | snp_B5799 | snp_B5821 | snp_B5826 | snp_B5857 | snp_B5873 | snp_B5900 | snp_B5977 | snp_B5985 | snp_B5991 | snp_B5993 | snp_B5994 | snp_B6006 | snp_B6010 | snp_B6019 | snp_B6041 | snp_B6156 | snp_B6157 | snp_B6161 | snp_B6269 | snp_B6354 | snp_B6358 | snp_B0005 | snp_B0016 | snp_B0022 | snp_B0045 | snp_B0047 | snp_B0069 | snp_B0118 | snp_B0144 | snp_B0194 | snp_B0198 | snp_B0250 | snp_B0297 | snp_B0301 | snp_B0304 | snp_B0316 | snp_B0335 | snp_B0345 | snp_B0347 | snp_B0353 | snp_B0355 | snp_B0374 | snp_B0408 | snp_B0416 | snp_B0475 | snp_B0491 | snp_B0518 |
|---|---|---|---|---|---|---|---|---|---|---|---|---|---|---|---|---|---|---|---|---|---|---|---|---|---|---|---|---|---|---|---|---|---|---|---|---|---|---|---|---|---|---|---|---|---|---|---|---|---|---|---|---|---|---|---|---|---|---|---|---|---|---|---|---|---|---|---|---|---|---|---|---|---|---|---|---|
| 2 | 2 | 2 | 2 | 2 | 2 | 2 | 2 | 2 | 2 | 2 | 2 | 2 | 2 | 2 | 2 | 2 | 2 | 2 | 2 | 2 | 2 | 2 | 2 | 2 | 2 | 2 | 2 | 2 | 2 | 2 | 2 | 2 | 2 | 2 | 2 | 2 | 2 | 2 | 2 | 2 | 2 | 2 | 2 | 2 | 2 | 2 | 2 | 2 | 2 | 2 | 3 | 3 | 3 | 3 | 3 | 3 | 3 | 3 | 3 | 3 | 3 | 3 | 3 | 3 | 3 | 3 | 3 | 3 | 3 | 3 | 3 | 3 | 3 | 3 | 3 | 3 |
| -- | -- | -- | -- | -- | -- | -- | -- | -- | -- | -- | -- | -- | -- | -- | -- | -- | -- | -- | LG1 | -- | -- | LG1 | -- | LG1 | -- | -- | -- | -- | LG1 | LG1 | -- | -- | -- | -- | -- | -- | -- | -- | -- | -- | LG1 | -- | -- | -- | -- | -- | -- | -- | -- | -- | -- | LG9 | -- | LG9 | -- | -- | -- | -- | -- | -- | -- | -- | LG9 | -- | -- | -- | -- | -- | -- | -- | -- | LG9 | LG9 | -- | -- | -- |
| [illegible] | | | | | | | | | | | | | | | | | | | | | | | | | | | | | | | | | | | | | | | | | | | | | | | | | | | | | | | | | | | | | | | | | | | | | | | | | | | | |

| snp_B0524 3 | snp_B0539 3 | snp_B0562 3 | snp_B0575 3 | snp_B0580 3 | snp_B0636 3 | snp_B0654 3 | snp_B0677 3 | snp_B0684 3 | snp_B0707 3 | snp_B0735 3 | snp_B0772 3 | snp_B0790 3 | snp_B0796 3 | snp_B0816 3 | snp_B0828 3 | snp_B0832 3 |
|---|---|---|---|---|---|---|---|---|---|---|---|---|---|---|---|---|
| -- | -- | -- | -- | -- | -- | -- | -- | -- | -- | -- | -- | -- | LG9 | -- | -- | -- |
| AA | AB | AB | AA | AA | AB | AB | AA | AB | AA | AA | AA | AB | AA | AA | AB | AA |
| AB | AA | AA | AB | AB | AA | AB | AA | AA | AB | AA | AA | AA | AA | AA | AA | AA |
| AA | AB | AB | AA | AA | AB | AB | AB | AA | AA | AA | AA | AB | AA | AA | AB | AB |
| AB | AA | AA | AA | AA | AA | AA | AA | AA | AA | AA | AA | AA | AB | AB | AA | AA |
| AB | AB | AB | AA | AA | AB | AB | AA | AB | AA | AB | AB | AB | AA | AB | AA | AA |
| AA | AB | AB | AA | - | AB | AB | AB | - | AA | AA | AB | AB | AA | AA | AB | AB |
| AB | AA | AA | AA | - | AA | AA | AA | AA | AA | AA | AA | AA | AA | AA | AA | AA |
| AB | AB | AB | AB | AB | AB | AB | AB | AB | AB | AB | AB | AB | AB | AB | AB | AB |
| AA | AB | AB | AA | AA | AB | AB | AA | AA | AB | AA | AB | AB | AB | AA | AB | AA |
| AA | AA | AA | AA | AA | AB | AB | AA | AA | AB | AA | AA | AA | AA | AA | AB | AA |
| AA | AA | AB | AB | AA | AB | AA | AA | AA | AB | AB | AB | AA | AB | AA | AB | AA |
| AA | AB | AB | AA | AB | AB | AB | AB | AA | AA | AA | AB | AB | AA | AA | AB | AA |
| AB | AA | AA | AA | AA | AA | AA | AB | AA | AA | AA | AA | AA | AA | AA | AA | AB |
| AB | AA | AA | AB | AB | AA | AA | AA | AB | AB | AB | AA | AA | AB | AB | AA | AB |
| AA | AB | AA | AB | AA | AB | AB | AB | AA | AB | AB | AB | AB | AA | AB | AB | AB |
| AB | AA | AB | AB | AB | AB | AB | AB | AA | AB | AB | AB | AB | AB | AB | AB | AB |
| AA | AB | AB | AB | AB | AB | AB | AA | AA | AB | AA | AB | AB | AB | AA | AB | AA |
| AA | - | - | AA | - | - | AB | AA | AB | AA | - | - | AA | - | - | AB | - |
| AA | AB | AA | AB | AB | AA | AB | AA | AA | AB | AA | AA | AA | AA | AA | AB | AA |
| AB | AA | AB | AB | AA | AA | AA | AB | AB | AB | AB | AA | AB | - | AA | AB | AB |
| AA | AB | AB | AB | AB | AB | AB | AA | AB | AB | AB | AA | AB | AB | AB | AB | AA |
| AB | AB | AB | AA | AB | AB | AB | AB | AB | AB | AB | AB | AB | AA | AB | AB | AB |
| AB | - | - | AB | - | - | - | AB | - | AB | - | - | AA | - | - | AA | - |
| AA | AA | AA | AA | AA | AA | AA | AB | AA | AA | AA | AA | AA | AA | AA | AA | AB |
| AA | AB | AA | AB | AB | AA | AA | AA | AB | AB | AB | AA | AA | AB | AB | AA | AA |
| AA | AB | AB | AA | AB | AB | AA | AA | AA | AB | AB | AB | AA | AA | AB | AB | AA |
| AA | AB | AA | AA | AA | AA | AA | AB | AB | AA | AA | AA | AA | - | AA | AA | AA |
| AA | AA | AB | AA | AA | AB | AB | AA | AA | AA | AA | AA | AB | - | AB | AB | AA |
| AA | AB | - | AA | - | - | AA | AB | AA | AA | AA | AA | AB | - | AA | AA | AA |
| AA | AA | - | AA | AA | AA | AA | AA | AB | AA | AB | AA | AA | AA | AB | AA | AA |
| AA | AB | AB | AA | AB | AB | AA | AB | AB | AB | AA | AB | AB | AA | AB | AB | AB |
| AA | - | AA | AA | - | AA | AA | AA | AA | AA | AA | AA | AA | AA | AA | AA | AA |
| AB | AA | AA | AA | - | AA | AA | AA | AB | AB | AA | AB | AA | - | - | AA | AA |
| AB | AA | AB | AA | AA | AB | AB | AA | AA | AA | - | AB | AB | AA | AB | AA | - |
| AB | AA | AA | AB | AB | - | AB | AB | AA | AB | AA | AA | AA | AB | - | AB | AB |
| AA | AA | AB | AA | AA | AA | AA | AB | AB | AB | AB | AA | AB | AA | AA | AB | AB |
| AB | AB | AB | AA | AB | AB | AA | AB | AB | AB | AB | AA | AB | AB | AA | AB | AB |
| AB | AA | AA | AB | AA | AA | AA | AB | AB | AB | AB | AA | AA | AB | AB | AA | AB |
| AB | AB | AB | AA | AA | AA | AA | AA | AB | AA | AB | AA | AB | - | - | AA | AA |
| AA | AA | AB | AB | AB | AB | AB | AB | AB | AA | AB | AA | AB | AB | AA | AA | AB |
| AA | AA | AB | AA | AB | AB | AB | AA | AB | AA | AB | AA | AB | AA | AA | AB | AA |
| AB | AA | AA | AA | AA | AA | AA | AA | AB | AA | AA | AA | AA | AB | AA | AA | AA |
| AA | AB | AA | AA | AA | AA | AA | AB | AA | AA | AA | AA | AA | AA | - | AA | AB |
| AA | AB | AB | AA | AA | AB | AB | AB | AA | AB | AA | AB | AA | AA | AA | AB | AA |
| AB | AB | AA | AA | AA | AB | AA | AA | AA | AA | AA | AA | AB | AA | AA | AA | AB |
| AA | AA | AB | AB | AB | AB | AB | AB | AB | AB | AB | AB | AB | AB | AB | AB | AB |
| AB | AA | AA | AB | AB | AA | AA | AB | AB | AB | AB | AA | AB | AB | AB | AA | AB |
| AA | AA | AB | AA | AA | AA | AA | AB | AA | AA | AB | AA | AB | AA | AB | AA | AB |
| AA | AB | AB | AB | AB | AB | AB | AB | AB | AA | AB | AA | AB | AB | AA | AA | AB |
| AA | - | AB | AA | AA | AB | AA | AA | AA | AA | AB | AB | AB | AA | AA | AB | AA |
| [illegible] | | | | | | | | | | | | | | | | |

| snp_B2001 | snp_B2016 | snp_B2019 | snp_B2020 | snp_B2021 | snp_B2024 | snp_B2033 | snp_B2060 | snp_B2091 | snp_B2097 | snp_B2099 | snp_B2116 | snp_B2125 | snp_B2197 | snp_B2199 | snp_B2200 | snp_B2202 |
|---|---|---|---|---|---|---|---|---|---|---|---|---|---|---|---|---|
| 3 | 3 | 3 | 3 | 3 | 3 | 3 | 3 | 3 | 3 | 3 | 3 | 3 | 3 | 3 | 3 | 3 |
| -- | -- | LG9 | -- | -- | LG9 | -- | -- | -- | -- | LG9 | -- | -- | -- | -- | -- | -- |
| AA | AB | AA | AB | AA | AA | AB | AB | AB | AA | AB | AA | AA | AB | AA | AA | AB |
| AB | AA | AA | AB | AB | AB | AA | AB | AB | AA | AA | AB | AA | AA | AA | AB | AA |
| AA | AB | AA | AB | AA | AA | AB | AB | AB | AA | AB | AA | AA | AB | AA | AA | AA |
| AA | AA | AB | AA | AA | AB | AA | AA | AA | AB | AA | AB | AB | AA | AA | AB | AA |
| AB | AA | AB | AB | AB | AB | AA | AA | AA | AA | AB | AA | AA | AB | AB | AA | AA |
| AA | - | AA | - | AA | - | AA | AA | AA | - | - | AB | - | AB | AA | AA | - |
| AA | AA | AA | AA | AA | AA | AA | AA | AA | AA | AA | AA | AA | AA | AA | AB | AA |
| AB | AB | AB | AB | AA | AB | AB | AB | AB | AB | AB | AB | AB | AB | AB | AB | AB |
| AA | AB | AA | AB | AA | AB | AB | AA | AA | AA | AB | AB | AA | AB | AB | AA | AB |
| AA | AA | AA | AB | AB | AB | AB | AA | AB | AB | AA | AB | AA | AB | AB | AA | AB |
| AB | AA | AA | AB | AA | AA | AB | AB | AA | AB | AB | AA | AA | AB | AB | AA | AB |
| AA | AA | AA | AB | AA | AA | AB | AB | AA | AA | AB | AB | AA | AB | AB | AA | AA |
| AA | AA | AA | AA | AA | AA | AA | AA | AB | AA | AA | AA | AA | AA | AB | AB | AA |
| AA | AA | AA | AA | AB | AB | AA | AA | AB | AB | AA | AA | AA | AA | AB | AB | AA |
| AA | AB | AA | AB | AB | AA | AB | AB | AB | AA | AB | AB | AA | AB | AA | AA | AB |
| AB | AB | AB | AB | AB | AB | AB | AB | AB | AB | AB | AB | AB | AB | AB | AB | AB |
| AA | AB | AB | AB | AA | AA | AB | AA | AA | AB | AB | AB | AB | AB | AB | AB | AB |
| AA | - | - | - | AB | - | - | - | - | - | - | AB | - | AB | AA | AA | - |
| AA | AB | AA | AA | AA | AA | AB | AB | AA | AA | AB | AA | AA | AA | AA | AB | AB |
| AB | AA | AB | AB | AA | AA | AA | AB | AB | - | AA | AB | - | AA | AA | AB | AA |
| AB | AB | AB | AB | AB | AB | AB | AA | AB | AB | AB | AB | AB | AB | AB | AB | AB |
| AB | AB | AB | AA | AA | AB | AA | AA | AA | AB | AB | AB | AA | AB | AB | AB | AA |
| AA | AA | - | - | AA | - | - | AA | AB | - | AA | AA | - | - | AA | AB | AA |
| AA | AA | AA | AB | AB | AA | AA | AB | AB | AB | AB | AA | AA | AA | AA | AA | AA |
| AB | AA | AB | AA | AA | AB | AB | AA | AA | AB | AA | AB | AB | AA | AB | AB | AB |
| AB | AA | AA | AA | AA | AB | AA | AA | AA | AB | AA | AB | AB | AA | AA | AB | AA |
| AA | AB | AA | AA | AA | AA | AB | AB | AA | AA | AB | AB | AA | AB | AA | AA | AB |
| AB | AA | AA | AB | AA | AA | AA | AB | AB | AA | AB | AA | AA | AA | AA | AA | AA |
| AB | AB | - | AB | AA | AA | AB | AA | AB | AB | AB | AB | - | AB | AA | AA | AB |
| AA | - | - | - | - | - | AA | AB | AA | - | - | AB | - | AA | AA | AA | - |
| AB | AA | AA | AA | AB | AB | AA | AA | AA | AA | AA | AA | AA | AA | AA | AB | AA |
| AB | AB | AB | AB | AB | AA | AB | AA | AB | AA | AB | AA | AB | AB | AB | AA | AB |
| AB | AA | - | - | AB | - | AA | AA | AB | AA | - | AA | AA | AA | AA | AA | AA |
| AB | AA | AB | AB | AA | AB | - | - | AB | - | - | AA | AA | AA | AB | AB | AA |
| AA | - | - | AA | AA | - | AA | AA | - | AA | - | AA | AA | AB | AB | AB | AA |
| AA | AB | - | AB | AB | AA | AB | AB | AB | AB | AA | AB | - | AA | AA | AB | AB |
| AB | AA | AB | AB | AB | AA | AA | AB | AA | AA | AA | AB | AA | AA | AA | AB | AA |
| AB | AB | AB | AB | AB | AB | AB | AB | AB | AB | AB | AB | AB | AB | AB | AB | AA |
| AB | AB | AB | AA | AB | AB | AA | AA | AB | AB | AA | AA | AB | AA | AA | AB | AB |
| AB | AA | - | AA | AA | AA | AA | - | AA | AA | - | AA | AA | AA | AA | AA | AA |
| AA | AA | AB | AB | AB | AA | AB | AB | AB | AB | AB | AA | AA | AB | AA | AA | AA |
| AB | AB | AB | AA | AB | AA | AB | AA | AA | AA | AB | AA | AB | AB | AB | AA | AB |
| AA | AA | AA | AA | AB | AA | AA | AA | AA | AB | AA | AB | AA | AA | AA | AB | AA |
| AB | AA | AA | - | AB | AA | AA | AB | AB | AA | - | AA | - | AA | AA | AA | AA |
| AB | AB | AA | AB | AA | AA | AB | AB | AB | AA | AB | AB | AA | AB | AB | AA | AB |
| AA | AB | AB | AA | AB | AB | AA | AB | AA | AA | AA | AB | AA | AA | AB | AB | AB |

| snp_B3005 | snp_B3052 | snp_B3065 | snp_B3070 | snp_B3081 | snp_B3087 | snp_B3100 | snp_B3133 | snp_B3139 | snp_B3143 | snp_B3168 | snp_B3169 | snp_B3213 | snp_B3218 | snp_B3268 | snp_B3270 | snp_B3301 | snp_B3303 | snp_B3309 | snp_B3312 | snp_B3322 | snp_B3324 | snp_B3328 | snp_B3340 | snp_B3343 | snp_B3356 | snp_B3364 | snp_B3388 | snp_B3411 | snp_B3414 | snp_B3417 | snp_B3424 | snp_B3426 | snp_B3437 | snp_B3450 | snp_B3457 | snp_B3467 | snp_B3486 | snp_B3494 | snp_B3505 | snp_B3514 | snp_B3532 | snp_B3548 | snp_B3575 | snp_B3594 | snp_B3596 | snp_B3598 | snp_B3626 | snp_B3633 | snp_B3640 | snp_B3662 | snp_B3684 | snp_B3694 | snp_B3698 | snp_B3707 | snp_B3751 | snp_B3752 | snp_B3761 | snp_B3777 | snp_B3782 | snp_B3792 | snp_B3797 | snp_B3815 | snp_B3819 | snp_B3827 | snp_B3842 | snp_B3848 | snp_B3857 | snp_B3861 | snp_B3867 | snp_B3870 | snp_B3875 | snp_B3922 | snp_B3936 | snp_B3949 | snp_B3951 | snp_B3958 |
|---|---|---|---|---|---|---|---|---|---|---|---|---|---|---|---|---|---|---|---|---|---|---|---|---|---|---|---|---|---|---|---|---|---|---|---|---|---|---|---|---|---|---|---|---|---|---|---|---|---|---|---|---|---|---|---|---|---|---|---|---|---|---|---|---|---|---|---|---|---|---|---|---|---|---|---|---|
| 3 | 3 | 3 | 3 | 3 | 3 | 3 | 3 | 3 | 3 | 3 | 3 | 3 | 3 | 3 | 3 | 3 | 3 | 3 | 3 | 3 | 3 | 3 | 3 | 3 | 3 | 3 | 3 | 3 | 3 | 3 | 3 | 3 | 3 | 3 | 3 | 3 | 3 | 3 | 3 | 3 | 3 | 3 | 3 | 3 | 3 | 3 | 3 | 3 | 3 | 3 | 3 | 3 | 3 | 3 | 3 | 3 | 3 | 3 | 3 | 3 | 3 | 3 | 3 | 3 | 3 | 3 | 3 | 3 | 3 | 3 | 3 | 3 | 3 | 3 | 3 | 3 |
| -- | LG9 | -- | -- | -- | LG9 | -- | -- | -- | -- | -- | -- | -- | -- | -- | LG9 | -- | -- | -- | -- | -- | -- | -- | -- | -- | -- | -- | -- | -- | -- | -- | -- | -- | -- | -- | -- | -- | -- | -- | -- | -- | -- | -- | -- | -- | -- | -- | -- | -- | -- | -- | LG9 | -- | -- | -- | -- | -- | -- | -- | -- | -- | -- | -- | -- | -- | -- | -- | -- | -- | -- | -- | -- | -- | -- | LG9 | -- | -- |
| [illegible] | [illegible] | [illegible] | [illegible] | [illegible] | [illegible] | [illegible] | [illegible] | [illegible] | [illegible] | [illegible] | [illegible] | [illegible] | [illegible] | [illegible] | [illegible] | [illegible] | [illegible] | [illegible] | [illegible] | [illegible] | [illegible] | [illegible] | [illegible] | [illegible] | [illegible] | [illegible] | [illegible] | [illegible] | [illegible] | [illegible] | [illegible] | [illegible] | [illegible] | [illegible] | [illegible] | [illegible] | [illegible] | [illegible] | [illegible] | [illegible] | [illegible] | [illegible] | [illegible] | [illegible] | [illegible] | [illegible] | [illegible] | [illegible] | [illegible] | [illegible] | [illegible] | [illegible] | [illegible] | [illegible] | [illegible] | [illegible] | [illegible] | [illegible] | [illegible] | [illegible] | [illegible] | [illegible] | [illegible] | [illegible] | [illegible] | [illegible] | [illegible] | [illegible] | [illegible] | [illegible] | [illegible] | [illegible] | [illegible] | [illegible] | [illegible] | [illegible] |

| snp_B3959 | snp_B3962 | snp_B3966 | snp_B3979 | snp_B3989 | snp_B4014 | snp_B4051 | snp_B4104 | snp_B4122 | snp_B4129 | snp_B4136 | snp_B4143 | snp_B4150 | snp_B4160 | snp_B4175 | snp_B4179 | snp_B4239 | snp_B4279 | snp_B4293 | snp_B4295 | snp_B4315 | snp_B4321 | snp_B4327 | snp_B4362 | snp_B4372 | snp_B4374 | snp_B4376 | snp_B4379 | snp_B4384 | snp_B4391 | snp_B4404 | snp_B4421 | snp_B4458 | snp_B4475 | snp_B4488 | snp_B4491 | snp_B4503 | snp_B4540 | snp_B4553 | snp_B4565 | snp_B4576 | snp_B4598 | snp_B4622 | snp_B4629 | snp_B4642 | snp_B4657 | snp_B4726 | snp_B4755 | snp_B4768 | snp_B4810 | snp_B4811 | snp_B4815 | snp_B4826 | snp_B4849 | snp_B4854 | snp_B4899 | snp_B4902 | snp_B4920 | snp_B4921 | snp_B4949 | snp_B4955 | snp_B4961 | snp_B4995 | snp_B5002 | snp_B5010 | snp_B5011 | snp_B5020 | snp_B5030 | snp_B5053 | snp_B5058 | snp_B5062 | snp_B5068 | snp_B5071 | snp_B5073 | snp_B5081 | snp_B5108 | snp_B5117 |
|---|---|---|---|---|---|---|---|---|---|---|---|---|---|---|---|---|---|---|---|---|---|---|---|---|---|---|---|---|---|---|---|---|---|---|---|---|---|---|---|---|---|---|---|---|---|---|---|---|---|---|---|---|---|---|---|---|---|---|---|---|---|---|---|---|---|---|---|---|---|---|---|---|---|---|---|---|
| 3 | 3 | 3 | 3 | 3 | 3 | 3 | 3 | 3 | 3 | 3 | 3 | 3 | 3 | 3 | 3 | 3 | 3 | 3 | 3 | 3 | 3 | 3 | 3 | 3 | 3 | 3 | 3 | 3 | 3 | 3 | 3 | 3 | 3 | 3 | 3 | 3 | 3 | 3 | 3 | 3 | 3 | 3 | 3 | 3 | 3 | 3 | 3 | 3 | 3 | 3 | 3 | 3 | 3 | 3 | 3 | 3 | 3 | 3 | 3 | 3 | 3 | 3 | 3 | 3 | 3 | 3 | 3 | 3 | 3 | 3 | 3 | 3 | 3 | 3 | 3 | 3 |
| -- | -- | -- | -- | -- | -- | -- | -- | -- | -- | -- | LG9 | -- | -- | -- | -- | -- | -- | -- | -- | -- | -- | -- | -- | -- | -- | LG9 | -- | -- | -- | -- | -- | -- | -- | -- | -- | -- | -- | -- | -- | -- | -- | -- | -- | -- | -- | -- | -- | LG9 | -- | -- | -- | -- | LG9 | -- | -- | -- | -- | -- | -- | -- | -- | -- | -- | -- | -- | -- | -- | -- | -- | LG9 | -- | -- | LG9 | -- | -- | -- |

| snp_B5150 | snp_B5152 | snp_B5153 | snp_B5178 | snp_B5198 | snp_B5200 | snp_B5210 | snp_B5216 | snp_B5221 | snp_B5247 | snp_B5248 | snp_B5263 | snp_B5268 | snp_B5292 | snp_B5300 | snp_B5308 | snp_B5339 | snp_B5347 | snp_B5388 | snp_B5400 |
|---|---|---|---|---|---|---|---|---|---|---|---|---|---|---|---|---|---|---|---|
| 3 | 3 | 3 | 3 | 3 | 3 | 3 | 3 | 3 | 3 | 3 | 3 | 3 | 3 | 3 | 3 | 3 | 3 | 3 | 3 |
| -- | -- | -- | -- | -- | -- | -- | -- | -- | -- | -- | -- | -- | -- | -- | -- | -- | -- | -- | -- |
| AB | AB | AB | AB | AB | AB | AB | AA | AB | AA | AB | AA | AB | AA | AA | AB | AA | AA | AA | AA |
| AB | AA | AA | AB | AA | AB | AB | AB | AB | AB | AA | AA | AB | AA | AB | AA | AB | AA | AB | AA |
| AB | AA | AB | AB | AB | AB | AB | AA | AB | AA | AB | AA | AB | AB | AA | AA | AA | AA | AA | AA |

| snp_B5424 | snp_B5428 | snp_B5448 | snp_B5456 | snp_B5474 | snp_B5518 | snp_B5521 | snp_B5560 | snp_B5570 | snp_B5573 | snp_B5575 | snp_B5581 | snp_B5588 | snp_B5600 | snp_B5613 | snp_B5626 | snp_B5648 | snp_B5653 | snp_B5697 |
|---|---|---|---|---|---|---|---|---|---|---|---|---|---|---|---|---|---|---|
| 3 | 3 | 3 | 3 | 3 | 3 | 3 | 3 | 3 | 3 | 3 | 3 | 3 | 3 | 3 | 3 | 3 | 3 | 3 |
| -- | -- | -- | -- | -- | -- | -- | -- | -- | -- | -- | -- | LG9 | -- | -- | -- | -- | -- | -- |
| AB | AB | AA | AB | AB | AA | AA | AB | AA | AB | AB | AB | AA | AA | AB | AA | AA | AB | AA |
| AA | AA | AA | AB | AB | AB | AA | AA | AA | AB | AA | AB | AA | AA | AA | AA | AB | AA | AA |
| AA | AB | AA | AB | AA | AA | AA | AA | AB | AA | AA | AA | AA | AB | AB | AA | AA | AB | AA |

| snp_B5704 | snp_B5736 | snp_B5738 | snp_B5740 | snp_B5752 | snp_B5766 | snp_B5793 | snp_B5794 | snp_B5862 | snp_B5929 | snp_B5942 | snp_B5963 | snp_B5984 | snp_B6009 | snp_B6020 | snp_B6152 | snp_B6155 | snp_B0010 | snp_B0011 |
|---|---|---|---|---|---|---|---|---|---|---|---|---|---|---|---|---|---|---|
| 3 | 3 | 3 | 3 | 3 | 3 | 3 | 3 | 3 | 3 | 3 | 3 | 3 | 3 | 3 | 3 | 3 | 4 | 4 |
| -- | -- | -- | -- | -- | -- | LG9 | -- | -- | -- | LG9 | -- | -- | -- | -- | -- | -- | -- | -- |
| AA | AB | AB | AA | AB | AB | AA | AB | AA | AB | AA | AA | AA | AA | AA | AA | AA | AB | AB |
| AA | AA | AA | AA | AB | AA | AA | AA | AB | AB | AB | AA | AA | AA | AB | AA | AB | AB | AB |
| AB | AB | AB | AA | AB | AB | AA | AB | AA | AB | AA | AA | AA | AA | AB | AB | AA | AB | AA |

| snp_B0030 | snp_B0068 | snp_B0110 | snp_B0165 | snp_B0172 | snp_B0178 | snp_B0211 | snp_B0220 | snp_B0221 | snp_B0230 | snp_B0246 | snp_B0263 | snp_B0276 | snp_B0290 | snp_B0377 | snp_B0396 | snp_B0445 | snp_B0450 | snp_B0503 |
|---|---|---|---|---|---|---|---|---|---|---|---|---|---|---|---|---|---|---|
| 4 | 4 | 4 | 4 | 4 | 4 | 4 | 4 | 4 | 4 | 4 | 4 | 4 | 4 | 4 | 4 | 4 | 4 | 4 |
| -- | -- | -- | -- | -- | -- | -- | -- | -- | -- | -- | -- | -- | -- | -- | -- | -- | -- | -- |
| AA | AB | AA | AA | AA | AA | AA | AB | AB | AA | AA | AB | AA | AA | AB | AA | AA | AB | AA |
| AB | AB | AB | AA | AB | AB | AB | AA | AA | AB | AB | AB | AB | AB | AA | AB | AB | AB | AB |
| AB | AB | AA | AA | AA | AB | AA | AA | AA | AB | AA | AB | AA | AB | AB | AB | AA | AB | AB |

| snp_B0531 | snp_B0563 | snp_B0567 | snp_B0579 | snp_B0591 | snp_B0607 | snp_B0644 | snp_B0651 | snp_B0701 | snp_B0759 | snp_B0767 | snp_B0769 | snp_B0797 | snp_B0799 | snp_B0820 | snp_B0835 | snp_B0846 | snp_B0852 | snp_B0853 | snp_B0870 | snp_B0879 | snp_B0885 | snp_B0894 | snp_B0924 | snp_B0926 | snp_B0930 | snp_B0947 | snp_B0949 | snp_B0959 | snp_B0960 | snp_B0994 | snp_B1004 | snp_B1007 | snp_B1029 | snp_B1034 | snp_B1048 | snp_B1050 | snp_B1051 | snp_B1054 | snp_B1063 | snp_B1070 | snp_B1098 | snp_B1109 | snp_B1128 | snp_B1143 | snp_B1152 | snp_B1163 | snp_B1179 | snp_B1191 | snp_B1216 | snp_B1224 | snp_B1230 | snp_B1258 | snp_B1280 | snp_B1301 | snp_B1302 | snp_B1338 | snp_B1360 | snp_B1365 | snp_B1375 | snp_B1380 | snp_B1395 | snp_B1398 | snp_B1414 | snp_B1445 | snp_B1458 | snp_B1463 | snp_B1468 | snp_B1491 | snp_B1498 | snp_B1506 | snp_B1604 | snp_B1611 | snp_B1612 | snp_B1627 | snp_B1637 | snp_B1639 |
|---|---|---|---|---|---|---|---|---|---|---|---|---|---|---|---|---|---|---|---|---|---|---|---|---|---|---|---|---|---|---|---|---|---|---|---|---|---|---|---|---|---|---|---|---|---|---|---|---|---|---|---|---|---|---|---|---|---|---|---|---|---|---|---|---|---|---|---|---|---|---|---|---|---|---|---|---|
| 4 | 4 | 4 | 4 | 4 | 4 | 4 | 4 | 4 | 4 | 4 | 4 | 4 | 4 | 4 | 4 | 4 | 4 | 4 | 4 | 4 | 4 | 4 | 4 | 4 | 4 | 4 | 4 | 4 | 4 | 4 | 4 | 4 | 4 | 4 | 4 | 4 | 4 | 4 | 4 | 4 | 4 | 4 | 4 | 4 | 4 | 4 | 4 | 4 | 4 | 4 | 4 | 4 | 4 | 4 | 4 | 4 | 4 | 4 | 4 | 4 | 4 | 4 | 4 | 4 | 4 | 4 | 4 | 4 | 4 | 4 | 4 | 4 | 4 | 4 | 4 | 4 |
| -- | -- | -- | -- | -- | -- | -- | -- | -- | LG6 | -- | -- | -- | -- | -- | LG6 | -- | -- | -- | -- | -- | -- | -- | -- | -- | -- | -- | -- | -- | -- | -- | -- | -- | -- | -- | -- | -- | -- | -- | -- | -- | -- | -- | -- | -- | -- | -- | -- | -- | -- | -- | -- | -- | -- | -- | -- | -- | -- | -- | -- | -- | -- | -- | -- | -- | -- | -- | -- | -- | LG6 | -- | LG6 | -- | -- | LG6 | -- | -- |
| [illegible] | | | | | | | | | | | | | | | | | | | | | | | | | | | | | | | | | | | | | | | | | | | | | | | | | | | | | | | | | | | | | | | | | | | | | | | | | | | | |

| snp_B1647 | snp_B1667 | snp_B1674 | snp_B1689 | snp_B1705 | snp_B1743 | snp_B1747 | snp_B1762 | snp_B1767 | snp_B1814 | snp_B1848 | snp_B1849 | snp_B1854 | snp_B1882 | snp_B1923 | snp_B1938 | snp_B2026 | snp_B2042 | snp_B2052 | snp_B2055 | snp_B2076 | snp_B2108 | snp_B2140 | snp_B2142 | snp_B2155 |
|---|---|---|---|---|---|---|---|---|---|---|---|---|---|---|---|---|---|---|---|---|---|---|---|---|
| 4 | 4 | 4 | 4 | 4 | 4 | 4 | 4 | 4 | 4 | 4 | 4 | 4 | 4 | 4 | 4 | 4 | 4 | 4 | 4 | 4 | 4 | 4 | 4 | 4 |
| -- | -- | -- | -- | LG6 | -- | -- | -- | -- | -- | -- | -- | -- | -- | -- | -- | -- | -- | -- | -- | LG6 | -- | -- | -- | -- |
| AB | AA | AB | AB | AA | AA | AB | AA | AB | AA | AB | AA | AB | AA | AA | AA | AB | AA | AB | AA | AA | AA | AB | AB | AA |
| [illegible] | [illegible] | [illegible] | [illegible] | [illegible] | [illegible] | [illegible] | [illegible] | [illegible] | [illegible] | [illegible] | [illegible] | [illegible] | [illegible] | [illegible] | [illegible] | [illegible] | [illegible] | [illegible] | [illegible] | [illegible] | [illegible] | [illegible] | [illegible] | [illegible] |

| snp_B2212 | snp_B2224 | snp_B2228 | snp_B2267 | snp_B2300 | snp_B2301 | snp_B2303 | snp_B2312 | snp_B2330 | snp_B2362 | snp_B2426 | snp_B2469 | snp_B2474 | snp_B2496 | snp_B2513 | snp_B2520 | snp_B2522 | snp_B2562 | snp_B2606 | snp_B2612 | snp_B2614 | snp_B2640 | snp_B2641 | snp_B2659 | snp_B2678 | snp_B2679 |
|---|---|---|---|---|---|---|---|---|---|---|---|---|---|---|---|---|---|---|---|---|---|---|---|---|---|
| 4 | 4 | 4 | 4 | 4 | 4 | 4 | 4 | 4 | 4 | 4 | 4 | 4 | 4 | 4 | 4 | 4 | 4 | 4 | 4 | 4 | 4 | 4 | 4 | 4 | 4 |
| -- | -- | -- | -- | -- | -- | -- | -- | -- | -- | -- | -- | -- | -- | -- | -- | LG6 | -- | -- | -- | -- | -- | -- | -- | -- | -- |
| [illegible] | [illegible] | [illegible] | [illegible] | [illegible] | [illegible] | [illegible] | [illegible] | [illegible] | [illegible] | [illegible] | [illegible] | [illegible] | [illegible] | [illegible] | [illegible] | [illegible] | [illegible] | [illegible] | [illegible] | [illegible] | [illegible] | [illegible] | [illegible] | [illegible] | [illegible] |

| snp_B2682 | snp_B2734 | snp_B2737 | snp_B2777 | snp_B2785 | snp_B2809 | snp_B2826 | snp_B2884 | snp_B2887 | snp_B2891 | snp_B2908 | snp_B2915 | snp_B2947 | snp_B2964 | snp_B2969 | snp_B2983 | snp_B2987 | snp_B2997 | snp_B3024 | snp_B3028 | snp_B3046 | snp_B3053 | snp_B3064 | snp_B3082 | snp_B3090 | snp_B3104 |
|---|---|---|---|---|---|---|---|---|---|---|---|---|---|---|---|---|---|---|---|---|---|---|---|---|---|
| 4 | 4 | 4 | 4 | 4 | 4 | 4 | 4 | 4 | 4 | 4 | 4 | 4 | 4 | 4 | 4 | 4 | 4 | 4 | 4 | 4 | 4 | 4 | 4 | 4 | 4 |
| -- | -- | -- | -- | -- | -- | -- | -- | -- | -- | -- | LG6 | -- | -- | -- | -- | -- | -- | -- | -- | -- | -- | -- | -- | -- | -- |
| AB | AB | AA | AB | AB | AA | AB | AA | AB | AB | AB | AA | AB | AB | AA | AB | AB | AA | AB | AA | AB | AB | AA | AB | AB | AB |
| [illegible] | [illegible] | [illegible] | [illegible] | [illegible] | [illegible] | [illegible] | [illegible] | [illegible] | [illegible] | [illegible] | [illegible] | [illegible] | [illegible] | [illegible] | [illegible] | [illegible] | [illegible] | [illegible] | [illegible] | [illegible] | [illegible] | [illegible] | [illegible] | [illegible] | [illegible] |

| snp_B3109 | snp_B3128 | snp_B3141 | snp_B3146 | snp_B3154 | snp_B3167 | snp_B3175 | snp_B3207 | snp_B3216 | snp_B3225 | snp_B3227 | snp_B3253 | snp_B3259 | snp_B3265 | snp_B3304 | snp_B3333 | snp_B3341 | snp_B3348 | snp_B3365 | snp_B3370 | snp_B3387 | snp_B3390 | snp_B3397 | snp_B3429 | snp_B3430 | snp_B3440 | snp_B3466 | snp_B3477 | snp_B3479 | snp_B3481 | snp_B3492 | snp_B3501 | snp_B3507 | snp_B3516 | snp_B3526 | snp_B3551 | snp_B3557 | snp_B3560 | snp_B3587 | snp_B3595 | snp_B3613 | snp_B3622 | snp_B3647 | snp_B3659 | snp_B3676 | snp_B3765 | snp_B3767 | snp_B3768 | snp_B3790 | snp_B3795 | snp_B3803 | snp_B3810 | snp_B3823 | snp_B3852 | snp_B3891 | snp_B3900 | snp_B3909 | snp_B3925 | snp_B3931 | snp_B3932 | snp_B3976 | snp_B3985 | snp_B3986 | snp_B4005 | snp_B4024 | snp_B4042 | snp_B4043 | snp_B4053 | snp_B4069 | snp_B4070 | snp_B4071 | snp_B4077 | snp_B4092 | snp_B4113 | snp_B4126 | snp_B4156 | snp_B4157 |
|---|---|---|---|---|---|---|---|---|---|---|---|---|---|---|---|---|---|---|---|---|---|---|---|---|---|---|---|---|---|---|---|---|---|---|---|---|---|---|---|---|---|---|---|---|---|---|---|---|---|---|---|---|---|---|---|---|---|---|---|---|---|---|---|---|---|---|---|---|---|---|---|---|---|---|---|---|
| 4 | 4 | 4 | 4 | 4 | 4 | 4 | 4 | 4 | 4 | 4 | 4 | 4 | 4 | 4 | 4 | 4 | 4 | 4 | 4 | 4 | 4 | 4 | 4 | 4 | 4 | 4 | 4 | 4 | 4 | 4 | 4 | 4 | 4 | 4 | 4 | 4 | 4 | 4 | 4 | 4 | 4 | 4 | 4 | 4 | 4 | 4 | 4 | 4 | 4 | 4 | 4 | 4 | 4 | 4 | 4 | 4 | 4 | 4 | 4 | 4 | 4 | 4 | 4 | 4 | 4 | 4 | 4 | 4 | 4 | 4 | 4 | 4 | 4 | 4 | 4 | 4 |
| -- | -- | -- | -- | -- | -- | -- | -- | -- | -- | -- | -- | -- | -- | -- | LG6 | -- | -- | -- | -- | -- | -- | -- | -- | LG6 | -- | -- | -- | -- | -- | -- | -- | -- | -- | -- | -- | -- | -- | -- | -- | -- | -- | -- | -- | -- | -- | -- | -- | -- | -- | LG6 | -- | -- | LG6 | -- | -- | -- | -- | -- | -- | -- | -- | -- | -- | -- | -- | -- | -- | -- | -- | -- | -- | -- | -- | -- | LG6 | -- |
| AA | AA | AA | AB | AB | AB | AA | AB | AB | AB | AA | AB | AB | AA | AA | AA | AB | AA | AB | AB | AA | AB | AA | AB | AA | AA | AB | AB | AA | AA | AA | AA | AB | AB | AB | AB | AA | AB | AA | AA | AB | AA | AA | AA | AA | AA | AA | AB | AB | AB | AA | AB | AB | AA | AA | AA | AB | AA | AA | AB | AB | AA | AA | AA | AA | AB | AA | AA | AA | AA | AA | AA | AA | AB | AA | AA | AA |
| AA | AB | AB | AA | AA | AA | AB | AB | AA | AB | AB | AB | AB | AB | AB | AB | AA | AA | AA | AB | AB | AA | AB | AA | AB | AB | AA | AA | AB | AB | AB | AB | AB | AB | AB | AA | AB | AB | AB | AB | AB | AB | AA | AA | AB | AB | AB | AB | AA | AB | AB | AA | AA | AB | AB | AA | AA | AA | AB | AB | AB | AB | AB | AA | AB | AB | AB | AB | AA | AB | AB | AB | AB | AB | AB | AB | AB |
| AB | AB | AA | AA | AB | AB | AA | AA | AB | AB | AB | AA | AB | AB | AB | AA | AB | AB | AA | AB | AB | AA | AB | AA | AA | AB | AB | AB | AA | AB | AA | AB | AA | AB | AA | AA | AA | AB | AB | AB | AA | AB | AB | AB | AA | AA | AB | AA | AB | AB | AA | AB | AB | AB | AA | AB | AB | AB | AB | AB | AA | AA | AA | AA | AA | AA | AA | AB | AA | AB | AB | AB | AB | AA | AB | AA | AA |
| AA | AB | AB | AA | AA | AB | AB | - | AB | AB | AB | AB | AB | AB | AA | AB | AA | AA | AA | AA | AA | AA | AB | AA | AB | AB | AB | AA | AA | AB | AB | AB | AB | AB | AB | AA | AB | AB | AB | AB | AB | AB | - | AA | AB | - | AB | AB | - | AB | AB | - | AA | AB | AB | AA | AB | AA | AB | AB | AB | AB | AB | AA | - | AB | AB | AB | AA | AB | AB | AB | AB | AB | AB | AB | AB |
| AB | AA | AB | AA | AB | AB | AA | AA | AA | AA | AA | AB | AA | AA | AA | AA | AB | AA | AA | AB | AB | AB | AA | AB | AA | AA | AA | AA | AA | AA | AA | AA | AA | AA | AA | AA | AA | AA | AA | AA | AA | AA | AA | AB | AA | AB | AA | AA | AA | AA | AA | AA | AB | AA | AB | AB | AA | AA | AA | AA | AA | AA | AA | AA | AA | AA | AA | AA | AA | AA | AA | AA | AA | AA | AA | AA | AA |

| snp_B4158 | snp_B4163 | snp_B4174 | snp_B4177 | snp_B4190 | snp_B4209 | snp_B4211 | snp_B4216 | snp_B4217 | snp_B4219 | snp_B4222 | snp_B4231 | snp_B4232 | snp_B4234 | snp_B4248 | snp_B4263 | snp_B4270 | snp_B4289 | snp_B4318 | snp_B4326 | snp_B4331 | snp_B4334 | snp_B4338 | snp_B4343 | snp_B4344 | snp_B4365 |
|---|---|---|---|---|---|---|---|---|---|---|---|---|---|---|---|---|---|---|---|---|---|---|---|---|---|
| 4 | 4 | 4 | 4 | 4 | 4 | 4 | 4 | 4 | 4 | 4 | 4 | 4 | 4 | 4 | 4 | 4 | 4 | 4 | 4 | 4 | 4 | 4 | 4 | 4 | 4 |
| -- | -- | LG6 | -- | -- | -- | -- | -- | -- | -- | -- | -- | -- | -- | -- | -- | -- | -- | -- | -- | -- | LG6 | -- | -- | -- | -- |
| AA | AB | AA | AA | AB | AB | AB | AB | AB | AB | AA | AB | AB | AA | AA | AB | AB | AA | AA | AB | AA | AA | AB | AB | AA | AA |
| AA | AB | AB | AB | AB | AB | AA | AB | AA | AB | AB | AA | AA | AA | AA | AA | AB | AA | AA | AA | AB | AB | AB | AA | AB | AA |
| AA | AA | AA | AA | AA | AA | AA | AA | AB | AB | AB | AA | AA | AA | AB | AB | AA | AA | AA | AB | AB | AA | AB | AA | AA | AA |

| snp_B4370 | snp_B4371 | snp_B4400 | snp_B4408 | snp_B4409 | snp_B4428 | snp_B4433 | snp_B4438 | snp_B4450 | snp_B4453 | snp_B4456 | snp_B4459 | snp_B4462 | snp_B4494 | snp_B4496 | snp_B4511 | snp_B4570 | snp_B4579 | snp_B4582 | snp_B4597 | snp_B4627 | snp_B4654 | snp_B4675 | snp_B4691 | snp_B4705 |
|---|---|---|---|---|---|---|---|---|---|---|---|---|---|---|---|---|---|---|---|---|---|---|---|---|
| 4 | 4 | 4 | 4 | 4 | 4 | 4 | 4 | 4 | 4 | 4 | 4 | 4 | 4 | 4 | 4 | 4 | 4 | 4 | 4 | 4 | 4 | 4 | 4 | 4 |
| -- | LG6 | -- | -- | -- | -- | -- | -- | -- | -- | -- | -- | -- | -- | -- | -- | -- | -- | -- | -- | -- | -- | -- | -- | -- |
| AB | AA | AA | AB | AB | AA | AB | AB | AA | AB | AB | AA | AA | AA | AB | AA | AB | AB | AA | AA | AA | AB | AB | AA | AA |
| AB | AB | AB | AA | AB | AB | AB | AB | AB | AA | AA | AB | AB | AB | AB | AB | AB | AB | AB | AB | AB | AB | AA | AB | AB |
| AB | AB | AB | AB | AA | AB | AB | AB | AB | AA | AB | AB | AB | AB | AB | AA | AA | AB | AB | AB | AB | AB | AB | AA | AB |

| snp_B4728 | snp_B4734 | snp_B4739 | snp_B4742 | snp_B4773 | snp_B4789 | snp_B4800 | snp_B4807 | snp_B4837 | snp_B4858 | snp_B4893 | snp_B4925 | snp_B4935 | snp_B4947 | snp_B5007 | snp_B5008 | snp_B5009 | snp_B5029 | snp_B5031 | snp_B5042 | snp_B5046 | snp_B5048 | snp_B5052 | snp_B5055 | snp_B5063 | snp_B5076 |
|---|---|---|---|---|---|---|---|---|---|---|---|---|---|---|---|---|---|---|---|---|---|---|---|---|---|
| 4 | 4 | 4 | 4 | 4 | 4 | 4 | 4 | 4 | 4 | 4 | 4 | 4 | 4 | 4 | 4 | 4 | 4 | 4 | 4 | 4 | 4 | 4 | 4 | 4 | 4 |
| -- | -- | -- | -- | -- | -- | -- | -- | -- | -- | -- | -- | -- | LG6 | -- | -- | -- | -- | -- | -- | -- | -- | -- | -- | -- | -- |
| AA | AB | AA | AA | AB | AB | AB | AB | AA | AA | AB | AB | AA | AA | AB | AB | AA | AB | AB | AA | AB | AA | AA | AB | AB | AB |
| AB | AB | AB | AB | AA | AB | AB | AB | AB | AB | AA | AB | AB | AB | AB | AA | AB | AB | AB | AB | AB | AB | AB | AA | AB | AA |
| AA | AB | AA | AB | AA | AB | AB | AB | AA | AB | AA | AB | AB | AA | AB | AB | AA | AB | AB | AA | AB | AA | AA | AB | AB | AB |

| snp_B5086 | snp_B5095 | snp_B5103 | snp_B5115 | snp_B5146 | snp_B5156 | snp_B5157 | snp_B5181 | snp_B5184 | snp_B5193 | snp_B5202 | snp_B5239 | snp_B5240 | snp_B5265 | snp_B5279 | snp_B5288 | snp_B5328 |
|---|---|---|---|---|---|---|---|---|---|---|---|---|---|---|---|---|
| 4 | 4 | 4 | 4 | 4 | 4 | 4 | 4 | 4 | 4 | 4 | 4 | 4 | 4 | 4 | 4 | 4 |
| -- | -- | -- | -- | -- | -- | -- | -- | -- | LG6 | -- | -- | -- | LG6 | -- | -- | -- |
| AB | AB | AB | AA | AB | AB | AA | AA | AA | AB | AA | AB | AA | AA | AB | AA | AB |
| AB | AB | AB | AA | AB | AA | AA | AA | AA | AB | AB | AB | AB | AA | AA | AB | AA |
| AB | AB | AB | AA | AB | AB | AA | AB | AA | AB | AA | AB | AA | AA | AB | AA | AB |
| AB | AB | AB | AA | AA | AB | AA | AA | AA | AB | AB | AB | - | AA | AA | AB | AA |
| AA | AA | AA | AA | AA | AA | AA | AA | AB | AB | AB | AA | AA | AA | AA | AB | AA |
| AA | AA | AA | AB | AA | - | - | AB | AB | - | - | AA | - | AA | - | AB | AA |
| AB | AA | AA | AB | AA | AB | AA | AB | AA | AA | AB | AA | AB | AB | AA | AB | AB |
| AB | AB | AA | AA | AB | AB | AA | AA | AA | AB | AA | AB | AB | AA | AA | AA | AB |
| AB | AB | AB | AA | AB | AB | AA | AA | AA | AA | AA | AB | AA | AA | AB | AB | AA |
| AB | AA | AB | AB | AA | AB | AB | AB | AA | AA | AB | AA | AB | AB | AA | AB | AA |
| AA | AB | AA | AB | AA | AA | AA | AA | AB | AA | AB | AA | AA | AB | AA | AA | AA |
| AB | AB | AB | AB | AB | AB | AA | AB | AA | AB | AA | AB | AA | AB | AA | AB | AA |
| AA | AB | AB | AB | AB | AA | AB | AB | AB | AB | AB | AB | AA | AB | AB | AB | AB |
| AA | AA | AA | AA | AA | AA | AA | AA | AB | AB | AB | AA | AB | AA | AA | AB | AB |
| AA | AB | AA | AB | AA | AA | AB | AA | AB | AA | AB | AA | AB | AB | AA | AA | AA |
| AB | AA | AB | AB | AA | AB | AB | AA | AB | AA | AB | AB | AB | AB | AA | AB | AA |
| AB | AA | AB | AA | AA | AB | AA | AA | AA | AA | AA | AA | AB | AB | AA | AA | AB |
| AB | AB | AB | AA | AB | AA | - | AB | AB | AB | AA | AB | - | AA | AA | AB | AB |
| AB | AB | AA | AA | AB | AA | AB | AA | AA | AB | AA | AB | AB | AB | AA | AA | AB |
| AB | AB | AB | AA | AB | AB | AB | AA | AA | AB | AA | AB | - | AA | AA | AA | AA |
| AB | AB | AB | AB | AB | AB | AB | AB | AB | AA | AB | AB | AB | AB | AB | AB | AB |
| AB | AA | AA | AA | AA | AA | AB | AA | AA | AA | AA | AA | AB | AB | AB | AA | AA |
| AB | AA | AB | AA | AB | AB | - | AA | AA | AA | AA | AB | - | AA | AB | AA | AB |
| AB | AB | AB | AA | AB | AB | AA | AA | AA | AB | AA | AB | AB | AA | AA | AA | AB |
| AB | AB | AB | AA | AB | AB | AB | AA | AA | AB | AA | AB | AB | AB | AB | AA | AB |
| AA | AA | AB | AB | AA | AA | AA | AA | AB | AB | AB | AA | AA | AA | AA | AA | AA |
| AB | AB | AB | AA | AB | AB | AB | AA | AA | AB | AA | AB | AB | AB | AA | AA | AB |
| AA | AA | AA | AB | AA | AB | AB | AB | AB | AA | AB | AA | AB | AB | AB | AB | AA |
| AB | AB | AB | AB | AB | AB | AA | AB | AB | AA | AA | AA | AB | - | AB | AA | AA |
| AB | AB | AA | AA | AB | AA | - | AA | AA | - | AA | AB | - | AB | - | AA | AB |
| AA | AB | AA | AB | AB | AA | AB | AB | AB | AA | AB | AB | AA | AA | AB | AB | AA |
| AB | AB | AB | AB | AB | AB | AB | AB | AB | AB | AB | AB | AB | AB | AB | AB | AB |
| AA | AA | AA | AB | AA | AA | AA | AB | AB | - | AB | AA | - | AA | AA | AB | AA |
| AB | AA | AA | AB | AA | AA | AB | AA | AA | AA | AB | AA | - | AB | AA | AB | AA |
| AA | AA | AA | AB | AA | AA | AA | AA | AB | AB | - | AA | AA | AA | AA | AA | AA |
| AB | AA | AB | AB | AB | AB | AB | AA | AB | AA | AB | AB | AB | AB | AB | AB | AB |
| AB | AA | AB | AA | AA | AB | AB | AA | AB | AA | AB | AA | AB | AA | AB | AA | AA |
| AB | AA | AB | AB | AA | AB | AA | AB | AB | AA | AB | AA | AB | AB | AB | AA | AA |
| AA | AA | AB | AB | AA | AB | AB | AB | AB | AA | AB | AA | AB | AA | AA | AB | AA |
| AB | AA | AA | AA | AA | AB | - | AA | AB | AA | AB | AA | AB | AB | AA | AB | AB |
| AB | AB | AB | AA | AB | AA | AA | AB | AA | AB | AA | AA | AA | AA | AA | AB | AB |
| AB | AA | AB | AB | AB | AA | AB | AB | AA | AB | AB | AA | AB | AA | AB | AB | AA |
| AB | AB | AA | AB | AB | AB | AB | AB | AA | AB | AB | AB | AB | AB | AA | AA | AB |
| AB | AB | AB | AB | AB | AB | AB | AB | AB | - | AA | AA | AA | AB | AB | AB | AB |
| AA | AB | AA | AB | AA | AA | AB | AB | AB | AB | AA | AB | AB | AB | AB | AB | AA |

| snp_B5343 | snp_B5358 | snp_B5363 | snp_B5368 | snp_B5435 | snp_B5458 | snp_B5471 | snp_B5484 | snp_B5523 | snp_B5524 | snp_B5558 | snp_B5562 | snp_B5564 | snp_B5596 | snp_B5602 | snp_B5620 | snp_B5634 | snp_B5639 | snp_B5649 | snp_B5652 | snp_B5713 | snp_B5727 | snp_B5735 | snp_B5762 |
|---|---|---|---|---|---|---|---|---|---|---|---|---|---|---|---|---|---|---|---|---|---|---|---|
| 4 | 4 | 4 | 4 | 4 | 4 | 4 | 4 | 4 | 4 | 4 | 4 | 4 | 4 | 4 | 4 | 4 | 4 | 4 | 4 | 4 | 4 | 4 | 4 |
| -- | -- | -- | -- | -- | -- | -- | -- | -- | -- | -- | -- | -- | LG6 | -- | -- | -- | -- | -- | LG6 | LG6 | LG6 | -- | -- |

| snp_B5763 | snp_B5769 | snp_B5779 | snp_B5816 | snp_B5820 | snp_B5828 | snp_B5831 | snp_B5835 | snp_B5907 | snp_B5914 | snp_B5930 | snp_B5935 | snp_B5952 | snp_B5954 | snp_B5966 | snp_B5992 | snp_B6053 | snp_B6162 | snp_B6172 |
|---|---|---|---|---|---|---|---|---|---|---|---|---|---|---|---|---|---|---|
| 4 | 4 | 4 | 4 | 4 | 4 | 4 | 4 | 4 | 4 | 4 | 4 | 4 | 4 | 4 | 4 | 4 | 4 | 4 |
| -- | -- | -- | -- | -- | -- | -- | -- | -- | -- | -- | -- | -- | -- | -- | -- | -- | -- | -- |

| snp_B0001 | snp_B0019 | snp_B0070 | snp_B0101 | snp_B0105 | snp_B0136 | snp_B0156 | snp_B0218 | snp_B0219 | snp_B0226 | snp_B0237 | snp_B0241 | snp_B0243 | snp_B0261 | snp_B0262 | snp_B0289 | snp_B0322 |
|---|---|---|---|---|---|---|---|---|---|---|---|---|---|---|---|---|
| 5 | 5 | 5 | 5 | 5 | 5 | 5 | 5 | 5 | 5 | 5 | 5 | 5 | 5 | 5 | 5 | 5 |
| -- | -- | -- | -- | -- | -- | -- | -- | -- | -- | -- | -- | -- | -- | -- | -- | -- |

| snp_B0343 | snp_B0348 | snp_B0357 | snp_B0370 | snp_B0394 | snp_B0428 | snp_B0430 | snp_B0437 | snp_B0441 | snp_B0444 | snp_B0458 | snp_B0466 | snp_B0467 | snp_B0486 | snp_B0492 | snp_B0506 | snp_B0512 | snp_B0514 | snp_B0519 | snp_B0583 | snp_B0604 | snp_B0616 | snp_B0628 | snp_B0633 | snp_B0643 | snp_B0675 | snp_B0724 | snp_B0729 |
|---|---|---|---|---|---|---|---|---|---|---|---|---|---|---|---|---|---|---|---|---|---|---|---|---|---|---|---|
| 5 | 5 | 5 | 5 | 5 | 5 | 5 | 5 | 5 | 5 | 5 | 5 | 5 | 5 | 5 | 5 | 5 | 5 | 5 | 5 | 5 | 5 | 5 | 5 | 5 | 5 | 5 | 5 |
| -- | -- | -- | -- | -- | -- | -- | -- | LG13 | -- | -- | -- | -- | -- | -- | -- | -- | -- | -- | -- | -- | -- | -- | -- | -- | -- | -- | -- |
| AA | AA | AB | AA | AB | AB | AA | AB | AB | AA | AB | AA | AA | AB | AB | AB | AB | AA | AA | AB | AB | AB | AB | AB | AA | AB | AA | AB |
| AA | AA | AB | AA | AA | AB | AA | AA | AA | AA | AB | AA | AA | AB | AA | AA | AA | AB | AA | AB | AA | AB | AB | AB | AA | AA | AB | AB |
| AA | AA | AA | AB | AA | AB | AB | AB | AA | AB | AA | AB | AA | AA | AB | AB | AA | AB | AB | AB | AB | AA | AB | AB | AA | AB | AB | AB |
| AB | AA | AA | AA | - | AB | - | AA | AA | AA | AA | AB | AA | - | AA | AA | AA | - | AA | AA | AB | AA | AB | AB | AA | AA | AB | AB |
| AB | AB | AB | AB | AB | AA | AB | AA | AA | AB | AA | AB | AA | AB | AB | AA | AB | AB | AA | AB | AB | AA | AA | AB | AA | AB | AB | AB |
| AB | AB | AA | AB | - | AB | - | AA | AB | AA | - | AB | AB | AB | - | AA | AB | - | AB | AA | - | AA | - | - | AA | AB | AB | AA |
| AB | AB | AB | AB | AA | AB | AB | AA | AB | AB | AA | AB | AB | AA | AA | AA | AB | AA | AB | AA | AB | AA | AA | AA | AB | AB | AB | AB |
| AA | AA | AA | AA | AA | AA | AA | AA | AA | AA | AA | AA | AA | AB | AA | AA | AA | AB | AA | AA | AA | AA | AA | AA | AA | AA | AA | AB |
| AB | AB | AA | AB | AB | AA | AB | AB | AA | AB | AA | AA | AA | AB | AB | AB | AB | AB | AB | AA | AB | AA | AA | AB | AB | AB | AB | AB |
| AB | AA | AB | AA | AB | AB | AA | AB | AB | AA | AB | AA | AA | AA | AB | AB | AA | AB | AB | AA | AB | AB | AB | AB | AA | AB | AB | AA |
| AB | AA | AA | AA | AB | AA | AB | AB | AB | AA | AB | AB | AB | AA | AA | AB | AA | AA | AB | AA | AB | AA | AB | AB | AA | AB | AA | AB |
| AB | AA | AB | AA | AB | AB | AB | AB | AA | AA | AB | AB | AA | AA | AB | AA | AA | AA | AA | AA | AB | AA | AB | AB | AB | AA | AB | AB |
| AA | AB | AB | AA | AA | AA | AB | AB | AA | AB | AB | AA | AA | AB | AA | AB | AA | AB | AA | AB | AA | AB | AB | AB | AA | AA | AA | AB |
| AB | AB | AA | AA | AB | AA | AB | AB | AA | AA | AB | AA | AB | AA | AB | AB | AA | AB | AB | AB | AB | AB | AB | AB | AA | AA | AB | AB |
| AB | AA | AB | AA | AB | AB | AA | AB | AB | AB | AA | AB | AB | AB | AB | AB | AB | AB | AB | AA | AB | AB | AB | AA | AB | AB | AA | AA |
| AB | AB | AA | AA | AB | AA | AB | AA | AA | AB | AB | AB | AB | AA | AB | AA | AA | AB | AA | AA | AB | AA | AA | AB | AA | AB | AB | AB |
| AA | AA | AA | AA | AA | AA | AA | AA | AB | AA | AA | AA | AA | AA | AA | AA | AA | AA | AA | AA | AA | AA | AA | AA | AA | AA | AA | AA |
| AA | AB | AB | AA | - | AB | AB | AB | AB | AA | - | AB | - | AB | AB | AA | AB | - | AB | - | AA | AA | AA | AA | AB | AA | - | AA |
| AB | AB | AB | AB | AA | AB | AB | AA | AB | AB | AB | AB | AB | AB | AA | AB | AB | AA | AB | AB | AB | AB | AB | AB | AB | AB | AB | AB |
| AB | AB | AB | AA | - | AB | AB | AB | AB | AB | AB | AA | - | AB | AB | AB | AB | - | AB | AB | AB | AB | AB | - | AA | AB | AA | AB |
| AB | AB | AB | AB | AB | AB | AB | AB | AB | AB | AB | AB | AB | AB | AA | AB | AB | AA | AB | AB | AB | AB | AB | AB | AB | AB | AB | AB |
| AB | AA | AB | AB | AA | AB | AB | AB | AB | AB | AB | AB | AB | AA | AA | AB | AB | AA | AB | AB | AB | AB | AB | AB | AB | AB | AB | AB |
| AB | AA | AA | AA | - | AA | - | AA | - | AA | - | AA | AB | - | - | - | AB | AB | - | AB | AB | AA | AA | - | AA | AB | - | AA |
| AA | AA | AB | AA | AA | AB | AB | AB | AB | AB | AB | AA | AA | AA | AA | AB | AB | AA | AB | AB | AA | AB | AB | AA | AA | AB | AA | AB |
| AA | AB | AA | AA | AB | AA | AA | AB | AA | AB | AB | AA | AA | AB | AB | AB | AA | AB | AA | AB | AA | AA | AB | AB | AA | AA | AA | AB |
| AB | AB | AB | AB | AB | AB | AB | AB | AB | AB | AB | AB | AB | AB | AB | AB | AB | AB | AB | AB | AB | AB | AB | AB | AB | AB | AB | AB |
| AA | AA | AA | AA | AA | AA | AA | AA | AA | AA | AA | AB | AA | AA | AA | AA | AA | AA | AA | AA | AA | AA | AA | AA | AA | AA | AB | AB |
| AA | AA | AB | AB | AA | AB | AA | AA | AB | AA | AB | AA | AA | AA | AA | AA | AB | AA | AA | AA | AB | AB | AB | AA | AB | AB | AA | AA |
| AA | AA | AA | AA | AA | AA | AB | AB | AA | AA | AA | AB | AB | AB | AA | AB | AA | AA | AA | AB | AB | AA | AA | AA | AB | AA | AB | AA |
| AA | AB | AA | - | - | AA | AA | AB | - | AB | - | AA | - | AB | - | - | AA | - | - | AB | - | AB | AB | - | AA | AA | - | AB |
| AB | AB | AB | AB | AB | AB | AB | AB | AB | AB | AB | AB | AB | AB | AB | AB | AB | AB | AB | AB | AB | AB | AB | AB | AB | AB | AB | AB |
| AA | AB | AA | AB | AA | AA | AA | AA | AA | AB | AA | AB | AA | AA | AA | AA | AA | AA | AA | AA | AA | AA | AA | AB | AA | AA | AB | AA |
| AB | AB | AA | - | - | AA | AB | AA | AA | AA | AA | AB | - | AB | AB | AA | AA | - | AA | AA | AA | AA | AA | AA | AB | AB | AB | AA |
| AA | AB | AB | AB | AA | AA | AA | AB | AB | AB | AB | AA | - | AB | AA | AA | AB | AA | AB | AA | AA | AA | AB | - | AB | AB | AB | AB |
| AB | AB | AA | AA | - | AA | AB | AA | AA | AB | AB | AB | AB | AA | AA | AA | - | AA | AA | AA | AB | AA | AA | - | AB | AB | AB | AB |
| AB | AB | AB | AB | AB | AB | AB | AB | AB | AB | AB | AA | - | AB | AB | AB | AB | - | AB | AB | AB | AB | AB | AB | AB | AB | AB | AB |
| AA | AA | AB | AB | AA | AA | AB | AA | AB | AB | AA | AB | AB | AA | AA | AA | AA | AA | AA | AA | AA | AA | AA | AA | AB | AA | AB | AA |
| AB | AB | AB | AB | AB | AB | AB | AB | AB | AB | AB | AB | AB | AB | AB | AB | AB | AB | AB | AB | AB | AB | AB | AB | AB | AB | AB | AB |
| AB | AB | AB | AB | AA | AB | AB | AB | AB | AB | AB | AB | AB | AB | AB | AB | AB | AA | AB | AB | AB | AB | AB | AB | AB | AB | AB | AB |
| AB | AB | AB | AB | AB | AA | AB | AB | AA | AB | - | AB | - | AB | AB | AB | AB | - | AA | AB | AB | AB | AB | - | AA | AB | AB | AB |
| AB | AB | AB | AB | AB | AB | AB | AA | AB | AB | AB | AB | AB | AA | AB | AA | AB | AB | AB | AB | AB | AB | AA | AB | AB | AB | AB | AB |
| AA | AA | AA | AA | AB | AA | AA | AB | AB | AA | AB | AA | AA | AB | AB | AB | AA | AB | AA | AB | AA | AA | AB | AB | AA | AA | AA | AB |
| AB | AA | AB | AA | AA | AB | AB | AA | AB | AB | AA | AA | AB | AB | AA | AB | AA | AA | AB | AA | AB | AA | AA | AA | AB | AB | AA | AB |
| AA | AB | AB | AA | AA | AA | AB | AA | AB | AB | AA | AA | AA | AB | AA | AB | AA | - | AA | AA | AA | AB | AB | - | AB | AB | AB | AB |
| AB | AA | AA | AB | AB | AA | AB | AB | AB | AA | AB | AA | AA | AA | AB | AB | AA | AB | AB | AA | AB | AB | AB | AB | AA | AB | AA | AB |
| AB | AB | AB | AB | AB | AB | AB | AB | AB | AB | AB | AB | AB | AB | AB | AB | AB | AB | AB | AB | AB | AB | AB | AB | AB | AB | AB | AB |
| AA | AA | AA | AA | AB | AA | AA | AA | AA | AA | AA | AA | AA | AA | AB | AA | AA | AB | AA | AA | AA | AA | AA | AA | AA | AA | AA | AA |
| AA | AA | AA | AA | AA | AA | AA | AA | AA | AA | AA | AA | AA | AA | AA | AA | AA | AA | AA | AA | AA | AA | AA | AA | AA | AA | AA | AA |
| AA | AA | AA | AB | AB | AA | AA | AB | AA | AA | AA | AB | AA | AB | AB | AB | AA | AB | AA | AB | AA | AA | AB | AB | AA | AA | AB | AB |
| AA | AA | AA | AA | AA | AB | AB | AA | AA | AA | AB | AB | AA | AB | AA | AB | AA | AA | AA | AA | AA | AA | AA | AA | AA | AA | AA | AA |
| AB | AA | AA | AA | AA | AB | AB | AA | AB | AA | AB | AB | AA | AB | AB | AB | AB | AB | AA | AB | AB | AA | AB | AB | AA | AA | AA | AA |
| AB | AA | AA | AA | - | AB | AA | AA | AB | AA | - | AA | AA | AB | AB | AA | AB | AB | AA | AB | AA | AA | AB | AB | AB | AA | AB | AB |
| AA | AB | AA | AA | AB | AA | AB | AB | AB | AA | AB | AA | AA | AB | AB | AB | AB | AB | AA | AB | AB | AB | AB | AB | AA | AB | AB | AB |
| AB | AB | AB | AB | AB | AB | AB | AB | AB | AB | AB | AA | AB | AB | AB | AB | AB | AB | AB | AB | AB | AB | AB | AB | AB | AB | AA | AB |
| AB | AA | AB | AA | AB | AB | AB | AB | AB | AA | AB | AB | AA | AB | AB | AB | AB | AB | AB | AB | AB | AB | AB | AA | AA | AB | AA | AB |
| AA | AB | AB | AB | AB | AB | AB | AA | AB | AA | AA | AA | AB | AA | AB | AB | AB | AB | AB | AA | AA | AB | AA | AA | AB | AA | AB | AA |
| AB | AA | AB | AA | AA | AB | AA | AA | AA | AA | AB | AB | AA | AA | AA | AB | AB | AB | AA | AB | AB | AB | AB | AB | AA | AB | AB | AB |
| AA | AA | AB | AB | AA | AB | AA | AB | AB | AB | AB | AB | AB | AB | AA | AA | AB | AA | AB | AB | AA | AB | AB | AA | AB | AA | AB | AB |
| AA | AA | AA | AB | AB | AA | AA | AA | AB | AA | AA | AA | AB | AB | AA | AB | AB | AB | AB | AA | AA | AB | AA | AB | AA | AA | AB | AA |
| AA | AA | AB | AB | AB | AA | AA | AA | AB | AA | AB | AA | AB | AB | AB | AB | AB | - | AB | AB | AA | AA | AB | AB | AA | AB | AB | AB |
| AB | AA | AB | AB | AB | AB | AB | AB | AB | AB | AB | AA | - | AA | AB | - | AB | AB | AB | AB | AA | AB | AB | AA | AB | AB | AA | AA |
| AB | AB | AB | AB | AB | AB | AB | AB | AB | AB | AB | AB | AB | AB | AB | AB | AB | AB | AB | AB | AB | AB | AB | AB | AA | AB | AA | AB |
| AB | AB | AB | AB | AB | AB | AB | AB | AB | AB | AB | AA | AB | AB | AB | AB | AB | AB | AB | AB | AB | AB | AB | AB | AB | AB | AA | AB |
| AA | AA | AB | AB | - | AB | AA | AA | AB | AA | - | AA | - | AB | AB | AA | AB | - | AA | AB | - | AB | AB | AA | AB | AB | AA | AA |
| AA | AB | AB | AA | AB | AA | AA | AB | AA | AB | AB | AB | AA | AA | AB | AB | AA | AA | AA | AA | AA | AB | AA | AA | AB | AA | AB | AA |
| AA | AB | AB | AA | AA | AB | AA | AB | AB | AA | AA | AB | AB | AA | AA | AB | AB | AA | AA | AB | AA | AB | AA | AA | AA | AB | AA | AA |
| AB | AB | AB | AA | AB | AB | AB | AB | AB | AB | AB | AB | AB | AA | AB | AB | AA | AB | AB | AB | AB | AB | AB | AA | AA | AB | AB | AA |
| AB | AB | AB | AB | AB | AB | AB | AB | AB | AB | AB | AB | AB | AB | AB | AB | AB | - | AB | AB | AB | AB | AB | - | AB | AB | AB | AB |
| AB | AA | AA | AB | AA | AB | AB | AB | AB | AA | AA | AA | AB | AA | AA | AA | AB | AA | AB | AA | AB | AA | AA | AB | AA | AB | AB | AA |
| AB | AA | AB | AA | AA | AB | AB | AA | AB | AA | AB | AA | AB | AA | AA | AA | AA | AA | AB | AA | AB | AA | AA | AB | AA | AB | AB | AA |
| AB | AB | AB | AB | AB | AB | AB | AB | AB | AB | AB | AB | AB | AB | AB | AB | AB | AB | AB | AB | AB | AB | AA | AB | AB | AB | AB | AB |
| AA | AB | AA | AB | AA | AA | AB | AA | AB | AA | AA | AB | AB | AA | AA | AA | AB | AA | AA | AB | AA | AB | AA | AA | AB | AA | AB | AB |
| AB | AB | AB | AA | AB | AB | AA | AB | AB | AB | AB | AA | AA | AA | AB | AA | AB | AB | AA | AB | AA | AB | AB | AB | AA | AB | AB | AB |
| AB | AB | AA | AA | AB | AA | AB | AA | AB | AB | AB | AB | AA | AA | AA | AA | AB | AA | AA | AA | AB | AA | AB | AB | AB | AB | AB | AB |
| AB | AA | AA | AB | AA | AA | AB | AB | AA | AA | AA | AB | AA | AB | AA | AB | AA | AB | AA | AB | AA | AA | AA | AA | AB | AB | AA | AA |
| AB | AB | AB | AA | AB | AB | AB | AB | AB | AA | AB | AA | AA | AB | AA | AB | AA | AB | AB | AB | AB | AB | AB | AA | AB | AB | AB | AB |
| AA | AB | AB | AB | AB | AB | AA | AA | AB | AB | AA | AA | AB | AB | AA | AA | AB | AA | AB | AB | AB | AB | AB | AB | AA | AA | AA | AB |
| AB | AA | AA | AA | AA | AA | AA | AB | AB | AA | AB | AA | - | AA | - | AB | AA | - | AA | AA | AB | AB | AB | - | AB | AA | AB | AB |
| AB | AB | AB | AB | AB | AB | AB | AB | AA | AB | AA | AA | AB | AB | AB | AB | AB | AB | AA | AB | AB | AB | AB | AB | AB | AB | AB | AB |
| AB | AB | AA | - | AB | AA | - | AA | AA | AA | - | AB | AA | - | - | AA | AA | - | AA | AA | AB | AA | - | - | AB | AA | AB | AA |
| AB | AB | AA | AA | AB | AA | AB | AA | AA | AA | AA | AB | AA | AB | AB | AA | AA | AB | AA | AA | AB | AA | AA | AB | AA | AA | AB | AA |
| AA | AA | AB | AB | AA | AB | AA | AB | AA | AB | AA | AB | AA | AB | AB | AA | AB | AA | AB | AA | AB | AA | AA | AB | AA | AA | AB | AA |
| AB | AB | AB | AB | AA | AB | AB | AA | AB | AB | AB | AB | AA | AB | AB | AB | AB | AB | AA | AA | AB | AA | AB | AB | AA | AA | AB | AB |
| AA | AB | AB | AB | AA | AB | AA | AB | AB | AB | AB | AA | AB | AB | AB | AB | AB | AA | AB | AB | AA | AB | AB | AA | AA | AB | AB | AB |
| AA | AB | AA | - | - | AA | - | AA | - | AA | AA | AA | - | - | AB | - | AA | - | AA | AA | AB | AA | AA | - | AB | AA | AA | AA |
| AA | AB | AA | AA | - | AA | - | AB | AA | AB | - | AB | AA | - | AB | - | - | - | AA | AB | AA | AA | AA | - | AA | AA | AB | AA |
| AB | AB | AA | AB | AB | AA | AB | AA | AA | AB | AB | AB | AB | AA | AB | AA | AA | AB | AA | AA | AB | AA | AA | AB | AA | AB | AB | AA |
| AA | AB | AA | AB | AA | AA | AA | AA | AB | AA | AA | AA | AA | AB | AA | AB | AA | AB | AA | AA | AA | AA | AA | AB | AA | AB | AA | AB |
| AB | AB | AB | AB | - | AB | - | AB | AB | AB | - | AB | - | AB | AA | AA | AB | AA | AB | AB | AB | AB | AB | AB | AB | AA | AB | AB |
| AA | AB | AA | AA | - | AA | AA | AA | AA | AB | AA | AA | AA | AB | AA | AB | AA | AB | AA | AB | AA | AB | AA | - | AA | AA | AA | AA |
| AA | AA | AA | AB | AA | AA | AA | AA | AA | AB | AB | AB | - | AB | AA | AB | AA | AA | AA | AB | AA | AB | AB | AB | AA | AA | AB | AA |
| AA | AA | AA | AA | AA | AA | AA | AA | AB | AA | AA | AB | AA | AA | AA | AA | AA | AA | AA | AB | AA | AA | AB | AA | AA | AA | AA | AA |
| AA | AB | AA | AA | AB | AA | AA | AB | AB | AB | AB | AA | AA | AB | AB | AB | AA | AB | AA | AB | AA | AA | AB | AA | AA | AA | AB | AB |
| AA | AA | AB | AA | - | AA | - | AA | - | AA | - | AA | AA | AB | AA | AA | AA | - | AA | AA | AA | AA | AA | - | AB | AA | AA | AA |
| AB | AA | AA | AB | AA | AA | AB | AA | AA | AA | AA | AA | AB | AA | AA | AA | AA | AA | AA | AA | AB | AA | AA | AA | AA | AA | AA | AA |
| AB | AA | AB | - | - | AB | AB | AA | AB | AA | AA | AA | - | - | - | - | AB | - | AB | AA | AB | AB | AA | - | AB | AB | AB | AA |
| AB | AA | AA | AB | AA | AB | AB | AB | AB | AA | AA | AB | AB | AA | AA | AB | AA | AA | AB | AA | AB | AB | AA | AB | AA | AB | AB | AB |
| AB | AB | AA | AA | AB | AA | AB | AA | AA | AA | AA | AA | AA | AB | AB | AA | AA | AB | AA | AA | AA | AA | AA | AA | AB | AA | AA | AA |
| AB | AB | AB | AB | - | AB | - | AA | AB | AB | AB | AB | - | AA | AB | AA | AB | - | AB | AA | AB | AA | AA | - | AB | AB | AA | AA |
| AB | AB | AB | AB | AB | AB | AB | AB | AB | AB | AB | AB | AB | AB | AB | AB | AB | AB | AB | AB | AB | AB | AB | AB | AB | AB | AB | AB |
| AA | AA | AA | AA | - | AA | AA | AA | AA | AA | - | AA | AA | AB | AA | AA | AA | AA | AA | AA | AA | AA | AA | AA | AA | AA | AA | AA |
| AB | AA | AA | AB | - | AB | AA | AA | AA | AA | - | AB | - | AA | AA | AA | AA | - | AB | AA | AB | AA | AB | - | AA | AA | AB | AA |
| AA | AB | AA | AA | AB | AA | AA | AA | AA | AA | AA | AB | AA | AB | AB | AA | AA | AB | AA | AB | AA | AA | AA | AB | AB | AA | AB | AA |
| AA | AB | AA | AA | AA | AA | AA | AB | AA | AB | AA | AB | AA | AB | AB | AA | AA | AB | AA | AA | AB | AA | AA | AB | AB | AA | AB | AA |
| AB | AA | AA | AA | AA | AA | AB | AA | AA | AA | AA | AA | AA | AA | AB | AA | AA | AA | AA | AA | AA | AA | AB | AA | AB | AA | AB | AA |
| AA | AB | AA | AA | AB | AA | AB | AA | AA | AA | AA | AA | AA | AB | AB | AA | AA | AB | AA | AA | AB | AA | AA | AB | AA | AA | AA | AA |
| AB | AA | AB | AA | AB | AB | AB | AB | AB | AB | AB | AA | AB | AB | AB | AB | AB | AB | AB | AB | AB | AB | AB | AA | AB | AA | AB | AB |
| AB | AB | AB | AB | AB | AB | AB | AB | AB | AB | AB | AB | AB | AB | AB | AB | AB | AB | AB | AB | AB | AB | AB | AB | AB | AB | AB | AB |
| AA | AB | AA | AB | AA | AA | AB | AA | AA | AA | AA | AB | AA | AB | AA | AA | AA | AA | AB | AA | AA | AB | AA | AB | AA | AB | AA | AA |
| AB | AB | AA | AB | AB | AA | AB | AA | AA | AA | AA | AA | AA | AB | AB | AA | AA | AB | AA | AB | AA | AA | AB | AA | AB | AB | AA | AA |
| AB | AA | AB | AB | AB | AB | AB | AA | AB | AB | AB | AB | AB | AB | AB | AB | AB | AA | AB | AB | AB | AB | AB | AB | AA | AB | AB | AB |
| AB | AB | AB | - | - | AB | AB | AB | AA | AB | AB | AB | - | - | AB | AB | AB | AB | - | AB | AB | AB | AB | AB | - | AB | AB | AB |
| AA | AB | AA | AB | AB | AA | AA | AB | AA | AB | AB | AB | AA | AB | AB | AB | AB | AA | AB | AA | AA | AA | AB | AB | AB | AA | AB | AB |
| AB | AB | AA | AA | AB | AA | AB | AA | AB | AA | AB | AA | AB | AA | AB | AA | AB | AA | AB | AA | AB | AA | AA | AB | AA | AB | AA | AA |
| AA | AB | AA | AA | AB | AA | AB | AA | AB | AB | AB | AB | AA | AB | AB | AA | AB | AA | AA | AB | AB | AB | AB | AA | AB | AA | AB | AB |
| AA | AA | AB | AA | AB | AA | AB | AA | AB | AA | AB | AA | AB | AA | AB | AA | AB | AA | AB | AA | AB | AA | AA | AB | AA | AA | AA | AA |
| AA | AB | AA | AA | AB | AA | AA | AB | AB | AB | AB | AA | AA | AB | AB | AB | AA | AA | AB | AA | AB | AB | AB | AA | AB | AA | AA | AB |
| AA | AA | AA | AA | AA | AA | AA | AA | AA | AA | AB | AA | AA | AA | AB | AA | AA | AA | AA | AA | AA | AB | AA | AA | AA | AA | AA | AB |
| AA | AB | AB | AB | AB | AA | AA | AB | AA | AB | AB | AA | AA | AB | AB | AB | AA | AB | AA | AB | AB | AB | AB | AA | AB | AA | AA | AB |
| AB | AA | AB | AA | AB | AA | AB | AA | AB | AA | AB | AA | AA | AB | AB | AA | AB | AA | AB | AA | AB | AA | AA | AA | AA | AB | AA | AA |
| AB | AB | AB | AB | AB | AB | AB | AB | AB | AB | AB | AB | AB | AB | AB | AB | AB | AB | AB | AB | AB | AB | AB | AB | AB | AB | AB | AB |
| AA | AB | AA | AB | AB | AA | AA | AB | AA | AB | AB | AB | AA | AB | AB | AB | AB | AB | AA | AB | AB | AB | AA | AB | AB | AB | AA | AB |
| AA | AA | AA | AA | AB | AA | AA | AB | AB | AB | AA | AA | AA | AA | AB | AB | AA | AA | AA | AB | AB | AB | AB | AA | AA | AA | AA | AA |
| AA | AA | AA | AA | AA | AA | AA | AA | AA | AA | AA | AA | AA | AA | AA | AA | AA | AA | AA | AA | AA | AA | AA | AA | AA | AA | AA | AA |
| AA | AA | AA | AA | AB | AA | AA | AB | AA | AB | AB | AA | AB | AA | AB | AB | AB | AB | AB | AA | AA | AB | AB | AA | AB | AA | AA | AA |
| AA | AA | AB | AB | AA | AB | AA | AB | AB | AB | AB | AA | AB | AA | AA | AB | AB | AA | AB | AB | AA | AB | AB | AA | AA | AB | AA | AB |
| AB | AA | AB | AB | AB | AB | AB | AA | AB | AB | AB | AA | AB | AB | AA | AB | AB | AB | AB | AB | AB | AB | AB | AB | AB | AB | AB | AB |
| AB | AA | AB | AB | AA | AB | AB | AA | AB | AA | AA | AB | AB | AA | AA | AA | AB | AB | AA | AB | AA | AB | AA | AB | AB | AB | AB | AB |
| AB | AB | AB | AB | AB | AB | AB | AB | AB | AB | - | AB | AB | AB | AB | AB | AB | AB | AB | AB | AB | AB | AB | - | AB | AB | AB | AB |
| AA | AB | AA | AB | AB | AA | AB | AA | AB | AB | AB | AA | AB | AB | AB | AB | AB | AB | AB | AA | AB | AA | AB | AB | AA | AB | AA | AB |
| AA | AA | AB | AB | AA | AB | AA | AB | AB | AB | - | AA | AB | AA | AB | AB | AB | AB | AA | AB | AA | AB | AB | AA | AA | AB | AA | AB |
| AA | AB | AA | AA | AB | AA | AA | AA | AB | AB | AB | AA | AA | AA | AB | AA | AA | AA | AB | AA | AB | AA | AA | AB | AA | AB | AA | AA |
| AA | AA | AA | AA | AB | AA | AA | AA | AA | AA | AA | AA | AA | AA | AA | - | AA | AA | - | AA | AA | AA | AA | AA | - | AA | AA | AA |
| AA | AB | AB | AA | - | AB | AA | AA | AA | AA | AA | AB | AB | AB | AB | AA | AB | AA | AB | AA | AA | AB | AA | AB | AA | AB | AA | AA |
| AB | AB | AB | AB | AA | AB | AB | AA | AB | AA | AB | AB | AB | AB | AA | AA | AB | AA | AB | AB | AB | AB | AA | AB | AA | AA | AB | AB |
| AA | AA | AB | AA | AA | AB | AB | AA | AB | AA | AA | AA | AB | AA | AB | AB | AA | AA | AB | AA | AA | AB | AA | AB | AA | AA | AA | AB |
| AB | AB | AB | AB | AB | AB | AB | AA | AB | AA | AB | AB | AA | AB | AB | AA | AB | AA | AB | AB | AB | AB | AB | AB | AB | AA | AB | AB |
| AB | AA | AA | AB | AA | AB | AB | AA | AA | AA | AA | AA | AA | AB | AA | AB | AA | AB | AA | AB | AB | AB | AA | AB | AA | AB | AB | AB |
| AB | AB | AB | AB | AA | AB | AB | AA | AB | AB | - | AB | AB | - | AA | AB | AB | AA | AB | AB | AB | AB | AA | AB | AB | AB | AB | AB |
| AB | AB | AA | - | AA | AB | AA | AB | AA | AA | AB | AB | AB | - | AA | AA | AA | AA | - | AA | AA | - | AA | AA | - | AB | AB | AA |
| AA | AB | AA | AA | AB | AB | AA | AB | AA | AB | - | AA | AB | AA | AB | AA | AA | AB | AA | AB | AA | AA | AB | AA | AB | AA | AB | AA |
| AB | AB | AB | AB | - | AB | AB | AA | AB | AB | AB | AB | - | AB | AB | AB | AB | AB | AB | AB | AB | AB | AB | - | AB | AB | AB | AB |
| AA | AB | AA | AB | AB | AA | AA | AB | AA | AB | AA | AA | AA | - | - | AB | AA | - | AA | AB | AA | AB | AB | AB | AB | AA | AB | AA |
| AA | AB | AB | AB | AB | AB | AA | AB | AB | AB | AB | AA | AA | AA | AB | AB | AB | AB | AA | AB | AA | AB | AB | AA | AB | AB | AA | AB |
| AA | AB | AB | AA | AB | AB | AA | AB | AB | AB | AB | AB | AA | AB | AB | AB | AA | AB | AA | AB | AB | AB | AB | AA | AB | AA | AB | AB |
| AA | AA | AA | AA | AA | AA | AA | AA | AA | AA | AA | AA | AA | AA | AA | AA | AA | AA | AA | AA | AA | AA | AA | AA | AA | AA | AA | AA |
| AA | AA | AA | AA | AA | AA | AA | AA | AB | AA | - | AA | AA | - | AA | - | AA | - | AA | AA | - | AA | AA | - | AA | AA | AA | AA |
| AB | AA | AA | AA | AA | AA | AA | AA | AA | AA | AA | AA | AA | AB | AA | AA | AA | AB | AA | AA | AB | AA | AA | AA | AA | AA | AB | AB |
| AA | AB | AA | AA | AB | AA | AB | AA | AA | AA | AA | AA | AB | AB | AA | AA | AB | AB | AB | AA | AB | AA | AA | AA | AA | AB | AA | AA |
| AB | AB | AB | AB | AB | AB | AB | AB | AB | AB | AB | AB | AB | AA | AB | AA | AB | AA | AB | AB | AB | AB | AB | AB | AB | AB | AB | AB |
| AA | AB | AA | AB | AB | AB | AA | AB | AB | AA | AB | AA | AA | AB | AA | AB | AB | AB | AB | AB | AA | AB | AB | AA | AB | AA | AB | AB |
| AA | AA | AA | AA | AA | AA | AA | AA | AB | AA | AA | AA | - | - | - | AA | AA | AA | AA | AA | AA | AA | AA | - | AA | AA | AA | AA |
| AB | AA | AB | AB | AA | AB | - | AA | AA | AA | AA | AB | - | - | - | AA | AB | - | AB | AA | AB | AA | AA | - | AA | AB | AB | AA |
| AA | AB | AB | AB | AB | AB | AA | AB | AB | AA | AB | AA | AB | AA | AB | AB | AA | AB | AB | AB | AA | AB | AA | AA | AB | AA | AB | AB |
| AA | AB | AB | AB | AB | AA | AB | AA | AB | AB | AB | AB | AB | AB | AB | AA | AB | AB | AB | AB | AB | AB | AB | AB | AB | AB | AB | AA |
| AB | AA | AB | AB | AA | AB | AB | AA | AB | AA | AA | AB | AB | AA | AA | AA | AB | AB | - | AB | AA | AB | AB | AA | AA | AB | AB | AB |
| AB | AA | AB | AA | AB | AB | AB | AA | AB | AA | AB | AB | AB | AB | AA | AB | AA | AB | AA | AA | AB | AB | AA | AB | AB | AB | AB | AB |
| AB | AB | AB | AB | AB | AB | AB | AB | AA | AB | AA | AB | AB | AB | AB | AB | AB | AB | AB | AA | AB | AB | AA | AA | AB | AA | AB | AB |
| AB | AB | AB | AB | - | AB | AB | AB | AA | AB | - | AB | AB | AB | AB | AA | AB | AB | AB | AB | AB | AB | AB | - | AB | AB | AB | AB |
| AA | AB | AA | AA | - | AA | - | AA | AA | AA | - | AB | - | AB | - | AA | AA | - | - | AB | AA | - | AA | AA | - | AA | AB | AA |
| AB | AA | AB | AA | AA | AB | AB | AA | AB | AA | AA | AB | AB | AB | AA | AA | AB | AA | AB | AA | AB | AB | AA | AA | AB | AA | AB | AA |
| AB | AB | AA | AB | AB | AA | AB | AA | AB | AA | AA | AB | AB | AB | AA | AB | AB | AB | AB | AA | AB | AA | AB | AB | AB | AB | AB | AB |
| AA | AA | AB | AA | AB | AB | AB | AB | AB | AB | AB | AA | AA | AA | AB | AA | AB | AB | AB | AB | AB | AB | AA | AB | AA | AB | AA | AB |
| AA | AA | AA | AA | AA | AA | AA | AA | AA | AA | AA | AA | AA | AA | AA | AA | AA | AB | AA | AA | AA | AA | AA | AA | AA | AA | AA | AB |
| AA | AB | AA | AA | AA | AB | AA | AB | AA | AB | AB | AB | AA | AA | AA | AB | AA | AA | AB | AA | AB | AA | AA | AB | AB | AB | AA | AA |
| AB | AA | AB | AB | AA | AB | AB | AA | AB | AB | AB | AB | AB | AB | AA | AA | AB | AA | AB | AB | AB | AA | AB | AA | AB | AB | AB | AA |
| AA | AB | AB | AB | AA | AB | AA | AA | AB | AA | AB | AB | AA | AA | AA | AB | AB | AA | AA | AA | AB | AB | AA | AA | AB | AB | AB | AB |
| AA | AB | AA | AB | AA | AB | AA | AA | AB | AA | AB | AB | AA | AA | AA | AB | AA | AA | AA | AB | AA | AB | AA | AB | AA | AB | AB | AA |
| AB | AA | AA | AA | AB | AA | AA | AB | AA | AB | AB | AA | AA | AA | AA | AB | AA | AA | AB | AA | AA | AB | AB | AA | AB | AA | AA | AB |
| AA | AA | AA | AA | AA | AA | AA | AB | AA | AA | AA | AB | AB | AA | AA | AA | AB | AA | AA | AA | AA | AB | AA | AB | AA | AA | AB | AA |
| AA | AA | AA | AB | AB | AA | AA | AB | AB | AB | AB | AA | AB | - | - | AB | AA | AA | AB | AA | AA | AA | AA | AB | AA | AA | AB | AA |
| - | AB | AA | AB | AB | AA | AB | AB | AA | AB | - | AB | - | AA | - | AB | AA | AB | AA | AA | - | AA | AA | - | AB | AB | AB | AB |
| AB | AB | AA | AA | AB | AA | AB | AA | AA | AA | AA | AB | AA | AB | AB | AA | AA | AB | AA | AA | AB | AA | AA | AB | AA | AA | AB | AA |
| AA | AA | AA | AA | AA | AB | AA | AA | AA | AA | AA | AB | AB | AA | AA | AB | AA | AA | AA | AA | AA | AB | AA | AB | AB | AA | AB | AA |
| AB | AB | AB | AB | AB | AB | AB | AA | AB | AB | AB | AB | AB | AB | AA | AA | AB | AA | AB | AB | AB | AB | AB | AB | AB | AA | AB | AB |
| AA | AB | AB | AA | AA | AB | AA | AA | AB | AA | AA | AB | AA | AB | AA | AB | AB | AA | AB | AA | AB | AA | AB | AA | AA | AB | AA | AB |
| AA | AB | AA | AB | AB | AA | AB | AB | AA | AB | AB | AB | AB | AA | AB | AA | AB | AA | AB | AA | AA | AA | AB | AA | AB | AB | AA | AA |
| AA | AB | AB | AA | AB | AB | AB | AB | AB | AB | AB | AB | AA | - | AB | AB | AB | AB | AB | AA | AA | - | AA | AA | - | AB | AB | AA |
| AB | AA | AA | AA | AB | AA | AB | AA | AB | AA | AB | AB | AB | AA | AB | AA | AB | AB | AA | AB | AB | AA | AB | AB | AA | AA | AB | AB |
| - | AA | AB | - | AA | AB | AB | AB | AB | AA | AA | AA | - | - | - | AB | AB | - | AB | AB | AB | AB | AA | AA | AA | AA | AB | AB |
| AA | AB | AA | AB | AA | AB | AA | AA | AA | AB | AA | AB | AA | AA | AA | AA | AA | AA | AA | AA | AA | AA | AA | AB | AB | AA | AB | AA |
| AB | AB | AB | AB | AB | AB | AB | AB | AB | AB | AB | AB | AB | AB | AB | AB | AB | AB | AB | AB | AB | AB | AB | AB | AB | AB | AB | AB |

| snp_B0736 | snp_B0765 | snp_B0789 | snp_B0812 | snp_B0829 | snp_B0839 | snp_B0844 | snp_B0857 | snp_B0860 | snp_B0882 | snp_B0895 | snp_B0934 | snp_B0995 | snp_B1065 | snp_B1091 | snp_B1096 | snp_B1138 | snp_B1157 | snp_B1164 | snp_B1183 | snp_B1184 |
|---|---|---|---|---|---|---|---|---|---|---|---|---|---|---|---|---|---|---|---|---|
| 5 | 5 | 5 | 5 | 5 | 5 | 5 | 5 | 5 | 5 | 5 | 5 | 5 | 5 | 5 | 5 | 5 | 5 | 5 | 5 | 5 |
| -- | -- | -- | -- | -- | -- | -- | -- | -- | -- | -- | -- | -- | -- | -- | -- | -- | -- | -- | -- | -- |
| AA | AA | AB | AA | AB | AB | AB | AB | AB | AA | AB | AB | AA | AA | AA | AB | AA | AA | AB | AB | AB |
| AB | AB | AB | AA | AB | AB | AB | AB | AB | AA | AB | AB | AA | AB | AA | AA | AB | AB | AB | AA | AB |
| AB | AB | AB | AA | AB | AB | AA | AB | AA | AA | AB | AA | AA | AA | AB | AA | AB | AB | AB | AA | AB |
| AB | AB | AA | AA | AA | - | AA | AA | AB | AA | AB | AB | AA | - | - | AA | AB | AB | AB | AA | AB |
| AB | AB | AB | AA | AB | AB | AA | AA | AA | AB | AA | AB | AB | AA | AA | AA | AA | AA | AA | AB | AB |
| AA | AB | - | - | - | - | AA | - | AB | AB | AA | AB | AB | - | - | AA | - | - | - | AA | - |
| AB | AB | AA | AA | AB | AB | AA | AB | AB | AB | AA | AB | AB | AB | AB | AA | AB | AB | AB | AB | AB |
| AB | AA | AB | AA | AA | AA | AB | AA | AB | AB | AA | AA | AA | AA | AA | AA | AA | AB | AA | AA | AA |
| AB | AA | AB | AA | AB | - | AA | AB | AA | AA | AB | AA | AA | AB | AB | AA | AB | AB | AB | AA | AB |
| AB | AB | AB | AB | AA | AA | AB | AB | AB | AA | AA | AA | AB | AB | AB | AA | AB | AA | AB | AA | AA |
| AB | AB | AB | AA | AB | AB | AB | AA | AB | AA | AB | AA | AA | AB | AA | AA | AB | AB | AA | AB | AA |
| AB | AB | AB | AB | AB | AB | AB | AA | AA | AA | AB | AA | AB | AA | AB | AA | AB | AA | AB | AA | AB |
| AB | AA | AB | AB | AB | AB | AB | AB | AB | AA | AB | AA | AB | AA | AA | AB | AA | AB | AA | AB | AA |
| AB | AA | AB | AA | AA | AB | AB | AA | AB | AA | AB | AB | AB | AA | AA | AB | AA | AB | AA | AA | AB |
| AB | AA | AB | AB | AA | AA | AA | AA | AB | AA | AB | AB | AA | AA | AA | AA | AB | AB | AA | AB | AA |
| AB | AB | AB | AB | AA | AB | AA | AB | AB | AA | AB | AB | AB | AA | AB | AB | AB | AB | AA | AA | AA |
| AA | AB | AA | AA | AA | AA | AA | AA | AA | AA | AA | AB | AA | AA | AA | AB | AA | AA | AA | AA | AA |
| - | AB | - | - | - | AA | AA | - | - | - | AA | AA | AB | - | - | AA | AA | - | - | - | AB |
| AB | AB | AB | AB | AB | AB | AB | AB | AB | AB | AB | AB | AB | AB | AB | AB | AB | AB | AB | AB | AB |
| AB | AA | AB | AB | AB | AB | AB | AB | AB | AB | AB | AB | AB | AB | AB | AB | AB | AB | AB | AB | AB |
| AB | AB | AB | AB | AB | AB | AB | AB | AB | AB | AB | AB | AB | AB | AB | AB | AB | AB | AB | AB | AB |
| AB | AB | AB | AB | AB | AB | AB | AB | AB | AB | AB | AB | AB | AB | AB | AB | AB | AB | AB | AB | AB |
| AB | AB | AA | - | - | - | AB | AA | - | - | AA | AB | AA | - | AB | AB | AB | AA | AA | AA | AA |
| AA | AA | AB | AB | AA | AB | AB | AB | AB | AB | AB | AA | AA | AA | AA | AB | AA | AA | AB | AA | AA |
| AB | AA | AB | AA | AA | AB | AB | AB | AA | AB | AA | AA | AA | AA | AA | AB | AA | AA | AA | AA | AA |
| AB | AB | AB | AB | AB | AB | AB | AB | AB | AA | AB | AB | AB | AB | AB | AB | AB | AB | AB | AA | AB |
| AB | AB | AB | AA | AA | AB | AB | AA | AA | AB | AA | AB | AB | AB | AB | AA | AB | AB | AB | AB | AA |
| AB | AA | AA | AA | AB | AB | AB | AB | AB | AB | AA | AB | AB | AB | AB | AB | AB | AB | AA | AB | AB |
| AB | AB | AA | AB | AB | AB | AB | AB | AA | AA | AB | AB | AB | - | AB | AB | AA | - | AA | AB | AB |
| - | AA | - | - | - | - | - | AA | AA | AA | AB | AA | AA | - | AA | AA | AA | - | - | AB | AA |
| AB | AB | AB | AB | AB | AB | AB | AB | AB | AB | AB | AB | AB | AB | AB | AB | AB | AB | AB | AB | AB |
| AB | AB | AB | AA | AB | AB | AB | AB | AA | AA | AB | AA | AA | AA | AA | AA | AB | AB | AB | AB | AA |
| - | AB | AA | AA | AB | - | AA | AB | - | AA | AA | AB | AB | - | AB | AA | AB | - | AA | AB | AB |
| AA | AA | AB | AA | AA | - | AB | AB | AA | AB | AA | AB | AA | AA | AA | AB | AB | - | AA | AA | AA |
| AB | AA | AB | AA | AA | - | AA | AB | - | AA | AA | AB | AA | AA | - | AA | - | AB | - | AB | - |
| AB | AB | AB | AB | AB | AB | AB | AB | AB | AB | AB | AB | AB | AB | AB | AB | AB | AB | AB | AB | AB |
| AB | AA | AB | AB | AA | AA | AA | AA | AB | AA | AB | AA | AB | AB | AA | AA | AB | AA | AB | AA | AB |
| AB | AB | AB | AB | AB | AB | AB | AB | AB | AB | AB | AB | AB | AB | AB | AB | AB | AB | AB | AB | AB |
| AB | AB | AB | AB | AB | AB | AB | AB | AB | AB | AB | AB | AB | AB | AB | AB | AB | AB | AB | AB | AB |
| AB | AB | AB | AB | AB | - | AB | - | - | AA | AB | AA | AB | - | - | AB | - | - | AA | AB | AB |
| AB | AB | AA | AB | AB | AB | AA | AB | AB | AB | AB | AA | AB | AB | AA | AB | AB | AB | AB | AB | AB |
| AB | AA | AB | AB | AA | AA | AA | AA | AA | AA | AB | AA | AB | AB | AA | AA | AA | AA | AA | AA | AA |
| AB | AB | AB | AA | AB | AB | AB | AA | AB | AB | AB | AA | AB | AB | AB | AA | AB | AA | AB | AA | AB |
| AA | AB | AB | AB | AB | AB | AB | AA | AB | AB | AB | AB | AB | AB | AB | AB | AA | AB | AB | AA | AA |
| AB | AA | AB | AB | AB | AB | AB | AB | AB | AB | AB | AA | AB | AA | AB | AB | AB | AA | AB | AA | AA |
| AB | AB | AB | AB | AB | AB | AB | AA | AB | AB | AB | AB | AB | AB | AB | AB | AB | AB | AB | AB | AB |
| AB | AA | AA | AA | AA | AA | AA | AA | AA | AA | AA | AA | AA | AA | AA | AA | AA | AB | AA | AA | AA |
| AB | AA | AA | AA | AA | AA | AA | AA | AA | AA | AA | AA | AA | AA | AA | AA | AA | AA | AA | AA | AB |
| AB | AB | AB | AA | AB | AB | AA | AB | AA | AA | AB | AA | AA | AA | AA | AA | AB | AB | AA | AB | AA |
| AA | AA | AB | AA | AA | AA | AA | AA | AA | AA | AB | AA | AA | AA | AA | AA | AA | AA | AA | AA | AA |
| AB | AB | AA | AA | AA | AB | AB | AA | AA | AA | AA | AA | AA | AA | AA | AA | AB | AB | AB | AA | AB |
| AB | - | AA | AA | - | - | AA | - | AA | AA | AB | AB | AA | - | - | AA | AA | AB | AB | AB | AB |
| AA | AA | AB | AB | AB | AA | AB | AB | AA | AB | AB | AA | AB | AA | AA | AB | AA | AB | AB | AA | AA |
| AA | AA | AB | AB | AA | AA | AB | AB | AB | AB | AB | AB | AB | AB | AB | AB | AA | AB | AB | AA | AA |
| AA | AA | AA | AA | AB | AA | AA | AB | AA | AB | AA | AB | AB | AA | AA | AB | AA | AA | AB | AA | AA |
| AA | AB | AB | AB | AA | AA | AA | AB | AA | AB | AA | AB | AA | AA | AB | AA | AB | AA | AB | AA | AA |
| AA | AA | AB | AB | AB | AB | AA | AB | AB | AA | AB | AB | AB | AA | AB | AB | AB | AA | AA | AA | AA |
| AA | AB | AA | AB | AB | AA | AB | AB | AB | AB | AB | AB | AB | AA | AB | AB | AA | AB | AA | AB | AB |
| AB | AA | AA | AA | AA | AA | AA | AB | AB | AB | AA | AB | AB | AA | AA | AB | AA | AA | AB | AA | AB |
| AA | AA | AA | AA | AB | AA | AB | AA | AB | AA | AB | AA | AB | AB | AA | AB | AA | AB | AA | AA | AA |
| - | AB | AB | AA | AB | AB | AA | AB | AB | AB | AB | AB | AB | AA | AB | AB | AA | AB | AA | AB | AB |
| AB | AA | AB | AB | AB | AB | AB | AB | AB | AB | AB | AA | AB | AA | AB | AB | AB | AB | AB | AB | AB |
| AB | AB | AB | AB | AB | AB | AB | AB | AB | AB | AB | AB | AB | AB | AB | AB | AB | AB | AB | AB | AB |
| AA | - | AA | AB | - | - | AB | - | AB | AB | AA | AA | AB | - | - | AB | AB | - | AB | AB | AA |
| AB | AB | AB | AB | AB | AA | AB | AB | AA | AA | AB | AB | AB | AB | AB | AB | AA | AB | AA | AB | AB |
| AB | AA | AB | AA | AA | AA | AB | AA | AB | AB | AB | AA | AB | AA | AB | AB | AA | AA | AA | AA | AA |
| AB | AA | AA | AA | AB | AA | AB | AB | AA | AB | AA | AB | AB | AA | AA | AB | AA | AA | AB | AB | AA |
| AA | AA | AB | AB | AB | AA | AB | AA | AB | AB | AB | AA | AB | AB | AB | AA | AB | AB | AB | AB | AB |
| AB | AA | AB | AB | AB | AA | AB | AA | AB | AB | AB | AB | AA | AB | AB | AA | AB | AA | AB | AB | AA |
| AB | AA | AB | AB | AB | AA | AB | AA | AB | AB | AB | AA | AB | AA | AB | AB | AA | AB | AB | AA | AA |
| AB | AB | AB | AB | AB | AB | AB | AB | AB | AB | AB | AB | AB | AB | AB | AB | AB | AB | AB | AB | AB |
| AA | AA | AA | AA | AA | AB | AA | AA | AB | AA | AB | AA | AA | AA | AB | AB | AA | AB | AA | AB | AB |
| - | AA | AB | AA | AA | AB | AA | AB | AB | AA | AA | AB | AB | AA | AB | AA | AB | AA | AB | AA | AA |
| AB | AA | AB | AA | AB | AB | AB | AB | AA | AA | AB | AB | AB | AA | AB | AA | AB | AB | AB | AA | AB |
| AB | AA | AB | AA | AB | AB | AA | AB | AA | AA | AB | AA | AA | AA | AB | AA | AA | AB | AB | AA | AA |
| AA | AB | AB | AB | AB | AA | AB | AA | AB | AB | AA | AB | AB | AA | AB | AA | AA | AB | AB | AA | AB |
| AB | AB | AB | AB | AB | AA | AB | AA | AB | AB | AB | AA | AB | AB | AB | AB | AB | AB | AA | AA | AB |
| AB | AB | AA | AB | AB | AB | AB | AA | AB | AB | AA | AB | AB | AB | AB | AB | AB | AB | - | AA | AB |
| - | AB | - | AA | AA | - | AB | AA | - | - | AA | AB | AB | - | - | AA | - | AB | AA | AB | AB |
| - | AB | - | AA | - | AB | AA | - | AA | AA | AA | AB | AB | - | - | AA | - | AA | AA | AB | AB |
| AB | AA | AA | AA | AB | AB | AA | AA | AA | AA | AA | AB | AB | AA | AB | AA | AB | AB | AA | AB | AB |
| AA | AB | AA | AB | AA | AA | AB | AB | AB | AB | AA | AB | AB | AB | AA | AB | AB | AA | AB | AA | AB |
| AA | AB | AB | AB | AA | AB | AB | AB | AB | AB | AA | AB | AB | AA | AB | AB | AB | AA | AB | AA | AB |
| AA | AB | AB | AB | AA | AB | AB | AB | AB | AB | AA | AB | AB | AA | AB | AB | AB | AA | AB | AB | AB |
| - | AA | - | AA | - | AA | AA | AA | - | AA | AA | AA | AA | AA | - | AA | AA | - | AA | AB | AA |
| AA | AB | AB | AA | AA | - | AA | AA | AA | AA | AB | AB | AB | AB | - | AA | AA | - | - | AB | AB |
| AB | AB | AB | AA | AA | AA | AB | AB | AB | AB | AB | AB | AB | AB | AA | AB | AA | AB | AB | AA | AB |
| AA | AA | AA | AA | AA | - | AA | AA | AA | AA | AB | AA | AA | AB | AA | AA | AA | AA | AA | AA | AA |
| AB | AB | AB | AB | - | - | AB | AB | AB | AB | AB | AB | AB | - | AB | AB | - | AB | AB | AB | AB |
| AA | AA | AA | AA | - | AA | AA | AA | AA | AA | AA | AA | AA | AA | AA | AA | AA | AA | AA | AA | AA |
| - | AB | AA | AB | AB | - | AB | AB | AA | AA | AA | AA | AA | AA | AA | AB | AB | AB | AA | AB | AA |
| AA | AB | AA | AA | AA | AA | AA | AB | AA | AB | AA | AA | AB | AB | AA | AA | AA | AA | AA | AA | AB |
| AA | AA | AB | AA | AA | AA | AB | AA | AA | AA | AA | AB | AA | AA | AA | AA | AA | AA | AA | AA | AA |
| AB | - | AA | AB | AB | AA | AA | - | - | AA | AA | AA | AA | - | - | AA | - | - | - | AA | AA |
| AB | AA | AA | AA | AA | AB | AA | AA | AA | AA | AA | AB | AA | AB | AB | AA | AA | AB | AA | AA | AA |
| - | AB | - | AB | AB | - | AA | AB | AB | AB | AB | AA | AB | AB | AA | AB | AB | AA | - | AB | AB |
| AA | AB | AB | AA | AA | AB | AA | AA | AB | AB | AB | AA | AB | AB | AB | AB | AB | AA | AB | AB | AB |
| AB | AB | AA | AA | AB | AB | AA | AA | AA | AA | AA | AA | AB | AB | AA | AB | AB | AB | AA | AB | AB |
| AB | AB | AB | AA | - | - | AA | - | AB | AB | AA | AB | AB | - | - | AA | AB | AB | AB | AB | AB |
| AB | AB | AB | AB | AB | AB | AB | AB | AB | AB | AB | AB | AA | AA | AB | AB | AB | AB | AB | AB | AB |
| AA | AA | AA | AA | AA | AA | AA | AA | AA | AA | AA | AA | AA | AB | AA | AA | AB | AB | AA | AA | AA |
| AA | AA | - | AA | - | AA | AA | - | AB | AB | AA | AB | AB | AB | AB | AA | AA | - | AB | AA | AA |
| AB | AB | AA | AB | AB | AB | AB | AA | AA | AA | AA | AA | AB | AA | AA | AA | AA | AB | AA | AB | AB |
| AB | AB | AA | AA | AB | AB | AA | AB | AA | AA | AA | AA | AB | AA | AA | AA | AB | AB | AA | AB | AB |
| AB | AB | AA | AA | AA | AA | AA | AA | AA | AA | AA | AB | AB | AA | AB | AA | AB | AA | AA | AA | AB |
| AA | AA | AA | AA | AA | AB | AA | AA | AA | AA | AA | AA | AB | AB | AA | AA | AA | AA | AA | AA | AA |
| AA | AA | AB | AB | AA | AA | AB | AB | AA | AB | AB | AB | AB | AB | AB | AB | AB | AA | AA | AB | AA |
| AB | AB | AB | AB | AB | AB | AB | AB | AB | AB | AB | AB | AB | AB | AB | AB | AB | AB | AB | AB | AB |
| AB | AA | AA | AA | AB | AA | AA | AA | AB | AA | AA | AA | AB | AA | AA | AA | AA | AA | AB | AB | AB |
| AB | AA | AA | AA | AB | AA | AA | AA | AB | AA | AA | AB | AB | AA | AB | AB | AA | AA | AB | AA | AB |
| AB | AB | AA | AA | AB | AB | AB | AB | AB | AB | AB | AA | AA | AB | AB | AB | AA | AB | AB | AA | AB |
| - | AB | AB | AB | AB | AB | AB | AB | AB | AB | AB | AB | AB | AB | AB | AB | AB | - | AB | AB | AB |
| AB | AB | AB | AA | AB | AB | AA | AB | AB | AA | AA | AB | AA | AA | AA | AA | AB | AB | AB | AA | AB |
| AB | AA | AA | AA | AB | AB | AA | AA | AA | AA | AA | AB | AA | AA | AA | AA | AB | AB | AB | AA | AB |
| AA | AB | AB | AB | AB | AA | AB | AB | AB | AB | AA | AB | AB | AB | AA | AB | AB | AB | AA | AB | AB |
| AB | AA | AB | AB | AB | AA | AA | AA | AA | AA | AB | AA | AB | AB | AB | AA | AB | AB | AB | AA | AA |
| AA | AA | AB | AB | AB | AA | AB | AB | AA | AA | AB | AA | AA | AA | AA | AB | AA | AB | AA | AB | AA |
| AA | AA | AA | AA | AB | AB | AA | AA | AA | AA | AA | AA | AA | AA | AA | AA | AA | AA | AA | AA | AA |
| AB | AB | AB | AB | AB | AB | AB | AB | AB | AA | AB | AB | AA | AB | AB | AB | AB | AB | AB | AB | AB |
| AA | AA | AA | AA | AB | AA | AA | AA | AA | AB | AA | AB | AA | AB | AB | AB | AA | AA | AB | AA | AB |
| AB | AB | AB | AB | AB | AB | AB | AB | AB | AB | AB | AB | AB | AB | AB | AB | AB | AB | AB | AB | AB |
| AB | AB | AB | AA | AB | AB | AB | AB | AB | AA | AB | AA | AA | AA | AA | AA | AB | AB | AA | AB | AA |
| AA | AA | AB | AA | AA | AA | AB | AA | AA | AA | AA | AB | AA | AA | AA | AA | AA | AA | AA | AA | AA |
| AA | AA | AA | AA | AA | AA | AA | AA | AA | AA | AA | AA | AA | AA | AA | AA | AA | AA | AA | AA | AA |
| AA | AA | AB | AB | AA | AA | AA | AB | AB | AB | AB | AB | AB | AA | AB | AB | AB | AB | AB | AA | AA |
| AA | AA | AB | AB | AA | AA | AB | AB | AB | AB | AA | AA | AA | AB | AB | AA | AB | AA | AB | AA | AA |
| AB | AB | AA | AA | AB | AB | AB | AB | AA | AB | AB | AB | AA | AB | AB | AB | AB | AB | AB | AB | AA |
| AB | AB | AB | AA | AB | AB | AB | AB | AA | AB | AB | AB | AA | AB | AB | AB | AB | AA | AB | AB | AA |
| AB | AB | AB | AB | AB | - | AB | AB | AB | AB | - | AB | AB | AB | AB | AB | AB | AB | - | AB | AB |
| AA | AA | AB | AA | AA | AA | AA | AA | AA | AA | AA | AB | AA | AA | AA | AA | AB | AA | AA | AA | AA |
| AA | AA | AB | AB | - | AA | AA | AB | AB | AB | AA | AB | AA | AA | AB | AA | AB | AA | - | AB | AA |
| AB | AA | AA | AA | AB | - | AA | AB | AB | AA | AA | AA | AA | AA | - | AB | AA | AA | AB | AA | AB |
| AA | AA | AA | AA | AA | AA | AA | AA | AA | AA | AA | AA | AA | AA | AB | AA | AA | AA | AA | AA | AA |
| AA | AA | AA | AA | AB | - | AA | AB | AA | AB | AB | AB | AB | AA | AB | AB | AB | AB | AB | AB | AA |
| AB | AB | AA | AA | AB | AB | AA | AA | AB | AB | AB | AB | AB | AB | AB | AB | AB | AB | AB | AB | AA |
| AB | AB | AA | AA | AB | AA | AA | AB | AA | AB | AA | AA | AB | AB | AB | AB | AB | AA | AB | AB | AB |
| AB | AB | AA | AA | AB | AB | AA | AB | AB | AB | AA | AA | AB | AB | AB | AB | AB | AA | AB | AA | AB |
| AB | AB | AA | AB | AB | AB | AA | AA | AB | AB | AA | AA | AB | AA | AB | AB | AB | AA | AB | AB | AB |
| AB | AB | AB | AB | - | - | AB | - | AB | AB | AB | AB | AB | - | AB | AB | - | AB | AB | AB | - |
| AB | AB | AA | AA | AA | AA | AA | AB | AB | AB | AA | AA | AB | AA | AB | AA | AB | AA | AA | AB | AA |
| AB | AA | AA | AA | AB | AB | AB | AA | AB | AA | AA | AA | AB | AA | AA | AA | AA | AB | AA | AA | AB |
| AB | AB | AA | - | - | - | - | AB | AB | AB | AB | AB | AB | AB | AB | AB | AB | - | AB | AB | AB |
| - | AA | AA | AB | - | AA | AA | AA | AA | AA | AA | AA | AB | AA | AA | AA | AA | - | AA | AB | AA |
| AA | AA | AB | AB | AA | AA | AB | AB | AB | AA | AA | AB | AB | AA | AB | AB | AB | AB | AB | AB | AA |
| AB | AA | AB | AB | AB | AA | AB | AB | AB | AA | AA | AB | AA | AA | AA | AB | AB | AA | AB | AA | AB |
| AA | AA | AA | AA | AA | - | AA | AA | AA | AA | AA | AA | AA | AA | AA | AA | AA | AA | AA | AA | AA |
| AA | AA | - | AA | - | AA | AA | AA | AA | AA | AA | AA | AA | AA | - | - | AA | AA | AA | AB | AA |
| AB | AB | AA | AA | AA | AA | AA | AA | AA | AA | AA | AB | AB | AA | AA | AA | AA | AA | AB | AA | AA |
| AB | AB | AA | AA | AA | AA | AA | AA | AA | AB | AB | AA | AA | AB | AB | AA | AA | AA | AB | AB | AA |
| AB | AB | AB | AB | AB | AB | AB | AB | AB | AB | AB | AB | AB | AB | AB | AB | AB | AB | AB | AB | AB |
| AA | AB | AA | AB | AA | AB | AA | AB | AB | AB | AA | AA | AB | AB | AA | AB | AB | AB | AB | AB | AA |
| - | AA | - | AA | - | AA | AA | AA | AA | AA | AA | AA | AA | AA | AA | AA | AA | AA | AA | AA | AA |
| AB | AB | AA | AA | AA | AB | AA | AA | AB | - | AA | AB | AB | AB | AB | AA | AB | - | AB | AA | AB |
| AA | AB | AB | AA | AA | AA | AB | AB | AB | AB | AB | AA | AB | AB | AA | AA | AB | AA | AB | AA | AB |
| AB | AB | AB | AB | AB | AB | AB | AB | AB | AB | AA | AB | AB | AB | AB | AB | AB | AA | AB | AA | AB |
| AB | AB | AA | AA | AB | AB | AA | AB | AB | AB | AB | AA | AB | AB | AA | AB | AA | AA | AB | AA | AB |
| AB | AB | AA | AB | - | AB | AA | AB | AB | AB | AA | AB | AB | - | AB | AB | AB | AA | AB | AA | AB |
| AB | AB | AB | AA | AB | AB | AB | AA | AA | AA | AA | AB | AB | AA | AA | AA | AA | AB | AB | AA | AB |
| AB | AB | AA | AB | AB | AA | AB | AB | AB | AB | AA | AB | AB | AB | AB | AA | AB | AB | AB | AB | AA |
| AA | AB | AA | AA | AA | - | AA | - | AA | - | AA | AA | AB | - | - | AA | - | - | - | AB | AB |
| AA | AB | AA | AB | AA | AA | AB | AA | AA | AB | AB | AB | AA | AB | AB | AA | AB | AA | AA | AB | AA |
| - | AB | AB | AB | AB | AA | AB | AA | AA | AB | AB | AB | AB | AA | AA | AB | AB | - | AA | AB | AB |
| AA | AA | AB | AA | AB | AA | AA | AA | AA | AB | AB | AA | AB | AA | AB | AB | AB | AA | AA | AB | AB |
| AA | AA | AA | AA | AB | AA | AA | AA | AA | AB | AA | AA | AA | AA | AB | AA | AA | AA | AA | AB | AA |
| AA | AA | AA | AB | AB | AB | AB | AB | AA | AA | AB | AA | AB | AA | AA | AA | AB | - | AA | AB | AA |
| AB | AB | AB | AA | AB | AB | AB | AB | AB | AB | AA | AB | AB | AB | AB | AB | AB | AB | AB | AB | AB |
| AA | AB | AB | AB | AA | AB | AA | AB | AB | AA | AA | AB | AA | AA | AB | AB | AB | AA | AB | AB | AA |
| AB | AB | AA | AB | AB | AB | AB | AB | AA | AB | AA | AB | AA | AB | AB | AB | AB | AB | AB | AB | AA |
| AB | AB | AA | AB | AB | AB | AA | AA | AA | AA | AB | AB | AB | AA | AA | AA | AB | AB | AB | AB | AB |
| AA | AB | AA | AB | AA | AA | AA | AA | AA | AA | AB | AB | AA | AB | AB | AA | AB | AB | AA | AB | AA |
| - | AB | AB | AA | AA | AA | AA | AA | AA | AA | AA | AB | AB | AB | AA | AA | AA | AB | AA | AB | AA |
| - | AA | - | AA | AA | - | AA | AB | AB | AA | AA | AB | AB | AA | - | - | AA | AB | - | AA | AB |
| AB | AB | AA | AA | AB | AB | AA | AA | AB | AA | AA | AA | AB | AB | AA | AB | AB | AB | AA | AB | AB |
| AB | AA | AB | AB | AA | AB | AA | AA | AA | AA | AB | AA | AB | AB | AA | AB | - | AB | AA | AA | AB |
| AB | AB | AB | AB | AB | AB | AB | AB | AB | AB | AB | AB | AB | AB | AB | AB | AB | AB | AB | AB | AB |
| AA | AB | AA | AB | AA | AB | AA | AA | AB | AA | AB | AA | AB | AA | AA | AA | AB | AA | AB | AA | AA |
| - | AB | AA | AA | AB | AB | AA | AA | - | AA | AA | AB | AB | AA | - | AA | AB | AB | - | AB | AB |
| AA | AB | AB | AB | AB | AB | AA | AB | AB | AB | AB | AA | AB | AA | AB | AA | AB | AA | AB | AB | AA |
| AB | AB | AB | AB | AB | AB | AA | AA | AA | AA | AB | AA | AB | AA | AB | AA | AB | AB | AB | AB | AA |
| - | AB | AB | AB | AB | AB | AB | AA | AA | AA | AB | AA | AB | AB | AB | AA | AB | AB | AB | AA | AB |
| AB | AB | AA | AA | AA | AA | AA | AA | AA | AA | AA | AB | AA | AA | AA | AA | AB | AB | AB | AA | AA |
| - | AB | AB | AB | AB | AB | AB | AB | AB | AB | AB | AB | AB | AB | AB | AB | AB | AB | AB | AB | AB |

| snp_B1215 | snp_B1226 | snp_B1240 | snp_B1243 | snp_B1256 | snp_B1263 | snp_B1294 | snp_B1295 | snp_B1300 | snp_B1317 | snp_B1339 | snp_B1349 | snp_B1368 | snp_B1403 | snp_B1406 | snp_B1410 | snp_B1476 | snp_B1479 | snp_B1483 | snp_B1488 | snp_B1497 | snp_B1504 | snp_B1517 | snp_B1518 | snp_B1536 | snp_B1539 | snp_B1540 | snp_B1551 |
|---|---|---|---|---|---|---|---|---|---|---|---|---|---|---|---|---|---|---|---|---|---|---|---|---|---|---|---|
| 5 | 5 | 5 | 5 | 5 | 5 | 5 | 5 | 5 | 5 | 5 | 5 | 5 | 5 | 5 | 5 | 5 | 5 | 5 | 5 | 5 | 5 | 5 | 5 | 5 | 5 | 5 | 5 |
| -- | -- | -- | -- | -- | LG13 | -- | -- | -- | -- | -- | -- | -- | -- | -- | -- | -- | -- | -- | -- | -- | -- | -- | -- | -- | -- | -- | -- |
| AA | AB | AA | AB | AA | AA | AB | AB | AB | AB | AA | AA | AB | AA | AB | AA | AB | AB | AA | AB | AB | AB | AB | AB | AB | AA | AA | AB |
| AB | AB | AB | AA | AA | AB | AB | AA | AB | AA | AB | AA | AB | AB | AA | AB | AB | AB | AA | AB | AA | AB | AB | AB | AA | AA | AB | AB |
| AB | AA | AA | AA | AA | AB | AA | AB | AB | AA | AB | AB | AA | AB | AB | AA | AB | AB | AB | AB | AA | AB | AB | AB | AA | AA | AB | AB |
| AB | AA | AB | - | AA | AA | AA | AA | AB | AA | AB | AB | AA | AA | AB | AA | AB | AB | AB | AB | AA | AB | - | AB | AA | AA | AB | AA |
| AB | AA | AB | AA | AB | AA | AA | AB | AA | AB | AA | AA | AB | AA | AB | AA | AB | AB | AA | AA | AB | AB | AB | AB | AA | AA | AB | AB |
| - | AA | AA | - | - | - | - | AB | AA | - | - | - | AB | AA | AA | - | AA | AA | - | AA | AB | AB | - | AB | AB | AA | - | AA |
| AB | AA | AB | AA | AB | AB | AB | AB | AA | - | AA | AB | AB | AB | AB | AB | AA | AA | AB | AB | AB | AB | AA | AB | AB | AB | AB | AA |
| AA | AA | AB | AA | AA | AA | AA | AA | AA | AA | AA | AA | AB | AA | AB | AB | AA | AA | AB | AB | AA | AA | AA | AA | AB | AA | AB | AA |
| AA | AA | AB | AA | AA | AA | AB | AA | AA | AB | AA | AB | AB | AB | AA | AB | AA | AB | AA | AA | AB | AA | AA | AA | AA | AB | AB | AA |
| AA | AA | AA | AB | AA | AB | AA | AB | AA | AB | AA | AB | AB | AB | AA | AB | AB | AB | AA | AB | AB | AA | AB | AB | AA | AB | AB | AA |
| AB | AA | AB | AB | AB | AB | AB | AB | AB | AA | AB | AA | AB | AB | AA | AB | AB | AB | AA | AB | AB | AB | AB | AA | AA | AB | AB | AB |
| AB | AB | AB | AB | AB | AA | AB | AB | AB | AB | AB | AA | AA | AA | AA | AB | AB | AB | AA | AB | AB | AB | AB | AB | AB | AB | AB | AB |
| AB | AB | AB | AA | AB | AA | AB | AB | AB | AB | AA | AA | AB | AB | AA | AA | AB | AA | AB | AB | AA | AA | AA | AA | AA | AB | AA | AB |
| AB | AA | AA | AB | AB | AA | AB | AA | AB | AB | AA | AA | AB | AB | AA | AB | AB | AA | AB | AB | AB | AA | AB | AB | AA | AB | AB | AB |
| AA | AB | AB | AB | AB | AB | AB | AA | AA | AB | AB | AB | AB | AB | AA | AB | AB | AB | AB | AA | AA | AB | AB | AB | AA | AB | AA | AA |
| AB | AB | AB | AB | AB | AB | AB | AB | AB | AA | AA | AB | AB | AB | AB | AA | AB | AB | AB | AA | AB | AA | AB | AA | AB | AB | AB | AA |
| AA | AA | AA | AA | AA | AA | AA | AA | AA | AA | AA | AA | AA | AA | AA | AA | AA | AA | AA | AA | AA | AA | AA | AA | AA | AA | AA | AA |
| - | AB | AA | - | AB | - | - | AB | AA | - | AA | - | AA | AB | - | - | AA | AA | AB | - | AA | AA | - | AB | AB | AA | AB | AA |
| AB | AB | AB | AB | AB | AB | AB | AB | AB | AB | AB | AB | AB | AB | AB | AB | AA | AA | AB | AB | AB | AA | AA | AB | AB | AB | AB | AB |
| AB | AB | AB | AB | AB | AB | AB | AB | AB | AB | AB | AB | AB | AB | AB | AB | AB | AB | AB | AB | AB | AB | AB | AB | AB | AB | AB | AB |
| AB | AB | AB | AB | AB | AB | AB | AB | AB | AB | AB | AB | AB | AB | AB | AB | AB | AB | AB | AB | AB | AB | AB | AB | AB | AB | AB | AB |
| AB | AB | AB | AB | AB | AB | AB | AB | AB | AB | AB | AB | AB | AB | AB | AB | AB | AB | AB | AB | AB | AB | AB | AB | AB | AB | AB | AB |
| AB | - | AB | - | AB | - | AB | AB | AB | - | - | - | AB | AB | AB | AB | AA | AA | AB | AB | AA | AA | - | AB | AB | AB | - | AB |
| AB | AB | AA | AB | AA | AB | AB | AA | AB | AB | AB | AB | AB | AA | AA | AA | AB | AB | AA | AB | AA | AA | AB | AB | AA | AA | AA | AB |
| AB | AA | AB | AB | AA | AB | AA | AB | AA | AB | AB | AB | AA | AA | AA | AA | AA | AA | AB | AB | AA | AA | AB | AA | AB | AA | AA | AB |
| AB | AB | AB | AB | AB | AB | AB | AB | AB | AB | AB | AB | AB | AB | AB | AB | AB | AB | AB | AB | AB | AB | AB | AB | AB | AB | AB | AB |
| AA | AA | AA | AA | AA | AA | AB | AA | AA | AA | AA | AA | AA | AA | AA | AB | AA | AA | AA | AA | AA | AB | AB | AA | AA | AA | AB | AA |
| AB | AB | AB | AB | AB | AB | AB | AB | AB | AB | AB | AB | AB | AB | AA | AB | AB | AB | AB | AB | AB | AB | AB | AB | AB | AB | AB | AB |
| AA | AA | AB | - | AA | - | AB | AB | AA | AB | AB | AB | AB | AB | AB | AB | AA | AA | AB | AA | AB | AA | AA | AA | AB | AA | AA | AA |
| - | - | AA | - | - | - | AA | AA | AA | AB | AB | - | AA | AB | AB | AA | AB | AB | AA | - | AA | AA | AB | AB | AA | AB | AA | AB |
| AB | AB | AB | AB | AB | AB | AB | AB | AB | AB | AB | AB | AB | AB | AB | AB | AB | AB | AB | AB | AB | AB | AB | AB | AB | AB | AB | AB |
| AB | AB | AB | AA | AB | AA | AB | AB | AB | AA | AA | AA | AA | AB | AB | AB | AA | AA | AA | AA | AB | AB | AA | AA | AB | AB | AA | AA |
| - | AA | AB | - | AB | AA | AB | AB | AA | AA | AA | - | AA | AB | AB | AA | AA | AA | AB | AA | AB | AB | - | AB | AB | AB | AA | AA |
| AA | AB | AA | AB | AA | - | AB | AB | AB | AB | AB | AB | AB | AA | AB | AB | AB | AB | AB | AB | AB | AA | - | AB | AB | AA | AB | AB |
| - | AB | AB | AB | AA | - | AB | AA | AB | AA | AB | AB | AB | AB | - | AB | AA | AB | AA | - | AA | AB | - | AB | AA | AB | - | AA |
| AB | AB | AB | AB | AB | - | AB | AB | AB | - | AB | AB | AB | AB | AB | AB | AB | AB | AB | AB | AB | AB | AB | AB | AB | AB | AB | AB |
| AA | AA | AA | AA | AA | AB | AA | AA | AA | AA | AA | AA | AA | AA | AA | AA | AA | AA | AA | AA | AB | AB | AB | AB | AB | AB | AB | AA |
| AB | AB | AB | AB | AB | AB | AB | AB | AB | AB | AB | AB | AB | AB | AB | AB | AB | AB | AB | AB | AB | AB | AB | AB | AB | AB | AB | AB |
| AB | AB | AB | AB | AB | AB | AB | AB | AB | AB | AB | AB | AB | AB | AB | AB | AB | AB | AB | AB | AB | AB | AB | AB | AB | AB | AB | AB |
| - | AB | AB | AA | AB | - | AB | AB | AB | - | AB | AA | AB | AB | AA | AA | AB | AB | AB | AB | - | AB | AB | AB | AA | AB | AB | AB |
| - | AB | AB | AB | AB | AB | AB | AB | AB | AA | AB | AB | AB | AB | AB | AB | AA | AB | AB | AB | AB | AB | AA | AB | AB | AB | AB | AA |
| AA | AB | AA | AA | AA | AB | AA | AA | AB | AB | AB | AA | AA | AA | AA | AA | AB | AB | AA | AA | AA | AB | AA | AA | AA | AA | AA | AB |
| AA | AB | AA | AB | AA | AB | AA | AA | AB | AA | AB | AB | AA | AA | AB | AA | AB | AB | AB | AA | AB | AA | AB | AA | AA | AA | AB | AA |
| AB | AB | AB | AB | AB | AB | AB | AA | AB | AA | AB | AA | AB | AA | AB | AA | AA | AA | AB | AA | AB | AA | AA | AB | AB | AA | AB | AB |
| AA | AB | AB | AA | AB | AB | AB | AB | AB | AB | AB | AA | AB | AB | AB | AB | AB | AB | AB | AA | AA | AB | AB | AA | AB | AB | AA | AA |
| AB | AB | AB | AB | AB | AB | AB | AB | AB | AB | AB | AB | AB | AB | AB | AB | AB | AB | AB | AB | AB | AB | AB | AB | AB | AB | AB | AB |
| AA | AA | AA | AA | AA | AA | AA | AA | AA | AA | AA | AA | AA | AA | AA | AA | AA | AA | AA | AA | AA | AA | AA | AA | AA | AA | AA | AA |
| AB | AA | AA | AA | AA | AA | AA | AA | AA | AA | AA | AA | AA | AA | AA | AA | AA | AA | AA | AA | AA | AA | AA | AA | AA | AA | AA | AA |
| - | AA | AB | AA | AB | AA | AB | AA | AA | AA | AA | AA | AA | AB | AB | AB | AA | AB | AB | AA | AA | AB | AB | AB | AA | AB | AA | AA |
| AA | AA | AB | AA | AA | AB | AA | AA | AA | AA | AA | AA | AA | AB | AB | AB | AA | AA | AB | AA | AA | AB | AB | AB | AA | AA | AA | AA |
| AB | AA | AB | AA | AA | AA | AB | AA | AB | AB | AB | AA | AB | AB | AB | AB | AA | AA | AB | AA | AA | AB | AB | AA | AB | AA | AB | AB |
| AB | AA | AB | - | AA | - | AA | AB | AA | AA | AA | AA | AB | AB | AB | AA | AA | AA | AB | AB | AB | AB | AA | AA | AB | AA | AB | AA |
| AB | AB | AB | AB | AA | AB | AB | AB | AB | AB | AB | AB | AB | AB | AB | AB | AB | AB | AA | AA | AB | AB | AA | AB | AB | AB | AB | AB |
| AB | AB | AA | AA | AA | AB | AB | AB | AB | AB | AB | AB | AB | AA | AA | AB | AB | AB | AB | AB | AB | AA | AB | AB | AB | AA | AB | AA |
| AB | AB | AA | AB | AA | AB | AB | AB | AA | AB | AB | AB | AB | AA | AA | AB | AB | AB | AB | AA | AA | AB | AB | AA | AA | AB | AB | AB |
| AB | AA | AB | AA | AA | AB | AB | AA | AB | AA | AA | AA | AB | AA | AA | AA | AB | AA | AA | AA | AB | AA | AA | AB | AA | AA | AA | AA |
| AA | AA | AA | AA | AA | AA | AA | AB | AA | AB | AB | AB | AB | AA | AA | AA | AB | AB | AA | AB | AB | AB | AB | AA | AA | AA | AB | AB |
| AB | AB | AA | AB | AA | AA | AB | AA | AB | AB | AB | AB | AB | AA | AA | AB | AB | AB | AA | AB | AB | AB | AB | AB | AA | AA | AB | AB |
| AB | AA | AA | AA | AB | AB | AB | AB | AA | AA | AA | AB | AA | AA | AA | AB | AA | AA | AA | AB | AA | AA | AB | AA | AB | AA | AA | AA |
| AB | AB | AA | AB | AB | AB | AB | AB | AB | AB | AA | AB | AB | AA | AA | AB | AA | AA | AB | AB | AB | AB | AB | AA | AB | AB | AA | AB |
| AA | AB | AB | AB | - | - | AB | AB | AB | AB | AA | AB | AB | AB | AA | AB | AB | AB | AB | AA | AA | AB | AB | AB | AB | AA | AA | AA |
| AA | AB | AB | AB | AB | AB | AB | AB | AB | AB | AB | AB | AB | AB | AB | AB | AB | AB | AB | AB | AB | AB | AB | AB | AB | AA | AA | AB |
| AB | AB | AB | AB | AB | AB | AB | AB | AB | AB | AB | AB | AB | AB | AA | AB | AB | AB | AB | AB | AB | AB | AB | AB | AB | AB | AA | AB |
| AB | AA | AB | - | - | - | AB | AA | AB | AB | AB | - | AB | AB | AA | - | AA | AA | AB | AB | AB | AA | - | AA | AA | AA | - | AB |
| AB | AB | AB | AB | AB | - | AB | AB | AB | AB | AB | AA | AA | AA | AB | AB | AB | AB | AB | AB | AB | AB | AA | AA | AB | AB | AB | AB |
| AA | AA | AA | AA | AA | AB | AA | AA | AA | AA | AB | AA | AA | AA | AA | AA | AA | AA | AA | AA | AA | AA | AB | AA | AB | AA | AA | AB |
| AA | AB | AB | AB | AB | AB | AB | AB | AB | AB | AB | AB | AB | AB | AB | AB | AA | AA | AB | AB | AB | AB | AB | AB | AB | AB | AA | AB |
| AB | AB | AB | AB | AB | AA | AB | AB | AB | AB | AB | AB | AB | AB | AB | AB | AB | AB | AB | AB | AB | AB | AB | AB | AB | AB | - | AB |
| AA | AB | AB | AB | AA | AB | AB | AA | AB | AA | AA | AB | AB | AB | AA | AB | AA | AA | AB | AA | AB | AB | AA | AB | AA | AA | AB | AA |
| AA | AB | AB | AB | AA | AB | AB | AB | AA | AB | AA | AA | AB | AB | AA | AB | AB | AA | AB | AA | AB | AB | AA | AA | AA | AB | AB | AA |
| AB | AB | AB | AB | AB | AB | AB | AB | AB | AB | AB | AB | AB | AB | AB | AB | AB | AB | AB | AB | AB | AB | AB | AB | AB | AB | AB | AB |
| AB | AA | AA | AA | AA | AA | AA | AA | AA | AA | AA | AA | AA | AA | AA | AA | AA | AA | AA | AA | AA | AA | AA | AA | AA | AA | AA | AA |
| AA | AA | AB | AA | AA | AB | AB | AA | AB | AB | AB | AB | AA | AB | AB | AB | AA | AB | AB | AB | AB | - | AB | AB | AB | AA | AA | AA |
| AB | AB | AA | AB | AB | AB | AB | AB | AA | AB | AA | AB | AB | AA | AB | AA | AA | AA | AB | AB | AB | AA | AA | AB | AB | AB | AB | AA |
| AA | AB | AA | AB | AB | AA | AB | AB | AA | AB | AB | AA | AB | AA | AA | AB | AA | AA | AB | AB | AA | AA | AB | AB | AA | AA | AA | AA |
| AB | AB | AB | AB | AA | AA | AB | AB | AB | AA | AB | AA | AA | AB | AA | AB | AA | AB | AA | AB | AA | AA | AB | AB | AA | AA | AA | AA |
| AB | AA | AB | AA | AA | AB | AB | AA | AA | AB | AB | AA | AB | AB | AB | AA | AB | AB | AB | AA | AB | AB | AB | AA | AB | AA | AA | AB |
| AB | AB | AB | AA | AB | AA | AB | AA | AA | - | AB | AA | AB | AA | AB | AB | AA | AB | AB | AB | AA | AB | AB | AA | AB | AB | AA | AB |
| AB | AB | AA | AA | AB | AB | AB | AB | AB | AA | AA | AB | AB | AB | AB | AB | AA | AB | AB | AB | AB | AA | AA | AB | AB | AB | AB | AB |
| AA | AA | AB | AA | AB | - | AA | AB | AA | - | - | AA | AA | AB | AB | AA | AA | AA | AB | AA | AB | AB | - | AB | AB | AB | AB | AA |
| AA | AA | AB | AA | AB | AA | AA | AB | AA | AA | AA | AA | AA | AB | AA | AA | AA | AA | AA | AB | AB | AB | - | AB | AB | AB | AB | AA |
| AB | AB | AA | AB | AA | AB | AB | AA | AB | AB | AB | AB | AB | AA | AA | AB | AA | AA | AB | AB | AA | AA | AA | AA | AB | AA | AB | AB |
| AB | AB | AA | AB | AA | AB | AB | AB | AB | AA | AB | AB | AB | AB | AA | AA | AB | AB | AB | AB | AA | AB | AB | AA | AB | AA | AB | AB |
| AA | AB | AA | AB | AB | AB | AB | AB | AB | AB | AB | AB | AB | AB | AA | AB | AB | AA | AB | AB | AB | AA | AA | AB | AA | AA | AB | AA |
| AA | - | AB | - | AB | - | AA | AB | - | - | AA | AA | AA | AB | AB | AB | AA | AA | AA | AB | AB | AA | - | AA | AB | AB | - | AA |
| - | AA | AA | - | - | - | - | AB | AA | AB | AB | AA | AA | AB | AA | AB | AB | AB | AA | AB | AA | AB | - | AA | AB | AA | AB | AA |
| AB | AB | AB | AB | AB | AA | AB | AB | AA | AA | AA | AB | AB | AB | AB | AA | AB | AB | AA | AA | AB | AB | AB | AA | AB | AA | AB | AA |
| AA | AA | AA | AA | AA | AA | AA | AA | AA | AA | AA | AA | AA | AA | AA | AA | AA | AA | AA | AA | AB | AA | AA | AA | AA | AA | AB | AA |
| AB | AB | AB | AB | - | - | AB | AA | AB | - | AB | AB | AB | AB | AB | AB | AB | AB | AB | - | AB | AB | AB | AB | AA | AB | - | AB |
| AB | AA | AA | - | AA | AA | AA | AB | AB | AA | - | AA | AA | AA | AB | AA | AA | AA | AA | AA | AA | AA | - | AA | AA | AA | AA | AB |
| AB | AB | AB | AB | AB | AA | AB | AA | AB | AB | AB | AB | AB | AA | AB | AA | AA | AB | AB | AA | AB | AB | AA | AA | AB | AA | AB | AB |
| AA | AA | AA | AA | AA | AB | AA | AA | AA | AA | AA | AA | AA | AA | AA | AA | AB | AB | AA | AA | AA | AB | AA | AA | AA | AA | AB | AA |
| AB | AA | AA | AA | AA | AB | AA | AB | AA | AB | AB | AB | AA | AA | AA | AA | AB | AB | AB | AA | AA | AA | AB | AA | AB | AA | AA | AB |
| AA | - | AB | - | AB | - | - | AA | AB | - | AA | - | AA | AA | AB | AB | AA | AA | AA | AA | AA | - | AA | AA | AA | AB | AB | AA |
| AA | AA | AA | AA | AA | AA | AA | AA | AA | AA | AA | AA | AA | AA | AA | AB | AA | AA | AA | AB | AA | AA | AA | AA | AA | AA | AA | AA |
| AA | AB | AB | AB | AA | AA | AB | AA | AB | - | AA | - | AB | - | AB | AB | AA | AB | AB | AB | AB | AA | AA | - | AA | AB | - | AA |
| AA | AA | AA | AA | AA | AB | AA | AA | AA | AA | AA | AB | AB | AA | AA | AB | AB | AB | AB | AA | AB | AB | AB | AB | AA | AA | AB | AA |
| AA | AA | AB | - | AB | AA | AA | AB | AA | AA | AA | AA | AA | AB | AB | AA | AA | AA | AB | AB | AA | - | AB | AB | AB | AB | AB | AA |
| AB | AB | AB | - | - | - | AB | AB | AA | AA | AA | AA | AA | AB | AB | AB | AA | AA | AB | - | AB | AB | - | AB | AB | AB | - | AB |
| AB | AB | AB | AB | AB | AB | AB | AB | AB | AB | AB | AB | AB | AB | AB | AA | AB | AB | AB | AB | AB | AB | AB | AA | AB | AB | AA | AB |
| AA | AA | AA | AA | AA | - | AA | AB | AA | AA | AA | AA | AA | AA | AA | AA | AA | AA | AA | AA | AA | AA | AA | AA | AB | AA | AA | AA |
| - | AA | AA | AA | - | - | AA | AA | AA | - | AA | - | AA | - | AA | AB | AA | AA | AB | - | AA | AB | - | AA | AA | AA | AA | AA |
| AB | AA | AA | AA | AA | AA | AA | AB | AA | AA | AB | AA | AA | AA | AB | AA | AA | AA | AA | AA | AB | AB | AB | AA | AB | AB | AB | AB |
| AB | AA | AB | AA | AB | AA | AA | AB | AA | AA | AB | AA | AA | AB | AB | AA | AA | AA | AA | AA | AB | AB | AB | AA | AA | AB | AB | AB |
| AA | AA | AA | AA | AB | AA | AA | AB | AA | AA | AA | AA | AA | AA | AA | AB | AA | AA | AB | AA | AB | AB | AA | AB | AA | AA | AB | AA |
| AA | AA | AA | AA | AA | AA | AA | AB | AA | AA | AA | AA | AA | AA | AA | AA | AA | AA | AA | AA | AA | AA | AA | AA | AB | AA | AA | AA |
| AA | AB | AA | AB | AA | AB | AA | AA | AB | AB | AB | AB | AB | AA | AA | AB | AB | AB | AB | AB | AB | AA | AA | AB | AA | AA | AA | AB |
| AB | AB | AB | AB | AB | AB | AB | AB | AB | AB | AB | AB | AB | AB | AB | AB | AB | AB | AB | AB | AB | AB | AB | AB | AB | AB | AB | AB |
| AB | AA | AA | AA | AB | AA | AA | AB | AB | AB | AA | AA | AA | AB | AB | AB | AA | AA | AB | AB | AB | AB | AB | AB | AA | AB | AA | AA |
| AB | AA | AB | AA | AB | AA | AA | AB | AA | AA | AA | AA | AA | AA | AB | AB | AA | AA | AA | AB | AA | AA | - | AA | AB | AB | AA | AA |
| AB | AB | AB | AB | AB | AB | AB | AB | AB | - | AB | AB | AB | AB | AB | AB | AB | AB | - | AB | AB | AB | AB | AB | AB | AB | - | AB |
| AB | AA | AB | AA | AB | AA | AB | AB | AB | AB | AA | AA | AA | AB | AB | AA | AB | AB | AA | AA | AB | AB | AB | AA | AB | AB | AA | AA |
| AA | AA | AA | AA | AA | AA | AA | AA | AB | AA | AA | AA | AA | AB | AB | AA | AA | AA | AB | AB | AB | AB | AA | AA | AB | AB | AA | AA |
| AB | AB | AB | AB | AB | AA | AB | AB | AB | AB | AB | AB | AB | AA | AB | AB | AB | AB | AB | AA | AA | AB | AB | AB | AB | AB | AB | AB |
| AB | AA | AB | AA | AB | AA | AB | AB | AB | AB | AA | AB | AA | AA | AB | AB | AA | AB | AB | AA | AB | AA | AB | AA | AB | AB | AB | AA |
| AB | AB | AB | AB | AB | AB | AB | AB | AB | AB | AB | AA | AA | AB | AB | AB | AA | AB | AB | AA | AA | AB | AA | AB | AB | AB | AB | AB |
| AA | AA | AA | AA | AA | AB | AA | AA | AA | AA | AA | AA | AA | AA | AA | AA | AA | AA | AA | AA | AA | AA | AA | AA | AA | AA | AA | AA |
| AB | AB | AB | AB | AB | AB | AB | AB | AB | AB | AB | AA | AA | AB | AB | AB | AB | AA | AB | AB | AB | AA | AA | AB | AB | AB | AB | AB |
| AB | AB | AA | AA | AA | AB | AA | AA | AA | AA | AA | AB | AB | AA | AA | AB | AA | AB | AB | AA | AB | AA | AB | AA | AB | AA | AB | AA |
| AB | AB | AB | AB | AB | AB | AB | AB | AB | AB | AB | AB | AB | AB | AB | AB | AB | AB | AB | AB | AB | AB | AB | AB | AB | AB | AB | AB |
| AB | AA | AB | AA | AB | AA | AB | AB | AB | AA | AB | AA | AA | AA | AB | AB | AA | AB | AB | AA | AA | AB | AB | AB | AA | AB | AB | AA |
| AA | AA | AA | AA | AA | AB | AA | AA | AA | AB | AB | AA | AA | AA | AA | AA | AA | AB | AB | AA | AA | AA | AB | AA | AA | AA | AA | AB |
| AA | AA | AA | AA | AA | AA | AA | AA | AA | AA | AA | AA | AA | AA | AA | AA | AA | AA | AA | AA | AA | AA | AA | AA | AA | AA | AA | AA |
| AA | AB | AA | AB | AA | AB | AB | AB | AB | AB | AA | AB | AB | AA | AA | AB | AB | AA | AB | AB | AB | AA | AA | AA | AB | AB | AA | AA |
| AB | AB | AA | AB | AA | AB | AB | AA | AB | AB | AA | AB | AA | AB | AA | AA | AB | AB | AA | AB | AB | AB | AA | AB | AA | AA | AA | AB |
| AB | AB | AB | AB | AB | AB | AB | AB | AB | AB | AB | AB | AB | AB | AB | AA | AB | AA | AB | AB | AB | AA | AA | AB | AB | AB | AB | AB |
| AA | AB | AB | AB | AB | AB | AB | AA | AB | AA | AA | AB | AB | AB | AA | AB | AB | AA | AA | AB | AB | AB | AB | AA | AB | AB | AB | AA |
| AB | AB | AB | AB | AB | - | AB | AB | AB | AB | - | AB | AB | AB | AB | AB | AB | AB | AB | AB | AB | AB | AB | - | AA | AB | AB | AB |
| AA | AA | AA | AA | AA | AB | AA | AA | AA | AA | AA | AA | AA | AA | AA | AA | AA | AA | AA | AA | AA | AA | AB | AB | AA | AA | AA | AA |
| AB | AB | AA | AB | AA | AB | AB | AA | AB | AB | AB | AB | AB | AA | AA | AA | AB | AB | AB | AB | AB | AA | AA | AB | AA | AA | AA | AA |
| AB | AB | AB | AB | AB | AB | AB | AB | AB | AA | AA | AA | AA | AA | AB | AB | AB | AA | AA | AA | AB | AB | AA | AA | AB | AB | AB | AA |
| AA | AA | AA | AA | AA | AA | AA | AB | AA | - | AA | AA | AA | AA | AA | AA | AA | AA | AA | AA | AA | AA | AA | AA | AB | AB | - | AA |
| - | AA | AA | AA | AA | - | AB | AB | AA | AA | AA | AA | AB | AB | AB | AB | AA | AA | AA | AB | AB | AA | AA | AA | AB | AB | AB | AA |
| AB | AB | AB | AA | AB | AB | AB | AB | AA | AA | AB | AA | AB | AB | AB | AB | AB | AA | AA | AB | AB | AB | AA | AB | AB | AB | AB | AA |
| AA | AA | AA | AA | AA | AA | AA | AA | AA | AA | AA | AA | AA | AA | AB | AA | AB | AA | AA | AA | AB | AA | AA | AA | AA | AA | AA | AB |
| AB | AB | AA | AB | AB | AB | AB | AB | AB | AA | AA | AB | AB | AB | AB | AB | AB | AA | AB | AB | AA | AB | AA | AB | AB | AB | AB | AB |
| AB | AA | AB | AB | AB | AB | AB | AB | AB | AA | AB | AA | AB | AB | AB | AB | AB | AA | AB | AB | AB | AA | AB | AB | AB | AB | AB | AB |
| AA | AA | AB | AA | AB | AB | AB | AB | AB | AA | AA | AB | AA | AB | AB | AB | AB | AA | AB | AB | AA | AB | AB | AA | AB | AB | AB | AA |
| - | AB | AB | AB | AB | AA | AB | AA | AB | - | AB | - | AB | AB | AB | AB | AB | AB | AB | AB | AB | AB | AB | AB | AB | AB | - | AB |
| AB | AB | AB | AB | AB | AA | AB | AB | AB | AB | AA | AA | - | AB | AB | AB | AA | AB | AB | AB | AB | AB | - | AA | AB | AB | - | AA |
| AA | AA | AA | AA | AB | AA | AA | AB | AB | AA | AB | AA | AA | AB | AB | AA | AA | AA | AA | AB | AA | AA | - | AB | AB | AB | AB | AB |
| AB | - | AB | AB | AB | - | AB | AB | AB | - | AB | AB | AB | AB | AB | AB | AB | AB | AB | - | AB | AB | - | AB | AB | AB | - | AB |
| AB | AA | AA | AA | AA | AB | AA | AB | AA | - | AB | AA | AA | AA | AB | AA | AB | AB | AA | AB | AA | AA | AB | AA | AB | AA | - | AB |
| AA | AA | AB | AB | AB | AB | AB | AB | AB | AB | AA | AB | AA | AB | AB | AB | AB | AB | AB | AB | AA | AA | AA | AA | AB | AA | AA | AB |
| AB | AB | AB | AB | AB | AB | AB | AB | AB | AB | AB | AB | AB | AB | AB | AB | AB | AB | AB | AB | AB | AB | AA | AB | AB | AB | AA | AB |
| AA | AA | AA | AA | AA | AA | AA | AA | AA | AA | AA | AA | AA | AA | AA | AA | AA | AA | AA | AA | AA | AA | AA | AA | AA | AA | AA | AA |
| - | AA | AA | AA | AA | - | AA | AA | AA | AA | AA | AA | AA | AA | AA | - | AA | AA | AA | - | AA | AA | AA | AA | AA | AA | AA | AA |
| AB | AA | AB | AA | AA | AA | AA | AA | AA | AA | AA | AA | AA | AA | AB | AA | AA | AA | AB | AA | AA | AA | AA | AB | AA | AA | AA | AA |
| AB | AA | AA | AA | AA | AB | AA | AA | AB | AA | AA | AA | AA | AB | AB | AB | AA | AA | AB | AA | AB | AA | AA | AB | AA | AB | AA | AA |
| AB | AB | AB | AB | AB | AB | AB | AB | AB | AB | AB | AB | AB | AB | AB | AB | AB | AB | AB | AB | AB | AB | AB | AB | AB | AB | AB | AB |
| AB | AB | AB | AB | AB | AB | AB | AA | AB | AB | AB | AA | AA | AB | AA | AB | AB | AB | AB | AB | AB | AB | AB | AB | AB | AB | AA | AB |
| AA | AA | AA | AA | AA | AB | AA | AA | AA | - | AA | AA | AA | - | AA | AA | AA | AA | AA | AA | AA | AA | AA | AA | AA | AA | - | AA |
| AA | AA | AA | AA | AA | AB | AA | AA | AA | AA | AA | AB | AB | AA | AA | AB | AA | AA | AB | AB | AB | AA | AB | AA | AA | AA | - | AA |
| AA | AA | AA | AA | AA | AB | AB | AB | AB | AB | AB | AA | AB | AA | AA | AB | AA | AB | AB | AB | AB | AA | AB | AB | AA | AA | AB | AB |
| AB | AB | AB | AA | AA | AB | AB | AB | AB | AB | AB | AA | AB | AB | AB | AA | AB | AB | AB | AA | AB | AA | AB | AB | AB | AB | AB | AB |
| AA | AB | AB | AB | AB | AB | AB | AA | AB | AA | AA | AB | AB | AB | AA | AB | AB | AA | AB | AB | AB | AA | - | AB | AA | AB | AB | AA |
| AA | AB | AB | AB | AB | AA | AB | AA | AB | AB | AB | AB | AB | AB | AB | AB | AB | AB | AB | AB | AA | AB | AB | AB | AA | AB | AB | AA |
| AB | AA | AA | AA | AB | - | AA | AA | AB | AB | AA | AB | AB | AB | AA | AB | AA | AB | AB | AB | AA | AB | AB | AA | AB | AB | AA | AB |
| AB | AB | AB | AB | - | - | AB | AA | AB | AB | AA | AA | AB | AB | AB | AB | AB | AA | AB | AB | AB | AB | - | AB | AA | AB | - | AB |
| AA | - | AA | - | - | - | AA | AB | AA | - | - | AA | AA | AB | AB | AA | AA | AA | AB | AA | AA | AB | - | AB | AB | AA | - | AA |
| AA | AB | AA | AB | AA | AB | AA | AA | AB | AA | AA | AB | AA | AA | AB | AB | AA | AA | AB | AB | AA | AA | AA | AB | AB | AB | AB | AB |
| AB | AA | AA | AB | AA | AB | AB | AB | AB | AB | AB | AB | AA | AB | AB | AB | AA | AB | AA | AB | AB | AB | AB | AB | AA | AA | AB | AB |
| AB | AB | AA | AB | AA | AB | AA | AB | AB | AB | AB | AA | AA | AB | AA | AB | AB | AA | AB | AA | AB | AB | AB | AB | AB | AA | AA | AA |
| AA | AB | AA | AA | AB | AA | AA | AA | AA | AA | AA | AA | AA | AA | AA | AB | AB | AA | AB | AB | AB | AA | AB | AB | AA | AA | AA | AA |
| AB | AB | AB | AA | AB | AB | AB | AB | AA | AB | AB | AB | AB | AA | AB | AB | AB | AA | AA | AA | AA | AA | - | AA | AA | AB | AB | AB |
| AB | AB | AB | AB | AB | AB | AB | AA | AB | AB | AB | AB | AB | AB | AB | AB | AA | AA | AB | AB | AB | AB | AA | AB | AA | AB | AB | AB |
| AA | AB | AB | AB | AB | AB | AA | AA | AB | AA | AB | AA | AB | AA | AA | AB | AA | AB | AB | AA | AB | AA | AB | AA | AA | AB | AA | AB |
| AB | AA | AA | AA | AB | - | AB | AB | AA | AA | AB | AA | AA | AB | AA | AB | AB | AB | AB | AA | AB | AA | AB | AA | AB | AA | AA | AB |
| AB | AA | AB | AA | AB | AA | AA | AA | AA | AA | AA | AB | AA | AB | AB | AB | AB | AB | AA | AB | AB | AB | AA | AA | AA | AB | AB | AA |
| AA | AA | AB | AA | AA | AB | AA | AA | AA | AA | AA | AA | AA | AA | AB | AA | AB | AA | AB | AA | AB | AB | AB | AA | AA | AB | AB | AB |
| AB | AA | AA | AA | AB | AB | AA | AA | AA | AA | AA | AB | AB | AA | AA | AB | AA | AA | AB | AA | AB | AB | AB | AB | AA | AA | AB | AB |
| - | - | AB | AB | AB | - | AB | AB | AB | - | AA | AB | AB | AB | AB | AB | AB | AB | AB | AB | AA | AB | - | AA | AB | AB | - | AA |
| AA | AA | AB | AA | AB | AA | AA | AB | AB | AA | AA | AB | AA | AB | AB | AA | AB | AA | AB | AB | AB | AA | AB | AB | AA | AB | AB | AB |
| AA | AB | AA | AB | AA | AB | AB | AA | AB | AA | AA | AB | AA | AB | AB | AB | AB | AA | AB | AB | AB | AA | AB | AB | AA | AB | AA | AB |
| AA | AB | AB | AB | AB | AB | AB | AB | AB | AB | AB | AB | AB | AB | AB | AB | AB | AB | AB | AB | AB | AB | AB | AB | AB | AB | AB | AB |
| AA | AA | AA | AA | AA | AB | AA | AA | AA | AA | AB | AA | AA | AA | AA | AA | AA | AA | AA | AA | AA | AB | AA | AB | AA | AA | AA | AB |
| AB | AB | AB | AB | AB | - | AB | AB | AA | AB | AB | AB | AB | AB | AB | AA | AA | AA | AA | AB | AA | AB | - | AA | AB | AB | AB | AA |
| AB | AB | AB | AB | AB | AB | AB | AB | AB | AB | AB | AB | AB | AB | AB | AB | AB | AB | AB | AB | AB | AA | AB | AB | AB | AA | AB | AB |
| AA | AB | AA | AB | AB | AB | AB | AB | AB | AB | AB | AB | AA | AA | AB | AA | AA | AB | AB | AA | AB | AA | AB | AB | AB | AB | AB | AB |
| AA | AA | AA | AA | AA | AB | - | AA | AA | - | AB | AA | AA | - | AA | AB | AB | AA | AB | AB | AA | AB | AA | - | AA | AA | - | AB |
| AB | AA | AB | AA | AB | AA | AB | AA | AA | AA | AA | AA | AA | AB | AA | AA | AA | AA | AA | AB | AB | AB | AA | AA | AB | AA | AA | AB |
| AB | AB | AB | AB | AB | AB | AB | AB | AB | AB | AB | AB | AB | AB | AB | AB | AB | AB | AB | AB | AB | AB | AB | AB | AB | AB | AB | AB |

| snp_B1556 | snp_B1594 | snp_B1608 | snp_B1618 | snp_B1656 | snp_B1657 | snp_B1658 | snp_B1676 | snp_B1702 | snp_B1721 | snp_B1745 | snp_B1761 | snp_B1768 | snp_B1776 | snp_B1779 | snp_B1808 | snp_B1812 | snp_B1820 | snp_B1845 | snp_B1864 | snp_B1889 | snp_B1892 | snp_B1901 | snp_B1920 | snp_B1928 | snp_B1929 | snp_B1944 | snp_B1961 | snp_B1979 | snp_B2015 | snp_B2043 | snp_B2044 | snp_B2047 | snp_B2067 | snp_B2083 | snp_B2089 | snp_B2106 | snp_B2120 | snp_B2148 | snp_B2154 | snp_B2171 | snp_B2195 | snp_B2217 | snp_B2235 | snp_B2259 | snp_B2270 | snp_B2277 | snp_B2281 | snp_B2285 | snp_B2290 | snp_B2294 | snp_B2328 | snp_B2329 | snp_B2342 | snp_B2351 | snp_B2364 | snp_B2368 | snp_B2382 | snp_B2394 | snp_B2414 | snp_B2430 | snp_B2443 | snp_B2463 | snp_B2466 | snp_B2475 | snp_B2524 | snp_B2534 | snp_B2535 | snp_B2552 | snp_B2575 | snp_B2590 | snp_B2604 | snp_B2615 | snp_B2619 | snp_B2631 | snp_B2636 | snp_B2667 |
|---|---|---|---|---|---|---|---|---|---|---|---|---|---|---|---|---|---|---|---|---|---|---|---|---|---|---|---|---|---|---|---|---|---|---|---|---|---|---|---|---|---|---|---|---|---|---|---|---|---|---|---|---|---|---|---|---|---|---|---|---|---|---|---|---|---|---|---|---|---|---|---|---|---|---|---|---|
| 5 | 5 | 5 | 5 | 5 | 5 | 5 | 5 | 5 | 5 | 5 | 5 | 5 | 5 | 5 | 5 | 5 | 5 | 5 | 5 | 5 | 5 | 5 | 5 | 5 | 5 | 5 | 5 | 5 | 5 | 5 | 5 | 5 | 5 | 5 | 5 | 5 | 5 | 5 | 5 | 5 | 5 | 5 | 5 | 5 | 5 | 5 | 5 | 5 | 5 | 5 | 5 | 5 | 5 | 5 | 5 | 5 | 5 | 5 | 5 | 5 | 5 | 5 | 5 | 5 | 5 | 5 | 5 | 5 | 5 | 5 | 5 | 5 | 5 | 5 | 5 | 5 |
| -- | -- | -- | -- | -- | -- | -- | -- | -- | -- | -- | -- | -- | -- | -- | -- | -- | -- | -- | -- | -- | -- | -- | -- | -- | -- | -- | -- | -- | -- | -- | -- | -- | -- | -- | -- | -- | -- | -- | -- | -- | -- | -- | -- | -- | -- | -- | -- | -- | -- | -- | LG13 | -- | -- | -- | -- | -- | -- | -- | -- | -- | -- | -- | -- | -- | -- | -- | -- | -- | -- | -- | -- | -- | -- | -- | -- | -- |

| snp_B2693 5 | snp_B2703 5 | snp_B2747 5 | snp_B2761 5 | snp_B2778 5 | snp_B2788 5 | snp_B2791 5 | snp_B2813 5 | snp_B2827 5 | snp_B2831 5 | snp_B2837 5 | snp_B2842 5 | snp_B2850 5 | snp_B2869 5 | snp_B2880 5 | snp_B2881 5 | snp_B2897 5 | snp_B2898 5 | snp_B2921 5 | snp_B2924 5 | snp_B2930 5 | snp_B2931 5 | snp_B2948 5 | snp_B2962 5 | snp_B2972 5 | snp_B3001 5 | snp_B3023 5 | snp_B3029 5 | snp_B3067 5 | snp_B3083 5 | snp_B3094 5 | snp_B3099 5 | snp_B3114 5 | snp_B3187 5 | snp_B3197 5 | snp_B3212 5 | snp_B3220 5 | snp_B3240 5 | snp_B3251 5 | snp_B3254 5 | snp_B3271 5 | snp_B3273 5 | snp_B3283 5 | snp_B3293 5 | snp_B3298 5 | snp_B3307 5 | snp_B3308 5 | snp_B3335 5 | snp_B3344 5 | snp_B3346 5 | snp_B3361 5 | snp_B3363 5 | snp_B3368 5 | snp_B3379 5 | snp_B3382 5 | snp_B3383 5 | snp_B3395 5 | snp_B3406 5 | snp_B3409 5 | snp_B3433 5 | snp_B3453 5 | snp_B3495 5 | snp_B3498 5 | snp_B3510 5 | snp_B3530 5 | snp_B3534 5 | snp_B3556 5 | snp_B3559 5 | snp_B3561 5 | snp_B3569 5 | snp_B3578 5 | snp_B3585 5 | snp_B3615 5 | snp_B3635 5 | snp_B3636 5 | snp_B3653 5 | snp_B3655 5 |
|---|---|---|---|---|---|---|---|---|---|---|---|---|---|---|---|---|---|---|---|---|---|---|---|---|---|---|---|---|---|---|---|---|---|---|---|---|---|---|---|---|---|---|---|---|---|---|---|---|---|---|---|---|---|---|---|---|---|---|---|---|---|---|---|---|---|---|---|---|---|---|---|---|---|---|---|---|
| -- | -- | -- | -- | -- | -- | -- | -- | -- | LG13 | -- | -- | -- | -- | -- | -- | -- | -- | -- | -- | -- | -- | -- | -- | -- | -- | -- | -- | -- | -- | -- | -- | -- | -- | -- | -- | -- | -- | -- | -- | -- | -- | -- | -- | -- | -- | -- | -- | -- | -- | -- | -- | -- | -- | -- | -- | -- | -- | -- | -- | -- | -- | -- | -- | -- | -- | -- | -- | -- | -- | -- | -- | -- | -- | -- | -- | -- |
| AB | AA | AB | AA | - | AA | AA | AA | AB | AB | AB | AA | AB | AA | AA | AA | AA | AB | AA | AA | AB | AB | AA | AB | AB | AB | AB | AB | AB | AB | AA | AA | AB | AA | AB | AA | AB | AB | AA | AA | AB | AB | AB | AB | AA | AB | AA | AA | AA | AA | AB | AA | AB | AA | AA | AB | AB | AB | AA | AA | AB | AA | AB | AB | AA | AA | AA | AB | AA | AA | AA | AB | AB | AA | AB | AB | AA |
| AA | AA | AA | AB | AB | AB | AB | AB | AB | AB | AA | AA | AB | AB | AA | AB | AA | AA | AB | AA | AB | AA | AA | AA | AB | AB | AB | AB | AB | AA | AA | AB | AA | AB | AB | AA | AA | AA | AA | AA | AA | AA | AA | AA | AB | AA | AA | AA | AA | AA | AA | AB | AB | AB | AA | AB | AB | AA | AB | AA | AB | AB | AB | AA | AA | AB | AA | AB | AB | AB | AB | AB | AB | AA | AA | AA | AB |

| snp_B3693 5 | snp_B3853 5 | snp_B3871 5 | snp_B3893 5 | snp_B3905 5 | snp_B3934 5 | snp_B3935 5 | snp_B3941 5 | snp_B3944 5 | snp_B3973 5 | snp_B3981 5 | snp_B4000 5 | snp_B4001 5 | snp_B4035 5 | snp_B4056 5 | snp_B4062 5 | snp_B4076 5 | snp_B4079 5 |
|---|---|---|---|---|---|---|---|---|---|---|---|---|---|---|---|---|---|
| -- | -- | -- | -- | -- | -- | -- | -- | -- | -- | -- | -- | -- | -- | -- | -- | -- | -- |
| AB | AA | AB | AA | AB | AB | AA | AB | AA | AB | AA | AB | AA | AB | AB | AA | AA | AA |
| AA | AA | AB | AA | AB | AB | AA | AB | AA | AB | AB | AB | AA | AB | AB | AA | AA | AA |
| AA | AB | AA | AA | AA | AB | AA | AB | AA | AA | AB | AA | AA | AB | AA | AA | AB | AA |
| AA | AB | AA | AA | AA | AB | AB | AB | AB | AB | AA | AA | AB | AB | AB | AB | AA | AB |
| AA | AB | AA | AA | AA | AB | AB | AB | AA | AB | AB | AA | AB | AB | AB | AB | AA | AA |
| AB | AB | AA | AB | AA | - | - | AA | - | AA | AA | AA | AB | AB | AB | AB | AA | AB |
| AB | AB | AB | AB | AA | AB | AA | AB | AB | AB | AB | AA | AB | AB | AB | AB | AA | AB |
| AA | AB | AA | AA | AA | AA | AB | AA | AA | AA | AB | AA | AA | AB | AB | AA | AA | AA |
| AB | AA | AB | AA | AA | AA | AB | AA | AB | AA | AB | AA | AA | AB | AB | AA | AB | AA |
| AB | AA | AA | AA | AB | AA | AB | AA | AB | AA | AA | AA | AA | AB | AB | AA | AA | AB |
| AA | AB | AA | AA | AA | AB | AA | AB | AA | AB | AB | AA | AA | AB | AA | AB | AA | AB |
| AA | AB | AB | AB | AB | AB | AB | AA | AB | AB | AA | AA | AB | AA | AB | AA | AB | AB |
| AA | AA | AB | AA | AB | AA | AA | AA | AA | AA | AB | AB | AA | AA | AA | AB | AB | AA |
| AA | AA | AA | AA | AA | AB | AA | AB | AA | AA | AA | AA | AA | AB | AB | AB | AA | AA |
| AB | AA | AB | AB | AB | AB | AA | AA | AB | AB | AA | AB | AA | AA | AB | AB | AA | AA |
| AA | AA | AB | AA | AA | AB | AB | AB | AA | AA | AB | AA | AA | AB | AA | AB | AB | AB |
| AA | AA | AA | AA | AA | AA | AA | AA | AA | AA | AA | AA | AA | AA | AA | AA | AA | AA |
| AB | AB | AB | - | - | AA | - | AA | AA | AB | AA | AA | - | AA | AB | AB | AB | - |
| AB | AB | AB | AB | AB | AB | AB | AA | AB | AB | AB | AB | AB | AB | AB | AB | AB | AB |
| AB | AB | AA | AB | AB | AA | AB | - | AB | AA | AA | AB | AB | AA | AB | AB | AB | AB |
| AB | AB | AB | AB | AB | AB | AB | AB | AB | AB | AB | AB | AB | AB | AB | AA | AB | AB |
| AB | AB | AB | AB | AB | AB | AA | AB | AB | AB | AB | AB | AB | AB | AB | AA | AB | AB |
| AA | - | AA | - | AA | AA | - | - | AB | AA | AB | AA | - | AB | AB | AB | AA | AB |
| AB | AA | AB | AB | AB | AA | AA | AA | AA | AB | AA | AB | AA | AA | AB | AA | AA | AB |
| AA | AA | AA | AA | AA | AA | AB | AA | AA | AA | AA | AB | AA | AA | AA | AA | AA | AA |
| AB | AB | AB | AB | AB | AB | AB | AB | AB | AB | AB | AB | AB | AB | AB | AB | AB | AB |
| AA | AB | AA | AA | AA | AB | AA | AA | AA | AA | AA | AA | AA | AA | AA | AA | AA | AA |
| AB | AB | AB | AB | AB | AA | AA | AA | AB | AA | AA | AB | AB | AA | AA | AA | AA | AA |
| AA | AA | AA | AA | AA | AA | - | AB | AA | AA | AB | AA | AA | AA | AB | AB | AA | AA |
| - | - | AA | - | - | AA | - | AA | - | AA | AA | AB | AA | AA | AA | AB | AA | AA |
| AB | AB | AB | AB | AB | AB | AB | AB | AB | AB | AB | AB | AB | AB | AB | AB | AB | AB |
| AA | AB | AB | AA | AA | AB | AA | AB | AA | AB | AB | AA | AA | AB | AB | AB | AB | AB |
| AA | AB | AA | - | AA | AB | - | AB | AB | AA | AB | AA | AB | AB | AB | AB | AB | AA |
| AB | AA | AB | AB | AB | AA | - | - | AB | AB | AB | AA | AA | AA | AB | AA | AA | AB |
| - | - | AB | - | - | AB | AA | AB | AA | AA | AB | AA | AA | AB | AA | AB | AA | AB |
| AB | AB | AB | AB | AB | AB | - | AB | AB | AB | AB | AB | AB | AB | AB | AB | AB | AB |
| AA | AB | AA | AB | AA | AB | AA | AA | AB | AA | AB | AA | AB | AB | AB | AA | AA | AB |
| AB | AB | AB | AB | AB | AB | AB | AB | AB | AB | AB | AB | AB | AB | AB | AB | AB | AB |
| AB | AB | AB | AB | AB | AB | AA | AB | AB | AB | AB | AB | AB | AB | AB | AB | AB | AB |
| AA | AB | AA | - | AA | - | AB | AB | AA | AB | AB | AB | AB | AB | AB | AB | AA | AA |
| AB | AA | AB | AB | AB | AB | AB | AB | AB | AB | AB | AB | AB | AB | AB | AB | AB | AB |
| AA | AA | AA | AA | AA | AA | AB | AA | AA | AA | AA | AA | AA | AA | AA | AA | AA | AA |
| AB | AB | AA | AB | AB | AA | AA | AA | AB | AA | AB | AA | AB | AA | AB | AA | AB | AB |
| AA | AA | AB | AA | AB | AA | AA | AB | AA | AA | AB | AB | AA | AB | AB | AB | AA | AB |
| AA | AA | AB | AA | AB | AA | AA | AB | AB | AB | AB | AB | AA | AB | AB | AB | AB | AA |
| AB | AB | AB | AB | AB | AA | AB | AB | AB | AB | AB | AB | AB | AB | AB | AB | AB | AB |
| AA | AA | AA | AA | AA | AA | AB | AA | AA | AA | AA | AA | AA | AA | AA | AA | AA | AA |
| AA | AA | AA | AA | AA | AA | AA | AA | AA | AA | AA | AA | AA | AA | AA | AA | AA | AA |
| AA | AA | AB | - | AA | AB | AB | AB | AA | AB | AB | AA | AA | AB | AA | AA | AA | AA |
| AA | AA | AA | AA | AA | AA | AA | AA | AA | AA | AA | AA | AA | AA | AA | AA | AA | AA |

| snp_B4084 5 | snp_B4098 5 | snp_B4112 5 | snp_B4118 5 | snp_B4151 5 | snp_B4153 5 | snp_B4180 5 | snp_B4187 5 | snp_B4224 5 | snp_B4240 5 | snp_B4282 5 | snp_B4284 5 | snp_B4336 5 | snp_B4369 5 | snp_B4393 5 | snp_B4396 5 | snp_B4411 5 | snp_B4418 5 | snp_B4437 5 | snp_B4440 5 | snp_B4446 5 | snp_B4471 5 | snp_B4477 5 | snp_B4481 5 | snp_B4482 5 | snp_B4506 5 |
|---|---|---|---|---|---|---|---|---|---|---|---|---|---|---|---|---|---|---|---|---|---|---|---|---|---|
| -- | -- | -- | -- | -- | -- | -- | -- | -- | -- | -- | -- | -- | -- | -- | -- | -- | -- | -- | -- | -- | -- | LG13 | -- | -- | -- |
| AB | AA | AA | AB | AB | AB | AA | AA | AB | AA | AB | AA | AB | AB | AB | AB | AB | AA | AB | AB | AB | AA | AB | AA | AA | AA |
| AB | AA | AA | AA | AA | AA | AB | AA | AA | AB | AA | AA | AB | AB | AA | AB | AA | AA | AA | AB | AB | AA | AA | AA | AA | AB |
| AB | AA | AA | AB | AB | AA | AB | AB | AA | AB | AB | AA | AA | AB | AB | AB | AA | AA | AB | AA | AA | AB | AB | AB | AB | AB |
| AB | AB | AA | AB | AB | AB | AA | AA | AA | AB | AB | AA | - | AA | AA | AB | AB | AA | AB | AA | AB | AA | AA | AA | - | AA |
| AB | AA | AA | AB | AA | AB | AB | AA | AB | AA | AB | AB | AA | AA | AB | AA | AB | AA | AA | AA | AA | AA | AA | AA | AA | AA |
| - | - | - | AA | AB | - | - | AB | - | - | AB | - | - | - | AB | - | AB | AB | AB | - | AA | AB | AA | AA | - | AA |
| AB | AB | AB | AA | AB | AB | AA | AB | AB | AA | AB | AB | AA | AA | AA | AB | AB | AB | AB | AA | AA | AB | AA | AB | AB | AB |
| AB | AA | AA | AA | AB | AA | AA | AB | AA | AA | AB | AA | AB | AA | AA | AB | AA | AA | AA | AA | AB | AA | AA | AB | AA | AA |
| AB | AB | AA | AB | AB | AB | AA | AB | AA | AB | AA | AA | AA | AA | AB | AA | AB | AA | AB | AA | AA | AB | AA | AB | AB | AB |
| AB | AA | AA | AA | AA | AB | AB | AA | AA | AA | AB | AB | AA | AB | AA | AB | AB | AA | AB | AB | AA | AB | AB | AA | AB | AA |
| AB | AA | AA | AB | AA | AB | AB | AA | AA | AB | AB | AB | AA | AB | AB | AB | AB | AA | AB | AA | AA | AB | AB | AB | AA | AB |
| AA | AB | AA | AB | AB | AB | AB | AB | AA | AA | AB | AA | AB | AB | AA | AB | AB | AA | AB | AB | AA | AB | AB | AB | AA | AA |
| AB | AA | AB | AA | AA | AA | AB | AA | AA | AB | AA | AA | AB | AB | AA | AB | AA | AA | AA | AB | AA | AA | AB | AA | AA | AB |
| AB | AA | AB | AA | AA | AA | AB | AA | AA | AB | AA | AA | AB | AB | AA | AB | AB | AA | AA | AB | AA | AA | AA | AB | AB | AA |
| AA | AA | AA | AB | AB | AB | AA | AB | AA | AA | AB | AB | AA | AA | AB | AA | AB | AB | AA | AB | AB | AB | AB | AA | AA | AA |
| AA | AB | AA | AA | AB | AA | AB | AB | AA | AB | AB | AB | AA | AA | AA | AB | AA | AA | AB | AA | AB | AA | AA | AB | AB | AB |
| AA | AA | AA | AA | AA | AA | AA | AA | AA | AA | AA | AA | AA | AA | AA | AA | AA | AA | AA | AA | AA | AA | AB | AA | AB | AA |
| AA | - | AB | AA | AA | - | - | - | - | - | AA | - | AA | - | - | - | AB | AB | - | - | AB | AB | - | AB | AA | AA |
| AB | AB | AB | AA | AB | AB | AB | AB | AB | AB | AB | AB | AB | AB | AA | AB | AB | AB | AB | AB | AB | AB | AB | AB | AB | AB |
| AA | AB | AB | AA | AB | AB | AB | AB | AB | AB | AB | AB | AA | AB | AA | AA | AB | AB | AB | AB | AB | AB | AB | AB | AA | AB |
| AB | AB | AB | AA | AB | AB | AB | AB | AB | AB | AB | AB | AB | AB | AA | AB | AB | AB | AB | AB | AB | AB | AB | AB | AB | AB |
| AB | AB | AA | AB | AB | AB | AB | AB | AB | AB | AB | AB | AB | AB | AA | AB | AB | AB | AB | AB | AB | AB | AB | AB | AB | AB |
| AB | AA | - | AB | AB | - | - | - | - | AB | AA | - | AA | AB | - | - | - | AB | - | AA | AA | AA | - | AB | - | AB |
| AB | AA | AA | AB | AB | AA | AB | AA | AA | AA | AA | AA | AB | AB | AA | AB | AA | AA | AA | AB | AB | AB | AB | AB | AB | AA |
| AA | AA | AB | AB | AA | AA | AA | AA | AA | AA | AA | AA | AA | AA | AB | AA | AA | AB | AA | AA | AA | AA | AB | AA | AA | AA |
| AB | AB | AB | AB | AB | AB | AB | AB | AB | AB | AB | AB | AB | AB | AB | AB | AB | AB | AB | AB | AB | AB | AB | AB | AB | AB |
| AA | AA | AA | AA | AA | AA | AA | AA | AB | AA | AB | AB | AA | AA | AA | AA | AA | AB | AA | AA | AA | AA | AA | AA | AA | AA |
| AA | AA | AA | AA | AA | AA | AB | AB | AB | AA | AB | AA | AB | AB | AA | AA | AB | AA | AA | AB | AB | AB | AB | AB | AB | AB |
| AB | AB | - | AA | AA | AA | AB | AA | AB | AA | AB | AB | - | AA | AA | AB | AA | AB | AA | AA | AB | AB | AB | AA | AA | AA |
| AA | AA | - | AA | AA | AA | - | - | - | AA | AA | - | AB | - | AA | - | AA | AA | - | - | AB | AA | AA | AA | - | AA |
| AB | AB | AB | AB | AB | AB | AB | AB | AB | AB | AB | AB | AB | AB | AB | AB | AB | AB | AB | AB | AB | AB | AB | AB | AB | AB |
| AB | AB | AB | AA | AA | AB | AB | AA | AA | AB | AB | AA | AA | AA | AB | AA | AB | AA | AB | AA | AA | AA | AA | AB | AB | AB |
| AA | - | AB | AA | AA | - | AA | AB | - | AB | AB | AA | AA | AA | AB | AA | AB | AB | AB | AA | AA | AA | - | AA | - | AB |
| AB | AA | AB | AA | AA | AB | AB | AB | AB | AA | AA | AB | AA | AB | AA | - | AA | AA | AA | AA | AB | AB | AB | AA | AB | AB |
| - | - | AA | AB | AB | - | AB | AB | - | AB | AB | - | AA | - | AB | AA | AA | AA | AA | AA | AA | AB | AA | AB | AB | AB |
| AB | AB | - | AB | AB | - | AB | AB | AB | AB | AB | AB | - | AB | AB | AB | AB | AB | AB | AB | AB | AB | AB | AB | - | AB |
| AA | AB | AA | AA | AA | AA | AA | AA | AB | AA | AB | AB | AA | AA | AA | AA | AA | AB | AA | AA | AA | AA | AB | AA | AB | AA |
| AB | AB | AB | AB | AB | AB | AB | AB | AB | AB | AB | AB | AB | AB | AB | AB | AB | AB | AB | AB | AB | AB | AB | AB | AB | AB |
| AB | AB | AB | AB | AB | AB | AB | AB | AB | AB | AB | AB | AB | AB | AB | AB | AB | AB | AB | AB | AB | AB | AB | AB | AB | AB |
| AB | - | - | AB | AA | AB | AB | AA | - | AA | AB | AB | AB | AA | AB | - | AB | AA | AA | AB | AA | AA | AA | AA | AA | AA |
| AB | AB | AA | AA | AB | AB | AB | AB | AB | AB | AB | AB | AA | AB | AA | AB | AB | AB | AB | AB | AA | AA | AA | AB | AB | AB |
| AA | AA | AA | AA | AB | AA | AB | AA | AA | AA | AA | AA | AB | AA | AA | AA | AA | AA | AA | AA | AA | AA | AA | AA | AA | AA |
| AA | AA | AA | AB | AA | AA | AB | AB | AB | AB | AA | AA | AA | AB | AA | AA | AB | AB | AA | AA | AA | AB | AB | AB | AA | AA |
| AB | AA | AB | AB | AA | AA | AB | AA | AA | AB | AA | AA | AB | AB | AB | AB | AA | AA | AA | - | AB | AA | AB | AB | - | AB |
| AB | AA | AA | AB | AA | AA | AB | AA | AA | AB | AA | AA | AB | AB | AA | AB | AA | AA | AB | AA | AA | AB | AB | AB | AB | AB |
| AB | AB | AB | AB | AB | AB | AB | AB | AB | AB | AA | AB | AB | AB | AB | AB | AB | AB | AB | AB | AB | AB | AA | AB | AB | AB |
| AA | AA | AA | AB | AA | AA | AA | AA | AA | AA | AA | AA | AB | AA | AB | AA | AA | AA | AA | AA | AA | AA | AA | AA | AA | AA |
| AB | AB | AA | AB | AA | AB | AA | AA | AA | AB | AB | AB | AA | AA | AB | AB | AA | AA | AB | AA | AA | AA | AA | AA | AB | AB |
| AB | AA | AB | AB | AA | AB | AA | AA | AA | AA | AA | AA | AB | AA | AB | AB | AA | AA | AA | AA | AA | AA | AB | AA | AA | AB |
| AB | AB | AA | AB | AA | AB | AA | AB | AA | AA | AA | AB | AA | AA | AB | AB | AA | AB | AA | AA | AB | AB | AA | AB | AA | AB |

| snp_B4523 5 | snp_B4525 5 | snp_B4536 5 | snp_B4560 5 | snp_B4589 5 | snp_B4607 5 | snp_B4623 5 | snp_B4644 5 | snp_B4662 5 | snp_B4664 5 | snp_B4686 5 | snp_B4693 5 | snp_B4711 5 | snp_B4786 5 |
|---|---|---|---|---|---|---|---|---|---|---|---|---|---|
| -- | -- | -- | -- | -- | -- | -- | -- | -- | -- | -- | -- | -- | -- |
| AA | AA | AB | AB | AA | AA | AB | AB | AB | AB | AA | AA | AA | AB |
| AA | AA | AA | AA | AA | AA | AB | AB | AB | AB | AB | AB | AB | AB |
| AA | AA | AA | AB | AA | AB | AA | AA | AB | AA | AB | AA | AA | AB |
| - | AA | AB | AA | AB | AA | - | AB | AB | AB | AA | AB | - | AA |
| AA | AA | AB | AB | AA | AA | AB | AA | AA | AB | AB | AB | AB | AB |
| - | AA | AA | AB | - | AA | - | AA | AB | AB | AA | AA | - | AA |
| AB | AA | AB | AA | AB | AB | AA | AA | AB | AA | AB | AB | AA | AB |
| AA | AA | AA | AA | AA | AA | AA | AB | AA | AB | AA | AB | AB | AA |
| AB | AB | AB | AB | AB | AA | AB | AB | AB | AB | AA | AB | AB | AB |
| AA | AB | AA | AB | AA | AA | AB | AB | AB | AA | AB | AA | AB | AA |
| AB | AA | AB | AA | AB | AB | AA | AB | AB | AA | AA | AB | AA | AB |
| AB | AB | AA | AB | AB | AB | AB | AA | AB | AA | AB | AA | AB | AB |
| AA | AA | AA | AA | AA | AB | AB | AB | AA | AB | AB | AB | AA | AB |
| AA | AB | AA | AB | AA | AB | AA | AA | AB | AA | AB | AA | AA | AB |
| AA | AB | AA | AB | AA | AA | AA | AB | AA | AB | AA | AB | AB | AA |
| AB | AA | AA | AB | AB | AA | AA | AA | AB | AA | AB | AB | AB | AA |
| AA | AA | AA | AA | AA | AA | AA | AA | AA | AA | AA | AA | AA | AA |
| - | AB | AA | AB | AA | AB | AA | AB | AB | - | - | AB | - | AA |
| AB | AA | AB | AA | AB | AB | AB | AB | AB | AB | AB | AB | AA | AB |
| - | AB | AA | AB | AA | AA | - | AB | AA | AB | AB | AA | AB | AB |
| AB | AB | AB | AB | AB | AB | AB | AB | AB | AB | AB | AB | AB | AB |
| AB | AB | AB | AA | AB | AB | AB | AB | AB | AA | AB | AB | AB | AA |
| AB | AB | - | - | - | AB | - | AA | - | - | AB | AB | AB | AB |
| AA | AB | AA | AA | AA | AB | AB | AB | AB | AA | AB | AA | AA | AA |
| AA | AA | AA | AB | AA | AA | AA | AA | AB | AB | AA | AA | AB | AA |
| AB | AB | AB | AB | AB | AB | AB | AB | AB | AB | AB | AB | AB | AB |
| AA | AA | AA | AA | AA | AA | AA | AA | AA | AA | AA | AA | AA | AA |
| AA | AA | AA | AB | AA | AA | AB | AA | AB | AB | AA | AB | AA | AB |
| AB | AA | AA | AA | AB | AB | AA | AB | AB | AA | AB | AB | AA | AA |
| AA | - | AA | AA | - | AA | - | - | - | - | - | AA | AB | AB |
| AB | AB | AB | AB | AB | AB | AB | AB | AB | AB | AB | AB | AB | AB |
| AB | AA | AB | AB | AB | AB | AA | AA | AB | AA | AB | AB | AA | AB |
| - | AA | AB | AB | AB | AA | AA | AA | AA | - | AA | AB | AB | AB |
| - | AB | AA | AA | - | AA | - | AB | AA | - | AB | AA | AB | AA |
| AB | AA | AA | AA | AB | AA | - | AA | AB | - | - | AB | - | AA |
| AB | AB | AB | AB | - | AB | AB | AB | AB | AB | AB | AB | - | AB |
| AB | AA | AA | AA | AB | AA | AA | AA | AA | AA | AA | AA | AA | AA |
| AB | AB | AB | AB | AB | AB | AB | AB | AB | AB | AB | AB | AB | AB |
| AB | AB | AB | AA | AB | AB | AB | AB | AB | AB | AB | AB | AB | AB |
| AB | AB | AB | AB | - | AA | AB | AA | AA | AB | AB | AB | AB | AB |
| AB | AA | AB | AB | AB | AA | AB | AB | AA | AA | AB | AB | AA | AB |
| AA | AB | AA | AB | AA | AA | AA | AA | AA | AB | AA | AA | AB | AA |
| AA | AA | AA | AB | AA | AA | AB | AB | AA | AA | AA | AA | AA | AA |
| AA | AB | AB | AB | AB | AB | AA | AA | AB | AB | AB | AB | AA | AB |
| AB | AA | AB | AB | AB | AB | AA | AB | AB | AB | AB | AB | AB | AB |
| AB | AB | AB | AB | AB | AB | AB | AB | AB | AB | AB | AB | AB | AB |
| AA | AA | AA | AB | AA | AA | AA | AA | AA | AA | AA | AA | AA | AA |
| AA | AA | AA | AA | AA | AA | AA | AA | AA | AA | AA | AA | AA | AA |
| AB | AB | AB | AB | AB | AB | AA | AA | AB | AB | AB | AA | AB | AB |
| AA | AB | AA | AA | AA | AA | AA | AA | AB | AA | AA | AA | AA | AA |

| snp_B4801 5 | snp_B4812 5 | snp_B4839 5 | snp_B4956 5 | snp_B4960 5 | snp_B4968 5 | snp_B5003 5 | snp_B5044 5 | snp_B5045 5 | snp_B5050 5 | snp_B5060 5 | snp_B5069 5 | snp_B5087 5 | snp_B5138 5 | snp_B5142 5 | snp_B5144 5 | snp_B5197 5 | snp_B5204 5 | snp_B5214 5 |
|---|---|---|---|---|---|---|---|---|---|---|---|---|---|---|---|---|---|---|
| -- | -- | -- | -- | -- | -- | -- | -- | -- | -- | -- | -- | -- | -- | -- | -- | -- | -- | -- |
| AB | AB | AB | AB | AB | AA | AB | AB | AA | AA | AB | AA | AA | AB | AB | AB | AB | AA | AB |
| AA | AA | AB | AA | AA | AA | AA | AB | AB | AA | AB | AA | AB | AA | AA | AB | AA | AB | AA |
| AA | AB | AB | AA | AB | AB | AA | AB | AB | AA | AA | AA | AB | AA | AB | AA | AB | AA | AA |
| AB | AA | AA | AB | AB | AA | AB | AB | AB | AA | - | AA | AA | AA | AA | AB | AB | AA | AB |
| AA | AA | AA | AA | AB | AA | AB | AB | AB | AA | AA | AA | AB | AA | AB | AB | AB | AA | AA |
| AA | AB | - | AB | AB | - | AB | AA | AA | AB | - | AB | - | - | - | AA | - | AA | - |
| AB | AB | AA | AB | AB | AB | AB | AA | AB | AB | AB | AB | AB | AB | AA | AA | AB | AA | AB |
| AA | AA | AA | AA | AA | AA | AA | AB | AB | AA | AA | AA | AA | AA | AA | AA | AB | AA | AA |
| AB | AB | AB | AB | AB | AB | AB | AA | AB | AA | AA | AA | AB | AA | AB | AB | AA | AA | AB |
| AA | AA | AB | AA | AB | AB | AA | AB | AA | AB | AB | AB | AA | AB | AA | AA | AB | AB | AA |
| AB | AA | AB | AA | AB | AA | AA | AA | AA | AA | AA | AB | AB | AA | AB | AB | AB | AA | AA |
| AB | AB | AB | AB | AB | AA | AA | AB | AA | AB | AB | AB | AA | AA | AB | AB | AB | AA | AB |
| AA | AA | AB | AA | AA | AA | AA | AA | AB | AA | AB | AA | AB | AA | AA | AA | AB | AB | AA |
| AA | AB | AB | AB | AA | AA | AB | AB | AA | AB | AA | AB | AA | AA | AB | AA | AB | AA | AB |
| AB | AB | AB | AB | AA | AB | AB | AA | AB | AB | AB | AB | AA | AA | AB | AB | AA | AB | AA |
| AA | AB | AA | AB | AB | AB | AB | AA | AB | AA | - | AA | AB | AA | AA | AA | AB | AA | AA |
| AA | AA | AA | AA | AA | AA | AA | AA | AA | AA | AA | AA | AA | AA | AA | AA | AA | AA | AA |
| AB | AA | - | AB | AA | - | AB | AA | AB | - | - | - | - | - | AB | AB | AB | AA | - |
| AB | AB | AB | AB | AB | AB | AB | AA | AB | AB | AB | AB | AB | AB | AA | AB | AB | AB | AB |
| AA | AA | AB | AA | AB | AB | AB | AA | AB | AB | AB | AA | AA | AB | AA | AB | AB | AB | AB |
| AB | AB | AB | AB | AB | AB | AB | AB | AB | AB | AB | AB | AB | AB | AA | AB | AB | AB | AB |
| AB | AB | AB | AB | AB | AB | AB | AB | AB | AB | AB | AB | AB | AB | AA | AB | AB | AB | AB |
| AA | AB | AA | AA | AB | - | AA | AA | AA | AB | - | AA | - | - | AB | - | - | AA | - |
| AB | AA | AB | AA | AA | AB | AA | AB | AA | AA | AB | AB | AA | AA | AA | AB | AB | AB | AB |
| AA | AA | AB | AA | AA | AA | AA | AB | AB | AA | AA | AA | AA | AA | AB | AA | AB | AA | AA |
| AB | AB | AB | AB | AB | AB | AB | AB | AB | AB | AB | AB | AB | AB | AB | AB | AB | AB | AB |
| AA | AA | AA | AA | AA | AA | AA | AA | AA | AA | AA | AA | AA | AB | AA | AA | AA | AA | AA |
| AA | AA | AA | AA | AA | AB | AB | AA | AB | AB | AB | AB | AA | AB | AA | AA | AB | AB | AA |
| AA | AB | AA | AA | AB | AA | AB | AB | AA | AA | AA | AA | AB | AB | AA | AA | AB | AA | AB |
| AA | - | AB | AA | AA | AA | AA | AB | AB | AA | - | AA | - | - | AA | AA | AB | - | - |
| AB | AB | AB | AB | AB | AB | AB | AB | AB | AB | AB | AB | AB | AB | AB | AB | AB | AB | AB |
| AB | AB | AA | AB | AB | AA | AB | AA | AB | AB | AA | AB | AB | AA | AA | AB | AB | AB | AA |
| AA | AB | AA | AB | AB | AA | AA | AA | AB | AA | - | AA | - | AA | AB | AA | AA | AA | AB |
| AB | - | AB | AB | AA | AA | AB | AA | AB | AB | - | AA | AA | - | AA | AB | AB | AA | AA |
| AA | AB | AA | AB | AB | AB | AB | AB | AB | AA | AA | AA | - | AA | AB | AB | AB | AA | AA |
| AB | AB | AB | AB | AB | AB | AB | AB | AB | AB | AB | AB | - | - | AB | AB | AB | AB | AB |
| AA | AA | AA | AB | AA | AA | AB | AA | AA | AB | AA | AA | AA | AB | AA | AA | AA | AA | AA |
| AB | AB | AB | AB | AB | AB | AB | AB | AB | AB | AB | AB | AB | AB | AB | AB | AB | AB | AB |
| AB | AB | AB | AB | AB | AB | AB | AB | AB | AB | AB | AB | AB | AB | AB | AB | AB | AB | AB |
| AA | AA | AB | AA | AB | - | AB | AB | AB | AA | - | AA | - | - | AB | AB | AB | AB | AB |
| AB | AB | AA | AB | AB | AB | AB | AA | AB | AB | AB | AB | AB | AB | AA | AB | AB | AB | AB |
| AA | AA | AB | AA | AA | AA | AA | AB | AA | AA | AA | AA | AA | AA | AB | AA | AA | AA | AA |
| AA | AA | AA | AB | AA | AB | AB | AA | AA | AB | AB | AA | AA | AB | AA | AA | AA | AA | AA |
| AA | AA | AB | AA | AA | AA | AA | AB | AB | AA | - | AA | - | AA | AA | AB | AB | AB | AA |
| AA | AA | AB | AB | AA | AA | AB | AB | AB | AB | AA | AB | AB | AA | AB | AA | AB | AA | AB |
| AB | AB | AB | AA | AB | AB | AB | AB | AB | AB | AB | AB | AB | AB | AB | AB | AB | AB | AB |
| AA | AA | AA | AA | AA | AA | AA | AA | AA | AA | AA | AA | AA | AA | AA | AA | AA | AA | AA |
| AA | AA | AA | AA | AA | AA | AA | AA | AA | AA | AA | AA | AA | AA | AA | AA | AA | AA | AA |
| AB | AB | AB | AB | AA | AA | AA | AB | AB | AA | AB | AB | AB | AA | AB | AB | AB | AA | AA |
| AA | AA | AB | AA | AB | AA | AA | AB | AA | AA | AA | AA | AA | AA | AB | AA | AB | AA | AA |

| snp_B5220 | snp_B5236 | snp_B5262 | snp_B5296 | snp_B5316 | snp_B5326 | snp_B5340 | snp_B5381 | snp_B5384 | snp_B5414 | snp_B5449 | snp_B5468 | snp_B5479 | snp_B5497 | snp_B5623 | snp_B5635 | snp_B5641 | snp_B5645 | snp_B5647 | snp_B5660 | snp_B5662 | snp_B5669 | snp_B5706 | snp_B5722 | snp_B5729 | snp_B5733 | snp_B5734 | snp_B5737 | snp_B5751 | snp_B5756 | snp_B5760 | snp_B5788 | snp_B5790 | snp_B5792 | snp_B5802 | snp_B5810 | snp_B5827 | snp_B5834 | snp_B5852 | snp_B5854 | snp_B5866 | snp_B5872 | snp_B5874 | snp_B5916 | snp_B5967 | snp_B5999 | snp_B6014 | snp_B6095 | snp_B6120 | snp_B6165 | snp_B0004 | snp_B0021 | snp_B0023 | snp_B0041 | snp_B0046 | snp_B0052 | snp_B0053 | snp_B0112 | snp_B0120 | snp_B0140 | snp_B0142 | snp_B0150 | snp_B0154 | snp_B0155 | snp_B0162 | snp_B0163 | snp_B0168 | snp_B0269 | snp_B0323 | snp_B0382 | snp_B0393 | snp_B0453 | snp_B0488 | snp_B0493 | snp_B0508 | snp_B0525 | snp_B0526 |
|---|---|---|---|---|---|---|---|---|---|---|---|---|---|---|---|---|---|---|---|---|---|---|---|---|---|---|---|---|---|---|---|---|---|---|---|---|---|---|---|---|---|---|---|---|---|---|---|---|---|---|---|---|---|---|---|---|---|---|---|---|---|---|---|---|---|---|---|---|---|---|---|---|---|---|---|---|
| 5 | 5 | 5 | 5 | 5 | 5 | 5 | 5 | 5 | 5 | 5 | 5 | 5 | 5 | 5 | 5 | 5 | 5 | 5 | 5 | 5 | 5 | 5 | 5 | 5 | 5 | 5 | 5 | 5 | 5 | 5 | 5 | 5 | 5 | 5 | 5 | 5 | 5 | 5 | 5 | 5 | 5 | 5 | 5 | 5 | 5 | 5 | 5 | 5 | 5 | 6 | 6 | 6 | 6 | 6 | 6 | 6 | 6 | 6 | 6 | 6 | 6 | 6 | 6 | 6 | 6 | 6 | 6 | 6 | 6 | 6 | 6 | 6 | 6 | 6 | 6 | 6 |
| -- | -- | -- | LG13 | -- | -- | -- | -- | -- | -- | -- | -- | -- | -- | -- | -- | -- | -- | -- | -- | -- | -- | -- | LG13 | -- | -- | -- | -- | -- | -- | -- | LG13 | -- | -- | -- | -- | -- | -- | -- | -- | -- | -- | -- | -- | -- | -- | -- | -- | -- | -- | -- | -- | -- | -- | -- | -- | -- | -- | -- | -- | -- | -- | -- | -- | -- | -- | -- | -- | -- | -- | -- | -- | -- | -- | -- | -- | -- |
| AA | AA | AA | AA | AB | AA | AB | AA | AB | AA | AB | AB | AB | AA | AB | AB | AB | AB | AA | AA | AB | AA | AB | AB | AA | AB | AB | AA | AA | AA | AB | AB | AB | AA | AA | AB | AB | AB | AA | AB | AA | AA | AA | AA | AA | AA | AB | AA | AB | AA | AA | AA | AA | AA | AB | AA | AA | AB | AA | AA | AA | AB | AA | AA | AA | AA | AA | AB | AA | AA | AA | AA | AA | AA | AB | AB | AB |

| snp_B0542 6 | snp_B0547 6 | snp_B0560 6 | snp_B0618 6 | snp_B0694 6 | snp_B0698 6 | snp_B0723 6 | snp_B0728 6 | snp_B0810 6 | snp_B0836 6 | snp_B0867 6 | snp_B0875 6 | snp_B0911 6 | snp_B0921 6 | snp_B0955 6 | snp_B0956 6 | snp_B0962 6 | snp_B0982 6 | snp_B1018 6 | snp_B1031 6 | snp_B1038 6 | snp_B1040 6 |
|---|---|---|---|---|---|---|---|---|---|---|---|---|---|---|---|---|---|---|---|---|---|
| -- | -- | -- | -- | LG8 | -- | -- | -- | -- | -- | -- | -- | -- | -- | -- | -- | LG8 | -- | -- | -- | LG8 | -- |
| AA | AA | AB | AA | AB | AA | AA | AB | AA | AA | AA | AA | AA | AA | AA | AA | AB | AB | AA | AB | AB | AA |
| AA | AA | AA | AB | AB | AA | AA | AB | AA | AA | AA | AA | AA | AA | AA | AA | AB | AB | AA | AB | AB | AA |
| AB | AB | AA | AB | AA | AA | AB | AA | AB | AB | AA | AB | AB | AB | AB | AB | AA | AA | AA | AB | AA | AB |
| AB | - | AA | AB | AA | AA | - | AB | AA | - | AA | AB | AB | AB | AB | AB | AA | AA | AB | AB | AA | AB |
| AB | AA | AB | AA | AB | AB | AB | AB | AB | AA | AB | AB | AB | AB | AA | AB | AB | AB | AB | AA | AB | AB |
| AA | - | - | AA | AA | AA | AA | AA | AB | - | AB | AB | AB | - | AA | AA | - | AA | AA | AA | AA | AB |
| - | AA | AB | AB | AB | AA | AB | AA | AB | AA | AB | AA | AA | AA | AB | AB | AA | AA | AA | AA | AB | AA |
| AB | AB | AA | AB | AA | AB | AB | AA | AB | AB | AA | AA | AA | AB | AB | AB | AA | AB | AB | AB | AA | AB |
| AB | AB | AA | AB | AA | AB | AB | AB | AA | AB | AA | AA | AA | AA | AB | AB | AB | AA | AB | AB | AB | AB |
| AB | AB | AA | AB | AA | AB | AA | AB | AA | AB | AA | AA | AA | AA | AB | AB | AA | AB | AA | AA | AA | AA |
| AB | AB | AA | AB | AB | AA | AB | AA | AB | AA | AA | AB | AB | AA | AB | AB | AB | AA | AB | AB | AA | AB |
| AB | AA | AA | AB | AB | AB | AB | AB | AA | AB | AA | AA | AA | AA | AB | AA | AA | AA | AA | AB | AB | AA |
| AB | AB | AA | AB | AB | AA | AB | AB | AA | AB | AA | AA | AA | AA | AB | AA | AA | AA | AA | AA | AB | AA |
| AB | AB | AA | AB | AA | AB | AB | AB | AA | AA | AA | AA | AA | AA | AA | AA | AA | AA | AA | AA | AB | AA |
| AB | AB | AA | AB | AA | AA | AB | AA | AB | AA | AA | AB | AB | AB | AA | AA | AA | AA | AB | AB | AA | AB |
| AB | AB | AA | AB | AA | AA | AB | AB | AB | AB | AA | AB | AB | AB | AB | AA | AA | AB | AB | AB | AA | AB |
| AB | AB | AA | AB | AA | AA | AB | AA | AB | AA | AB | AA | AA | AA | AA | AA | AA | AA | AA | AA | AA | AA |
| - | AA | - | AA | AB | AB | AA | AA | AA | AA | AB | AA | AA | - | AA | AA | AB | AA | - | AA | - | AA |
| AB | AB | AB | AB | AB | AB | AB | AB | AB | AB | AB | AB | AB | AB | AB | AB | AB | AA | AB | AB | AB | AB |
| AB | AB | AB | AB | AB | AB | - | AB | AA | AA | AB | AA | AA | AA | AB | - | AB | AB | AA | AA | AB | AA |
| AB | AB | AB | AB | AB | AB | AA | AB | AB | AB | AB | AA | AB | AB | AB | AB | AA | AB | AB | AB | AB | AB |
| AA | AA | AB | AB | AB | AB | AA | AB | AA | AA | AB | AA | AA | AA | AB | AA | AA | AB | AB | AB | AB | AA |
| - | - | - | AA | AB | AA | - | AA | AB | - | AB | AB | - | AB | AA | - | AB | AA | AA | AB | - | AB |
| AA | AA | AA | AA | AB | AB | AA | AB | AA | AB | AB | AB | AB | AB | AA | AA | AA | AB | AB | AB | AB | AB |
| AB | AB | AB | AB | AA | AA | AB | AA | AB | AB | AB | AB | AA | AA | AB | AB | AA | AA | AA | AA | AA | AA |
| AA | AA | AB | AB | AB | AA | AA | AA | AA | AA | AB | AA | AB | AA | AA | AB | AB | AA | AA | AA | AB | AA |
| AB | AB | AB | AB | AA | AA | AB | AA | AB | AB | AB | AB | AB | AB | AB | AB | AA | AA | AB | AB | AA | AB |
| AB | AB | AA | AB | AA | AA | AB | AA | AB | AB | AB | AB | AB | AB | AB | AB | AA | AA | AB | AB | AA | AB |
| AA | - | AA | AB | AA | AA | - | AB | AA | AA | AA | AA | AB | AA | AB | - | AA | AB | AB | AB | AA | AA |
| - | - | - | - | AA | AA | - | AB | AB | - | - | AB | AB | AB | AA | - | AB | AA | AB | AB | - | AB |
| AB | - | AA | AB | AA | AB | AB | AB | AA | AA | AB | AA | AA | AA | AB | AA | AA | AB | AA | AA | AA | AA |
| AA | AA | AB | AA | AB | AB | AB | AA | AB | AA | AA | AA | AA | AA | AB | AB | AB | AB | AA | AA | AB | AA |
| - | - | AB | AA | AA | AB | AA | AB | AB | - | AB | AB | AB | AB | AA | - | AB | AB | - | AB | - | AB |
| AA | - | - | AB | AA | AA | AA | AA | AB | - | AB | AA | AA | AA | AA | AA | AB | AA | - | AA | - | AA |
| AB | AB | AA | AB | AA | AA | AB | AB | AA | - | AA | AB | AA | AB | AB | AA | AA | AA | AB | AB | AA | AB |
| AB | AB | - | AB | AB | AB | AB | AB | AB | AB | AB | AB | AB | AB | AB | AB | AB | AB | AB | AB | - | AB |

| snp_B1092 6 | snp_B1106 6 | snp_B1113 6 | snp_B1119 6 | snp_B1139 6 | snp_B1155 6 | snp_B1180 6 | snp_B1186 6 | snp_B1194 6 | snp_B1212 6 | snp_B1254 6 | snp_B1255 6 | snp_B1289 6 | snp_B1296 6 | snp_B1313 6 | snp_B1314 6 | snp_B1325 6 | snp_B1367 6 | snp_B1377 6 | snp_B1404 6 | snp_B1423 6 | snp_B1439 6 | snp_B1443 6 | snp_B1444 6 | snp_B1460 6 | snp_B1462 6 |
|---|---|---|---|---|---|---|---|---|---|---|---|---|---|---|---|---|---|---|---|---|---|---|---|---|---|
| -- | -- | -- | -- | -- | -- | -- | -- | -- | -- | -- | LG8 | -- | -- | -- | -- | -- | -- | -- | -- | -- | LG8 | -- | -- | -- | -- |
| AB | AB | AA | AB | AB | AA | AB | AB | AA | AA | AB | AB | AB | AA | AB | AA | AA | AA | AA | AB | AA | AB | AA | AA | AB | AA |
| AB | AB | AB | AB | AB | AB | AA | AA | AA | AA | AA | AB | AA | AA | AB | AB | AA | AA | AA | AA | AA | AB | AA | AB | AB | AA |
| AB | AA | AB | AB | AA | AA | AB | AA | AA | AB | AA | AA | AA | AB | AA | AA | AB | AB | AB | AA | AB | AA | AB | AA | AA | AB |
| AA | AA | - | AA | AA | AA | AA | AB | AB | AB | AA | AA | AA | AB | AA | AA | AB | AB | AB | AA | AB | AA | AB | AB | - | AB |
| AB | AB | AB | AB | AB | AB | AB | AB | AB | AB | AB | AB | AB | AA | AB | AB | AB | AB | AA | AB | AB | AB | AA | AA | AB | AA |
| AA | AA | AB | AA | AA | - | - | AA | AA | AB | AB | AA | - | AA | - | AA | AB | - | AA | - | - | - | AA | - | - | - |
| AB | AA | AA | AB | AA | AB | AA | AA | AB | AA | AB | AA | AB | AA | AA | AB | AA | AB | AA | AA | AA | AA | AA | AB | AB | AB |
| AA | AA | AB | AA | AA | AA | AB | AB | AB | AB | AA | AA | AA | AA | AA | AA | AB | AB | AB | AB | AB | AA | AB | AA | AA | AB |
| AB | AA | AA | AA | AB | AB | AA | AB | AA | AB | AA | AB | AA | AB | AB | AB | AA | AA | AB | AB | AB | AA | AB | AA | AB | AB |
| AB | AA | AA | AA | AA | AB | AA | AB | AA | AA | AB | AB | AA | AB | AB | AB | AA | AA | AB | AB | AB | AA | AA | AA | AB | AB |
| AA | AA | AB | AA | AB | AB | AB | AA | AB | AB | AA | AA | AA | AA | AA | AA | AB | AB | AB | AA | AB | AA | AA | AB | AA | AB |
| AB | AB | AA | AA | AB | AA | AA | AB | AA | AA | AB | AB | AA | AB | AB | AA | AA | AA | AB | AA | AB | AB | AA | AB | AA | AB |
| AA | AB | AA | AA | AB | AA | AA | AB | AA | AA | AA | AB | AA | AB | AB | AA | AA | AA | AB | AA | AB | AB | AA | AB | AB | AB |
| AA | AB | AA | AA | AB | AB | AA | AB | AB | AA | AA | AB | AA | AA | AA | AA | AA | AA | AB | AB | AA | AB | AB | AA | AA | AA |
| AA | AB | AB | AA | AB | AA | AB | AB | AB | AB | AA | AA | AA | AA | AA | AA | AB | AB | AB | AB | AB | AA | AB | AA | AB | AB |
| AB | AB | AB | AA | AB | AA | AB | AB | AB | AA | AA | AB | AA | AB | AB | AB | AB | AB | AB | AB | AB | AA | AB | AA | AB | AA |
| AB | AA | AA | AA | AA | AA | AA | AA | AA | AA | AA | AA | AA | AA | AA | AA | AA | AB | AA | AA | AA | AA | AA | AB | AA | AA |
| - | AB | - | AB | AB | - | AB | AA | AA | - | - | AB | - | AA | - | AA | AA | - | AA | - | AA | AB | AA | - | AB | AB |
| AB | AB | AB | AB | AB | AB | AB | AB | AB | AB | AB | AB | AB | AB | AB | AB | AB | AA | AB | AB | AB | AB | AB | AB | AB | AB |
| AB | AA | AA | AB | AB | AB | AA | AB | AB | AA | AB | AB | AB | AA | AB | AB | AB | AB | AB | AA | AB | AB | AB | AB | - | AB |
| AB | AB | AB | AA | AB | AB | AB | AB | AB | AA | AB | AB | AB | AA | AB | AB | AB | AB | AB | AB | AB | AB | AB | AB | AB | AB |
| AB | AB | AA | AA | AB | AB | AB | AA | AB | AA | AB | AB | AB | AA | AB | AB | AA | AA | AA | AA | AB | AB | AA | AB | AA | AB |
| AB | AB | AB | AB | AB | - | - | AB | AB | AB | AA | AB | - | AA | - | AA | AB | - | - | - | - | - | AB | - | AB | AA |
| AB | AB | AB | AB | AB | AB | AB | AB | AB | AB | AA | AB | AA | AB | AB | AB | AB | AB | AA | AB | AB | AB | AB | AB | AA | AA |
| AA | AA | AB | AA | AB | AA | AA | AB | AA | AB | AB | AA | AB | AB | AA | AB | AA | AB | AB | AA | AB | AA | AB | AB | AA | AB |
| AA | AA | AB | AA | AB | AA | AB | AB | AA | AB | AB | AA | AB | AB | AA | AA | AA | AA | AB | AA | AB | AB | AA | AB | AA | AB |
| AA | AA | AB | AA | AB | AA | AB | AB | AA | AB | AB | AA | AB | AB | AA | AB | AB | AB | AB | AA | AB | AA | AA | AB | AB | AB |
| AA | AA | AB | AA | AB | AA | AB | AA | AB | AA | AB | AB | AA | AB | AB | AA | AB | AB | AB | AB | AA | AB | AA | AA | AB | AB |
| - | - | AB | AA | AB | AA | AA | AA | AA | AB | AB | AB | AA | AB | AB | AB | AB | AA | AB | AB | AB | AA | AB | AB | AA | AB |
| - | AA | - | AB | AA | - | - | AB | AB | AB | AB | AA | AB | AB | - | AB | AB | - | AB | - | AB | AA | AB | - | - | AB |
| AB | AA | AA | AB | AB | AA | AA | AB | AB | AA | AA | AB | AA | AA | AB | AA | AA | AA | AA | AA | AB | AB | AB | AB | AA | AB |
| AA | AA | AA | AB | AB | AB | AA | AA | AA | AB | AB | AB | AB | AA | AB | AB | AA | AB | AA | AA | AB | AB | AB | AA | AB | AA |
| - | - | AB | AB | AB | AB | AB | AB | AA | AB | AB | AB | AB | AB | AB | AB | AA | AB | AB | AA | AB | AB | AB | - | - | AA |
| - | AA | AB | AB | AA | AA | AB | AB | AA | AB | AB | AA | - | AA | AA | AA | AA | AB | AA | AA | AA | AB | AA | - | - | AA |
| - | AB | AB | AA | AB | - | AA | AB | AA | AB | AA | AB | - | AB | AB | AB | AB | - | AB | - | AB | AA | AB | - | - | AA |
| AB | AB | AB | AB | AB | AB | AB | AB | AB | AB | AB | AB | - | AB | AB | AB | AB | - | AB | AB | AB | AB | AB | AB | - | AB |

| snp_B1478 6 | snp_B1496 6 | snp_B1503 6 | snp_B1514 6 | snp_B1546 6 | snp_B1560 6 | snp_B1595 6 | snp_B1606 6 | snp_B1609 6 | snp_B1625 6 | snp_B1630 6 | snp_B1677 6 | snp_B1698 6 | snp_B1707 6 | snp_B1765 6 | snp_B1804 6 | snp_B1810 6 | snp_B1836 6 | snp_B1877 6 | snp_B1909 6 | snp_B1914 6 | snp_B1916 6 | snp_B1930 6 | snp_B1932 6 | snp_B1949 6 | snp_B1995 6 | snp_B1998 6 | snp_B2010 6 | snp_B2011 6 |
|---|---|---|---|---|---|---|---|---|---|---|---|---|---|---|---|---|---|---|---|---|---|---|---|---|---|---|---|---|
| -- | -- | -- | -- | -- | -- | -- | -- | -- | -- | -- | -- | -- | -- | -- | -- | -- | -- | -- | -- | -- | -- | -- | -- | -- | -- | -- | -- | -- |
| AB | AB | AB | AB | AB | AA | AB | AA | AA | AA | AB | AB | AB | AA | AB | AA | AA | AB | AB | AA | AA | AA | AA | AB | AB | AB | AA | AB | AA |
| AA | AA | AB | AA | AB | AA | AA | AA | AA | AA | AB | AA | AA | AB | AA | AB | AA | AB | AA | AA | AA | AA | AB | AA | AA | AA | AA | AB | AA |
| AA | AA | AB | AA | AB | AA | AA | AA | AA | AB | AA | AA | AB | AB | AA | AB | AA | AB | AA | AB | AA | AA | AB | AA | AA | AA | AA | AB | AA |
| AB | AA | AA | AA | AB | AB | AA | AA | AB | - | AA | AA | AB | AA | AA | AB | AA | AB | - | AB | AB | AA | AB | AA | AB | AA | AA | AB | AB |
| AB | AB | AB | AB | AB | AB | AA | AB | AB | AB | AB | AB | AB | AB | AA | AA | AB | AB | AB | AA | AB | AB | AA | AB | AB | AB | AA | AB | AB |
| AB | AB | AA | - | AA | AA | AA | - | AA | - | AA | - | - | AA | AB | - | - | AB | - | - | - | AB | AA | AA | AB | AB | AB | - | AA |
| AA | AB | AB | AB | AA | AB | AB | AA | AA | AA | AB | AA | AA | AB | AB | AB | AA | AA | AA | AB | AA | AB | AB | AA | AA | AB | AB | AA | AA |
| AB | AB | AA | AB | AA | AB | AB | AA | AB | AB | AA | AB | AB | AA | AB | AB | AA | AA | AB | AB | AB | AA | AB | AA | AB | AA | AA | AB | AB |
| AB | AB | AA | AB | AA | AB | AB | AA | AB | AB | AA | AB | AB | AA | AB | AB | AA | AA | AB | AB | AB | AA | AB | AB | AB | AA | AA | AA | AB |
| AA | AA | AA | AA | AB | AB | AB | AB | AB | AB | AA | AB | AA | AA | AB | AB | AB | AB | AA | AB | AB | AA | AB | AB | AA | AA | AB | AA | AB |
| AA | AB | AA | AA | AB | AB | AA | AB | AA | AA | AA | AA | AA | AB | AB | AB | AB | AA | AB | AB | AA | AB | AB | AA | AA | AA | AB | AB | AA |
| AB | AA | AB | AA | AA | AB | AB | AB | AB | AB | AB | AB | AA | AA | AA | AB | AA | AB | AA | AB | AA | AA | AA | AB | AB | AB | AA | AA | AA |
| AA | AA | AA | AA | AA | AB | AA | AA | AA | AA | AA | AA | AA | AA | AA | AB | AA | AA | AB | AB | AA | AA | AA | AA | AA | AA | AA | AA | AA |
| AB | AA | AA | AA | AB | AA | AB | AB | AA | AB | AB | AB | AB | AB | AB | AB | AA | AA | AB | AA | AA | AA | AB | AA | AB | AA | AA | AA | AA |
| AB | AB | AA | AB | AB | AB | AA | AB | AA | AB | AA | AA | AA | AA | AB | AB | AB | AA | AB | AB | AA | AA | AB | AB | AA | AA | AA | AB | AB |
| AB | AA | AB | AB | AA | AB | AA | AA | AB | AB | AB | AB | AB | AB | AB | AB | AB | AB | AB | AB | AA | AA | AB | AA | AB | AB | AA | AB | AB |
| AA | AB | AA | AB | AA | AA | AB | AA | AA | AA | AA | AA | AA | AB | AA | AA | AA | AA | AA | AA | AA | AA | AB | AA | AB | AA | AA | AA | AA |
| AA | AA | AA | - | AA | AB | AB | AB | AA | - | AA | - | AB | AA | AB | - | AA | - | - | - | AA | - | AA | AB | - | AA | AB | AB | AA |
| AB | AB | AB | AB | AB | AB | AB | AB | AB | AB | AB | AB | AB | AB | AB | AB | AB | AB | AB | AB | AB | AB | AB | AA | AB | AB | AB | AB | AB |
| AB | AB | AB | AB | AB | AA | AA | AB | AB | AB | AB | AA | AB | AB | AB | AB | AA | AB | AA | AB | AA | AB | AB | AA | AB | AB | AB | AA | AA |
| AB | AB | AB | AB | AB | AB | AB | AB | AB | AB | AB | AB | AB | AA | AB | AB | AB | AB | AB | AB | AB | AB | AB | AA | AB | AB | AB | AB | AB |
| AA | AB | AB | AA | AB | AB | AB | AB | AB | AB | AB | AB | AB | AA | AB | AB | AB | AB | AB | AB | AB | AB | AB | AA | AB | AA | AA | AA | AB |
| - | AB | AA | - | AB | AA | AB | - | AA | - | AB | - | - | AA | AA | - | - | AA | AB | - | - | AA | AA | AB | - | AA | AB | - | AB |
| AB | AB | AA | AA | AB | AB | AA | AB | AB | AB | AB | AB | AB | AB | AB | AB | AB | AB | AB | AB | AB | AA | AA | AA | AB | AA | AA | AB | AB |
| AA | AB | AB | AA | AA | AB | AA | AB | AA | AA | AB | AA | AA | AB | AB | AA | AB | AA | AA | AA | AB | AA | AB | AA | AA | AB | AA | AA | AA |
| AB | AB | AA | AA | AA | AB | AA | AB | AB | AB | AA | AB | AA | AA | AB | AA | AB | AA | AA | AA | AB | AB | AB | AB | AB | AA | AB | AA | AB |
| AA | AB | AA | AB | AB | AA | AA | AA | AB | AA | AB | AA | AA | AA | AA | AA | AA | AA | AB | AB | AB | AB | AB | AB | AA | AB | AA | AA | AB |
| AA | AB | AA | AB | AB | AA | AA | AA | AB | AA | AB | AA | AA | AA | AB | AA | AA | AA | AB | AB | AB | AB | AB | AB | AB | AA | AB | AA | AB |
| AB | AB | AA | - | AA | AA | AA | AA | AB | AB | AA | AA | AB | AA | AA | AB | - | AB | AB | AB | AB | AA | AB | AA | AA | AB | AA | AA | AB |
| - | AB | AA | - | AB | AA | AA | - | AA | - | AB | AA | - | AB | AB | - | - | AA | - | AA | - | AB | AB | AB | - | AA | AB | - | AB |
| AB | AB | AB | AB | AA | AA | AA | AB | AB | AB | AB | AA | AB | AA | AA | AB | AA | AB | AA | AA | AA | AA | AA | AA | AB | AB | AA | AA | AA |
| AB | AB | AA | AB | AA | AA | AA | AB | AB | AB | AA | AA | AB | AB | AA | AA | AA | AB | AB | AA | AA | AA | AB | AB | AB | AA | AB | AB | AA |
| AB | AB | AA | AB | AB | AB | AB | - | AB | - | AA | - | AB | AA | AB | AA | - | AB | AB | AA | AB | AB | AA | AB | - | AB | AB | AB | AB |
| AA | AB | AB | AB | AA | AA | AA | - | AA | AA | AA | AA | AA | AA | AB | AA | - | AA | - | AA | AA | AB | AA | AB | AA | AB | AB | - | AA |
| AB | AA | AB | - | AB | AA | AB | - | AA | - | AB | AA | AB | AB | AB | - | - | AB | - | - | AA | AA | AB | AA | - | AB | AA | - | AA |
| AB | AB | AB | AB | AB | AB | AB | AA | AB | AB | AB | AB | AB | AB | AB | AB | - | AB | AB | AB | AB | AB | AB | AA | AB | AB | AB | AB | AB |

| snp_B2053 | snp_B2057 | snp_B2062 | snp_B2074 | snp_B2079 | snp_B2090 | snp_B2098 | snp_B2101 | snp_B2109 | snp_B2121 | snp_B2133 | snp_B2134 | snp_B2147 | snp_B2150 | snp_B2159 | snp_B2172 | snp_B2182 | snp_B2190 | snp_B2221 | snp_B2232 |
|---|---|---|---|---|---|---|---|---|---|---|---|---|---|---|---|---|---|---|---|
| 6 | 6 | 6 | 6 | 6 | 6 | 6 | 6 | 6 | 6 | 6 | 6 | 6 | 6 | 6 | 6 | 6 | 6 | 6 | 6 |
| -- | -- | -- | -- | -- | -- | LG8 | -- | -- | -- | LG8 | -- | -- | -- | -- | -- | -- | -- | -- | -- |
| AA | AB | AB | AA | AA | AB | AB | AB | AA | AA | AB | AA | AA | AA | AB | AA | AA | AB | AB | AA |
| AA | AA | AB | AA | AA | AB | AB | AA | AA | AA | AB | AA | AB | AA | AA | AA | AA | AA | AA | AB |
| AB | AB | AA | AB | AB | AB | AA | AA | AB | AB | AA | AB | AB | AB | AA | AA | AB | AA | AA | AB |
| AB | AB | AA | AA | AB | AB | AA | - | AB | AB | AA | AB | AA | AB | AA | AA | AA | AA | AA | AA |
| AA | AA | AB | - | AB | AB | AB | AB | AB | AB | AB | AA | AA | AA | AA | AB | AB | AB | AB | AA |
| AA | AA | - | AA | AA | AB | AA | - | - | - | AA | - | - | AA | AA | AA | - | - | AA | AA |
| AB | AB | AB | AA | AB | AA | AA | AB | - | AA | AB | AA | - | AB | AA | AA | AA | AB | AB | AB |
| AB | AB | AA | AA | AB | AB | AA | AA | AB | AB | AA | AB | AA | AB | AB | AA | AB | AA | AA | AA |
| AA | AA | AA | AA | AA | AA | AB | AB | AA | AA | AB | AB | AA | AB | AB | AB | AA | AA | AA | AB |
| AB | AA | AB | AA | AB | AA | AB | AA | AA | AB | AA | AA | AB | AB | AB | AB | AA | AA | AB | AB |
| AB | AB | AB | AB | AB | AA | AA | AA | AB | AB | AB | AA | AB | AB | AA | AA | AB | AA | AA | AA |
| AB | AB | AA | AB | AA | AA | AB | AB | AA | AB | AB | AB | AB | AB | AB | AB | AA | AA | AA | AA |
| AB | AB | AA | AA | AB | AA | AB | AA | AA | AB | AB | AB | AB | AB | AA | AA | AA | AA | AA | AA |
| AA | AB | AA | AB | AB | AA | AB | AA | AA | AB | AA | AB | AB | AA | AA | AB | AA | AA | AA | AA |
| AB | AA | AA | AB | AB | AA | AA | AB | AB | AB | AA | AB | AB | AB | AB | AA | AB | AA | AA | AB |
| AB | AA | AA | AB | AB | AB | AA | AB | AB | AB | AA | AA | AB | AB | AB | AB | AB | AA | AB | AB |
| AA | AB | AA | AA | AB | AA | AA | AA | AB | AA | AA | AA | AA | AA | AA | AA | AA | AA | AA | AA |
| - | AB | - | - | AB | - | - | - | - | AA | - | - | AA | AA | AB | - | AA | - | AB | AB |
| AB | AB | AB | AA | AB | AB | AB | AB | AB | AB | AB | AB | AB | AB | AB | AB | AB | AB | AB | AB |
| AB | AB | AB | - | AB | AA | AB | AB | AA | AA | AB | AB | AB | AB | AA | - | AA | AB | AB | AB |
| AB | AA | AB | AB | AB | AB | AB | AA | AB | AB | AB | AB | AB | AB | AB | AB | AB | AB | AB | AB |
| AA | AB | AB | AB | AA | AA | AB | AB | AA | AB | AB | AB | AA | AB | AB | AB | AA | AB | AB | AB |
| AA | AA | AB | - | AA | AB | AA | - | - | AB | - | AA | - | AB | AA | - | - | - | AB | AA |
| AA | AB | AB | AB | AA | AB | AB | AA | AB | AB | AB | AB | AA | AB | AB | AB | AB | AB | AA | AB |
| AB | AA | AA | AA | AA | AB | AA | AB | AA | AB | AA | AA | AA | AB | AA | AA | AA | AB | AA | AA |
| AA | AB | AB | AB | AA | AA | AB | AA | AA | AA | AB | AB | AA | AA | AB | AB | AA | AB | AA | AA |
| AB | AB | AA | AA | AB | AA | AA | AA | AB | AB | AA | AB | AB | AB | AA | AA | AB | AA | AA | AB |
| AB | AB | AA | AA | AB | AA | AA | AA | AB | AB | AA | AB | AB | AB | AA | AA | AB | AB | AA | AB |
| AB | AB | AA | AA | AB | AA | AB | - | - | AB | AA | AA | AB | AB | AA | - | AB | AB | AB | AB |
| AA | AA | AB | AA | AB | AB | AA | - | - | AB | AA | - | - | AA | AB | - | AB | - | AB | AB |
| AB | AB | AA | AA | AB | AA | AB | AA | AA | AA | AB | AB | AB | AA | AA | AA | AA | AA | AA | AB |
| AA | AA | AB | AA | AA | AA | AB | AB | AA | AA | AA | AB | AA | AA | AA | AB | AA | AB | AB | AA |
| AA | AA | AA | AA | AA | AB | - | AA | AB | AB | - | AB | AA | AB | AB | AB | AB | AB | AB | AA |
| AA | AA | AA | AA | AA | AA | AA | AB | - | AA | AA | AA | - | AA | AA | - | - | AB | AB | AA |
| AB | AA | - | AA | AB | AB | AA | AB | - | - | AA | AA | - | AB | AA | AA | - | - | AB | AB |
| AB | AB | AB | - | AB | AB | AB | AB | AB | AB | AB | AB | AB | AB | AB | - | AB | - | AB | AB |
| AB | AB | AB | AB | AB | AB | AB | AB | AB | AB | AB | AB | AB | AB | AB | AB | AB | AA | AB | AB |
| AB | AB | AB | AA | AB | AB | AB | AB | AB | AB | AB | AB | AB | AB | AB | AB | AB | AB | AB | AB |
| AA | AA | AA | AA | AA | AA | AA | AA | AA | AA | AA | AA | AA | AA | AA | AA | AA | AB | AA | AA |
| AB | AB | AA | - | AA | AB | AB | AA | AA | AA | AB | - | AB | AB | AB | - | AB | - | AA | AB |
| AB | AA | AA | AA | AB | AB | AA | AB | AB | AB | AA | AA | AB | AB | AA | AA | AB | AA | AB | AB |
| AB | AB | AA | AB | AB | AA | AA | AA | AA | AA | AB | AA | AB | AB | AB | AB | AA | AA | AA | AB |
| AA | AA | AA | AB | AA | AB | AA | AA | AB | AB | AB | AA | AB | AB | AA | AB | AA | AB | AA | AA |
| AA | AB | AA | AA | AB | AA | AA | AB | AA | AA | AA | AA | - | AA | AA | AA | - | AB | AB | AB |
| AA | AA | AB | AA | AA | AB | AA | AB | AB | AA | AB | AB | AA | AA | AA | AA | AA | AB | AB | AB |
| AA | AA | AB | AA | AA | AB | AB | AB | AA | AA | AB | AA | AA | AA | AB | AB | AA | AB | AB | AA |
| AA | AA | AB | AA | AA | AB | AB | AA | AB | AB | AB | AA | AA | AB | AB | AA | AB | AB | AB | AB |
| AA | AA | AA | AA | AA | AB | AA | AA | AB | AA | AA | AA | AA | AB | AA | AA | AB | AB | AA | AB |
| AA | AA | AA | AA | AA | AA | AA | AA | AA | AA | AA | AA | AA | AA | AA | AA | AB | AA | AA | AA |
| AA | AA | AA | AA | AA | AA | AA | AA | AA | AA | AA | AA | AA | AA | AA | AA | AA | AA | AA | AA |
| AB | AB | AA | AB | AA | AA | AA | AA | AB | AB | AA | AA | AB | AB | AA | AB | AB | AA | AA | AA |
| AB | AB | AA | AB | AB | AB | AA | AA | - | AA | AA | AB | AB | AB | AA | AB | - | - | AA | AA |
| AB | AB | AB | AB | AB | AB | AB | AB | AB | AB | AB | AB | AB | AB | AB | AB | AB | AB | AB | AB |
| AB | AB | AB | AB | AB | AB | AB | AB | AB | AB | AB | AB | AB | AB | AB | AB | AB | AB | AB | AB |
| [illegible] | | | | | | | | | | | | | | | | | | | |

| snp_B3258 6 | snp_B3269 6 | snp_B3339 6 | snp_B3353 6 | snp_B3377 6 | snp_B3394 6 | snp_B3400 6 | snp_B3420 6 | snp_B3439 6 | snp_B3443 6 | snp_B3462 6 | snp_B3520 6 | snp_B3525 6 | snp_B3527 6 | snp_B3576 6 | snp_B3592 6 | snp_B3597 6 | snp_B3600 6 | snp_B3627 6 | snp_B3639 6 | snp_B3670 6 | snp_B3681 6 |
|---|---|---|---|---|---|---|---|---|---|---|---|---|---|---|---|---|---|---|---|---|---|
| -- | -- | -- | -- | -- | -- | -- | -- | -- | -- | -- | -- | -- | -- | -- | -- | -- | -- | -- | -- | -- | -- |
| AA | AA | AA | AB | AB | AA | AA | AB | AA | AB | AA | AA | AB | AB | AB | AA | AA | AB | AA | AB | AB | AA |
| AA | AA | AA | AA | AA | AA | AA | AA | AA | AB | AA | AA | AB | AB | AB | AA | AA | AB | AA | AA | AB | AA |
| AB | AA | AB | AA | AA | AA | AB | AA | AA | AA | AA | AB | AB | AA | AB | AB | AB | AB | AB | AA | AA | AB |
| AB | AA | AA | AA | AA | AA | AB | AA | AA | AB | AA | AB | AB | AB | AA | AB | AB | AA | AB | AB | AA | AB |
| AA | AB | AB | AB | AB | AB | AB | AB | AA | AB | AB | AB | AB | AA | AA | AB | AA | AB | AA | AA | AB | AB |
| AA | AB | AB | AA | AA | - | AB | AA | AA | AA | AA | AA | AB | AA | AB | AA | AA | AB | AB | AA | - | - |
| AA | AB | AB | AA | AA | AA | AA | AA | AA | AB | AA | AA | AA | AB | AB | AA | AA | AB | AA | AA | AB | AA |
| AB | AA | AB | AB | AB | AB | AB | AB | AB | AA | AB | AA | AB | AA | AB | AB | AB | AA | AB | AB | AA | AB |
| AA | AB | AB | AB | AA | AA | AA | AB | AA | AA | AA | AA | AA | AB | AB | AB | AB | AA | AB | AA | AA | AB |
| AA | AA | AB | AB | AB | AA | AA | AB | AA | AB | AB | AA | AA | AB | AA | AA | AB | AB | AB | AA | AA | AA |
| AB | AA | AB | AB | AA | AB | AB | AA | AB | AA | AA | AB | AB | AA | AB | AA | AB | AA | AB | AA | AA | AB |
| AB | AA | AA | AB | AA | AA | AA | AB | AB | AB | AA | AA | AA | AB | AA | AB | AB | AA | AB | AB | AB | AB |
| AB | AA | AB | AA | AA | AA | AA | AA | AA | AB | AA | AA | AA | AB | AA | AA | AB | AA | AB | AA | AB | AA |
| AA | AA | AB | AB | AB | AB | AA | AB | AA | AA | AA | AA | AA | AA | AB | AA | AA | AA | AB | AB | AA | AA |
| AB | AA | AB | AA | AA | AB | AB | AA | AB | AA | AB | AB | AA | AB | AA | AB | AB | AA | AB | AA | AB | AA |
| AB | AA | AB | AB | AB | AB | AB | AB | AB | AA | AB | AB | AB | AA | AA | AB | AB | AA | AB | AB | AB | AB |
| AA | AB | AB | AA | AA | AA | AA | AA | AA | AA | AA | AA | AA | AB | AA | AA | AA | AA | AB | AA | AB | AA |
| AA | AB | AA | AB | AB | AB | AA | AA | - | AA | AB | - | AA | - | AB | - | AA | AB | AA | AA | - | - |
| AB | AB | AA | AB | AB | AB | AB | AB | AB | AB | AB | AB | AB | AB | AB | AB | AB | AB | AB | AB | AB | AB |
| AA | AB | AA | AB | AB | - | AA | AA | AA | AB | AA | AA | AA | AB | AB | AA | AB | AB | AB | AB | AB | - |
| AB | AB | AA | AB | AB | AB | AB | AB | AB | AB | AB | AB | AB | AB | AB | AB | AB | AB | AB | AB | AB | AB |
| AA | AA | AA | AB | AB | AB | AB | AA | AB | AA | AA | AB | AB | AA | AA | AB | AB | AB | AB | AA | AA | AB |
| AA | AB | AA | AA | AA | - | - | AA | AA | - | AA | - | AA | AB | AB | AB | AB | AB | AA | AB | - | AB |
| AA | AA | AA | AB | AB | AB | AB | AB | AB | AB | AB | AB | AB | AB | AB | AB | AA | AA | AA | AB | AB | AB |
| AB | AA | AA | AB | AA | AA | AA | AA | AB | AA | AB | AA | AB | AB | AA | AB | AA | AB | AA | AB | AA | AB |
| AA | AB | AB | AB | AA | AA | AA | AB | AB | AA | AB | AA | AA | AB | AB | AA | AB | AB | AB | AA | AA | AA |
| AB | AA | AB | AA | AB | AA | AB | AA | AB | AA | AA | AA | AB | AA | AB | AB | AB | AA | AB | AB | AB | AA |
| AB | AB | AB | AA | AB | AA | AB | AA | AB | AA | AA | AA | AB | AA | AA | AB | AB | AA | AB | AB | AB | AA |
| AB | AA | AA | AA | AA | AB | AB | AA | AA | - | AA | AA | AB | AA | AA | AB | AB | AA | AB | AA | AA | AB |
| AB | AB | AA | AA | AA | - | - | AA | AA | - | AB | - | AB | AA | AB | AB | AA | AB | - | - | AA | AA |
| AA | AB | AB | AB | AB | AA | AA | AA | AA | AA | AA | AA | AA | AB | AA | AA | AB | AA | AA | AB | AB | AA |
| AA | AA | AA | AB | AA | AA | AA | AA | AA | AB | AA | AA | AA | AA | AA | AA | AB | AA | AA | AB | AA | AA |
| AB | AB | AB | AB | AB | AB | AB | AB | AB | AA | - | AB | AB | AA | AB | AB | AB | AA | AB | - | - | AB |
| AA | AB | AB | AA | AA | - | AA | AA | AA | - | AA | AA | AA | AB | AB | AA | AB | AB | AB | AA | AB | AA |
| AB | AA | AB | AA | AB | AB | AB | AA | AA | AA | AA | - | AB | AA | AA | AB | AB | AA | AB | AA | - | AB |
| AB | AB | AA | AB | AB | AB | AB | AB | AB | - | AB | AB | AB | AB | AB | AB | AB | AB | AA | AB | AB | AB |
| AB | AA | AB | AB | AB | AB | AB | AB | AB | AB | AB | AB | AB | AB | AA | AB | AB | AB | AA | AB | AA | AB |
| AB | AA | AB | AB | AB | - | AB | AB | AB | AB | AB | AB | AB | AB | AB | AB | AB | AB | AB | AB | AB | AB |
| AA | AB | AA | AA | AA | AA | AA | AA | AA | AA | AA | AA | AA | AA | AB | AA | AA | AB | AA | AA | AA | AA |
| AB | AA | AB | AA | AB | - | AB | AB | AB | AA | AB | AA | AB | AB | AB | AB | AB | AA | AA | AB | AA | AA |
| AB | AA | AA | AA | AA | AB | AB | AA | AA | AA | AA | AB | AB | AA | AA | AB | AA | AA | AB | AB | AB | AB |
| AB | AB | AB | AB | AB | AA | AB | AA | AB | AA | AA | AA | AB | AB | AA | AA | AB | AA | AB | AA | AB | AA |
| AB | AA | AB | AA | AA | AB | AB | AB | AB | AA | AB | AB | AB | AA | AA | AB | AA | AA | AB | AA | AA | AB |
| AA | AA | AB | AA | AA | AA | AA | AA | AA | AB | AA | - | AA | AB | AB | AA | AA | AB | AB | AA | AB | AB |
| AB | AB | AA | AB | AA | AA | AB | AA | AB | AA | AA | AA | AB | AA | AB | AA | AA | AB | AB | AA | AB | AB |
| AA | AB | AA | AB | AA | AB | AA | AB | AA | AB | AB | AB | AA | AB | AB | AA | AB | AB | AA | AA | AA | AB |
| AA | AA | AA | AA | AA | AB | AB | AA | AB | AA | AB | AB | AA | AB | AB | AB | AB | AB | AA | AB | AA | AA |
| AB | AB | AB | AB | AA | AA | AB | AA | AB | AA | AA | AA | AB | AB | AB | AA | AB | AB | AA | AB | AB | AB |
| AA | AA | AB | AA | AA | AA | AA | AA | AA | AA | AA | AA | AA | AA | AA | AA | AA | AB | AA | AA | AA | AA |
| AA | AB | AB | AA | AA | AA | AA | AA | AA | AA | AA | AA | AA | AA | AA | AA | AA | AA | AA | AA | AA | AA |
| AA | AA | AB | AB | AB | AB | AB | AB | AB | AB | AB | AB | AB | AB | AB | AB | AA | AB | AB | AB | AB | AB |
| AB | AB | AB | AA | AA | AB | AB | AA | AB | - | AA | AB | AB | AA | AB | AA | AA | AA | AA | AA | - | AB |
| AB | AB | AB | AB | AB | AB | AB | AB | AB | AB | AB | AB | AB | AB | AB | AB | AB | AB | AB | AB | AB | AB |
| AB | AB | AB | AB | AB | AB | AB | AB | AB | AB | AB | AB | AB | AB | AB | AB | AB | AB | AB | AB | AB | AB |
| [illegible] | [illegible] | [illegible] | [illegible] | [illegible] | [illegible] | [illegible] | [illegible] | [illegible] | [illegible] | [illegible] | [illegible] | [illegible] | [illegible] | [illegible] | [illegible] | [illegible] | [illegible] | [illegible] | [illegible] | [illegible] | [illegible] |

[illegible]

| snp_B4405 6 | snp_B4420 6 | snp_B4422 6 | snp_B4425 6 | snp_B4427 6 | snp_B4430 6 | snp_B4442 6 | snp_B4445 6 | snp_B4451 6 | snp_B4455 6 | snp_B4466 6 | snp_B4474 6 | snp_B4514 6 | snp_B4526 6 | snp_B4535 6 | snp_B4541 6 | snp_B4554 6 | snp_B4564 6 | snp_B4571 6 | snp_B4661 6 | snp_B4743 6 | snp_B4752 6 |
|---|---|---|---|---|---|---|---|---|---|---|---|---|---|---|---|---|---|---|---|---|---|
| -- | -- | -- | -- | -- | -- | -- | LG8 | -- | -- | -- | -- | -- | -- | -- | -- | -- | -- | -- | -- | -- | -- |
| AA | AA | AA | AA | AA | AA | AB | AB | AA | AA | AA | AB | AB | AA | AA | AA | AB | AA | AA | AA | AB | AB |
| AA | AA | AA | AB | AA | AA | AA | AB | AA | AA | AA | AB | AA | AA | AA | AA | AA | AA | AA | AA | AB | AB |
| AB | AB | AB | AA | AB | AB | AA | AA | AA | AB | AB | AB | AB | AA | AB | AB | AB | AB | AB | AA | AA | AA |
| AB | AB | AB | - | AB | AB | AA | - | AA | AB | AB | AA | AA | AB | - | AB | AA | AB | AB | - | AA | AB |
| AB | AB | AB | AB | AB | AA | AB | AB | AB | AA | AA | AA | AB | AB | AB | AB | AB | AA | AA | AB | AB | AB |
| AB | - | AB | AB | AB | AB | AB | - | AA | AA | AA | AA | AA | - | - | - | - | - | AA | AA | AA | AA |
| AA | AA | AA | AB | AA | AA | AB | AA | AA | AA | AB | AB | AB | AB | AA | AA | AB | AB | AA | AA | AB | AA |
| AB | AB | AB | AA | AB | AB | AA | AA | AB | AB | AB | AA | AB | AA | AB | AB | AA | AB | AB | AB | AA | AB |
| AA | AA | AA | AA | AA | AA | AB | AB | AB | AB | AB | AA | AB | AB | AB | AB | AB | AB | AB | AB | AA | AB |
| AA | AA | AA | AA | AB | AA | AA | AB | AB | AA | AB | AB | AA | AA | AA | AA | AA | AB | AA | AB | AA | AA |
| AB | AB | AB | AA | AB | AB | AA | AA | AA | AB | AB | AB | AA | AB | AB | AB | AA | AB | AB | AB | AA | AB |
| AA | AA | AA | AB | AA | AB | AA | AB | AB | AB | AB | AB | AA | AA | AA | AB | AB | AB | AB | AB | AA | AB |
| AA | AA | AA | AB | AA | AA | AA | AB | AA | AA | AB | AA | AA | AB | AA | AA | AB | AB | AA | AA | AA | AA |
| AA | AA | AA | AA | AA | AA | AA | AB | AB | AA | AA | AA | AA | AA | AA | AA | AB | AA | AA | AA | AA | AB |
| AA | AB | AB | AB | AB | AB | AA | AA | AA | AB | AB | AA | AA | AB | AB | AB | AA | AB | AB | AB | AA | AB |
| AB | AB | AB | AA | AB | AB | AA | AB | AB | AB | AB | AB | AA | AA | AB | AB | AA | AB | AB | AB | AA | AB |
| AA | AA | AA | AB | AA | AA | AA | AA | AA | AA | AA | AB | AB | AA | AA | AA | AA | AA | AA | AA | AA | AA |
| AB | - | - | AA | AA | AA | AB | - | AA | - | AA | AA | AA | - | - | AA | - | - | AA | - | AB | AB |
| AB | AB | AB | AB | AB | AB | AB | AB | AB | AB | AB | AB | AB | AB | AB | AB | AB | AB | AB | AB | AB | AB |
| AA | AA | AA | AB | AA | AA | AB | AB | AB | AA | AB | AB | AB | AB | AA | AA | AB | AB | AA | - | AB | AA |
| AB | AB | AB | AB | AB | AB | AB | AB | AB | AB | AB | AB | AA | AA | AB | AB | AB | AB | AB | AB | AA | AB |
| AA | AA | AA | AB | AA | AA | AA | AB | AB | AA | AA | AA | AB | AA | AA | AA | AB | AA | AA | AB | AB | AB |
| - | - | - | - | AB | AA | AB | AA | AA | - | AA | AA | - | - | - | AB | - | - | AB | - | - | AA |
| AB | AB | AB | AB | AB | AB | AA | AB | AB | AB | AA | AA | AB | AA | AB | AA | AB | AA | AB | AB | AB | AB |
| AA | AA | AA | AB | AA | AA | AA | AA | AA | AB | AB | AB | AA | AB | AB | AA | AB | AB | AA | AA | AA | AA |
| AA | AA | AA | AB | AA | AA | AB | AA | AB | AA | AB | AA | AB | AB | AB | AA | AB | AB | AA | AB | AB | AB |
| AB | AB | AB | AA | AB | AB | AA | AA | AA | AB | AB | AB | AB | AB | AB | AA | AA | AB | AB | AA | AA | AA |
| AB | AB | AB | AB | AB | AB | AB | AA | AB | AB | AB | AB | AA | AB | AB | AB | AB | AB | AB | AA | AA | AA |
| AA | AA | AA | AB | AA | AA | AA | AA | AB | AB | AB | AB | AA | AB | AB | AB | AA | AB | AB | AA | AA | AA |
| - | AB | AB | AA | AA | AB | AB | AA | AA | AB | AB | AA | AB | AB | AB | - | AB | - | AB | AA | - | AA |
| AA | AA | AA | AB | AA | AA | AA | AB | AB | AA | AB | AB | AB | AA | AA | AA | AA | AB | AA | AA | AA | AA |
| AA | AA | AA | AA | AA | AA | AB | AB | AB | AA | AA | AA | AA | AB | AA | AA | AB | AA | AA | AA | AB | AA |
| AB | AB | - | AB | AB | AB | AB | - | AB | AB | AA | AA | AA | AA | AB | AB | AA | - | AB | - | AA | AB |
| AA | AA | AA | AB | AA | AA | AB | AA | AA | AA | AA | AA | AA | AA | AA | AA | - | AA | AA | - | AA | AA |
| - | - | AB | AA | - | AB | AA | - | AA | AB | AB | AB | AA | - | AA | AB | - | - | AA | AA | - | AA |
| AB | AB | AB | AB | AB | AB | AB | - | AB | AB | AB | AB | AB | AB | AB | AB | AB | AB | AB | - | AB | AB |
| AB | AB | AB | AA | AB | AB | AB | AB | AB | AB | AB | AB | AB | AB | AB | AB | AB | AB | AB | AB | AB | AB |
| AB | AB | AB | AB | AB | AB | AB | AB | AB | AB | AB | AB | AB | AB | AB | AB | AB | AB | AB | AB | AB | AB |
| AA | AA | AA | AA | AA | AA | AB | AA | AA | AA | AA | AA | AB | AA | AA | AA | AA | AB | AB | AA | AA | AA |
| AA | AB | AA | AA | AA | AB | AA | - | AB | AB | AB | AB | AA | AA | AB | AB | - | - | AB | AB | - | AB |
| AB | AB | AB | AA | AB | AB | AA | AB | AA | AB | AB | AB | AA | AA | AA | AB | AA | AB | AA | AA | AA | AA |
| AA | AA | AA | AB | AA | AA | AA | AA | AB | AB | AB | AB | AB | AA | AA | AA | AA | AB | AA | AB | AA | AB |
| AB | AB | AB | AA | AB | AB | AA | AA | AA | AB | AA | AA | AA | AA | AB | AB | AA | AA | AB | AB | AA | AB |
| AA | AA | AA | AB | AA | AA | AB | AA | AA | AA | AA | AB | AB | AB | AA | AA | AB | AA | AA | AA | AB | AA |
| AA | AB | AA | AB | AA | AA | AB | AB | AA | AB | AA | AB | AB | AB | AA | AA | AB | AA | AA | AA | AB | AA |
| AA | AA | AA | AB | AA | AA | AA | AB | AB | AA | AA | AA | AA | AA | AA | AB | AB | AA | AA | AB | AB | AB |
| AB | AB | AB | AB | AA | AA | AB | AB | AA | AB | AA | AA | AB | AB | AA | AB | AB | AA | AB | AB | AB | AB |
| AB | AB | AB | AB | AB | AB | AB | AA | AA | AB | AA | AA | AA | AA | AB | AB | AA | AA | AB | AA | AA | AA |
| AA | AA | AA | AA | AA | AA | AA | AA | AA | AA | AA | AA | AA | AA | AA | AA | AA | AA | AA | AA | AA | AA |
| AA | AA | AA | AA | AA | AA | AB | AA | AA | AA | AA | AA | AA | AA | AA | AA | AA | AA | AA | AA | AA | AA |
| AB | AB | AB | AB | AB | AB | AA | AA | AB | AB | AA | AA | AA | AB | AB | AB | AA | AB | AB | AB | AA | AB |
| AB | AB | AB | AB | - | AB | AB | - | AA | AB | AB | AB | AA | AA | - | AB | AA | - | AB | AA | AA | AA |
| AB | AB | AB | AB | AB | AB | AB | AB | AB | AB | AB | AB | AB | AB | AB | AB | AB | AB | AB | AB | AB | AB |

| snp_B5552 | snp_B5555 | snp_B5556 | snp_B5563 | snp_B5565 | snp_B5578 | snp_B5579 | snp_B5632 | snp_B5656 | snp_B5663 | snp_B5676 | snp_B5682 | snp_B5708 | snp_B5742 | snp_B5768 | snp_B5784 | snp_B5789 | snp_B5795 | snp_B5797 | snp_B5798 | snp_B5805 |
|---|---|---|---|---|---|---|---|---|---|---|---|---|---|---|---|---|---|---|---|---|
| 6 | 6 | 6 | 6 | 6 | 6 | 6 | 6 | 6 | 6 | 6 | 6 | 6 | 6 | 6 | 6 | 6 | 6 | 6 | 6 | 6 |
| -- | -- | -- | -- | -- | -- | -- | -- | -- | LG8 | -- | -- | -- | -- | LG8 | -- | -- | -- | -- | -- | -- |

| snp_B5837 | snp_B5840 | snp_B5841 | snp_B5843 | snp_B5869 | snp_B5912 | snp_B5925 | snp_B5931 | snp_B5940 | snp_B5957 | snp_B5983 | snp_B5987 | snp_B6154 | snp_B6159 | snp_B6250 | snp_B6364 | snp_B0003 | snp_B0034 | snp_B0039 |
|---|---|---|---|---|---|---|---|---|---|---|---|---|---|---|---|---|---|---|
| 6 | 6 | 6 | 6 | 6 | 6 | 6 | 6 | 6 | 6 | 6 | 6 | 6 | 6 | 6 | 6 | 7 | 7 | 7 |
| -- | -- | -- | -- | -- | -- | -- | -- | -- | LG8 | -- | -- | LG8 | -- | -- | -- | LG5 | -- | -- |

| snp_B0119 | snp_B0125 | snp_B0148 | snp_B0166 | snp_B0171 | snp_B0201 | snp_B0204 | snp_B0210 | snp_B0224 | snp_B0248 | snp_B0283 | snp_B0285 | snp_B0307 | snp_B0310 | snp_B0320 | snp_B0326 | snp_B0336 | snp_B0352 | snp_B0359 |
|---|---|---|---|---|---|---|---|---|---|---|---|---|---|---|---|---|---|---|
| 7 | 7 | 7 | 7 | 7 | 7 | 7 | 7 | 7 | 7 | 7 | 7 | 7 | 7 | 7 | 7 | 7 | 7 | 7 |
| -- | -- | -- | -- | -- | -- | -- | -- | -- | -- | -- | -- | LG5 | -- | -- | -- | -- | -- | -- |

| snp_B0384 | snp_B0412 | snp_B0443 | snp_B0448 | snp_B0455 | snp_B0490 | snp_B0494 | snp_B0548 | snp_B0576 | snp_B0578 | snp_B0645 | snp_B0653 | snp_B0657 | snp_B0659 | snp_B0721 | snp_B0732 | snp_B0745 | snp_B0783 |
|---|---|---|---|---|---|---|---|---|---|---|---|---|---|---|---|---|---|
| 7 | 7 | 7 | 7 | 7 | 7 | 7 | 7 | 7 | 7 | 7 | 7 | 7 | 7 | 7 | 7 | 7 | 7 |
| -- | -- | -- | -- | -- | -- | -- | -- | -- | LG5 | -- | -- | -- | LG5 | -- | -- | -- | -- |

| snp_B0804 | snp_B0814 | snp_B0855 | snp_B0856 | snp_B0889 | snp_B0901 | snp_B0908 | snp_B0920 | snp_B0938 | snp_B0946 | snp_B0958 | snp_B0968 | snp_B0991 | snp_B1044 | snp_B1047 | snp_B1055 | snp_B1057 | snp_B1060 | snp_B1069 | snp_B1076 | snp_B1085 | snp_B1094 | snp_B1097 | snp_B1108 | snp_B1129 | snp_B1133 | snp_B1136 | snp_B1150 | snp_B1153 | snp_B1167 | snp_B1200 | snp_B1228 | snp_B1231 | snp_B1244 | snp_B1253 | snp_B1273 | snp_B1308 | snp_B1316 | snp_B1332 | snp_B1374 | snp_B1382 | snp_B1390 | snp_B1407 | snp_B1416 | snp_B1419 | snp_B1428 | snp_B1432 | snp_B1452 | snp_B1455 | snp_B1457 | snp_B1490 | snp_B1523 | snp_B1528 | snp_B1542 | snp_B1547 | snp_B1548 | snp_B1553 | snp_B1564 | snp_B1567 | snp_B1596 | snp_B1598 | snp_B1603 | snp_B1629 | snp_B1634 | snp_B1650 | snp_B1652 | snp_B1664 | snp_B1665 | snp_B1691 | snp_B1704 | snp_B1716 | snp_B1720 | snp_B1722 | snp_B1740 | snp_B1754 | snp_B1772 | snp_B1781 |
|---|---|---|---|---|---|---|---|---|---|---|---|---|---|---|---|---|---|---|---|---|---|---|---|---|---|---|---|---|---|---|---|---|---|---|---|---|---|---|---|---|---|---|---|---|---|---|---|---|---|---|---|---|---|---|---|---|---|---|---|---|---|---|---|---|---|---|---|---|---|---|---|---|---|---|---|---|
| 7 | 7 | 7 | 7 | 7 | 7 | 7 | 7 | 7 | 7 | 7 | 7 | 7 | 7 | 7 | 7 | 7 | 7 | 7 | 7 | 7 | 7 | 7 | 7 | 7 | 7 | 7 | 7 | 7 | 7 | 7 | 7 | 7 | 7 | 7 | 7 | 7 | 7 | 7 | 7 | 7 | 7 | 7 | 7 | 7 | 7 | 7 | 7 | 7 | 7 | 7 | 7 | 7 | 7 | 7 | 7 | 7 | 7 | 7 | 7 | 7 | 7 | 7 | 7 | 7 | 7 | 7 | 7 | 7 | 7 | 7 | 7 | 7 | 7 | 7 | 7 | 7 |
| -- | -- | -- | -- | -- | -- | -- | -- | -- | -- | -- | -- | -- | -- | -- | -- | -- | -- | -- | -- | -- | -- | -- | -- | -- | -- | -- | -- | -- | LG5 | -- | LG5 | -- | LG5 | -- | -- | LG5 | -- | -- | -- | -- | LG5 | LG5 | -- | -- | -- | -- | -- | -- | -- | -- | LG5 | -- | -- | -- | -- | -- | -- | LG5 | -- | -- | -- | -- | -- | -- | -- | -- | -- | -- | -- | -- | -- | -- | -- | -- | -- | -- |

| snp_B1786 | snp_B1791 | snp_B1793 | snp_B1838 | snp_B1840 | snp_B1866 | snp_B1880 | snp_B1913 | snp_B1933 | snp_B1943 | snp_B1987 | snp_B1989 | snp_B1994 | snp_B2050 | snp_B2058 | snp_B2072 | snp_B2095 | snp_B2102 | snp_B2110 | snp_B2115 | snp_B2124 | snp_B2135 | snp_B2178 | snp_B2209 | snp_B2210 | snp_B2216 | snp_B2227 | snp_B2250 | snp_B2253 | snp_B2257 | snp_B2273 | snp_B2279 | snp_B2304 | snp_B2313 | snp_B2398 | snp_B2410 | snp_B2429 | snp_B2442 | snp_B2454 | snp_B2485 | snp_B2488 | snp_B2493 | snp_B2527 | snp_B2545 | snp_B2560 | snp_B2567 | snp_B2574 | snp_B2582 | snp_B2639 | snp_B2672 | snp_B2716 | snp_B2749 | snp_B2756 | snp_B2789 | snp_B2801 | snp_B2812 | snp_B2839 | snp_B2848 | snp_B2860 | snp_B2863 | snp_B2865 | snp_B2920 | snp_B2943 | snp_B2956 | snp_B2968 | snp_B2985 | snp_B3002 | snp_B3008 | snp_B3019 | snp_B3026 | snp_B3041 | snp_B3050 | snp_B3057 | snp_B3105 | snp_B3131 | snp_B3147 | snp_B3151 |
|---|---|---|---|---|---|---|---|---|---|---|---|---|---|---|---|---|---|---|---|---|---|---|---|---|---|---|---|---|---|---|---|---|---|---|---|---|---|---|---|---|---|---|---|---|---|---|---|---|---|---|---|---|---|---|---|---|---|---|---|---|---|---|---|---|---|---|---|---|---|---|---|---|---|---|---|---|
| 7 | 7 | 7 | 7 | 7 | 7 | 7 | 7 | 7 | 7 | 7 | 7 | 7 | 7 | 7 | 7 | 7 | 7 | 7 | 7 | 7 | 7 | 7 | 7 | 7 | 7 | 7 | 7 | 7 | 7 | 7 | 7 | 7 | 7 | 7 | 7 | 7 | 7 | 7 | 7 | 7 | 7 | 7 | 7 | 7 | 7 | 7 | 7 | 7 | 7 | 7 | 7 | 7 | 7 | 7 | 7 | 7 | 7 | 7 | 7 | 7 | 7 | 7 | 7 | 7 | 7 | 7 | 7 | 7 | 7 | 7 | 7 | 7 | 7 | 7 | 7 | 7 |
| -- | -- | -- | -- | -- | -- | LG5 | -- | -- | -- | -- | LG5 | -- | LG5 | -- | -- | -- | LG5 | LG5 | -- | -- | -- | -- | -- | LG5 | LG5 | -- | -- | -- | -- | -- | -- | -- | LG5 | -- | -- | -- | -- | LG5 | -- | LG5 | -- | LG5 | LG5 | -- | -- | -- | LG5 | -- | -- | -- | -- | -- | -- | -- | -- | -- | -- | -- | -- | -- | -- | -- | -- | LG5 | -- | -- | -- | -- | -- | LG5 | -- | LG5 | -- | LG5 | -- | -- |

| snp_B1786 | snp_B1791 | snp_B1793 | snp_B1838 | snp_B1840 | snp_B1866 | snp_B1880 | snp_B1913 | snp_B1933 | snp_B1943 | snp_B1987 | snp_B1989 | snp_B1994 | snp_B2050 | snp_B2058 | snp_B2072 | snp_B2095 | snp_B2102 | snp_B2110 |
|---|---|---|---|---|---|---|---|---|---|---|---|---|---|---|---|---|---|---|
| AB | AB | AB | AB | AA | AA | AA | AB | AA | AA | AA | AA | AB | AB | AB | AA | AA | AB | AB |
| AB | AB | AB | AA | AA | AB | AB | AB | AB | AB | AA | AB | AB | AB | AB | AB | AB | AB | AB |
| AA | AA | AA | AA | AB | AB | AB | AA | AB | AB | AB | AB | AB | AA | AA | AB | AB | AB | AB |
| - | AB | AB | - | AA | AB | AB | - | - | AB | - | AB | AB | - | AB | AB | AB | AB | AB |
| AB | AA | AA | AB | AB | AA | AB | AA | AB | AA | AA | AB | AA | AB | AA | AA | AA | AB | AB |
| - | AB | AB | - | AA | AA | AA | - | - | - | - | AA | AB | AA | AB | AA | AB | AA | AA |
| AA | AB | AB | AA | AA | AA | AA | AA | AB | AA | AB | AA | AB | AA | AA | AA | AB | AA | AA |
| AB | AB | AB | AB | AB | AB | AB | AB | AB | AB | AA | AB | AB | AB | AB | AB | AB | AB | AB |
| - | AA | AB | AA | AA | AB | AB | AA | AB | - | AA | AB | AB | AA | AA | AB | AA | AB | AB |
| AA | AB | AB | AA | AA | AA | AA | AA | AB | AB | AA | AA | AA | AA | AA | AB | AA | AB | AA |
| AB | AA | AB | AA | AB | AB | AA | AB | AA | AA | AB | AA | AB | AA | AA | AB | AB | AB | AB |
| AB | AB | AB | AB | AB | AB | AB | AB | AB | AA | AA | AB | AB | AB | AB | AB | AB | AB | AB |
| AA | AB | AB | AA | AA | AB | AB | AB | AB | AB | AA | AA | AB | AA | AA | AB | AB | AB | AB |
| AA | AB | AB | AA | AA | AA | AA | AB | AA | AA | AB | AA | AA | AA | AA | AA | AA | AA | AA |
| AA | AB | AA | AA | AA | AA | AA | AB | AB | AB | AA | AB | AA | AB | AB | AA | AA | AA | AA |
| AB | AB | AB | AB | AB | AB | AB | AA | AB | AB | AB | AB | AB | AB | AB | AB | AB | AB | AB |
| AA | AB | AB | AA | AA | AA | AA | AB | AA | AB | AA | AA | AB | AA | AB | AA | AA | AA | AA |
| - | AA | AB | - | AB | AB | - | AA | - | - | - | AB | AB | AB | - | AB | AA | AB | AB |
| AB | AB | AB | AB | AB | AB | AB | AB | AB | AB | AB | AB | AB | AB | AB | AB | AB | AB | AB |
| AA | AA | AA | AA | AA | AA | AA | AA | AB | AA | AB | AA | AA | AA | - | - | AA | AA | AA |
| AA | AB | AB | AA | AA | AA | AA | AB | AB | AB | AB | AA | AB | AA | AB | AB | AB | AA | AA |
| AA | AA | AA | AA | AA | AA | AA | AA | AB | AA | AB | AA | AA | AA | AA | AA | AA | AA | AA |
| - | AA | AA | - | - | AB | AB | - | - | - | - | AB | AB | - | - | - | - | AB | AA |
| AA | AB | AB | AA | AA | AA | AA | AB | AB | AA | AB | AA | AB | AA | AB | AA | AB | AA | AA |
| AB | AB | AB | AB | AB | AB | AB | AB | AB | AB | AA | AB | AB | AB | AB | AB | AB | AB | AB |
| AB | AA | AA | AB | AA | AA | AA | AB | AA | AA | AB | AA | AA | AB | AB | AA | AA | AA | AA |
| AA | AA | AA | AA | AA | AA | AA | AA | AA | AA | AA | AA | AA | AA | AA | AA | AA | AA | AA |
| AA | AB | AB | AA | AA | AB | AB | AA | AB | AB | AA | AB | AA | AB | AA | AB | AA | AB | AB |
| AB | AB | AA | AB | AA | AB | AA | AB | - | AA | AB | AA | AB | AB | AA | AB | AB | AA | AA |
| - | AA | - | - | AB | AB | AB | - | - | - | - | AB | AA | - | - | AB | - | AB | AB |
| AA | AB | AB | AA | AA | AA | AA | AB | AB | AA | AB | AA | AB | AA | AB | AA | AB | AA | AA |
| AB | AA | AA | AB | AB | AA | AB | AA | AA | AB | AA | AB | AA | AB | AA | AB | AA | AB | AB |
| - | AA | AA | AB | AB | AB | AB | AA | AB | - | - | AB | AA | - | - | - | - | AB | - |
| AB | AB | AB | - | AA | AA | AA | - | - | AA | - | AB | - | AB | AA | AB | AB | AA | AA |
| - | AA | AA | AA | AB | AA | AA | AA | AA | - | AB | AA | AA | AB | AB | AA | AA | AA | AA |
| - | AB | AB | AB | AB | AB | AB | AB | AB | AB | AB | AB | AB | AB | AB | AB | AB | AB | AB |
| AA | AB | AB | AA | AA | AB | AB | AA | AB | AB | AB | AA | AB | AA | AA | AB | AB | AA | AA |
| AA | AB | AB | AA | AA | AB | AA | AB | AA | AA | AA | AA | AB | AA | AB | AA | AB | AA | AA |
| AA | AA | AB | AA | AA | AB | AB | AB | AA | AB | AB | AB | AB | AA | AA | AB | AA | AB | AB |
| - | AA | AA | AB | AB | AB | AB | - | AA | AA | AA | AB | AA | - | AA | AB | AA | AB | AB |
| AB | AB | AB | AB | AB | AB | AB | AB | AB | AB | AA | AB | AB | AB | AB | AB | AB | AB | AB |
| AB | AA | AA | AB | AB | AA | AA | AA | AA | AA | AA | AB | AA | AB | AB | AA | AA | AB | AB |
| AB | AA | AA | AB | AB | AB | AB | AB | AB | AA | AB | AB | AB | AB | AB | AA | AA | AB | AB |
| AB | AA | AA | AB | AB | AB | AB | AA | AA | AB | - | AB | AB | AB | - | - | AA | AB | AB |
| AB | AA | AB | AB | AA | AA | AB | AA | AA | AA | AB | AA | AA | AB | AB | AB | AA | AB | AA |
| AA | AA | AA | AA | AA | AA | AA | AA | AA | AA | AB | AA | AA | AB | AB | AA | AB | AA | AA |
| AA | AB | AB | AA | AB | AB | AB | AA | AA | AB | AB | AB | AB | AA | AA | AB | AB | AB | AB |
| AB | AA | AB | AB | AB | AB | AB | AA | AA | AB | AA | AB | AB | AB | AB | AB | AB | AB | AB |
| AB | AB | AB | AB | AB | AB | AB | AA | AB | AB | AA | AB | AB | AB | AB | AB | AB | AB | AB |
| AA | AB | AB | AA | AB | AB | AB | AA | AB | AB | AA | AB | AB | AA | AA | AB | AB | AB | AB |

| snp_B3174 | snp_B3177 | snp_B3205 | snp_B3210 | snp_B3219 | snp_B3229 | snp_B3234 | snp_B3260 | snp_B3284 | snp_B3342 | snp_B3362 | snp_B3405 | snp_B3435 | snp_B3444 | snp_B3446 | snp_B3472 | snp_B3515 | snp_B3543 | snp_B3544 | snp_B3589 | snp_B3591 | snp_B3599 | snp_B3611 | snp_B3616 | snp_B3682 | snp_B3683 | snp_B3687 | snp_B3711 | snp_B3723 | snp_B3724 | snp_B3730 | snp_B3737 | snp_B3749 | snp_B3750 | snp_B3763 | snp_B3791 | snp_B3798 | snp_B3802 | snp_B3847 | snp_B3878 | snp_B3897 | snp_B3902 | snp_B3903 | snp_B3945 | snp_B3947 | snp_B3961 | snp_B3963 | snp_B3964 | snp_B3991 | snp_B4008 | snp_B4020 | snp_B4028 | snp_B4041 | snp_B4096 | snp_B4141 | snp_B4149 | snp_B4168 | snp_B4184 | snp_B4195 | snp_B4204 | snp_B4210 | snp_B4223 | snp_B4226 | snp_B4229 | snp_B4256 | snp_B4259 | snp_B4267 | snp_B4271 | snp_B4277 | snp_B4292 | snp_B4306 | snp_B4307 | snp_B4330 | snp_B4332 | snp_B4349 | snp_B4366 | snp_B4377 |
|---|---|---|---|---|---|---|---|---|---|---|---|---|---|---|---|---|---|---|---|---|---|---|---|---|---|---|---|---|---|---|---|---|---|---|---|---|---|---|---|---|---|---|---|---|---|---|---|---|---|---|---|---|---|---|---|---|---|---|---|---|---|---|---|---|---|---|---|---|---|---|---|---|---|---|---|---|
| 7 | 7 | 7 | 7 | 7 | 7 | 7 | 7 | 7 | 7 | 7 | 7 | 7 | 7 | 7 | 7 | 7 | 7 | 7 | 7 | 7 | 7 | 7 | 7 | 7 | 7 | 7 | 7 | 7 | 7 | 7 | 7 | 7 | 7 | 7 | 7 | 7 | 7 | 7 | 7 | 7 | 7 | 7 | 7 | 7 | 7 | 7 | 7 | 7 | 7 | 7 | 7 | 7 | 7 | 7 | 7 | 7 | 7 | 7 | 7 | 7 | 7 | 7 | 7 | 7 | 7 | 7 | 7 | 7 | 7 | 7 | 7 | 7 | 7 | 7 | 7 | 7 |
| -- | -- | -- | -- | -- | -- | -- | -- | -- | -- | -- | -- | -- | -- | -- | -- | -- | LG5 | -- | -- | LG5 | -- | -- | LG5 | LG5 | -- | -- | LG5 | -- | -- | -- | -- | -- | -- | LG5 | -- | LG5 | -- | -- | -- | -- | -- | -- | -- | -- | -- | -- | LG5 | -- | -- | -- | -- | -- | -- | -- | -- | -- | -- | -- | LG5 | -- | -- | LG5 | -- | -- | -- | -- | -- | LG5 | LG5 | -- | -- | -- | -- | -- | -- | -- |

| snp_B43827 | snp_B44077 | snp_B44247 | snp_B44447 | snp_B44687 | snp_B44707 | snp_B44737 | snp_B44877 | snp_B44957 | snp_B44997 | snp_B45437 | snp_B45467 | snp_B45817 | snp_B45927 | snp_B46207 | snp_B46217 | snp_B46387 | snp_B46507 | snp_B46517 | snp_B46747 |
|---|---|---|---|---|---|---|---|---|---|---|---|---|---|---|---|---|---|---|---|
| -- | -- | -- | -- | -- | -- | -- | -- | -- | -- | -- | -- | -- | -- | -- | -- | -- | -- | -- | -- |
| AB | AA | AB | AA | AA | AB | AB | AA | AA | AB | AA | AB | AB | AA | AB | AB | AA | AA | AB | AB |
| AA | AB | AA | AB | AB | AA | AB | AA | AA | AB | AA | AB | AA | AB | AA | AB | AA | AA | AA | AB |
| AB | AB | AA | AB | AB | AA | AB | AB | AB | AB | AA | AA | AA | AB | AA | AA | AA | AA | AA | AA |
| AB | AB | AA | AB | - | AA | AB | AA | AA | AB | AA | - | AB | AB | AA | AB | AB | AA | AA | - |
| AA | AA | AB | AB | AB | AB | AA | AB | AB | AA | AB | AA | AA | AA | AB | AA | AB | AB | AB | AA |
| - | AB | AA | AA | - | AA | AB | - | - | AB | AB | - | - | AB | AA | AB | AB | - | AA | - |
| AB | AB | AA | AA | AA | AA | AB | AA | AA | AB | AB | AA | AB | AB | AA | AB | AB | AA | AA | AA |
| AA | AB | AB | AB | AB | AB | AA | AB | AB | AB | AA | AB | AA | AB | AB | AB | AB | AA | AB | AB |
| AB | AA | AB | AB | AA | AB | AB | AB | AB | AB | AA | AA | AB | AA | AA | AB | AB | AA | AA | AB |
| AB | AA | AA | AB | AB | AA | AB | AB | AA | AA | AA | AB | AB | AA | AA | AB | AA | AA | AA | AB |
| AA | AB | AA | AB | AB | AB | AA | AB | AB | AA | AB | AA | AA | AB | AA | AB | AB | AA | AA | AA |
| AA | AB | AB | AB | AB | AB | AA | AB | AB | AB | AA | AB | AA | AB | AB | AB | AB | AA | AB | AB |
| AA | AB | AA | AB | AB | AB | AA | AA | AB | AB | AA | AB | AB | AB | AA | AB | AB | AA | AA | AB |
| AB | AA | AA | AA | AB | AB | AA | AB | AA | AB | AB | AB | AB | AA | AB | AB | AB | AA | AA | AB |
| AA | AB | AB | AB | AA | AA | AA | AA | AA | AA | AA | AB | AA | AB | AA | AA | AA | AB | AB | AB |
| AB | AB | AB | AB | AB | AB | AB | AB | AB | AB | AB | AB | AB | AB | AB | AB | AB | AB | AB | AA |
| AA | AA | AB | AB | AA | AB | AA | AA | AA | AB | AA | AB | AB | AA | AA | AB | AA | AB | AB | AB |
| - | AA | AB | AA | - | AB | AA | - | - | AB | AB | - | AB | AA | - | - | AB | AB | AB | - |
| AB | AB | AB | AB | AB | AB | AB | AB | AB | AB | AB | AB | AB | AB | AB | AB | AB | AB | AB | AB |
| AB | AB | AB | AA | AB | AA | AB | AB | AB | AA | AB | AA | AB | AB | AA | AA | AA | AA | AB | AB |
| AB | AB | AA | AB | AB | AA | AB | AB | AA | AA | AB | AB | AB | AB | AA | AA | AB | AA | AB | AB |
| AB | AB | AA | AA | AA | AA | AB | AA | AA | AA | AB | AA | AB | AB | AA | AA | AA | AA | AA | AA |
| - | AA | AB | AA | - | AB | AA | - | AB | AB | - | - | - | - | AB | AA | AB | - | AB | - |
| AB | AB | AA | AA | AA | AA | AB | AA | AA | AB | AB | AB | AB | AB | AA | AB | AB | AA | AA | AA |
| AB | AB | AB | AB | AB | AA | AB | AB | AB | AB | AA | AB | AB | AB | AB | AB | AB | AB | AA | AB |
| AB | AA | AA | AA | AA | AA | AB | AA | AA | AA | AB | AB | AB | AA | AB | AA | AB | AA | AA | AB |
| AB | AA | AA | AA | AA | AA | AB | AA | AA | AA | AB | AA | AB | AA | AA | AA | AA | AA | AA | AA |
| AA | AA | AA | AB | AA | AA | AA | AA | AA | AB | AB | AA | AB | AA | AA | AB | AB | AB | AA | AB |
| AB | AA | AA | AA | AA | AA | AB | - | - | AB | AA | AB | - | AA | AB | AA | AB | - | AB | AB |
| AA | - | AB | AB | - | AA | AA | - | AB | AB | AB | - | - | AA | AA | AB | AA | - | AB | - |
| AB | AB | AA | AA | AB | AA | AB | AB | AB | AB | AB | AB | AB | AB | AA | AB | AB | AA | AA | AA |
| AA | AA | AA | AB | AB | AB | AA | AB | AB | AA | AA | AA | AB | AA | AB | AA | AA | AB | AA | AB |
| AA | AA | AB | AB | AB | AB | AA | - | AB | AA | AA | AA | AB | AA | AB | AA | AA | - | AB | AB |
| AB | AB | AA | AA | - | AA | AB | AA | AA | AB | AA | - | AB | AB | AA | AB | AB | AA | AA | - |
| AB | AA | AB | AA | - | AB | AB | - | AB | AA | AB | - | AB | AA | AA | - | AA | AA | AB | - |
| AB | AB | AB | AB | AB | AB | AB | AB | AB | AB | AB | AB | AB | AB | AB | AB | AB | AB | AB | AB |
| AB | AB | AA | AB | AA | AB | AB | AA | AA | AB | AB | AA | AB | AB | AA | AB | AB | AB | AA | AA |
| AA | AA | AB | AA | AA | AB | AA | AA | AA | AB | AA | AB | AA | AA | AA | AB | AB | AA | AB | AB |
| AA | AA | AA | AB | AA | AB | AA | AA | AB | AB | AB | AA | AB | AA | AB | AB | AA | AB | AB | AB |
| AA | AA | AB | AB | - | AB | AA | - | AB | AA | AB | - | AA | AA | AB | - | AA | AB | AB | - |
| AA | AB | AB | AB | AB | AB | AA | AB | AB | AB | AA | AB | AA | AB | AB | AB | AA | AA | AB | AB |
| AA | AA | AA | AA | AB | AB | AA | AB | AB | AA | AA | AA | AA | AA | AB | AA | AA | AA | AA | AA |
| AB | AB | AB | AB | AB | AB | AB | AB | AB | AA | AB | AB | AB | AB | AB | AA | AA | AA | AB | AB |
| AA | AB | AA | AB | AB | AB | AA | - | AB | AB | AB | AA | AB | AB | AA | AB | AA | - | AA | AA |
| AA | AB | AA | AA | AA | AB | AA | AA | AA | AB | AB | AB | AA | AB | AB | AB | AA | AA | AB | AB |
| AB | AB | AA | AA | AA | AA | AB | AA | AA | AA | AA | AA | AA | AB | AA | AA | AA | AB | AA | AB |
| AB | AA | AB | AA | AB | AB | AB | AB | AB | AA | AA | AA | AB | AA | AB | AB | AA | AB | AB | AB |
| AA | AA | AB | AB | AB | AB | AA | AB | AB | AB | AA | AA | AA | AB | AA | AB | AB | AB | AB | AB |
| AB | AB | AA | AB | AB | AB | AB | AB | AB | AA | AB | AA | AB | AB | AA | AB | AB | AB | AA | - |
| AA | AB | AB | AB | AB | AB | AA | AB | AB | AB | AA | AA | AA | AB | AB | AB | AB | AB | AB | AA |
| AA | AB | AB | AB | AB | AB | AA | AB | AB | AB | AA | AB | AA | AB | AB | AB | AA | AB | AB | AB |
| AA | AB | AB | AB | AB | AB | AA | - | AB | AB | AB | - | AA | AB | AB | AB | AA | AB | AB | AB |
| AB | AA | AB | AA | AB | AA | AA | AA | AA | AA | AA | AB | AB | AA | AA | AA | AB | AB | AB | AB |

[illegible]

| snp_B5877 | snp_B5887 | snp_B5908 | snp_B5939 | snp_B5951 | snp_B5956 | snp_B5964 | snp_B5997 | snp_B6032 | snp_B6163 | snp_B6293 | snp_B6305 | snp_B6325 | snp_B0007 | snp_B0008 | snp_B0009 | snp_B0012 | snp_B0017 | snp_B0018 | snp_B0058 | snp_B0085 | snp_B0097 |
|---|---|---|---|---|---|---|---|---|---|---|---|---|---|---|---|---|---|---|---|---|---|
| 7 | 7 | 7 | 7 | 7 | 7 | 7 | 7 | 7 | 7 | 7 | 7 | 7 | 8 | 8 | 8 | 8 | 8 | 8 | 8 | 8 | 8 |
| -- | -- | -- | -- | -- | -- | LG5 | -- | -- | -- | -- | -- | -- | -- | -- | -- | -- | -- | -- | -- | -- | -- |
| AA | AB | AB | AB | AA | AB | AA | AB | AB | AB | AB | AA | AB | AB | AA | AB | AA | AB | AA | AA | AA | AA |
| AB | AB | AB | AB | AA | AB | AB | AB | AA | AA | AB | AA | AB | AB | AA | AB | AA | AA | AA | AA | AA | AA |
| AB | AB | AB | AA | AA | AB | AB | AA | AB | AB | AB | AA | AA | AA | AA | AB | AA | AB | AA | AA | AB | AA |
| AA | AB | AA | AA | AA | AB | AB | AB | AB | AA | AB | AB | - | AB | AB | AB | AA | - | AB | AB | AA | AA |
| AB | AA | AA | AA | AB | AB | AB | AB | AA | AA | AA | AB | AB | AB | AA | AB | AA | AB | AB | AB | AA | AA |
| AB | AB | AA | AB | AB | AB | AA | AB | - | AB | - | - | - | AB | AB | AA | - | - | AA | - | AA | - |
| AB | AB | AA | AA | AA | AA | AA | AA | AB | AB | AB | AA | AA | AA | AB | AA | AB | AA | AA | AA | AB | AA |
| AB | AB | AB | AB | AA | AB | AB | AB | AA | AA | AA | AB | AB | AA | AA | AB | AA | AA | AA | AA | AA | AB |
| AB | AA | AA | AA | AA | AA | AB | AB | AB | AB | AB | AB | AA | AA | AA | AB | AA | AA | AA | AA | AB | AB |
| AB | AB | AB | AB | AA | AA | AA | AA | AA | AB | AB | AB | AA | AA | AA | AA | AA | AB | AB | AB | AB | AA |
| AA | AA | AA | AA | AA | AB | AB | AA | AA | AA | AA | AA | AA | AB | AB | AB | AA | AA | AB | AB | AB | AA |
| AA | AA | AB | AB | AA | AB | AB | AB | AA | AA | AA | AA | AB | AA | AB | AB | AA | AA | AA | AA | AA | AA |
| AB | AB | AA | AB | AA | AA | AB | AB | AA | AA | AA | AB | AB | AA | AA | AB | AA | AB | AB | AB | AA | AB |
| AB | AB | AA | AB | AA | AB | AA | AB | AB | AB | AB | AB | AB | AB | AB | AA | AB | AB | AA | AA | AB | AB |
| AA | AA | AB | AB | AA | AA | AB | AB | AA | AA | AA | AA | AB | AB | AB | AB | AA | AB | AB | AB | AB | AA |
| AB | AB | AB | AB | AA | AB | AB | AB | AB | AB | AB | AB | AB | AB | AB | AB | AA | AB | AB | AB | AB | AB |
| AA | AA | AB | AB | AB | AB | AB | AB | AA | AA | AA | AB | AA | AB | AB | AB | AB | AB | AB | AB | AB | AB |
| AB | AA | AB | AA | AA | AA | AA | AA | - | AB | - | - | - | AB | AA | AB | AB | - | AA | - | - | AB |
| AB | AB | AB | AB | AA | AB | AB | AB | AB | AB | AB | AB | AB | AA | AA | AA | AB | - | AA | AA | AA | AA |
| AB | AB | AA | AA | AA | AA | AA | AA | AB | AB | AB | AA | AA | AB | AB | AB | AB | AB | AB | AB | AB | AB |
| AB | AB | AA | AB | AB | AB | AA | AB | AB | AB | AB | AB | AB | AA | AB | AB | AB | AB | AB | AA | AB | AB |
| AB | AB | AA | AA | AA | AA | AA | AA | AB | AB | AB | AA | AA | AA | AA | AA | AA | AA | AA | AA | AA | AA |
| AB | AA | AB | AA | AB | AB | AB | AA | - | AA | - | - | - | AA | AB | AA | - | - | AA | - | AB | - |
| AB | AB | AA | AB | AB | AB | AA | AB | AB | AB | AB | AA | AB | AA | AA | AB | AB | AB | AB | AB | AB | AB |
| AB | AB | AA | AB | AA | AB | AB | AB | AA | AB | AA | AB | AB | AA | AA | AB | AB | AB | AB | AB | AB | AB |
| AA | AB | AB | AB | AA | AA | AA | AA | AB | AB | AB | AA | AA | AA | AA | AA | AA | AB | AA | AA | AB | AA |
| AA | AA | AA | AA | AA | AA | AA | AA | AB | AB | AB | AA | AA | AB | AA | AB | AA | AA | AB | AA | AB | AB |
| AB | AA | AB | AA | AB | AB | AB | AA | AA | AA | AA | AA | AA | AA | AB | AB | AB | AB | AA | AB | AB | AB |
| AB | AB | AA | AA | AB | AA | AB | AA | AB | AB | AB | AA | AA | AB | AB | AB | AB | - | AB | AB | AB | - |
| - | AA | AB | AA | AB | - | - | AA | - | AB | - | - | - | AA | AA | AB | AA | - | AA | - | - | - |
| AB | AB | AB | AB | AA | AB | AA | AB | AB | AB | AB | AA | AA | AB | AB | AB | AB | AB | AB | AB | AB | AB |
| AA | AB | AB | AB | AA | AB | AA | AB | AA | AA | AB | AA | AB | AB | AA | AB | AB | AA | AA | AA | AB | AB |
| AA | AA | AB | AA | AA | AA | - | AA | AA | AA | AA | - | - | AB | AB | AA | AA | - | AA | AA | - | AA |
| AB | AB | AB | AB | AA | AB | - | AB | AB | AB | AB | - | - | AB | AB | AB | AB | AB | AB | - | AB | AB |
| AA | AB | AB | AB | AA | AA | AA | AA | - | AB | - | - | - | AA | AA | AB | AA | - | AA | AA | AB | AA |
| AB | AB | AB | AB | AB | AB | AB | AB | AB | AB | AB | - | AB | AB | AB | AB | AB | AB | AB | AB | AB | AB |
| AB | AB | AA | AA | AB | AA | AB | AA | AB | AB | AB | AB | AA | AA | AA | AA | AA | AA | AA | AA | AA | AA |
| AA | AA | AB | AB | AA | AB | AA | AB | AA | AA | AA | AA | AA | AB | AB | AB | AA | AB | AB | AB | AA | AA |
| AB | AA | AB | AB | AB | AA | AB | AA | AA | AB | AA | AB | AA | AB | AB | AB | AB | AB | AB | AB | AB | AA |
| AB | AA | AA | AA | AB | - | AB | AB | AA | AA | - | - | - | AA | AB | AB | AB | AA | AA | AA | - | - |
| AA | AB | AB | AB | AA | AB | AB | AB | AA | AA | AA | AB | AB | AB | AB | AA | AA | AB | AB | AB | AB | AA |
| AA | AA | AA | AA | AA | AA | AA | AB | AA | AA | AA | AA | AB | AB | AA | AA | AA | AB | AB | AB | AA | AA |
| AB | AB | AB | AB | AA | AB | AB | AB | AB | AB | AB | AB | AB | AB | AB | AA | AB | AB | AB | AB | AB | AA |
| AA | AB | AA | AA | AB | AA | AB | AB | AA | AA | AA | AB | AB | AA | AA | AB | AB | AA | AB | AB | AB | - |
| AA | AB | AA | AA | AB | AA | AB | AB | AA | AA | AA | AB | AB | AA | AA | AB | AB | AA | AA | AB | AB | AB |
| AA | AB | AB | AA | AA | AA | AA | AB | AB | AB | AB | AB | AB | AB | AB | AA | AB | AB | AB | AB | AA | AA |
| AB | AA | AB | AA | AB | AA | AB | AA | AB | AB | AB | AB | AA | AA | AA | AA | AB | AB | AA | AA | AA | AA |
| AA | AA | AB | AA | AB | AA | AB | AA | AA | AB | AA | AB | AA | AA | AA | AA | AA | AB | AB | AB | AA | AA |
| AB | AB | AA | AA | AB | AB | AB | AB | AB | AB | AB | AB | AB | AB | AA | AA | AB | AB | AB | AB | AA | AA |
| AB | AB | AA | AB | AB | AB | AB | AB | AA | AA | AA | AB | AB | AB | AB | AA | AB | AB | AB | AB | AB | AA |
| AB | AB | AB | AB | AB | AB | AB | AB | AA | AA | AA | AB | AB | AB | AA | AB | AB | AB | AB | AB | AB | AA |
| AA | AA | AA | AA | AB | AB | AB | AB | AA | AA | AA | - | - | AA | AB | AA | AA | - | AA | - | AA | AB |
| AA | AA | AB | AB | AB | AB | AB | AA | AB | AB | AA | AB | AA | AA | AA | AA | AA | AA | AA | AA | AA | AA |

[illegible]

| snp_B10728 | snp_B10738 | snp_B10938 | snp_B10998 | snp_B11048 | snp_B11078 | snp_B11448 | snp_B11458 | snp_B11608 | snp_B11878 | snp_B11908 | snp_B12138 | snp_B12328 | snp_B12468 | snp_B12528 | snp_B12578 | snp_B12598 | snp_B12628 | snp_B12648 | snp_B13128 | snp_B13158 | snp_B13698 | snp_B13788 | snp_B13968 | snp_B14058 | snp_B14308 | snp_B14368 | snp_B14378 | snp_B14388 | snp_B14408 | snp_B14468 | snp_B14648 | snp_B14708 | snp_B14878 | snp_B15018 | snp_B15298 | snp_B15328 | snp_B15458 |
| --- | --- | --- | --- | --- | --- | --- | --- | --- | --- | --- | --- | --- | --- | --- | --- | --- | --- | --- | --- | --- | --- | --- | --- | --- | --- | --- | --- | --- | --- | --- | --- | --- | --- | --- | --- | --- | --- |
| -- | -- | -- | -- | -- | -- | -- | -- | -- | -- | -- | -- | -- | -- | -- | -- | -- | -- | -- | -- | -- | -- | -- | -- | -- | -- | -- | -- | -- | LG10 | -- | -- | -- | -- | -- | -- | -- | -- |
| [illegible] | | | | | | | | | | | | | | | | | | | | | | | | | | | | | | | | | | | | | |

| snp_B15578 | snp_B15698 | snp_B15738 | snp_B15898 | snp_B15938 | snp_B15998 | snp_B16138 | snp_B16208 | snp_B16328 | snp_B16468 | snp_B17068 | snp_B17758 | snp_B17828 | snp_B18008 | snp_B18068 | snp_B18178 | snp_B18308 | snp_B18418 | snp_B18478 | snp_B18738 | snp_B18868 | snp_B19568 | snp_B19888 | snp_B20078 | snp_B20388 | snp_B20418 | snp_B20548 | snp_B20708 | snp_B20828 | snp_B20928 | snp_B21178 | snp_B21418 | snp_B21838 | snp_B21858 | snp_B21868 | snp_B21968 | snp_B22148 | snp_B22188 | snp_B22348 |
| --- | --- | --- | --- | --- | --- | --- | --- | --- | --- | --- | --- | --- | --- | --- | --- | --- | --- | --- | --- | --- | --- | --- | --- | --- | --- | --- | --- | --- | --- | --- | --- | --- | --- | --- | --- | --- | --- | --- |
| -- | -- | -- | -- | -- | -- | -- | -- | -- | -- | -- | -- | -- | -- | -- | -- | -- | -- | -- | -- | -- | -- | -- | -- | LG10 | LG10 | -- | -- | -- | -- | LG10 | -- | -- | -- | -- | -- | -- | -- | -- |
| [illegible] | | | | | | | | | | | | | | | | | | | | | | | | | | | | | | | | | | | | | | |

| snp_B22668 | snp_B22988 | snp_B23028 | snp_B23158 | snp_B23228 | snp_B23338 | snp_B23618 | snp_B23658 | snp_B23868 | snp_B23998 | snp_B24048 | snp_B24068 | snp_B24088 | snp_B24368 | snp_B24558 | snp_B24768 | snp_B24918 |
|---|---|---|---|---|---|---|---|---|---|---|---|---|---|---|---|---|
| -- | -- | -- | -- | -- | -- | -- | -- | -- | LG10 | -- | -- | -- | -- | -- | -- | -- |
| AB | AA | AB | AB | AB | AA | AB | AA | AA | AB | AA | AB | AA | AA | AA | AA | AB |
| AA | AA | AA | AB | AB | AA | AB | AA | AA | AB | AA | AB | AB | AB | AA | AB | AA |
| AA | AA | AA | AB | AB | AB | AB | AA | AA | AB | AB | AB | AB | AB | AA | AB | AB |
| AA | AB | AB | AB | AB | - | AB | AA | AB | AA | AB | AB | AB | AB | AA | AB | AB |
| AB | AB | AA | AB | AB | AA | AB | AA | AA | AB | AA | AB | AB | AA | AA | AB | AB |
| AA | - | - | AB | AA | - | AB | AA | AA | AA | - | AA | AA | - | AA | AA | AA |
| AA | AA | AA | AB | AA | AA | AA | AA | AB | AA | AA | AB | AA | AB | AA | AA | AA |
| AB | AA | AA | AB | AA | AA | AA | AA | AA | AA | AA | AB | AA | AA | AB | AA | AA |
| AB | AA | AB | AA | AA | AA | AB | AA | AB | AA | AB | AB | AA | AA | AB | AB | AA |
| AB | AA | AA | AB | AB | AA | AB | AA | AB | AB | AA | AB | AA | AA | AB | AB | AA |
| AB | AB | AB | AB | AA | AA | AB | AB | AA | AA | AB | AB | AA | AA | AA | AA | AB |
| AA | AA | AB | AB | AA | AA | AA | AA | AA | AB | AA | AA | AA | AA | AA | AA | AA |
| AA | AB | AB | AB | AA | AB | AA | AA | AB | AA | AA | AA | AA | AA | AB | AA | AB |
| AA | AB | AB | AA | AA | AB | AB | AA | AB | AB | AA | AA | AA | AA | AB | AB | AA |
| AB | AB | AB | AA | AB | AA | AB | AA | AA | AA | AB | AB | AA | AA | AA | AB | AB |
| AA | AB | AB | AB | AB | AA | AB | AA | AB | AA | AB | AA | AB | AB | AA | AB | AB |
| AA | AB | AB | AB | AB | AB | AB | AB | AB | AA | AB | AA | AB | AB | AB | AB | AB |
| AA | - | AA | AA | AB | - | - | AB | - | AA | - | AA | AA | AA | - | AB | - |
| AA | AA | AA | AA | AA | AA | AA | AA | AA | AA | AA | AA | AA | AA | AA | AA | AA |
| AB | AB | AA | AA | AB | AA | AA | AA | AB | AA | AA | AA | AB | AB | AB | AA | AB |
| AB | AB | AB | AB | AB | AB | AB | AB | AB | AB | AB | AB | AB | AB | AB | AB | AA |
| AB | AA | AA | AA | AA | AA | AA | AA | AA | AA | AA | AB | AA | AA | AA | AA | AA |
| AA | - | - | - | AA | - | - | AB | AA | AB | AB | AB | AA | - | - | AB | AA |
| AA | AA | AA | AA | AB | AB | AA | AB | AB | AA | AB | AB | AB | AA | AB | AB | AB |
| AB | AA | AA | AA | AB | AB | AA | AB | AB | AA | AB | AB | AB | AA | AB | AB | AB |
| AB | AA | AA | AB | AB | AA | AB | AA | AA | - | AB | AB | AA | AB | AB | AB | AB |
| AA | AB | AA | AB | AB | AA | AB | AA | AA | AA | AB | AB | AB | AA | AB | AA | AB |
| AB | AB | AB | AB | AA | AB | AB | AB | AB | AB | AB | AB | AB | AB | AB | AB | AA |
| AA | AB | AB | AA | AB | AB | AA | AB | AB | AA | AB | AB | AB | AA | - | AA | AA |
| AB | - | - | AB | AA | AA | AA | AA | AA | - | AA | - | AA | AA | - | AA | - |
| AB | AB | AB | AB | AB | AB | AB | AB | AB | AB | AB | AB | AB | AB | AB | AB | AB |
| AB | AA | AB | AB | AB | AB | AB | AB | AA | AB | AB | AB | AB | AB | AB | AB | AB |
| AB | AB | AB | - | AA | - | AB | AA | AB | AA | AB | AB | AA | AB | AA | AB | AA |
| AB | - | AB | - | AA | AB | AB | AB | AB | AA | AB | AA | AB | AB | - | AB | AA |
| AB | - | AA | AB | AA | - | AB | AB | AA | AB | AB | AB | AA | AB | AB | AB | AA |
| AB | AB | AB | AB | AB | AB | AB | AB | AB | AB | AB | AB | AB | AB | AB | AB | AB |
| AA | AA | AA | AA | AA | AA | AA | AA | AA | AA | AA | AA | AA | AA | AA | AA | AA |
| AA | AB | AA | AA | AA | AA | AA | AA | AA | AA | AA | AA | AB | AA | - | AA | AA |
| AB | AA | AB | AA | AA | AB | AA | AB | AB | AB | AB | AB | AA | AB | AB | AA | AB |
| AB | - | AA | AB | AA | - | AA | AA | AA | AA | AA | AB | AA | AB | - | AA | AA |
| AB | AA | AA | AB | AB | AA | AB | AA | AA | AB | AB | AB | AA | AB | AB | AB | AB |
| AA | AB | AB | AB | AB | AB | AB | AA | AB | AB | AB | AB | AB | AA | AA | AB | AB |
| AA | AB | AB | AB | AB | AB | AB | AB | AB | AB | AB | AB | AB | AB | AA | AB | AB |
| AB | AB | AA | AB | AB | - | AA | AB | AB | AA | AB | AA | AA | AA | - | AA | AB |
| AB | AB | AA | AA | AA | AB | AA | AA | AB | AA | AA | AA | AA | AA | AA | AA | AB |
| AB | AA | AB | AA | AB | AB | AA | AB | AB | AB | AA | AA | AA | AB | AB | AB | AB |
| AA | AB | AA | AA | AB | AA | AB | AA | AB | AB | AA | AB | AA | AA | AA | AB | AA |
| AB | AA | AA | AA | AB | AB | AA | AB | AB | AA | AB | AB | AB | AA | AA | AB | AA |
| AA | AA | AB | AB | AB | AB | AB | AB | AA | AA | AB | AB | AB | AA | AA | AB | AB |
| AA | AB | AB | AA | AB | AB | AB | AB | AA | AA | AB | AA | AB | AB | AA | AB | AB |
| AA | AB | AB | AA | AB | AB | AB | AB | AB | AA | AB | AB | AB | AB | AA | AB | AB |

| snp_B3456 | snp_B3463 | snp_B3469 | snp_B3489 | snp_B3500 | snp_B3537 | snp_B3571 | snp_B3574 | snp_B3582 | snp_B3590 | snp_B3623 | snp_B3654 | snp_B3700 | snp_B3701 | snp_B3800 | snp_B3813 | snp_B3837 | snp_B3854 | snp_B3856 | snp_B3868 | snp_B3884 | snp_B3908 | snp_B3919 | snp_B3953 | snp_B4003 | snp_B4016 | snp_B4038 | snp_B4061 | snp_B4083 | snp_B4119 | snp_B4131 | snp_B4135 | snp_B4166 | snp_B4178 | snp_B4182 | snp_B4213 | snp_B4235 | snp_B4262 | snp_B4304 | snp_B4305 | snp_B4313 | snp_B4316 | snp_B4340 | snp_B4399 | snp_B4401 | snp_B4416 | snp_B4429 | snp_B4448 | snp_B4484 | snp_B4485 | snp_B4486 | snp_B4498 | snp_B4534 | snp_B4555 | snp_B4634 | snp_B4648 | snp_B4659 | snp_B4682 | snp_B4689 | snp_B4698 | snp_B4708 | snp_B4723 | snp_B4750 | snp_B4762 | snp_B4856 | snp_B4884 | snp_B4895 | snp_B4900 | snp_B4906 | snp_B4923 | snp_B4951 | snp_B4969 | snp_B4980 | snp_B4993 | snp_B4998 | snp_B5097 | snp_B5132 |
|---|---|---|---|---|---|---|---|---|---|---|---|---|---|---|---|---|---|---|---|---|---|---|---|---|---|---|---|---|---|---|---|---|---|---|---|---|---|---|---|---|---|---|---|---|---|---|---|---|---|---|---|---|---|---|---|---|---|---|---|---|---|---|---|---|---|---|---|---|---|---|---|---|---|---|---|---|
| 8 | 8 | 8 | 8 | 8 | 8 | 8 | 8 | 8 | 8 | 8 | 8 | 8 | 8 | 8 | 8 | 8 | 8 | 8 | 8 | 8 | 8 | 8 | 8 | 8 | 8 | 8 | 8 | 8 | 8 | 8 | 8 | 8 | 8 | 8 | 8 | 8 | 8 | 8 | 8 | 8 | 8 | 8 | 8 | 8 | 8 | 8 | 8 | 8 | 8 | 8 | 8 | 8 | 8 | 8 | 8 | 8 | 8 | 8 | 8 | 8 | 8 | 8 | 8 | 8 | 8 | 8 | 8 | 8 | 8 | 8 | 8 | 8 | 8 | 8 | 8 | 8 |
| -- | -- | -- | -- | LG10 | -- | -- | -- | -- | -- | -- | -- | LG10 | -- | -- | -- | -- | -- | -- | -- | -- | -- | -- | -- | -- | -- | -- | -- | -- | -- | -- | -- | -- | -- | -- | -- | -- | -- | -- | -- | -- | -- | -- | -- | -- | -- | -- | -- | -- | -- | LG10 | -- | -- | -- | -- | -- | -- | -- | -- | -- | -- | -- | -- | -- | -- | -- | -- | -- | -- | -- | -- | -- | -- | -- | -- | -- | -- |
| AB | AB | AB | AA | AA | AB | AB | AB | AB | AA | AA | AB | AB | AA | AB | AA | AA | AA | AB | AB | AA | AA | AB | AB | AB | AA | AA | AB | AA | AB | AB | AB | AB | AB | AA | AA | AA | AB | AA | AB | AA | AA | AA | AB | AB | AB | AB | AB | AA | AB | AA | AB | AA | AB | AA | AA | AB | AA | AA | AB | AA | AA | AA | AA | AB | AB | AB | AA | AA | AA | AB | AA | AA | AA | AA | AA | AB |

| snp_B5143 | snp_B5171 | snp_B5175 | snp_B5255 | snp_B5297 | snp_B5309 | snp_B5314 | snp_B5364 | snp_B5380 | snp_B5382 | snp_B5395 | snp_B5409 | snp_B5425 | snp_B5442 | snp_B5465 | snp_B5473 | snp_B5488 | snp_B5537 | snp_B5540 | snp_B5557 | snp_B5571 | snp_B5583 | snp_B5586 | snp_B5624 | snp_B5630 | snp_B5637 | snp_B5664 | snp_B5687 | snp_B5718 | snp_B5725 | snp_B5753 | snp_B5757 | snp_B5759 | snp_B5898 | snp_B5936 | snp_B6001 | snp_B6150 | snp_B6160 | snp_B0002 | snp_B0006 | snp_B0084 | snp_B0090 | snp_B0091 | snp_B0115 | snp_B0173 | snp_B0190 | snp_B0222 | snp_B0238 | snp_B0305 | snp_B0403 | snp_B0465 | snp_B0481 | snp_B0523 | snp_B0532 | snp_B0533 | snp_B0555 | snp_B0577 | snp_B0600 | snp_B0617 | snp_B0641 | snp_B0649 | snp_B0662 | snp_B0710 | snp_B0713 | snp_B0719 | snp_B0739 | snp_B0741 | snp_B0755 | snp_B0764 | snp_B0793 | snp_B0864 | snp_B0876 | snp_B0902 | snp_B0905 | snp_B0922 | snp_B0952 | snp_B0980 |
|---|---|---|---|---|---|---|---|---|---|---|---|---|---|---|---|---|---|---|---|---|---|---|---|---|---|---|---|---|---|---|---|---|---|---|---|---|---|---|---|---|---|---|---|---|---|---|---|---|---|---|---|---|---|---|---|---|---|---|---|---|---|---|---|---|---|---|---|---|---|---|---|---|---|---|---|---|
| 8 | 8 | 8 | 8 | 8 | 8 | 8 | 8 | 8 | 8 | 8 | 8 | 8 | 8 | 8 | 8 | 8 | 8 | 8 | 8 | 8 | 8 | 8 | 8 | 8 | 8 | 8 | 8 | 8 | 8 | 8 | 8 | 8 | 8 | 8 | 8 | 8 | 8 | 9 | 9 | 9 | 9 | 9 | 9 | 9 | 9 | 9 | 9 | 9 | 9 | 9 | 9 | 9 | 9 | 9 | 9 | 9 | 9 | 9 | 9 | 9 | 9 | 9 | 9 | 9 | 9 | 9 | 9 | 9 | 9 | 9 | 9 | 9 | 9 | 9 | 9 | 9 |
| -- | -- | -- | -- | -- | -- | -- | -- | -- | -- | -- | -- | -- | -- | -- | -- | -- | -- | -- | -- | -- | -- | -- | -- | LG10 | -- | -- | -- | -- | -- | -- | -- | -- | -- | -- | -- | -- | -- | -- | -- | -- | -- | -- | -- | -- | LG2 | -- | -- | -- | -- | -- | LG2 | -- | -- | -- | -- | LG2 | -- | -- | -- | -- | -- | -- | -- | -- | -- | -- | -- | -- | -- | -- | -- | -- | -- | -- | -- | -- |
| AA | AA | AB | AB | AB | AB | AA | AA | AA | AB | AA | AA | AA | AB | AA | AA | AB | AB | AB | AA | AB | AA | AB | AB | AB | AA | AB | AA | AA | AA | AA | AB | AA | AB | AA | AB | AA | AA | AA | AA | AA | AB | AA | AA | AA | AB | AB | AB | AA | AB | AB | AB | AA | AB | AB | AA | AB | AA | AA | AB | AA | AA | AA | AA | AA | AA | AB | AB | AA | AA | AB | AB | AA | AA | AB | AA | AA |
| AA | AB | AB | AB | AA | AA | AB | AA | AB | AA | AA | AA | AA | AB | AB | AB | AB | AA | AA | AA | AB | AB | AA | AA | AB | AA | AB | AA | AB | AB | AA | AB | AA | AB | AA | AA | AA | AB | AB | AA | AB | AB | AB | AA | AA | AB | AB | AB | AA | AB | AB | AB | AB | AB | AB | AA | AB | AA | AB | AB | AA | AA | AA | AB | AA | AB | AB | AB | AA | AB | AB | AB | AB | AA | AB | AB | AB |
| AA | AB | AB | AB | AB | AA | AB | AA | AA | AA | AA | AB | AB | AB | AA | AB | AA | AA | AA | AB | AA | AA | AA | AA | AB | AA | AA | AB | AB | AA | AA | AA | AB | AA | AB | AA | AB | AB | AA | AB | AA | AB | AA | AB | AB | AA | AB | AB | AB | AA | AB | AA | AB | AA | AB | AA | AA | AB | AA | AA | AB | AA | AB | AA | AB | AA | AB | AB | AA | AB | AB | AB | AA | AB | AA | AB | AB |
| AB | AB | - | AB | AA | AA | AB | AA | AA | AB | AA | - | AA | AB | AA | AB | AB | - | AA | AA | AB | AB | AA | AA | AB | AB | AB | AA | AB | AA | AB | AB | - | AB | AA | AB | AA | AB | - | AA | AA | AB | AB | AB | - | AA | AA | AA | AB | AB | AB | AB | AB | - | AB | AB | - | AA | AB | AA | AB | - | AB | AB | AB | - | AA | AB | AB | AB | AB | AA | AA | AB | AB | AB | AB |
| AA | AB | AB | AB | AB | AA | AB | AB | AB | AB | AB | AB | AA | AB | AA | AB | AB | AB | AA | AB | AB | AA | AB | AA | AB | AA | AA | AA | AB | AB | AA | AA | AB | AA | AA | AA | AA | AB | AA | AA | AB | AB | AB | AA | AA | AB | AA | AB | AA | AA | AB | AA | AB | AB | AB | AA | AA | AA | AB | AB | AA | AB | AB | AA | AA | AB | AB | AA | AA | AA | AB | AA | AA | AB | AB | AB | AB |
| AA | AA | - | AB | AB | AB | AA | - | AB | AB | AA | - | AA | - | AB | AA | AA | AB | AA | AA | AA | AA | AB | AB | AB | - | AA | AA | AB | AB | AB | AB | - | AA | AA | AB | AA | - | - | - | AA | AA | AB | - | - | AA | AB | AA | - | AA | - | - | AA | AA | AB | AB | - | AB | - | - | AB | AA | AA | - | AB | AA | AA | AB | AB | AB | - | - | - | AB | AB | AB | AB |
| AB | AA | AA | AA | AA | AA | AA | AA | AA | AA | AB | AA | AA | AB | AA | AA | AB | AB | AA | AA | AA | AA | AA | AA | AA | AB | AA | AA | AA | AA | AB | AA | AA | AA | AB | AB | AB | AA | AB | AB | AA | AB | AB | AB | AB | AB | AB | AB | AB | AB | AB | AB | AB | AB | - | AB | AB | AB | AB | AA | AB | AB | AB | AB | AB | AB | AB | AB | AB | AB | AA | AB | AB | AB | AB | AB | AB |
| AA | AA | AA | AA | AA | AA | AA | AA | AB | AA | AB | AA | AA | AB | AB | AA | AB | AA | AA | AA | AA | AA | AA | AA | AA | AA | AA | AA | AA | AB | AB | AA | AA | AA | AA | AA | AB | AA | AA | AB | AB | AA | AA | AB | AB | AA | AB | AA | AB | AA | AA | AA | AB | AA | AA | AB | AA | AB | AA | AB | AB | AB | AA | AA | AB | AB | AA | AA | AB | AA | AB | AB | AB | AB | AA | AA | AA |
| AA | AB | AA | AB | AA | AB | AB | AA | AA | AB | AA | AA | AA | AA | AA | AA | AB | AA | AB | AA | AA | AB | AB | AA | AB | AA | AB | AA | AA | AA | AA | AA | AA | AA | AA | AB | AA | AA | AB | AB | AA | AA | AA | AB | AB | AA | AB | AA | AB | AA | AB | AA | AB | AA | AA | AB | AA | AB | AA | AB | AA | AB | AA | AA | AB | AA | AA | AA | AB | AA | AB | AB | AA | AB | AA | AA | AA |
| AA | AB | AA | AB | AB | AA | AB | AA | AA | AB | AB | AA | AA | AB | AA | AB | AB | AB | AA | AA | AA | AA | AA | AA | AB | AA | AB | AA | AB | AB | AA | AA | AA | AA | AA | AA | AA | AB | AA | AA | AA | AB | AB | AB | AA | AA | AA | AA | AA | AB | AA | AB | AA | AA | AB | AA | AA | AA | AA | AB | AA | AB | AB | AB | AB | AA | AA | AA | AA | AA | AB | AA | AA | AB | AA | AA | AA |
| AB | AB | AA | AA | AB | AB | AA | AA | AA | AA | AB | AA | AA | AA | AA | AA | AB | AA | AB | AA | AB | AA | AB | AA | AA | AB | AB | AA | AA | AA | AB | AB | AA | AB | AA | AB | AB | AA | AB | AB | AB | AA | AA | AA | AB | AA | AA | AA | AB | AA | AB | AB | AA | AA | AA | AA | AB | AA | AB | AB | AB | AB | AB | AB | AA | AA | AA | AA | AA | AA | AB | AB | AA | AB | AB | AB | AB |
| AA | AA | AA | AA | AA | AB | AA | AA | AB | AA | AA | AA | AA | AA | AB | AA | AB | AA | AB | AA | AA | AA | AB | AA | AA | AA | AA | AA | AA | AB | AB | AA | AA | AA | AA | AA | AB | AA | AA | AA | AA | AA | AA | AA | AA | AA | AA | AA | AB | AB | AA | AB | AA | AA | AA | AA | AB | AB | AB | AA | AB | AA | AA | AA | AA | - | AA | AB | AA | AB | AA | AA | AA | AB | AA | AA | AA |
| AB | AB | AA | AA | AB | AB | AA | AB | AA | AB | AA | AB | AB | AA | AA | AB | AB | AB | AA | AB | AA | AA | AB | AB | AA | AB | AA | AB | AB | AA | AA | AA | AB | AA | AA | AB | AA | AA | AA | AA | AB | AA | AB | AA | AA | AB | AB | AB | AA | AB | AB | AB | AA | AB | AB | AA | AB | AA | AB | AB | AA | AA | AB | AB | AA | AB | AB | AB | AA | AB | AB | AB | AB | AA | AB | AB | AB |
| AA | AB | AB | AB | AA | AB | AB | AA | AB | AB | AB | AA | AB | AA | AB | AB | AB | AA | AB | AB | AA | AA | AB | AB | AB | AA | AA | AA | AB | AB | AB | AA | AA | AA | AB | AB | AB | AA | AB | AB | AA | AB | AB | AA | AB | AA | AB | AA | AA | AA | AB | AB | AB | AA | AB | AA | AB | AA | AB | AB | AB | AB | AB | AB | AB | AA | AA | AB | AA | AB | AB | AA | AB | AA | AB | AB | AB |
| AA | AA | AA | AA | AA | AA | AB | AB | AA | AB | AA | AA | AA | AA | AA | AB | AA | AB | AA | AA | AA | AB | AB | AB | AA | AA | AB | AA | AA | AA | AA | AA | AB | AA | AA | AB | AA | AA | AA | AA | AA | AA | AA | AA | AA | AB | AB | AB | AA | AA | AA | AB | AA | AB | AA | AA | AB | AA | AA | AB | AA | AB | AA | AB | AA | AB | AA | AB | AA | AB | AB | AA | AA | AA | AA | AA | AA |
| AB | AB | AB | AB | AB | AA | AB | AA | AB | AA | AA | AB | AA | AA | AA | AB | AB | AB | AB | AB | AB | AB | AB | AB | AB | AB | AB | AB | AB | AB | AA | AA | AB | AB | AB | AB | AA | AB | AB | AA | AB | AA | AB | AA | AA | AA | AB | AB | AB | AA | AA | AB | AA | AB | AB | AB | AB | AA | AA | AB | AA | AA | AB | AA | AA | AB | AB | AA | AB | AA | AB | AA | AB | AA | AB | AA | AA |
| AB | AA | AB | AB | AB | AB | AB | AB | AA | AB | AB | AB | AB | AA | AA | AB | AB | AB | AB | AB | AB | AB | AB | AB | AB | AB | AB | AB | AB | AA | AA | AA | AB | AB | AB | AB | AA | AB | AB | AA | AB | AA | AB | AB | AB | AB | AA | AA | AA | AB | AA | AA | AB | AA | AB | AA | AA | AB | AA | AB | AA | AA | AB | AA | AB | AA | AA | AA | AB | AA | AB | AA | AA | AA | AB | AA | AA |
| AA | AB | - | AB | - | AA | AA | AA | AB | AA | AB | - | AB | AB | AB | AB | AA | AA | AA | - | AB | AA | AA | AA | AB | - | AA | AB | AB | AB | AA | AA | AB | AA | AB | AA | AA | - | - | AB | AA | AB | AB | - | - | - | AB | AA | AA | AA | - | AB | AA | - | - | AB | - | AA | - | - | AB | - | AB | - | AB | AA | AA | AB | AB | AB | - | - | - | AA | AB | AB | - |
| AA | AA | AA | AA | AA | AA | AA | AA | AA | AA | AA | AA | AA | AA | AA | AA | AA | AA | AA | AA | AA | AA | AA | AA | AA | AA | AA | AA | AA | AA | AA | AA | AA | AA | AA | AA | AA | AA | AA | AA | AA | AA | AA | AA | AA | AA | AA | AA | AB | AB | AA | AB | AA | AA | AA | AB | AB | AB | AB | AA | AB | AA | AA | AA | AA | AB | AA | AB | AB | AA | AA | AA | AA | AB | AB | AA | AA |
| AB | AB | AB | AA | AA | AA | AA | AA | AA | AA | AA | - | AA | - | AA | AB | AA | AB | AA | AA | AA | AA | AA | AA | AA | AB | AB | AA | AB | AA | AA | AA | AA | AA | AB | AB | AA | AA | AB | AB | AA | AA | AB | AA | AA | AB | AA | AA | AB | AA | AA | AB | AB | AB | AB | AB | - | AA | AB | AA | AB | AA | AB | AB | AA | - | AA | AB | AB | AA | AA | AA | AA | AB | AB | AA | AA |
| AB | AB | AA | AB | AB | AB | AB | AB | AB | AB | AB | AB | AB | AB | AB | AB | AB | AB | AB | AB | AB | AB | AB | AB | AB | AB | AB | AB | AB | AB | AB | AB | AB | AA | AB | AB | AB | AB | AB | AB | AA | AB | AB | AB | AB | AA | AA | AA | AA | AA | AB | AA | AA | AA | AA | AB | AA | AA | AB | AA | AB | AA | AA | AB | AB | AB | AA | AA | AB | AA | AB | AA | AA | AA | AB | AA | AA | AA |
| AA | AA | AA | AA | AA | AA | AA | AA | AB | AA | AA | AA | AA | AB | AB | AA | AA | AA | AA | AA | AA | AA | AA | AA | AA | AA | AA | AA | AA | AA | AB | AB | AA | AA | AA | AA | AA | AB | AA | AA | AB | AA | AB | AA | AA | AA | AB | AB | AB | AB | AA | AB | AA | AB | AB | AA | AA | AA | AB | AB | AA | AB | AA | AB | AA | AA | AA | AB | AA | AA | AB | AA | AB | AB | AA | AB | AA | AA |
| - | AB | - | AB | - | AA | AB | AB | AA | AA | AA | - | AA | - | AA | AA | AB | AA | AA | AA | AB | AB | AB | AA | AB | AA | AB | AA | AA | AA | AA | - | AA | AA | AA | AB | AA | AB | AB | AB | AA | AB | AB | - | AB | - | AA | AA | AA | AA | - | AB | AA | - | AB | AA | - | AA | - | AA | AB | AA | AB | - | AA | - | AA | AA | AA | AB | AB | AA | AB | AA | AB | AB | AB |
| AA | AB | AB | AA | AA | AA | AB | AB | AA | AA | AB | AB | AB | AA | AA | AB | AB | AA | AB | AB | AA | AB | AA | AA | AA | AA | AB | AB | AA | AA | AA | AA | AB | AB | AB | AA | AB | AA | AB | AA | AB | AA | AA | AA | AA | AA | AB | AA | AA | AA | AB | AA | AA | AA | AA | AA | AA | AB | AA | AB | AA | AA | AB | AA | AA | AA | AA | AB | AA | AB | AB | AB | AB | AB | AB | AB | AB |
| AA | AB | AB | AA | AA | AA | AB | AB | AB | AA | AB | AB | AB | AB | AB | AB | AB | AA | AB | AB | AA | AB | AA | AA | AA | AA | AA | AB | AA | AB | AB | AA | AB | AA | AA | AA | AB | AA | AB | AA | AB | AB | AB | AA | AB | AA | AA | AA | AB | AA | AB | AB | AA | AA | AB | AA | AA | AB | AB | AB | AA | AA | AB | AB | AA | AA | AA | AB | AA | AB | AB | AA | AB | AB | AA | AB | AB |
| AA | AB | AA | AB | AB | AA | AB | AA | AA | AB | AA | AA | AA | AB | AB | AB | AB | AB | AA | AB | AA | AB | AA | AB | AB | AA | AA | AB | AB | AA | AB | AA | AB | AA | AA | AA | AB | AB | AA | AA | AB | AB | AA | AA | AA | AA | AA | AB | AA | AA | AB | AA | AB | AB | AA | AB | AA | AA | AA | AA | AA | AA | AB | AA | AA | AB | AB | AA | AB | AA | AA | AB | AB | AA | AB | AB | AB |
| AA | AB | AB | AB | AA | AA | AB | AA | AA | AB | AA | AB | AA | AB | AA | AB | AB | AB | AB | AA | AB | AB | AA | AB | AA | AA | AA | AB | AB | AA | AB | AA | AB | AA | AA | AA | AB | AB | AA | AA | AB | AB | AA | AA | AA | AA | AB | AB | AB | AB | AA | AB | AB | AB | AA | AA | AB | AB | AB | AA | AA | AB | AA | AB | AA | AB | AB | AB | AA | AB | AB | AB | AB | AA | AB | AA | AB |
| AB | AA | AA | AB | AB | AB | AB | AB | AA | AB | AB | AB | AB | AB | AB | AB | AB | AA | AB | AB | AB | AB | AB | AB | AB | AA | AA | AB | AB | AB | AB | AA | AB | AB | AB | AB | AB | AA | AB | AB | AB | AB | AB | AA | AB | AB | AB | AB | AB | AB | AB | AB | AB | AB | AB | AB | AB | AB | AB | AB | AA | AB | AA | AB | AA | AB | AB | AB | AB | AB | AB | AB | AB | AA | AB | AA | AB |
| AB | AA | AA | AA | AB | AB | AA | AA | AB | AA | AA | AB | AB | AA | - | AB | AA | AA | AB | AB | AB | AA | AA | AA | - | AB | AB | AB | AB | AA | AB | AB | AB | AB | AB | AB | AA | AB | AB | AB | AB | AB | AB | AB | AB | - | AA | AB | AB | AB | AB | AA | AB | AA | AA | AB | AA | AB | AB | AB | AB | AB | AB | AB | AA | - | AA | AB | AA | AB | AA | AB | AB | AA | AB | AB | AA |
| - | AA | - | AA | AA | AB | AA | - | AA | AA | AA | AA | AA | AA | - | AA | AB | - | AA | AB | AA | AA | AA | AA | - | AA | AB | AA | AA | AA | AA | - | AA | AA | AA | AA | AA | - | - | - | AB | AA | AA | - | - | - | AB | AB | AA | AB | - | AB | AA | - | - | - | - | AB | - | AB | AB | AB | - | - | AA | AA | AB | AB | - | AB | - | AB | AA | AB | AB | AB | - |
| AB | AB | AB | AB | AB | AB | AB | AB | AB | AB | AB | AB | AB | AB | AB | AB | AB | AB | AB | AB | AB | AB | AB | AB | AB | AB | AB | AB | AB | AB | AB | AB | AB | AB | AB | AB | AB | AB | AB | AB | AB | AB | AA | AB | AA | AA | AA | AB | AB | AA | AA | AB | AB | AB | AB | AB | AB | AA | AB | AB | AB | AB | AA | AB | AA | AB | AB | AB | AB | AB | AB | AA | AA | AB | AB | AA | AA |

| snp_B1019 | snp_B1077 | snp_B1089 | snp_B1111 | snp_B1116 | snp_B1149 | snp_B1169 | snp_B1172 | snp_B1182 | snp_B1185 | snp_B1188 | snp_B1196 | snp_B1197 | snp_B1209 | snp_B1268 | snp_B1276 | snp_B1297 | snp_B1309 | snp_B1318 | snp_B1324 | snp_B1334 | snp_B1345 | snp_B1353 | snp_B1373 | snp_B1384 | snp_B1402 | snp_B1409 | snp_B1417 | snp_B1420 | snp_B1425 | snp_B1461 | snp_B1494 | snp_B1527 | snp_B1544 | snp_B1549 | snp_B1554 | snp_B1576 | snp_B1605 | snp_B1628 | snp_B1633 | snp_B1654 | snp_B1686 | snp_B1690 | snp_B1694 | snp_B1751 | snp_B1763 | snp_B1787 | snp_B1799 | snp_B1829 | snp_B1907 | snp_B1937 | snp_B1951 | snp_B1958 | snp_B2032 | snp_B2045 | snp_B2075 | snp_B2084 | snp_B2096 | snp_B2128 | snp_B2165 | snp_B2189 | snp_B2239 | snp_B2240 | snp_B2246 | snp_B2341 | snp_B2355 | snp_B2369 | snp_B2381 | snp_B2384 | snp_B2472 | snp_B2483 | snp_B2497 | snp_B2498 | snp_B2512 | snp_B2539 | snp_B2565 | snp_B2570 |
|---|---|---|---|---|---|---|---|---|---|---|---|---|---|---|---|---|---|---|---|---|---|---|---|---|---|---|---|---|---|---|---|---|---|---|---|---|---|---|---|---|---|---|---|---|---|---|---|---|---|---|---|---|---|---|---|---|---|---|---|---|---|---|---|---|---|---|---|---|---|---|---|---|---|---|---|---|
| 9 | 9 | 9 | 9 | 9 | 9 | 9 | 9 | 9 | 9 | 9 | 9 | 9 | 9 | 9 | 9 | 9 | 9 | 9 | 9 | 9 | 9 | 9 | 9 | 9 | 9 | 9 | 9 | 9 | 9 | 9 | 9 | 9 | 9 | 9 | 9 | 9 | 9 | 9 | 9 | 9 | 9 | 9 | 9 | 9 | 9 | 9 | 9 | 9 | 9 | 9 | 9 | 9 | 9 | 9 | 9 | 9 | 9 | 9 | 9 | 9 | 9 | 9 | 9 | 9 | 9 | 9 | 9 | 9 | 9 | 9 | 9 | 9 | 9 | 9 | 9 | 9 |
| LG2 | -- | -- | -- | -- | -- | -- | -- | -- | -- | -- | -- | -- | LG2 | -- | -- | -- | -- | -- | -- | LG2 | -- | -- | -- | -- | -- | -- | LG2 | -- | -- | LG2 | -- | -- | -- | -- | -- | -- | -- | -- | -- | -- | -- | -- | -- | -- | -- | -- | -- | -- | -- | -- | -- | -- | -- | -- | -- | -- | -- | -- | -- | -- | -- | -- | -- | -- | -- | -- | LG2 | -- | -- | -- | -- | -- | -- | -- | -- | -- |
| AB | AB | AB | AA | AB | AB | AB | AB | AB | AB | AA | AB | AA | AB | AA | AB | AB | AB | AA | AA | AA | AA | AB | AA | AB | AB | AA | AB | AB | AA | AB | AA | AB | AB | AA | AB | AA | AA | AB | AB | AA | AA | AA | AB | AA | AB | AB | AB | AA | AB | AA | AA | AA | AA | AB | AB | AA | AB | AA | AB | AB | AA | AA | AA | AB | AA | AA | AA | AB | AB | AA | AA | AA | AA | AB | AA | AB |
| AB | AB | AB | AB | AB | AB | AB | AB | AB | AA | AB | AA | AB | AB | AB | AB | AB | AA | AA | AB | AB | AA | AB | AB | AB | AB | AB | AB | AB | AA | AB | AB | AB | AB | AA | AB | AB | AA | AB | AB | AA | AA | AA | AA | AB | AB | AB | AA | AB | AB | AA | AB | AB | AB | AA | AA | AA | AB | AA | AB | AA | AB | AB | AB | AB | AB | AB | AB | AA | AB | AB | AA | AA | AB | AB | AA | AB |
| AA | AB | AA | AA | AA | AA | AA | AA | AA | AA | AB | AA | AA | AB | AB | AB | AB | AA | AA | AA | AA | AB | AA | AB | AA | AB | AB | AA | AB | AA | AA | AA | AA | AA | AB | AA | AB | AB | AA | AB | AB | AB | AA | AB | AA | AA | AB | AA | AA | AA | AB | AB | AB | AA | AA | AA | AA | AA | AB | AB | AB | AA | AB | AB | AB | AA | AB | AB | AA | AA | AA | AB | AB | AA | AB | AB | AB |
| AB | AA | AB | AB | AB | AB | AB | AB | AB | AB | AB | - | - | AA | AA | AB | AB | AA | AB | AB | AA | AB | AB | AB | AB | AA | AB | - | AB | AA | AB | AA | AB | - | AB | AB | AB | AA | AB | AA | AB | AA | AA | AA | AB | AB | AA | AA | AA | AB | AA | AA | AB | AB | AB | AB | - | AB | AA | AB | AB | AA | AB | AA | AB | AA | AB | - | AB | AB | AB | AA | AA | - | AB | AA | AB |
| AA | AB | AA | AA | AB | AA | AB | AA | AA | AA | AA | AA | AA | AA | AA | AA | AA | AA | AA | AA | AB | AA | AA | AB | AA | AA | AA | AA | AB | AA | AB | AA | AA | AA | AA | AB | AA | AA | AA | AA | AA | AA | AA | AA | AB | AB | AB | AA | AA | AA | AA | AA | AA | AB | AA | AB | AA | AA | AA | AA | AA | AB | AB | AB | AA | AA | AB | AA | AA | AA | AB | AA | AA | AA | AB | AA | AA |

| snp_B2595 | snp_B2617 | snp_B2689 | snp_B2719 | snp_B2728 | snp_B2740 | snp_B2775 | snp_B2782 | snp_B2832 | snp_B2835 | snp_B2867 | snp_B2879 | snp_B2909 | snp_B2934 | snp_B2938 | snp_B2942 | snp_B2952 | snp_B2961 | snp_B2965 | snp_B2967 | snp_B2982 | snp_B2996 | snp_B2998 | snp_B3014 | snp_B3022 | snp_B3030 | snp_B3036 | snp_B3037 | snp_B3039 | snp_B3040 | snp_B3055 | snp_B3066 | snp_B3091 | snp_B3116 | snp_B3125 | snp_B3127 | snp_B3156 | snp_B3176 | snp_B3189 | snp_B3199 | snp_B3200 | snp_B3201 | snp_B3223 | snp_B3226 | snp_B3281 | snp_B3286 | snp_B3323 | snp_B3347 | snp_B3360 | snp_B3367 | snp_B3455 | snp_B3460 | snp_B3488 | snp_B3491 | snp_B3521 | snp_B3535 | snp_B3550 | snp_B3562 | snp_B3565 | snp_B3581 | snp_B3628 | snp_B3631 | snp_B3641 | snp_B3657 | snp_B3669 | snp_B3713 | snp_B3719 | snp_B3721 | snp_B3729 | snp_B3738 | snp_B3773 | snp_B3788 | snp_B3835 | snp_B3849 | snp_B3851 | snp_B3859 | snp_B3889 |
|---|---|---|---|---|---|---|---|---|---|---|---|---|---|---|---|---|---|---|---|---|---|---|---|---|---|---|---|---|---|---|---|---|---|---|---|---|---|---|---|---|---|---|---|---|---|---|---|---|---|---|---|---|---|---|---|---|---|---|---|---|---|---|---|---|---|---|---|---|---|---|---|---|---|---|---|---|
| 9 | 9 | 9 | 9 | 9 | 9 | 9 | 9 | 9 | 9 | 9 | 9 | 9 | 9 | 9 | 9 | 9 | 9 | 9 | 9 | 9 | 9 | 9 | 9 | 9 | 9 | 9 | 9 | 9 | 9 | 9 | 9 | 9 | 9 | 9 | 9 | 9 | 9 | 9 | 9 | 9 | 9 | 9 | 9 | 9 | 9 | 9 | 9 | 9 | 9 | 9 | 9 | 9 | 9 | 9 | 9 | 9 | 9 | 9 | 9 | 9 | 9 | 9 | 9 | 9 | 9 | 9 | 9 | 9 | 9 | 9 | 9 | 9 | 9 | 9 | 9 | 9 |
| -- | -- | -- | -- | -- | LG2 | -- | LG2 | -- | -- | -- | -- | -- | -- | -- | -- | -- | -- | -- | -- | -- | -- | -- | -- | -- | -- | -- | -- | -- | -- | -- | -- | -- | -- | -- | -- | -- | -- | -- | -- | LG2 | -- | -- | -- | -- | -- | -- | -- | -- | LG2 | -- | -- | LG2 | -- | -- | LG2 | -- | -- | -- | LG2 | -- | LG2 | -- | -- | -- | -- | -- | -- | -- | -- | -- | -- | -- | LG2 | -- | LG2 | -- |

| snp_B3916 9 | snp_B3930 9 | snp_B3939 9 | snp_B3970 9 | snp_B3994 9 | snp_B3995 9 | snp_B4029 9 | snp_B4031 9 | snp_B4033 9 | snp_B4046 9 | snp_B4047 9 | snp_B4068 9 | snp_B4075 9 | snp_B4090 9 | snp_B4095 9 | snp_B4103 9 | snp_B4134 9 | [illegible] |
|---|---|---|---|---|---|---|---|---|---|---|---|---|---|---|---|---|---|
| -- | -- | -- | -- | -- | -- | -- | -- | -- | -- | -- | -- | -- | -- | -- | -- | -- | [illegible] |
| AA | AA | AB | AB | AA | AB | AB | AA | AA | AA | AA | AB | AB | AA | AA | AA | AA | [illegible] |
| AA | AB | AB | AB | AB | AB | AB | AB | AA | AA | AB | AB | AA | AB | AA | AA | AB | [illegible] |
| AB | AB | AB | AA | AA | AA | AA | AA | AB | AA | AA | AA | AA | AA | AB | AB | AA | [illegible] |
| - | AA | AA | AB | AB | - | AB | AB | AB | AA | AB | AB | AA | AB | AA | AA | AB | [illegible] |
| AA | AB | AA | AA | AA | AA | AA | AA | AA | AA | AB | AA | AA | AA | AA | AA | AA | [illegible] |
| AB | AA | AA | AA | AB | - | AB | AB | AB | AB | AA | AB | AB | - | AB | AB | AB | [illegible] |
| AA | AB | AB | AB | AB | AB | AB | AB | AB | AB | AB | AB | AB | - | AB | AB | AB | [illegible] |
| AA | AA | AB | AB | AB | AA | AA | AA | AB | AB | AB | AA | AB | AA | AB | AB | AB | [illegible] |
| AA | AB | AB | AB | AB | AA | AA | AA | AB | AA | AB | AA | AA | AB | AB | AB | AB | [illegible] |
| AB | AA | AA | AA | AB | AA | AA | AA | AA | AB | AB | AA | AA | AB | AB | AB | AA | [illegible] |
| AA | AA | AA | AA | AB | AB | AB | AB | AA | AB | AA | AA | AB | AB | AA | AB | AA | [illegible] |
| AB | AA | AA | AA | AA | AB | AB | AB | AA | AB | AA | AA | AB | AA | AB | AB | AA | [illegible] |
| AA | AB | AB | AA | AA | AB | AB | AB | AA | AA | AA | AB | AA | AB | AA | AA | AA | [illegible] |
| AA | AB | AA | AB | AB | AB | AB | AA | AB | AA | AB | AB | AB | AA | AA | AA | AB | [illegible] |
| AA | AB | AB | AB | AA | AB | AA | AB | AA | AA | AA | AB | AB | AA | AA | AB | AB | [illegible] |
| AA | AA | AA | AA | AB | AB | AB | AB | AA | AB | AB | AB | AB | AA | AA | AB | AB | [illegible] |
| AA | AA | AA | AB | AB | AA | AA | AA | AB | AB | AB | AA | AA | AA | AB | AB | AB | [illegible] |
| - | AA | AB | AB | AA | - | AB | AB | AB | AA | AB | AA | AB | - | AA | - | - | [illegible] |
| AA | AA | AA | AA | AA | AB | AB | AB | AA | AB | AA | AA | AB | AA | AB | AB | AA | [illegible] |
| AA | AA | AA | AB | AB | AB | AB | AB | AA | AA | AB | AB | AB | AA | AA | AA | AB | [illegible] |
| AA | AA | AA | AA | AB | AB | AB | AB | AB | AB | AB | AA | AB | AB | AB | AB | AA | [illegible] |
| AB | AB | AB | AB | AA | AB | AB | AB | AA | AB | AA | AB | AB | AA | AB | AB | AA | [illegible] |
| AA | AB | AA | AA | AB | AB | AB | AB | AA | AA | AB | AB | AA | - | AB | AA | - | [illegible] |
| AB | AA | AB | AA | AA | AB | AB | AB | AA | AB | AA | AA | AB | AA | AB | AB | AA | [illegible] |
| AB | AA | AA | AA | AB | AB | AB | AB | AB | AB | AA | AB | AB | AB | AB | AB | AA | [illegible] |
| AA | AA | AB | AA | AA | AA | AA | AA | AA | AA | AA | AB | AA | AB | AA | AA | AA | [illegible] |
| AA | AB | AB | AA | AA | AB | AB | AB | AB | AB | AA | AB | AA | AA | AA | AA | AA | [illegible] |
| AA | AB | AB | AB | AB | AB | AB | AB | AA | AB | AB | AA | AA | AB | AA | AA | AB | [illegible] |
| AA | AB | AB | AB | AB | AA | AA | AA | AB | AB | AB | AA | AA | AA | - | AA | AB | [illegible] |
| AA | AB | AB | AB | AA | - | AA | AA | AA | AB | AB | AB | AB | - | AB | AB | AA | [illegible] |
| AA | AB | AA | AA | AB | AB | AB | AB | AA | AA | AB | AB | AB | AA | AA | AA | AB | [illegible] |
| AB | AB | AB | AA | AA | AA | AA | AA | AA | AB | AA | AB | AA | AB | AB | AB | AA | [illegible] |
| AB | AB | AA | AB | AA | - | AB | AB | AB | AB | AA | AB | AB | - | - | AB | - | [illegible] |
| AA | AB | AB | AA | AB | - | AB | AB | AA | AA | AA | AB | AA | - | - | AA | AB | [illegible] |
| AA | AB | AB | AB | AB | - | AA | AA | AB | AA | AB | AB | AA | - | AA | AA | AB | [illegible] |
| AB | AA | AA | AA | AA | AA | AA | AA | AA | AA | AA | AA | AA | - | AA | AA | - | [illegible] |
| AB | AB | AA | AB | AB | AB | AB | AB | AB | AB | AB | AB | AB | AB | AB | AB | AB | [illegible] |
| AB | AA | AA | AB | AB | AA | AA | AB | AA | AB | AB | AA | AA | AA | AB | AB | AB | [illegible] |
| AB | AA | AB | AA | AA | AA | AA | AA | AB | AB | AA | AA | AB | AA | AB | AB | AA | [illegible] |
| AA | AA | AA | AA | AA | - | AA | AA | AA | AA | AA | AA | AA | - | AA | AA | - | [illegible] |
| AA | AA | AB | AA | AA | AA | AA | AA | AB | AB | AA | AA | AA | AB | AB | AA | AA | [illegible] |
| AB | AA | AA | AA | AA | AA | AA | AA | AA | AA | AA | AA | AA | AA | AB | AA | AA | [illegible] |
| AB | AA | AA | AB | AB | AA | AA | AA | AA | AA | AB | AA | AA | AA | AA | AA | AB | [illegible] |
| AB | AB | AA | AB | AB | AA | AA | AA | AA | AA | AA | AA | AA | AA | AA | AA | AA | [illegible] |
| AA | AB | AA | AB | AB | AB | AB | AB | AB | AB | AB | AB | AA | AB | AA | AA | AA | [illegible] |
| AA | AB | AB | AB | AA | AA | AA | AA | AA | AA | AA | AA | AA | AA | AA | AA | AA | [illegible] |
| [illegible] | [illegible] | [illegible] | [illegible] | [illegible] | [illegible] | [illegible] | [illegible] | [illegible] | [illegible] | [illegible] | [illegible] | [illegible] | [illegible] | [illegible] | [illegible] | [illegible] | [illegible] |

| snp_B5490 | snp_B5499 | snp_B5505 | snp_B5528 | snp_B5535 | snp_B5559 | snp_B5576 | snp_B5577 | snp_B5589 | snp_B5594 | snp_B5607 | snp_B5625 | snp_B5643 | snp_B5644 | snp_B5677 | snp_B5696 | snp_B5705 |
|---|---|---|---|---|---|---|---|---|---|---|---|---|---|---|---|---|
| 9 | 9 | 9 | 9 | 9 | 9 | 9 | 9 | 9 | 9 | 9 | 9 | 9 | 9 | 9 | 9 | 9 |
| -- | -- | -- | -- | -- | -- | -- | -- | -- | -- | LG2 | -- | -- | LG2 | -- | -- | -- |
| AB | AB | AA | AA | AB | AB | AB | AA | AB | AA | AB | AA | AB | AB | AA | AA | AB |
| AB | AB | AA | AA | AB | AB | AA | AB | AB | AB | AB | AB | AB | AA | AA | AA | AB |
| AA | AB | AA | AA | AB | AA | AA | AA | AB | AA | AA | AA | AA | AB | AA | AB | AB |
| AB | AB | AB | AA | AB | AA | AB | AB | AB | AA | AB | AA | AB | AB | AA | AA | AB |
| AA | AB | AA | AA | AB | AA | AA | AB | AA | AB | AA | AA | AB | AB | AA | AA | AA |
| AA | AA | AB | - | AA | AB | AA | AA | AA | - | AA | AB | AA | AA | AB | AA | AA |
| AB | AB | AB | AB | AB | AA | AB | AB | AB | AB | AB | AA | AB | AB | AB | AB | AA |
| AB | AA | AB | AB | AA | AB | AB | AA | AA | AA | AA | AB | AA | AB | AB | AB | AB |
| AB | AA | AB | AA | AA | AB | AA | AA | AA | AA | AA | AA | AA | AB | AA | AB | AB |
| AA | AA | AB | AA | AB | AA | AB | AA | AA | AA | AA | AB | AA | AA | AB | AA | AB |
| AA | AB | AB | AB | AB | AB | AB | AA | AB | AB | AB | AB | AA | AA | AA | AB | AB |
| AA | AA | AA | AB | AA | AA | AA | AA | AB | AA | AB | AA | AA | AA | AB | AB | AA |
| AA | AB | AA | AA | AB | AB | AA | AB | AB | AB | AB | AB | AB | AA | AA | AA | AB |
| AB | AA | AA | AA | AB | AA | AB | AA | AA | AB | AB | AB | AA | AA | AA | AB | AA |
| AA | AB | AA | AA | AA | AA | AB | AB | AB | AA | AB | AB | AB | AB | AA | AA | AB |
| AA | AB | AA | AB | AA | AB | AA | AA | AB | AB | AB | AB | AA | AB | AA | AB | AA |
| AA | AA | AB | AB | AB | AB | AB | AA | AA | AA | AA | AB | AA | AA | AB | AA | AA |
| AB | AA | - | - | AB | AA | - | - | AB | AB | AB | - | AA | - | AA | AA | AA |
| AA | AA | AA | AB | AA | AA | AA | AA | AB | AA | AB | AA | AA | AA | AB | AB | AA |
| AA | AB | AA | AA | AB | AA | AA | AB | AB | AA | AB | AA | AA | AB | AA | AB | AA |
| AA | AA | AB | AB | AB | AA | AB | AA | AA | AA | AA | AB | AA | AA | AB | AA | AA |
| AB | AB | AA | AB | AB | AB | AB | AB | AA | AB | AA | AB | AB | AB | AB | AA | AB |
| AA | AB | AB | AA | AB | AB | AB | - | AA | AB | AB | - | - | - | AA | AB | - |
| AA | AA | AA | AB | AA | AB | AA | AA | AA | AA | AA | AB | AA | AA | AB | AA | AA |
| AA | AA | AA | AB | AA | AB | AB | AA | AB | AA | AB | AA | AA | AA | AB | AB | AA |
| AA | AB | AA | AA | AB | AB | AA | AA | AA | AB | AA | AA | AB | AB | AA | AA | AB |
| AA | AB | AA | AA | AB | AB | AA | AB | AB | AA | AB | AB | AB | AB | AA | AA | AB |
| AB | AB | AA | AA | AB | AB | AA | AB | AB | AB | AB | AB | AB | AA | AA | AA | AB |
| AB | AA | AA | AA | AB | AB | AA | AA | AB | AB | AB | AA | AA | AA | AB | AA | AB |
| AB | AA | AA | AB | AA | AA | AA | AB | AB | - | AB | AA | AB | AB | AB | AA | AB |
| AB | AB | AA | AA | AB | AB | AA | AB | AB | AB | AB | AB | AB | AB | AA | AB | AA |
| AB | AB | AA | AB | AA | AA | AA | AB | AA | AB | AA | AA | AB | AB | AB | AA | AB |
| AB | AB | AB | AB | AA | AA | AB | AB | AB | AB | AB | AA | AA | - | AB | AB | AA |
| AA | AB | AA | AA | AB | AB | AA | AB | AB | AB | AB | AB | AB | AB | AA | AB | AB |
| AB | AB | AB | AA | AB | AB | AB | - | AA | AB | AB | - | AA | AB | AA | AA | AB |
| AA | AA | AA | AA | AA | AA | AA | AA | AA | AA | AA | AA | AA | AA | AA | AA | AA |
| AB | AB | AB | AB | AB | AB | AB | AB | AB | AB | AB | AA | AB | AB | AB | AB | AA |
| AA | AA | AB | AB | AA | AB | AB | AA | AA | AA | AA | AA | AA | AA | AB | AA | AA |
| AA | AB | AB | AB | AA | AB | AB | AA | AB | AB | AA | AB | AA | AA | AB | AA | AA |
| AA | AA | AB | AA | AA | AA | AB | AA | AA | AA | AA | AA | - | AA | AA | AA | AA |
| AA | AA | AB | AA | AB | AB | AB | AB | AA | AB | AA | AA | AA | AA | AB | AA | AB |
| AA | AA | AA | AA | AA | AB | AA | AA | AA | AA | AA | AA | AA | AA | AA | AA | AB |
| AA | AA | AA | AA | AB | AB | AA | AA | AA | AA | AA | AB | AA | AA | AA | AA | AA |
| AA | AA | AA | AA | AA | AB | AA | AA | AA | AB | AA | AB | AA | AA | AA | AA | AB |
| AB | AB | AB | AA | AB | AB | AB | AA | AA | AB | AB | AA | AA | AA | AA | AA | AB |
| AB | AB | AA | AA | AA | AB | AA | AB | AA | AB | AA | AB | AB | AA | AA | AA | AB |

| snp_B1250 | snp_B1251 | snp_B1278 | snp_B1293 | snp_B1306 | snp_B1326 | snp_B1342 | snp_B1363 | snp_B1366 | snp_B1401 | snp_B1421 | snp_B1447 | snp_B1448 | snp_B1453 | snp_B1515 | snp_B1525 | snp_B1537 | snp_B1550 | snp_B1579 | snp_B1580 | snp_B1583 | snp_B1587 | snp_B1651 | snp_B1678 | snp_B1701 | snp_B1728 | snp_B1731 |
|---|---|---|---|---|---|---|---|---|---|---|---|---|---|---|---|---|---|---|---|---|---|---|---|---|---|---|
| 10 | 10 | 10 | 10 | 10 | 10 | 10 | 10 | 10 | 10 | 10 | 10 | 10 | 10 | 10 | 10 | 10 | 10 | 10 | 10 | 10 | 10 | 10 | 10 | 10 | 10 | 10 |
| -- | LG4 | -- | -- | -- | -- | -- | -- | -- | -- | -- | -- | -- | -- | -- | -- | -- | LG4 | -- | LG4 | -- | -- | -- | -- | -- | -- | -- |
| AB | AA | AA | AB | AA | AA | AA | AA | AB | AB | AB | AA | AB | AA | AA | AA | AB | AA | AB | AB | AA | AB | AB | AB | AA | AB | AB |

| snp_B1733 | snp_B1735 | snp_B1785 | snp_B1823 | snp_B1844 | snp_B1850 | snp_B1852 | snp_B1856 | snp_B1857 | snp_B1878 | snp_B1899 | snp_B1900 | snp_B1905 | snp_B1910 | snp_B1918 | snp_B1921 | snp_B1945 | snp_B1954 | snp_B1960 | snp_B1963 | snp_B1971 | snp_B1993 | snp_B2002 | snp_B2080 | snp_B2094 | snp_B2114 | snp_B2156 |
|---|---|---|---|---|---|---|---|---|---|---|---|---|---|---|---|---|---|---|---|---|---|---|---|---|---|---|
| 10 | 10 | 10 | 10 | 10 | 10 | 10 | 10 | 10 | 10 | 10 | 10 | 10 | 10 | 10 | 10 | 10 | 10 | 10 | 10 | 10 | 10 | 10 | 10 | 10 | 10 | 10 |
| -- | -- | -- | -- | -- | -- | -- | -- | -- | -- | -- | -- | -- | -- | -- | -- | -- | -- | LG4 | -- | -- | -- | -- | -- | -- | -- | -- |
| AB | AA | AA | AA | AB | AA | AB | AA | AA | AB | AA | AA | AA | AB | AA | AA | AB | AB | AA | AA | AA | AA | AA | AB | AA | AB | AB |

| snp_B2158 | snp_B2169 | snp_B2175 | snp_B2184 | snp_B2188 | snp_B2204 | snp_B2215 | snp_B2231 | snp_B2236 | snp_B2272 | snp_B2288 | snp_B2289 | snp_B2299 | snp_B2306 | snp_B2307 | snp_B2338 | snp_B2352 | snp_B2380 | snp_B2395 | snp_B2397 | snp_B2422 | snp_B2427 | snp_B2440 |
|---|---|---|---|---|---|---|---|---|---|---|---|---|---|---|---|---|---|---|---|---|---|---|
| 10 | 10 | 10 | 10 | 10 | 10 | 10 | 10 | 10 | 10 | 10 | 10 | 10 | 10 | 10 | 10 | 10 | 10 | 10 | 10 | 10 | 10 | 10 |
| -- | -- | -- | -- | -- | LG4 | -- | -- | LG4 | -- | -- | -- | -- | -- | -- | -- | -- | -- | -- | -- | -- | -- | -- |
| AA | AB | AB | AA | AA | AA | AA | AA | AA | AA | AA | AB | AA | AA | AA | AB | AB | AA | AB | AB | AB | AA | AB |

| snp_B2457 | snp_B2499 | snp_B2503 | snp_B2504 | snp_B2518 | snp_B2528 | snp_B2532 | snp_B2566 | snp_B2589 | snp_B2629 | snp_B2671 | snp_B2673 | snp_B2676 | snp_B2684 | snp_B2710 | snp_B2720 | snp_B2723 | snp_B2796 | snp_B2803 | snp_B2816 | snp_B2836 | snp_B2841 | snp_B2853 | snp_B2861 | snp_B2866 | snp_B2885 | snp_B2919 | snp_B2979 | snp_B3003 | snp_B3031 | snp_B3047 | snp_B3068 | snp_B3076 | snp_B3079 | snp_B3118 | snp_B3122 | snp_B3173 | snp_B3275 | snp_B3299 | snp_B3300 | snp_B3336 | snp_B3369 | snp_B3396 | snp_B3403 | snp_B3422 | snp_B3425 | snp_B3504 | snp_B3617 | snp_B3619 | snp_B3648 | snp_B3650 | snp_B3661 | snp_B3667 | snp_B3668 | snp_B3673 | snp_B3674 | snp_B3699 | snp_B3716 | snp_B3717 | snp_B3743 | snp_B3745 | snp_B3770 | snp_B3786 | snp_B3822 | snp_B3855 | snp_B3865 | snp_B3880 | snp_B4006 | snp_B4067 | snp_B4088 | snp_B4107 | snp_B4121 | snp_B4125 | snp_B4147 | snp_B4171 | snp_B4230 | snp_B4276 |
|---|---|---|---|---|---|---|---|---|---|---|---|---|---|---|---|---|---|---|---|---|---|---|---|---|---|---|---|---|---|---|---|---|---|---|---|---|---|---|---|---|---|---|---|---|---|---|---|---|---|---|---|---|---|---|---|---|---|---|---|---|---|---|---|---|---|---|---|---|---|---|---|---|---|---|---|---|
| 10 | 10 | 10 | 10 | 10 | 10 | 10 | 10 | 10 | 10 | 10 | 10 | 10 | 10 | 10 | 10 | 10 | 10 | 10 | 10 | 10 | 10 | 10 | 10 | 10 | 10 | 10 | 10 | 10 | 10 | 10 | 10 | 10 | 10 | 10 | 10 | 10 | 10 | 10 | 10 | 10 | 10 | 10 | 10 | 10 | 10 | 10 | 10 | 10 | 10 | 10 | 10 | 10 | 10 | 10 | 10 | 10 | 10 | 10 | 10 | 10 | 10 | 10 | 10 | 10 | 10 | 10 | 10 | 10 | 10 | 10 | 10 | 10 | 10 | 10 | 10 | 10 |
| -- | -- | -- | -- | -- | -- | -- | -- | LG4 | -- | LG4 | -- | LG4 | -- | LG4 | -- | -- | LG4 | -- | -- | -- | LG4 | -- | -- | -- | -- | -- | LG4 | -- | LG4 | LG4 | -- | -- | -- | -- | -- | -- | -- | -- | -- | -- | -- | -- | LG4 | -- | -- | -- | -- | -- | -- | -- | -- | -- | LG4 | -- | LG4 | -- | -- | -- | -- | -- | -- | -- | -- | -- | -- | -- | -- | -- | LG4 | -- | -- | -- | -- | -- | -- | -- |
| AB | AA | AB | AA | AA | AB | AB | AB | AA | AA | AA | AB | AB | AA | AA | AA | AA | AB | AA | AA | AB | AA | AA | AA | AA | AB | AB | AB | AB | AA | AA | AA | AB | AB | AB | AA | AA | AA | AB | AA | AA | AA | AA | AA | AB | AB | AB | AA | AA | AB | AA | AA | AA | AA | AA | AA | AB | AA | AB | AA | AA | AA | AB | AB | AB | AA | AB | AB | AA | AA | AB | AA | AA | AB | AB | AA | AA |
| AA | AA | AB | AA | AB | AB | AA | AA | AA | AA | AA | AA | AA | AB | AA | AA | AA | AA | AA | AA | AA | AA | AA | AB | AA | AA | AA | AA | AA | AA | AA | AB | AA | AA | AB | AA | AA | AA | AB | AA | AA | AA | AA | AA | AA | AB | AB | AA | AA | AA | AA | AA | AB | AA | AA | AA | AA | AA | AA | AA | AA | AB | AA | AA | AA | AA | AA | AA | AA | AA | AA | AA | AA | AA | AB | AA | AA |
| AB | AB | AB | AB | AB | AB | AA | AA | AA | AB | AA | AB | AB | AB | AA | AA | AB | AA | AA | AA | AA | AA | AA | AB | AA | AA | AB | AA | AB | AA | AA | AB | AA | AA | AB | AA | AA | AB | AB | AA | AA | AA | AA | AA | AA | AB | AB | AA | AA | AA | AB | AB | AA | AA | AA | AA | AA | AB | AB | AA | AA | AB | AA | AB | AA | AB | AA | AB | AB | AA | AB | AA | AA | AB | AA | AB | AA |
| AB | AB | AA | AB | AA | AA | AB | AB | AB | AA | AB | AA | - | AB | AB | AB | AB | AB | AB | AB | AB | AB | - | - | AB | AA | AA | AB | AB | AB | AB | AA | AB | AA | AA | AB | AB | AA | AA | AB | AB | AB | AA | AB | AB | AA | AA | AB | AB | AB | AB | AA | AA | AA | AB | AA | AB | AA | AB | - | - | - | AB | AB | AB | AB | AA | AB | AB | AB | AB | AB | AB | AB | AB | AA | AB |
| AB | AB | AB | AB | AA | AB | AB | AA | AB | AB | AB | AA | AB | AB | AB | AB | AB | AB | AB | AB | AB | AB | AB | AB | AB | AA | AA | AB | AB | AB | AB | AB | AA | AB | AA | AB | AB | AB | AB | AA | AA | AB | AA | AB | AB | AB | AB | AB | AB | AB | AB | AA | AB | AB | AB | AB | AA | AB | AA | AA | AB | AB | AA | AB | AB | AB | AB | AB | AB | AB | AB | AB | AB | AA | AB | AB | AB |
| AB | AA | AA | - | AB | AA | AB | - | AA | AA | AA | AA | - | AB | AB | - | AB | - | - | AB | AA | - | - | - | AA | AA | AB | AA | AB | - | - | AB | AA | AB | AA | - | AB | - | - | - | AB | - | AB | AA | - | - | AA | AB | AA | AB | AB | AA | AA | AB | AA | - | AA | - | - | - | - | AB | AA | - | AA | AA | AA | AB | AB | - | - | AB | AA | - | AA | - | AB |
| AA | AB | AA | AA | AB | AB | AB | AB | AB | AB | AB | AA | AB | AB | AB | AA | AA | AB | AB | AA | AB | AB | - | AB | AB | AB | AB | AA | AA | AB | AB | AB | AB | AB | AB | AB | AB | AA | AB | AB | AB | AB | AA | AB | AA | AB | AB | AB | AB | AB | AA | AA | AB | AB | AA | AB | AB | AA | AB | AB | AB | AA | AB | AA | AB | AB | AA | AA | AB | AB | AB | AB | AA | AB | AA | AA | AB |
| AA | AA | AA | AA | AB | AA | AA | AA | AA | AB | AA | AB | AA | AA | AA | AA | AA | AA | AA | AA | AA | AA | AB | AB | AA | AB | AB | AA | AA | AA | AA | AB | AA | AA | AB | AA | AA | AA | AB | AA | AB | AA | AA | AA | AA | AA | AB | AA | AA | AA | AA | AB | AA | AA | AA | AA | AA | AA | AB | AA | AA | AA | AA | AA | AA | AA | AB | AA | AA | AA | AA | AA | AA | AB | AB | AA | AB |
| AA | AB | AB | AA | AB | AB | AB | AB | AA | AB | AA | AB | AA | AA | AA | AA | AB | AA | AB | AA | AA | AA | AA | AA | AB | AA | AB | AB | AB | AA | AA | AA | AB | AA | AA | AB | AB | AB | AA | AA | AA | AA | AA | AA | AB | AB | AA | AB | AA | AB | AA | AB | AB | AA | AB | AA | AB | AA | AB | AA | AA | AB | AA | AA | AA | AB | AA | AB | AB | AA | AA | AA | AB | AB | AA | AB | AA |
| AA | AA | AA | AB | AA | AA | AA | AB | AB | AA | AB | AB | AB | AA | AB | AA | AA | AB | AB | AA | AA | AB | AB | AA | AB | AB | AB | AB | AB | AB | AB | AB | AB | AA | AB | AA | AB | AA | AA | AA | AB | AA | AB | AB | AB | AA | AB | AB | AB | AA | AB | AA | AB | AA | AA | AA | AB | AB | AB | AB | AB | AB | AB | AA | AA | AB | AB | AA | AA | AB | AA | AA | AA | AB | AB | AB | AA |
| AB | AA | AB | AB | AB | AA | AB | AA | AA | AA | AA | AB | AA | AA | AA | AA | AA | AB | AA | AA | AA | AA | AB | AA | AB | AB | AA | AA | AA | AA | AA | AA | AA | AA | AA | AA | AB | AA | AA | AB | AB | AA | AA | AA | AB | AB | AB | AA | AA | AA | AA | AB | AA | AA | AB | AA | AA | AA | AB | AB | AA | AB | AA | AA | AA | AB | AA | AA | AA | AA | AA | AA | AA | AB | AA | AB | AA |
| AB | AB | AB | AB | AA | AB | AB | AB | AB | AB | AB | AA | AB | AB | AB | AA | AB | AA | AB | AA | AB | AB | AB | AB | AB | AB | AB | AB | AB | AB | AB | AB | AA | AB | AA | AB | AB | AB | AB | AB | AA | AB | AB | AB | AB | AB | AB | AB | AB | AB | AB | AB | AB | AB | AB | AB | AA | AA | AA | AA | AB | AB | AA | AB | AB | AB | AA | AB | AB | AB | AB | AB | AB | AA | AA | AB | AB |
| AB | AA | AB | AB | AB | AB | AA | AA | AA | AB | AA | AB | AB | AB | AB | AB | AB | AA | AA | AB | AA | AA | AB | AB | AA | AB | AB | AA | AB | AA | AA | AB | AA | AB | AB | AB | AA | AB | AB | AB | AB | AA | AA | AA | AB | AB | AB | AA | AB | AA | AB | AB | AB | AB | AA | AB | AA | AA | AB | AA | AA | AB | AA | AA | AB | AA | AB | AB | AB | AA | AB | AB | AB | AB | AB | AB | AB |
| AA | AA | AB | AB | AB | AA | AB | AA | AA | AB | AA | AB | AA | AA | AA | AA | AA | AB | AA | AA | AA | AA | AA | AA | AA | AA | AA | AA | AA | AA | AA | AA | AA | AA | AA | AA | AB | AB | AB | AA | AA | AA | AA | AA | AB | AB | AB | AA | AA | AB | AB | AB | AA | AA | AA | AA | AA | AA | AA | AA | AB | AB | AA | AB | AA | AA | AB | AA | AB | AA | AA | AA | AA | AB | AB | AB | AA |
| AB | AB | AA | AA | AA | AB | AA | AB | AB | AA | AB | AA | AB | AB | AB | AA | AA | AB | AB | AA | AB | AB | AB | AA | AB | AA | AB | AB | AA | AB | AB | AA | AB | AB | AB | AA | AA | AB | AB | AA | AA | AB | AB | AB | AA | AA | AA | AA | AB | AA | AA | AB | AB | AB | AB | AB | AB | AA | AB | AB | AB | AB | AB | AA | AA | AB | AB | AA | AA | AB | AA | AB | AA | AA | AB | AA | AA |
| AA | AB | AB | AA | AB | AB | AB | AB | AB | AA | AB | AB | AA | AB | AB | AA | AB | AB | AB | AA | AB | AB | AB | AB | AB | AA | AB | AB | AA | AB | AB | AB | AA | AA | AB | AA | AB | AA | AB | AA | AA | AB | AA | AB | AA | AB | AB | AB | AB | AB | AA | AA | AB | AA | AB | AB | AB | AB | AA | AB | AB | AB | AB | AB | AB | AB | AB | AA | AA | AB | AA | AA | AA | AA | AB | AB | AA |
| AB | AA | AB | AB | AB | AA | AB | AB | AA | AA | AA | AB | AA | AA | AA | AA | AB | AA | AA | AA | AA | AA | AB | AA | AB | AA | AB | AB | AB | AA | AA | AA | AA | AB | AA | AA | AA | AB | AA | AA | AB | AB | AA | AA | AB | AA | AB | AB | AB | AB | AB | AB | AA | AA | AB | AA | AB | AA | AB | AB | AA | AB | AB | AB | AA | AB | AA | AB | AA | AA | AA | AA | AA | AB | AB | AB | AA |
| AB | AB | AA | - | AA | AB | AB | - | AB | - | AB | AA | - | AA | AA | - | AA | - | - | AB | AB | - | - | - | AA | AB | AA | AA | AA | - | - | - | AB | AB | AB | AB | AA | - | - | - | AB | - | - | AB | - | AA | AA | AB | AA | AA | AA | AB | AB | AB | AA | - | AA | - | - | - | AB | AA | AB | AA | AA | AA | AA | AA | AA | - | - | - | AB | AA | AB | AA | AB |
| AB | AB | AB | AB | AB | AB | AB | AB | AB | AB | AB | AA | AB | AB | AB | AB | AB | AB | AB | AB | AB | AB | AB | AB | AB | AB | AB | AB | AB | AB | AB | AB | AB | AB | AB | AB | AB | AB | AB | AB | AB | AB | AB | AB | AB | AB | AB | AB | AB | AB | AB | AB | AB | AB | AB | AA | AB | AB | AB | AB | AB | AB | AB | AB | AB | AB | AB | AB | AB | AB | AB | AB | AB | AB | - | AB | AB |
| AA | AA | AB | AA | AB | AA | AB | AB | AA | AB | AA | AA | AB | - | AA | AB | AB | AA | - | AB | AA | AA | - | AA | AA | AB | AA | AA | AA | - | AA | AA | AA | AA | AA | AB | AA | AB | AA | AB | AA | AB | AA | AA | AB | AA | AA | AA | AA | AB | AA | AA | AA | AB | AB | AB | AB | AA | - | AB | AA | AB | AB | AB | AA | AA | AB | AB | AA | AA | - | AB | AB | AA | AB | AB | AB |
| AB | AB | AB | AB | AB | AB | AB | AB | AB | AB | AB | AB | AB | AB | AB | AB | AB | AB | AB | AB | AB | AB | AB | AB | AB | AB | AB | AB | AB | AB | AB | AB | AB | AB | AB | AB | AB | AB | AB | AB | AB | AB | AB | AB | AB | AB | AB | AB | AB | AB | AB | AB | AB | AB | AB | AB | AB | AA | AB | AB | AB | AB | AB | AB | AB | AB | AB | AB | AB | AB | AA | AB | AB | AB | AB | AB | AB |
| AB | AB | AB | AA | AA | AB | AB | AA | AB | AA | AA | AA | AB | AB | AB | AB | AB | AA | AB | AB | AB | AB | AA | AA | AB | AA | AB | AA | AB | AA | AB | AA | AA | AB | AA | AB | AA | AB | AB | AA | AB | AA | AB | AB | AB | AB | AA | AB | AB | AA | AB | AA | AB | AB | AB | AB | AA | AB | AA | AA | AB | AB | AA | AB | AA | AB | AA | AB | AB | AA | AA | AA | AA | AA | AA | AA | AB |
| AB | AB | AA | - | AB | AB | - | AB | AB | - | AB | AA | - | - | AB | AB | AA | - | AB | AB | - | AB | - | - | AA | AA | AA | AB | AA | - | AB | - | AB | AA | AB | - | AB | - | - | AB | AA | AB | AA | AB | AB | - | AB | AB | AB | AB | AB | AA | AB | AA | - | - | AB | - | - | - | - | - | AB | AA | AB | AA | AA | AA | AB | - | - | - | AA | AA | - | AA | AB |
| AB | AB | AB | AB | AA | AB | AB | AB | AB | AB | AB | AA | AB | AB | AB | AB | AB | AB | AB | AB | AB | AB | AB | AA | AB | AB | AB | AB | AB | AB | AB | AA | AB | AB | AB | AB | AB | AB | AB | AB | AB | AB | AB | AB | AB | AB | AB | AB | AB | AA | AB | AB | AB | AB | AB | AB | AB | AA | AB | AA | AB | AB | AB | AB | AB | AB | AB | AB | AB | AB | AB | AB | AB | AB | AA | AB | AB |
| AA | AB | AB | AB | AB | AB | AB | AA | AA | AB | AA | AB | AA | AA | AA | AA | AA | AA | AB | AA | AA | AA | AB | AB | AA | AB | AA | AA | AA | AA | AA | AB | AA | AA | AA | AA | AB | AA | AB | AB | AA | AA | AB | AA | AA | AB | AA | AB | AB | AB | AA | AB | AB | AA | AB | AA | AA | AB | AB | AB | AB | AA | AA | AA | AB | AB | AB | AA | AA | AA | AA | AA | AB | AB | AA | AB | AA |
| AB | AA | AB | AA | AA | AA | AA | AA | AA | AA | AA | AB | AA | AB | AA | AA | AB | AA | AA | AA | AA | AA | AA | AA | AB | AB | AB | AA | AB | AA | AA | AA | AA | AA | AA | AA | AA | AB | AB | AB | AB | AA | AB | AA | AB | AB | AA | AB | AA | AB | AB | AB | AA | AA | AA | AA | AA | AA | AA | AB | AA | AB | AA | AB | AA | AB | AA | AB | AA | AA | AB | AA | AA | AB | AA | AB | AA |
| AB | AB | AB | AB | AB | AB | AB | AA | AB | AB | AB | AB | AB | AB | AB | AB | AB | AA | AB | AB | AB | AB | AB | AB | AB | AA | AB | AA | AB | AB | AB | AB | AA | AB | AA | AB | AB | AB | AB | AB | AA | AB | AB | AB | AB | AB | AB | AB | AB | AB | AB | AB | AB | AB | AB | AB | AA | AB | AA | AA | AB | AB | AA | AB | AB | AB | AA | AB | AB | AB | AB | AB | AA | AB | AB | AB | AB |
| AA | AB | AB | AB | AA | AB | AB | AA | AB | AA | AB | AB | AB | AB | AB | AA | AB | AB | AB | AB | AB | AB | AA | AA | AB | AB | AB | AB | AB | AB | AB | AA | AA | AB | AA | AA | AA | AA | AB | AA | AA | AA | AA | AB | AB | AB | AA | AB | AB | AB | AB | AB | AB | AA | AB | AA | AA | AB | AB | AA | AB | AA | AA | AB | AB | AB | AA | AB | AA | AA | AB | AB | AA | AB | AB | AB | AA |
| AA | AA | AA | AB | AB | AB | AA | AA | AB | AB | AB | AA | - | AA | AB | AB | AB | AA | AA | AB | AB | AB | AA | AA | AB | AA | AA | AB | AB | AB | AB | AA | AA | AB | AA | AB | AB | AB | AA | AA | AB | AB | AB | AB | AA | AB | AA | AB | AA | AB | AA | AB | AA | AB | AA | - | AA | AB | AA | AB | - | AA | AA | AA | AB | AB | AB | AA | AB | AB | - | AA | AB | AA | AB | AA | AB |
| AB | AB | AB | - | AA | AB | - | - | AA | AA | AA | - | - | - | AA | AB | AA | - | AB | AB | AA | - | AB | - | AA | AB | AA | AA | AA | AA | AA | AA | AA | AB | AA | AB | AB | - | - | AB | AB | AB | AB | AA | - | AB | AA | AB | AB | AB | AA | AB | AA | AA | AB | - | AA | AB | AB | AA | - | AB | AA | AA | AB | AB | AB | AA | AA | - | AA | AA | AA | AB | - | AB | AB |
| [illegible] | [illegible] | [illegible] | [illegible] | [illegible] | [illegible] | [illegible] | [illegible] | [illegible] | [illegible] | [illegible] | [illegible] | [illegible] | [illegible] | [illegible] | [illegible] | [illegible] | [illegible] | [illegible] | [illegible] | [illegible] | [illegible] | [illegible] | [illegible] | [illegible] | [illegible] | [illegible] | [illegible] | [illegible] | [illegible] | [illegible] | [illegible] | [illegible] | [illegible] | [illegible] | [illegible] | [illegible] | [illegible] | [illegible] | [illegible] | [illegible] | [illegible] | [illegible] | [illegible] | [illegible] | [illegible] | [illegible] | [illegible] | [illegible] | [illegible] | [illegible] | [illegible] | [illegible] | [illegible] | [illegible] | [illegible] | [illegible] | [illegible] | [illegible] | [illegible] | [illegible] | [illegible] | [illegible] | [illegible] | [illegible] | [illegible] | [illegible] | [illegible] | [illegible] | [illegible] | [illegible] | [illegible] | [illegible] | [illegible] | [illegible] | [illegible] | [illegible] |

| snp_B4283 | snp_B4290 | snp_B4300 | snp_B4310 | snp_B4324 | snp_B4325 | snp_B4333 | snp_B4341 | snp_B4342 | snp_B4346 | snp_B4350 | snp_B4360 | snp_B4363 | snp_B4395 | snp_B4412 | snp_B4472 | snp_B4490 | snp_B4545 | snp_B4566 | snp_B4569 | snp_B4588 | snp_B4603 | snp_B4604 | snp_B4612 | snp_B4613 | snp_B4618 | snp_B4643 | snp_B4665 | snp_B4668 | snp_B4678 | snp_B4716 | snp_B4744 | snp_B4748 | snp_B4775 | snp_B4806 | snp_B4818 | snp_B4845 | snp_B4927 | snp_B4930 | snp_B4932 | snp_B4946 | snp_B4971 | snp_B4979 |
|---|---|---|---|---|---|---|---|---|---|---|---|---|---|---|---|---|---|---|---|---|---|---|---|---|---|---|---|---|---|---|---|---|---|---|---|---|---|---|---|---|---|---|
| 10 | 10 | 10 | 10 | 10 | 10 | 10 | 10 | 10 | 10 | 10 | 10 | 10 | 10 | 10 | 10 | 10 | 10 | 10 | 10 | 10 | 10 | 10 | 10 | 10 | 10 | 10 | 10 | 10 | 10 | 10 | 10 | 10 | 10 | 10 | 10 | 10 | 10 | 10 | 10 | 10 | 10 | 10 |
| -- | -- | -- | -- | -- | -- | -- | -- | -- | -- | -- | -- | LG4 | -- | -- | -- | -- | -- | -- | -- | -- | -- | -- | -- | -- | -- | -- | -- | -- | LG4 | LG4 | LG4 | -- | -- | -- | -- | -- | -- | -- | -- | -- | -- | -- |
| AB | AA | AB | AA | AB | AA | AA | AA | AB | AB | AA | AA | AA | AB | AA | AA | AB | AB | AA | AA | AB | AA | AB | AB | AB | AB | AB | AB | AB | AB | AA | AA | AB | AA | AA | AB | AA | AB | AA | AB | AB | AA | AA |
| AA | AB | AA | AA | AA | AA | AA | AA | AA | AA | AB | AA | AA | AA | AA | AA | AA | AA | AA | AA | AA | AA | AA | AB | AA | AB | AB | AA | AA | AA | AA | AA | AB | AA | AA | AA | AA | AA | AA | AA | AA | AA | AB |
| AB | AB | AA | AB | AB | AA | AB | AB | AA | AB | AB | AA | AB | AB | AB | AB | AA | AA | AB | AA | AB | AA | AA | AB | AA | AB | AB | AB | AB | AA | AB | AA | AB | AA | AA | AB | AA | AA | AB | AB | AA | AA | AB |
| - | AA | AA | AB | AA | AB | AA | AB | AB | AB | AA | AB | AB | AB | AA | AB | AA | AB | AA | AB | AA | AB | AA | AA | AA | AB | AA | AB | AB | AB | AB | AB | AB | AB | AB | AB | AA | AA | AA | AB | AA | AB | AB |
| AB | AB | AA | AB | AA | AA | AA | AB | AA | AA | AB | AB | AB | AB | AA | AB | AA | AB | AA | AB | AB | AB | AB | AB | AB | AB | AB | AB | AB | AA | AB | AB | AA | AB | AB | AA | AB | AB | AB | AB | AB | AB | AB |
| - | AA | AA | - | AB | AA | AA | AB | AA | AA | AA | - | AA | AB | - | AB | AA | AB | - | AB | AA | AA | AB | - | AB | AA | AA | AA | AA | AA | AB | - | - | AA | AA | - | AA | AA | AA | AB | - | AB | AB |

| snp_B4990 | snp_B5037 | snp_B5057 | snp_B5110 | snp_B5136 | snp_B5147 | snp_B5162 | snp_B5169 | snp_B5222 | snp_B5253 | snp_B5275 | snp_B5277 | snp_B5285 | snp_B5293 | snp_B5315 | snp_B5319 | snp_B5349 | snp_B5371 | snp_B5412 | snp_B5419 | snp_B5422 | snp_B5454 | snp_B5501 | snp_B5527 | snp_B5529 | snp_B5542 | snp_B5580 | snp_B5582 | snp_B5587 | snp_B5606 | snp_B5617 | snp_B5619 | snp_B5628 | snp_B5642 |
|---|---|---|---|---|---|---|---|---|---|---|---|---|---|---|---|---|---|---|---|---|---|---|---|---|---|---|---|---|---|---|---|---|---|
| 10 | 10 | 10 | 10 | 10 | 10 | 10 | 10 | 10 | 10 | 10 | 10 | 10 | 10 | 10 | 10 | 10 | 10 | 10 | 10 | 10 | 10 | 10 | 10 | 10 | 10 | 10 | 10 | 10 | 10 | 10 | 10 | 10 | 10 |
| -- | -- | -- | -- | -- | -- | LG4 | -- | LG4 | -- | -- | -- | -- | -- | -- | -- | -- | -- | -- | -- | -- | -- | -- | -- | -- | -- | LG4 | -- | LG4 | -- | -- | -- | -- | -- |

| snp_B5665 | snp_B5670 | snp_B5672 | snp_B5675 | snp_B5719 | snp_B5723 | snp_B5758 | snp_B5783 | snp_B5848 | snp_B5856 | snp_B5863 | snp_B5888 | snp_B5897 | snp_B5899 | snp_B5903 | snp_B5917 | snp_B5923 |
|---|---|---|---|---|---|---|---|---|---|---|---|---|---|---|---|---|
| 10 | 10 | 10 | 10 | 10 | 10 | 10 | 10 | 10 | 10 | 10 | 10 | 10 | 10 | 10 | 10 | 10 |
| -- | -- | -- | -- | -- | -- | -- | -- | -- | -- | -- | -- | -- | -- | -- | LG4 | -- |
| AB | AA | AA | AB | AA | AA | AB | AA | AA | AA | AA | AA | AA | AA | AB | AB | AA |
| AB | AB | AA | AA | AA | AA | AA | AA | AA | AA | AA | AA | AA | AA | AA | AB | AA |
| AB | AA | AA | AB | AB | AA | AA | AA | AB | AA | AA | AB | AA | AB | AA | AB | AB |
| AA | AA | AB | - | AA | AA | AB | AB | - | AA | AA | AB | AB | - | AB | AA | AB |
| AA | AB | AB | AB | AB | AB | AB | AB | AB | AB | AA | AA | AA | AA | AA | AB | AB |
| - | AB | - | - | AA | AA | AB | AA | - | AB | - | AA | AA | - | AA | AA | AA |
| AB | AB | AB | AB | AB | AB | AB | AB | AB | AB | AB | AB | AB | AB | AB | AB | AA |
| AB | AB | AA | AA | AA | AA | AA | AA | AB | AB | AB | AA | AA | AB | AB | AA | AA |
| AB | AB | AB | AA | AA | AB | AA | AA | AA | AA | AB | AB | AB | AA | AA | AA | AB |
| AA | AA | AA | AA | AA | AA | AA | AA | AA | AA | AA | AA | AA | AA | AB | AA | AB |
| AA | AB | AA | AA | AA | AA | AA | AA | AA | AA | AB | AA | AB | AB | AB | AA | AB |
| AA | AB | AB | AB | AB | AB | AB | AB | AB | AB | AA | AA | AA | AA | AA | AB | AB |
| AB | AB | AA | AB | AB | AB | AB | AB | AB | AB | AB | AA | AA | AB | AA | AB | AA |
| AA | AB | AA | AA | AA | AA | AA | AB | AA | AB | AA | AA | AB | AA | AA | AA | AA |
| AB | AA | AB | AA | AA | AB | AB | AB | AB | AB | AA | AA | AA | AA | AB | AB | AB |
| AB | AB | AB | AA | AB | AA | AB | AB | AA | AA | AB | AB | AB | AA | AB | AA | AB |
| AA | AB | AB | AB | AA | AA | AA | AA | AB | AB | AA | AB | AA | AB | AA | AB | AB |
| - | - | - | - | - | - | AB | AA | AA | - | AB | AB | AA | - | - | AA | AA |
| AB | AB | AB | AB | AB | AB | AB | AB | AB | AB | AB | AB | AB | AB | AB | AB | AB |
| AA | AB | AB | AA | AA | AB | AA | AB | AA | AB | AB | - | AA | AA | AA | AB | AB |
| AB | AB | AB | AB | AA | AB | AB | AB | AB | AB | AB | AB | AB | AB | AB | AB | AB |
| AA | AA | AA | AA | AB | AA | AB | AB | AB | AB | AA | AA | AA | AB | AA | AB | AB |
| - | - | AA | - | - | - | AB | - | - | AB | AA | AB | AB | - | - | AB | AA |
| AB | AA | AA | AB | AB | AB | AB | AB | AB | AB | AA | AB | AB | AB | AB | AB | AA |
| AA | AB | AB | AB | AA | AB | AA | AA | AA | AA | AB | AA | AA | AA | AA | AA | AB |
| AA | AB | AB | AA | AA | AA | AA | AA | AA | AA | AB | AA | AB | AA | AA | AA | AA |
| AA | AB | AB | AB | AB | AB | AB | AB | AB | AB | AB | AB | AB | AB | AA | AB | AB |
| AA | AA | AA | AA | AA | AB | AB | AB | AA | AB | AA | AA | AA | AA | AA | AB | AA |
| AB | - | AA | - | AB | AB | - | AB | AB | AB | AB | AA | AA | AA | AA | AB | AB |
| - | AB | - | - | - | - | - | AA | AA | - | - | AA | AA | - | AA | AA | AB |
| AB | AA | AA | AB | AB | AA | AB | AA | AA | AB | AA | AB | AB | AB | AB | AB | AA |
| AA | AB | AB | AB | AB | AB | AB | AA | AA | AB | AB | AA | AA | AA | AA | AB | AA |
| - | AB | AB | - | AA | AB | - | AA | AA | - | AB | AB | AB | - | AB | AA | AA |
| AA | AB | AB | - | AA | AA | AA | AB | AB | - | AB | - | AA | AB | AA | AA | AA |
| - | - | - | - | - | - | AA | AA | - | AB | - | AA | AA | - | - | AA | AA |
| AB | AA | AB | AB | AA | AA | AB | AB | AA | AB | AB | AB | AB | - | AB | AB | AB |
| AB | AB | AB | AB | AB | AB | AB | AA | AB | AB | AB | AB | AB | AB | AB | AB | AA |
| AB | AA | AA | AB | AB | AA | AB | AA | AB | AA | AA | AB | AB | AB | AB | AB | AA |
| AA | AA | AA | AA | AB | AA | AA | AA | AA | AA | AA | AA | AA | AA | AB | AB | AA |
| - | AB | AB | AA | AA | AB | AA | AA | - | AB | AA | AA | AA | AA | AA | AA | AB |
| AA | AB | AB | AA | AA | AB | AA | AA | AB | AB | AB | AA | AB | AB | AA | AB | AA |
| AA | AB | AB | AB | AA | AB | AB | AB | AA | AB | AB | AA | AA | AA | AA | AB | AB |
| AA | AA | AA | AB | AA | AA | - | AA | AA | AA | AB | AA | AA | AA | AA | AA | AA |
| AA | AB | AA | AA | AA | AB | - | AA | AB | AA | AB | AA | AB | AB | AA | AA | AA |
| AB | AB | AA | AB | AB | AA | AB | AB | AB | AB | AB | AB | AB | AA | AA | AB | AA |
| AA | AA | AA | AB | AA | AB | AB | AA | AB | AB | AA | AA | AB | AA | AB | AB | AB |
| AB | AB | AB | AB | AB | AB | AB | AB | AB | AB | AB | AB | AB | AB | AB | AB | AB |
| AB | AA | AA | AB | AB | AB | AB | AB | AB | AB | AB | AB | AB | AB | AB | AB | AA |
| AB | AB | AA | AB | AB | AB | AB | AB | AA | AB | AB | AB | AB | AA | AB | AB | AA |
| AB | AB | AB | AA | AA | AB | AA | AB | AB | AB | AB | AB | AB | AB | AB | AB | AA |
| AB | AB | AA | AA | AA | AA | AA | AA | AB | AB | AB | AA | AA | AB | AA | AA | AA |

| snp_B1298 | snp_B1361 | snp_B1372 | snp_B1379 | snp_B1381 | snp_B1450 | snp_B1534 | snp_B1566 | snp_B1588 | snp_B1617 | snp_B1662 | snp_B1697 | snp_B1703 | snp_B1715 | snp_B1730 | snp_B1748 | snp_B1764 | snp_B1796 | snp_B1802 | snp_B1813 | snp_B1858 | snp_B1888 | snp_B1915 | snp_B1925 | snp_B1940 |
|---|---|---|---|---|---|---|---|---|---|---|---|---|---|---|---|---|---|---|---|---|---|---|---|---|
| 11 | 11 | 11 | 11 | 11 | 11 | 11 | 11 | 11 | 11 | 11 | 11 | 11 | 11 | 11 | 11 | 11 | 11 | 11 | 11 | 11 | 11 | 11 | 11 | 11 |
| -- | -- | -- | -- | -- | -- | -- | -- | LG11 | -- | -- | -- | -- | LG11 | LG11 | -- | LG12 | -- | -- | -- | LG11 | -- | -- | -- | -- |
| [illegible] | [illegible] | [illegible] | [illegible] | [illegible] | [illegible] | [illegible] | [illegible] | [illegible] | [illegible] | [illegible] | [illegible] | [illegible] | [illegible] | [illegible] | [illegible] | [illegible] | [illegible] | [illegible] | [illegible] | [illegible] | [illegible] | [illegible] | [illegible] | [illegible] |

| snp_B1947 | snp_B1972 | snp_B2030 | snp_B2161 | snp_B2179 | snp_B2194 | snp_B2223 | snp_B2264 | snp_B2291 | snp_B2319 | snp_B2320 | snp_B2324 | snp_B2343 | snp_B2374 | snp_B2376 | snp_B2420 | snp_B2421 | snp_B2432 | snp_B2462 | snp_B2468 | snp_B2473 | snp_B2507 | snp_B2551 | snp_B2616 | snp_B2646 |
|---|---|---|---|---|---|---|---|---|---|---|---|---|---|---|---|---|---|---|---|---|---|---|---|---|
| 11 | 11 | 11 | 11 | 11 | 11 | 11 | 11 | 11 | 11 | 11 | 11 | 11 | 11 | 11 | 11 | 11 | 11 | 11 | 11 | 11 | 11 | 11 | 11 | 11 |
| LG11 | -- | -- | LG11 | -- | -- | -- | LG12 | -- | -- | -- | -- | -- | -- | -- | -- | -- | -- | LG12 | -- | -- | -- | -- | -- | -- |
| [illegible] | [illegible] | [illegible] | [illegible] | [illegible] | [illegible] | [illegible] | [illegible] | [illegible] | [illegible] | [illegible] | [illegible] | [illegible] | [illegible] | [illegible] | [illegible] | [illegible] | [illegible] | [illegible] | [illegible] | [illegible] | [illegible] | [illegible] | [illegible] | [illegible] |

| snp_B2647 | snp_B2651 | snp_B2656 | snp_B2666 | snp_B2711 | snp_B2722 | snp_B2738 | snp_B2745 | snp_B2752 | snp_B2762 | snp_B2771 | snp_B2787 | snp_B2805 | snp_B2852 | snp_B2873 | snp_B2941 | snp_B2949 | snp_B2970 | snp_B2980 | snp_B2988 | snp_B3007 | snp_B3013 | snp_B3051 | snp_B3059 | snp_B3077 | snp_B3121 | snp_B3134 |
|---|---|---|---|---|---|---|---|---|---|---|---|---|---|---|---|---|---|---|---|---|---|---|---|---|---|---|
| 11 | 11 | 11 | 11 | 11 | 11 | 11 | 11 | 11 | 11 | 11 | 11 | 11 | 11 | 11 | 11 | 11 | 11 | 11 | 11 | 11 | 11 | 11 | 11 | 11 | 11 | 11 |
| -- | -- | -- | LG11 | -- | -- | -- | -- | -- | -- | -- | -- | -- | -- | -- | -- | -- | -- | -- | -- | -- | LG12 | -- | -- | -- | -- | -- |
| [illegible] | [illegible] | [illegible] | [illegible] | [illegible] | [illegible] | [illegible] | [illegible] | [illegible] | [illegible] | [illegible] | [illegible] | [illegible] | [illegible] | [illegible] | [illegible] | [illegible] | [illegible] | [illegible] | [illegible] | [illegible] | [illegible] | [illegible] | [illegible] | [illegible] | [illegible] | [illegible] |

| snp_B3160 | snp_B3184 | snp_B3195 | snp_B3237 | snp_B3256 | snp_B3277 | snp_B3296 | snp_B3334 | snp_B3366 | snp_B3386 | snp_B3415 | snp_B3448 | snp_B3483 | snp_B3497 | snp_B3509 | snp_B3513 | snp_B3602 | snp_B3604 | snp_B3605 | snp_B3606 | snp_B3624 | snp_B3644 | snp_B3646 | snp_B3675 | snp_B3712 | snp_B3718 | snp_B3720 | snp_B3728 | snp_B3733 | snp_B3839 | snp_B3901 | snp_B3923 | snp_B3960 | snp_B4004 | snp_B4012 | snp_B4049 | snp_B4057 | snp_B4065 | snp_B4124 | snp_B4137 | snp_B4200 | snp_B4220 | snp_B4252 | snp_B4260 | snp_B4274 | snp_B4291 | snp_B4317 | snp_B4320 | snp_B4355 | snp_B4359 | snp_B4394 | snp_B4423 | snp_B4441 | snp_B4461 | snp_B4542 | snp_B4552 | snp_B4641 | snp_B4684 | snp_B4687 | snp_B4709 | snp_B4721 | snp_B4740 | snp_B4767 | snp_B4784 | snp_B4804 | snp_B4922 | snp_B4936 | snp_B4945 | snp_B4950 | snp_B4963 | snp_B5034 | snp_B5070 | snp_B5074 | snp_B5094 | snp_B5105 | snp_B5113 | snp_B5128 |
|---|---|---|---|---|---|---|---|---|---|---|---|---|---|---|---|---|---|---|---|---|---|---|---|---|---|---|---|---|---|---|---|---|---|---|---|---|---|---|---|---|---|---|---|---|---|---|---|---|---|---|---|---|---|---|---|---|---|---|---|---|---|---|---|---|---|---|---|---|---|---|---|---|---|---|---|---|
| 11 | 11 | 11 | 11 | 11 | 11 | 11 | 11 | 11 | 11 | 11 | 11 | 11 | 11 | 11 | 11 | 11 | 11 | 11 | 11 | 11 | 11 | 11 | 11 | 11 | 11 | 11 | 11 | 11 | 11 | 11 | 11 | 11 | 11 | 11 | 11 | 11 | 11 | 11 | 11 | 11 | 11 | 11 | 11 | 11 | 11 | 11 | 11 | 11 | 11 | 11 | 11 | 11 | 11 | 11 | 11 | 11 | 11 | 11 | 11 | 11 | 11 | 11 | 11 | 11 | 11 | 11 | 11 | 11 | 11 | 11 | 11 | 11 | 11 | 11 | 11 | 11 |
| -- | -- | -- | -- | -- | LG12 | LG11 | -- | -- | -- | LG11 | -- | -- | -- | -- | -- | -- | -- | -- | -- | -- | -- | -- | -- | -- | LG12 | -- | -- | -- | -- | -- | -- | -- | -- | -- | -- | -- | -- | -- | -- | -- | -- | -- | -- | -- | -- | -- | LG11 | LG12 | -- | -- | LG12 | -- | -- | -- | -- | -- | -- | -- | -- | -- | LG11 | -- | LG12 | -- | -- | -- | -- | -- | -- | LG11 | -- | -- | LG12 | -- | -- | -- |

| snp_B5149 | snp_B5158 | snp_B5203 | snp_B5241 | snp_B5307 | snp_B5311 | snp_B5365 | snp_B5376 | snp_B5378 | snp_B5383 | snp_B5387 | snp_B5429 | snp_B5447 | snp_B5516 | snp_B5532 | snp_B5566 | snp_B5621 | snp_B5622 | snp_B5654 | snp_B5674 | snp_B5694 | snp_B5698 | snp_B5730 | snp_B5739 |
|---|---|---|---|---|---|---|---|---|---|---|---|---|---|---|---|---|---|---|---|---|---|---|---|
| 11 | 11 | 11 | 11 | 11 | 11 | 11 | 11 | 11 | 11 | 11 | 11 | 11 | 11 | 11 | 11 | 11 | 11 | 11 | 11 | 11 | 11 | 11 | 11 |
| -- | -- | -- | -- | -- | LG11 | LG11 | -- | -- | -- | -- | -- | LG12 | -- | -- | LG11 | -- | -- | -- | -- | -- | -- | -- | -- |
| AA | AB | AB | AA | AB | AA | AB | AB | AB | AB | - | AA | AA | AA | AA | AA | AA | AA | AB | AA | AB | AB | AB | AA |
| AB | AA | AB | AB | AA | AA | AB | AB | AB | AA | AB | AB | AB | AA | AA | AA | AA | AA | AB | AA | AB | AB | AB | AA |
| AB | AA | AB | AB | AB | AA | AB | AB | AA | AA | AB | AA | AB | AB | AA | AA | AA | AA | AA | AA | AB | AB | AB | AB |
| AB | AB | - | AB | AB | AA | AB | - | AB | AB | AB | AB | AA | - | AB | AB | AB | - | - | AA | AB | AB | AB | - |
| AA | AB | AB | AB | AB | AA | AB | AB | AB | AB | AB | AA | AB | AB | AB | AB | AB | AB | AA | AB | AB | AB | AB | AB |
| AB | - | - | - | AA | AA | AB | AA | AA | AB | AA | AB | AA | AB | AB | - | - | - | AA | - | - | AB | AB | AB |
| AB | AB | AB | AA | AB | AA | AA | AA | AB | AA | AB | AA | AB | AA | AB | AA | AA | AA | AA | AB | AA | AA | AA | AA |
| AA | AB | AB | AA | AB | AB | AA | AA | AB | AA | AB | AA | AB | AA | AA | AB | AB | AB | AA | AB | AA | AA | AA | AA |
| AB | AA | AB | AB | AB | AB | AA | AA | AA | AA | AB | AB | AB | AA | AA | AB | AA | AB | AA | AB | AA | AA | AA | AB |
| AB | AA | AA | AB | AA | AB | AA | AB | AA | AB | AA | AB | AA | AA | AB | AB | AA | AB | AA | AB | AA | AA | AB | AB |
| AB | AA | AB | AA | AA | AB | AA | AA | AB | AA | AB | AB | AA | AB | AA | AB | AA | AB | AA | AB | AA | AB | AB | AB |
| AA | AB | AA | AB | AB | AB | AA | AB | AA | AB | AA | AA | AB | AB | AB | AB | AB | AA | AB | AA | AB | AB | AB | AB |
| AB | AB | AB | AA | AB | AB | AB | AB | AA | AA | AB | AB | AB | AA | AA | AB | AB | AB | AB | AB | AA | AA | AB | AB |
| AB | AA | AB | AA | AB | AB | AA | AB | AB | AA | AB | AB | AB | AB | AA | AB | AA | AB | AA | AB | AB | AA | AB | AB |
| AB | AA | AB | AA | AB | AA | AA | AB | AB | AA | AB | AB | AA | AB | AA | AB | AA | AB | AB | AB | AB | AB | AB | AB |
| AA | AA | AA | AA | AA | AB | AA | AB | AB | AB | AA | AA | AA | AB | AB | AA | AA | AA | AA | AB | AB | AA | AB | AA |
| AB | AB | AA | AA | AB | AB | AB | AA | AA | AB | AB | AB | AA | AB | AB | AB | AB | AB | AB | AB | AB | AB | AB | AB |
| AB | - | - | AA | AA | AA | AA | - | AB | - | AA | AA | - | AB | AA | - | - | AA | - | AB | - | - | AA | - |
| AA | AB | AB | AB | AB | AA | AA | AB | AB | AA | AB | AA | AB | AA | AA | AA | AA | AA | AA | AB | AA | AB | AB | AA |
| AA | AB | AB | AB | AB | AB | AA | AA | AB | AA | AB | AB | AB | AA | AA | AB | AB | AB | AB | AB | AB | AA | AB | AB |
| AB | AA | AA | AB | AA | AB | AB | AB | AA | AB | AA | AB | AA | AB | AB | AB | AB | AB | AA | AB | AB | AB | AB | AA |
| AA | AB | AB | AB | AB | AA | AA | AA | AA | AA | AB | AA | AB | AA | AA | AA | AA | AA | AA | AB | AA | AA | AA | AA |
| AA | - | - | - | - | - | AB | - | AA | AA | AA | AB | AA | - | AA | AA | - | - | - | - | - | AA | AB | - |
| AA | AB | AB | AA | AB | AA | AA | AA | AB | AA | AB | AB | AB | AB | AB | AA | AA | AA | AA | AB | AB | AA | AA | AA |
| AA | AA | AA | AB | AA | AA | AA | AA | AB | AA | AB | AB | AA | AA | AB | AA | AA | AA | AB | AB | AA | AA | AA | AA |
| AB | AA | AA | AA | AB | AA | AB | AB | AB | AB | AA | AB | AA | AA | AB | AA | AB | AA | AA | AB | AB | AB | AB | AA |
| AB | AB | AB | AB | AB | AA | AB | AB | AB | AB | AB | AB | AB | AB | AB | AB | AB | AB | AB | AB | AB | AB | AB | AA |
| AB | AB | AB | AB | AB | AB | AB | AB | AB | AB | AB | AB | AB | AB | AB | AB | AB | AB | AB | AB | AB | AB | AB | AB |
| AB | AA | AA | AB | AA | AB | AA | AA | AB | AB | - | AA | AA | - | AB | AB | AA | AA | AB | AA | AB | AB | AB | AB |
| AA | AA | - | - | AA | AB | AA | - | AB | AA | - | AA | - | - | AA | - | - | AA | - | - | AA | AA | AA | AA |
| AB | AB | AB | AB | AB | AB | AB | AB | AB | AB | AB | AB | AB | AB | AB | AB | AB | AB | AB | AB | AB | AB | AB | AB |

| snp_B5743 | snp_B5750 | snp_B5761 | snp_B5771 | snp_B5780 | snp_B5791 | snp_B5800 | snp_B5801 | snp_B5817 | snp_B5829 | snp_B5851 | snp_B5879 | snp_B5895 | snp_B5910 | snp_B5913 | snp_B5955 | snp_B5958 | snp_B5990 | snp_B6000 | snp_B6016 | snp_B6149 | snp_B6167 | snp_B6244 | snp_B6170 | snp_B6018 | snp_B6005 |
|---|---|---|---|---|---|---|---|---|---|---|---|---|---|---|---|---|---|---|---|---|---|---|---|---|---|
| 11 | 11 | 11 | 11 | 11 | 11 | 11 | 11 | 11 | 11 | 11 | 11 | 11 | 11 | 11 | 11 | 11 | 11 | 11 | 11 | 11 | 11 | 11 | 12 | 12 | 12 |
| -- | -- | -- | -- | -- | -- | -- | LG12 | -- | LG12 | -- | -- | -- | -- | -- | -- | -- | -- | -- | -- | -- | -- | LG11 | -- | -- | -- |
| AA | AA | AA | AA | AA | AA | AA | AA | AA | AA | AA | AA | AB | AB | AA | AA | AA | AA | AA | AA | AB | AA | AA | AA | AA | AA |
| AA | AB | AB | AB | AA | AA | AA | AB | AB | AB | AA | AA | AA | AA | AB | AA | AA | AA | AA | AA | AB | AA | AA | AB | AB | AB |
| AB | AA | AB | AA | AB | AA | AB | AB | AB | AB | AA | AA | AA | AB | AB | AA | AA | AA | AA | AA | AB | AA | AA | AA | AA | AA |
| AB | AB | AB | - | AA | AB | AA | - | AA | AA | AB | AB | - | - | AB | AB | AB | AB | AB | AB | AB | AB | AB | AB | AB | AB |
| AB | AB | AB | AA | AB | AB | AA | AB | AB | AB | AB | AB | AB | AA | AB | AB | AA | AA | AA | AB | AB | AB | AB | AB | AB | AB |
| AB | - | AA | - | AB | AB | AA | - | - | - | AB | AB | AB | - | AB | - | AB | AB | - | AB | AA | AB | - | AA | - | AB |
| AB | AA | AA | AA | AB | AA | AA | AA | - | AA | AB | AB | AA | AA | AB | AA | AB | AA | AB | AA | AB | AA | AA | AB | AA | AB |
| AA | AB | AB | AA | AA | AB | AB | AB | AB | AB | AA | AB | AA | AB | AA | AB | AA | AB | AB | AA | AB | AB | AB | AA | AA | AA |
| AA | AB | AA | AB | AA | AB | AB | AB | AB | AB | AA | AB | AA | AA | AB | AA | AA | AB | AB | AA | AB | AB | AB | AB | AB | AB |
| AA | AA | AB | AA | AA | AB | AB | AB | AA | AB | AA | AA | AB | AA | AB | AA | AB | AB | AB | AA | AB | AA | AB | AA | AA | AA |
| AA | AB | AB | AA | AB | AB | AB | AB | AA | AA | AB | AB | AB | AA | AB | AB | AB | AB | AB | AA | AB | AA | AB | AA | AA | AA |
| AB | AA | AA | AA | AA | AB | AB | AB | AB | AB | AA | AB | AB | AA | AB | AA | AB | AB | AA | AB | AB | AA | AB | AB | AA | AB |
| AB | AB | AB | AB | AA | AB | AB | AB | AB | AB | AB | AB | AB | AA | AB | AB | AB | AB | AB | AB | AB | AB | AB | AB | AB | AB |
| AB | AB | AB | AB | AA | AB | AB | AB | AB | AB | AB | AB | AB | AA | AB | AB | AB | AB | AB | AA | AB | AB | AB | AB | AA | AB |
| AB | AB | AB | AB | AA | AB | AA | AA | AB | AB | AB | AA | AB | AB | AA | AB | AB | AB | AB | AA | AB | AB | AB | AB | AA | AB |
| AA | AB | AB | AA | AB | AA | AB | AA | AB | AB | AB | AB | AB | AA | AB | AB | AA | AA | AB | AB | AB | AB | AB | AB | AA | AB |
| AB | AA | AA | AB | AB | AB | AB | AA | AA | AA | AB | AB | AB | AB | AB | AB | AB | AB | AB | AB | AB | AB | AB | AB | AB | AB |
| AA | - | AB | AB | AB | AA | AA | - | - | - | AA | AB | - | - | AA | AB | - | AA | - | AA | AA | AB | - | AA | - | AA |
| AA | AB | AB | AA | AB | AB | AA | AB | AB | AB | AA | AA | AB | AA | AA | AA | AA | AA | AA | AA | AB | AA | AA | AB | AB | AB |
| AA | AB | AB | AA | AB | AB | AB | AB | AB | AB | AA | AA | AB | AA | AB | AB | AA | AB | AB | AB | AB | AB | AA | AB | AB | AB |
| AB | AA | AB | AB | AB | AB | AB | AA | AA | AA | AB | AA | AB | AB | AA | AB | AB | AB | AB | AA | AB | AB | AB | AB | AB | AB |
| AA | AA | AA | AA | AA | AA | AA | AB | AB | AB | AA | AA | AA | AA | AA | AA | AA | AA | AA | AA | AA | AA | AA | AA | AA | AA |
| AB | AB | AB | AA | - | AB | AA | - | - | AA | AB | - | - | AB | AB | - | AA | AA | AA | AA | AA | AA | - | AB | - | AB |
| AA | AB | AB | AB | AA | AA | AB | AB | AB | AB | AA | AA | AB | AA | AB | AB | AA | AA | AA | AB | AB | AA | AB | AA | AB | AB |
| AA | AA | AA | AB | AA | AA | AA | AA | AA | AA | AA | AA | AA | AB | AA | AA | AB | AB | AA | AB | AA | AA | AA | AB | AA | AA |
| AA | AA | AA | AA | AA | AA | AA | AB | AB | AB | AA | AA | AA | AB | AA | AB | AA | AA | AB | AA | AB | AA | AA | AA | AB | AA |
| AB | AB | AB | AB | AB | AB | AB | AB | AB | AB | AB | AB | AA | AB | AA | AB | AB | AB | AB | AB | AA | AB | AB | AA | AA | AB |
| AB | AB | AB | AB | AB | AB | AB | AB | AB | AB | AB | AB | AB | AB | AB | AB | AB | AB | AB | AB | AB | AB | AB | AA | AA | AB |
| AB | AB | AB | AB | AA | AB | AA | AA | AA | AB | AA | AB | - | AA | AB | AB | AB | AB | AB | AA | AB | AB | AB | AA | AA | AB |
| AA | - | AB | AA | AA | AA | - | AA | - | - | AA | AB | - | - | AA | AA | - | AA | - | AA | AB | AB | - | AA | AB | AB |
| AB | AB | AB | AB | AB | AB | AB | AB | AB | AB | AB | AB | AB | AB | AB | AB | AB | AB | AB | AB | AB | AB | AB | AB | AB | AB |

| snp_B5982 | snp_B5980 | snp_B5978 | snp_B5973 | snp_B5959 | snp_B5919 | snp_B5894 | snp_B5889 | snp_B5832 | snp_B5777 | snp_B5732 | snp_B5721 | snp_B5711 | snp_B5703 | snp_B5700 | snp_B5699 | snp_B5633 | snp_B5611 | snp_B5534 | snp_B5519 | snp_B5513 | snp_B5496 | snp_B5494 | snp_B5489 | snp_B5463 | snp_B5457 | snp_B5453 |
|---|---|---|---|---|---|---|---|---|---|---|---|---|---|---|---|---|---|---|---|---|---|---|---|---|---|---|
| 12 | 12 | 12 | 12 | 12 | 12 | 12 | 12 | 12 | 12 | 12 | 12 | 12 | 12 | 12 | 12 | 12 | 12 | 12 | 12 | 12 | 12 | 12 | 12 | 12 | 12 | 12 |
| -- | -- | LG7 | -- | -- | LG7 | -- | -- | -- | -- | -- | -- | -- | -- | -- | -- | -- | -- | -- | -- | -- | -- | LG7 | -- | -- | -- | LG7 |
| AA | AB | AA | AB | AA | AA | AA | AA | AA | AB | AA | AA | AA | AA | AA | AA | AA | AB | AB | AA | AA | AA | AA | AA | AA | AB | AA |
| AA | AB | AB | AA | AB | AB | AA | AA | AB | AB | AA | AB | AA | AA | AA | AB | AB | AA | AA | AB | AA | AA | AB | AA | AA | AA | AB |
| AA | AB | AA | AA | AA | AA | AA | AB | AA | AA | AA | AB | AA | AA | AA | AB | AA | AA | AB | AA | AA | AA | AA | AA | AA | AB | AA |
| AA | AB | AB | AA | AB | AB | AB | AA | AB | AB | AA | - | AB | AA | AA | AA | AA | AA | AA | AB | AA | - | AB | AA | AB | AB | - |
| AB | AB | AB | AB | AB | AB | AB | AB | AB | AB | AB | AA | AB | AB | AB | AB | AA | AB | AB | AB | AB | AB | AB | AB | AB | AB | AB |
| AA | AB | - | AB | AB | - | AB | AA | AB | AB | AB | - | AB | AB | AB | AB | AB | AB | AB | AB | AB | - | - | AA | AB | - | - |
| AB | AB | AB | AB | AA | AB | AB | AB | AB | AB | AA | AB | AB | AB | AB | AB | AA | AA | AB | AA | AA | AB | AB | AA | AA | AA | AB |
| AB | AA | AA | AB | AB | AA | AB | AB | AA | AB | AA | AA | AB | AA | AA | AB | AA | AB | AB | AA | AA | AA | AA | AA | AB | AB | AA |
| AB | AA | AB | AB | AB | AA | AB | AB | AA | AB | AB | AA | AB | AA | AA | AB | AA | AB | AA | AB | AB | AA | AB | AB | AB | AB | AB |
| AA | AA | AA | AA | AA | AA | AA | AB | AA | AA | AA | AA | AA | AA | AB | AA | AB | AB | AA | AA | AA | AB | AA | AA | AA | AB | AA |
| AA | AA | AA | AA | AA | AA | AA | AA | AA | AA | AB | AA | AA | AA | AA | AA | AB | AB | AB | AA | AA | AB | AA | AA | AA | AB | AA |
| AB | AA | AB | AB | AB | AA | AB | AA | AB | AB | AB | AB | AB | AA | AB | AB | AB | AA | AB | AB | AB | AB | AB | AB | AB | AA | AB |
| AB | AA | AB | AB | AA | AB | AB | AA | AA | AB | AB | AB | AA | AB | AB | AA | AB | AA | AA | AB | AA | AA | AB | AA | AB | AA | AB |
| AA | AA | AA | AA | AA | AA | AA | AA | AA | AA | AB | AA | AA | AA | AA | AA | AB | AB | AA | AA | AA | AA | AA | AA | AA | AB | AA |
| AB | AB | AB | AA | AB | AB | AB | AB | AB | AB | AB | AB | AB | AA | AB | AB | AB | AA | AA | AA | AB | AB | AB | AB | AB | AB | AB |
| AB | AB | AB | AB | AB | AB | AB | AB | AB | AB | AA | AB | AB | AB | AB | AB | AA | AB | AB | AA | AA | AB | AB | AB | AB | AB | AB |
| AA | AA | AA | AA | - | AA | AA | AB | - | AA | AA | AA | AA | AA | AA | AA | AA | AA | AA | AA | AA | - | - | AA | AA | AB | AA |
| AB | AB | AB | AB | AB | AB | AB | AB | AB | AB | AB | AB | AB | AB | AB | AB | AA | AA | AA | AA | AB | AB | AB | AB | AB | AB | AB |
| AA | AB | AA | AA | AA | AB | AB | AB | AB | AA | AB | AB | AA | AA | AA | AA | AB | AA | AA | AB | AA | - | AA | AA | AA | AA | AB |
| AB | AB | AB | AB | AA | AB | AB | AB | AB | AB | AB | AB | AB | AA | AB | AB | AB | AA | AB | AB | AB | AB | AB | AA | AB | AB | AB |
| AB | AA | AA | AA | AA | AA | AA | AB | AA | AA | AA | AA | AB | AA | AB | AB | AB | AB | AA | AB | AB | AA | AA | AA | AA | AB | AA |
| AB | AB | - | AB | - | - | AB | AA | AB | AA | AB | - | AB | AB | - | AB | AB | AA | AB | AA | - | - | - | - | - | - | - |
| AB | AB | AB | AA | AB | AB | AA | AA | AB | AB | AA | AB | AA | AB | AA | AA | AA | AA | AB | AB | AA | AB | AB | AA | AB | AA | AB |
| AB | AB | AA | AB | AB | AB | AA | AA | AB | AB | AA | AB | AB | AA | AB | AB | AB | AA | AB | AB | AB | AB | AB | AB | AB | AB | AB |
| AB | AB | AA | AB | AB | AB | AB | AB | AB | AB | AA | AB | AB | AA | AB | AB | AB | AA | AB | AB | AB | AB | AB | AB | AB | AB | AB |
| AB | AA | AA | AB | AA | AA | AA | AB | AA | AA | AA | AA | AB | AB | AB | AB | AB | AA | AB | AB | AB | AA | AA | AA | AB | AB | AA |
| AA | AA | AA | AB | AB | AA | AA | AB | AA | AA | AB | AA | AB | AA | AA | AB | AB | AA | AB | AA | AB | AA | AA | AA | AB | AB | AA |
| AB | AB | - | AB | AB | AB | AB | AB | AB | AB | AB | AB | AB | AA | - | AB | AA | AA | AB | AA | AB | AA | AB | AB | - | AA | AB |
| AB | AA | - | AB | AB | AB | AA | AA | AB | AB | AA | AB | AB | AB | - | AB | AA | AA | AA | AB | AA | AB | - | AB | - | - | AB |
| AB | AA | AB | AB | AA | AB | AA | AA | AB | AB | AB | AB | AB | AA | AB | AB | AB | AB | AB | AA | AA | AA | AB | AB | AA | AB | AA |

| snp_B5436 | snp_B5417 | snp_B5411 | snp_B5399 | snp_B5394 | snp_B5392 | snp_B5374 | snp_B5334 | snp_B5305 | snp_B5274 | snp_B5273 | snp_B5264 | snp_B5250 | snp_B5217 | snp_B5192 | snp_B5164 | snp_B5151 | snp_B5106 | snp_B5084 | snp_B5024 | snp_B5021 | snp_B4983 | snp_B4965 | snp_B4937 | snp_B4907 | snp_B4901 | snp_B4860 | snp_B4850 | snp_B4797 | snp_B4759 | snp_B4725 | snp_B4645 | snp_B4637 | snp_B4635 | snp_B4593 | snp_B4586 | snp_B4572 | snp_B4460 | snp_B4398 | snp_B4390 |
|---|---|---|---|---|---|---|---|---|---|---|---|---|---|---|---|---|---|---|---|---|---|---|---|---|---|---|---|---|---|---|---|---|---|---|---|---|---|---|---|
| 12 | 12 | 12 | 12 | 12 | 12 | 12 | 12 | 12 | 12 | 12 | 12 | 12 | 12 | 12 | 12 | 12 | 12 | 12 | 12 | 12 | 12 | 12 | 12 | 12 | 12 | 12 | 12 | 12 | 12 | 12 | 12 | 12 | 12 | 12 | 12 | 12 | 12 | 12 | 12 |
| -- | -- | -- | -- | -- | -- | -- | -- | -- | -- | LG7 | -- | -- | -- | -- | -- | -- | -- | -- | -- | -- | -- | -- | -- | -- | -- | -- | LG7 | -- | -- | -- | LG7 | -- | -- | -- | -- | -- | -- | -- | LG7 |
| [illegible] | [illegible] | [illegible] | [illegible] | [illegible] | [illegible] | [illegible] | [illegible] | [illegible] | [illegible] | [illegible] | [illegible] | [illegible] | [illegible] | [illegible] | [illegible] | [illegible] | [illegible] | [illegible] | [illegible] | [illegible] | [illegible] | [illegible] | [illegible] | [illegible] | [illegible] | [illegible] | [illegible] | [illegible] | [illegible] | [illegible] | [illegible] | [illegible] | [illegible] | [illegible] | [illegible] | [illegible] | [illegible] | [illegible] | [illegible] |

| snp_B4388 | snp_B4352 | snp_B4339 | snp_B4328 | snp_B4309 | snp_B4280 | snp_B4237 | snp_B4225 | snp_B4212 | snp_B4196 | snp_B4188 | snp_B4089 | snp_B4023 | snp_B3990 | snp_B3987 | snp_B3957 | snp_B3956 | snp_B3950 | snp_B3921 | snp_B3914 | snp_B3906 | snp_B3899 | snp_B3894 | snp_B3879 | snp_B3828 | snp_B3826 | snp_B3794 | snp_B3772 | snp_B3742 | snp_B3735 | snp_B3722 | snp_B3695 | snp_B3690 | snp_B3679 | snp_B3672 | snp_B3620 | snp_B3579 |
|---|---|---|---|---|---|---|---|---|---|---|---|---|---|---|---|---|---|---|---|---|---|---|---|---|---|---|---|---|---|---|---|---|---|---|---|---|
| 12 | 12 | 12 | 12 | 12 | 12 | 12 | 12 | 12 | 12 | 12 | 12 | 12 | 12 | 12 | 12 | 12 | 12 | 12 | 12 | 12 | 12 | 12 | 12 | 12 | 12 | 12 | 12 | 12 | 12 | 12 | 12 | 12 | 12 | 12 | 12 | 12 |
| -- | -- | -- | -- | -- | -- | -- | -- | -- | -- | LG7 | LG7 | -- | -- | -- | -- | -- | -- | -- | LG7 | -- | -- | -- | -- | -- | -- | -- | -- | -- | -- | LG7 | -- | -- | -- | -- | -- | -- |
| [illegible] | [illegible] | [illegible] | [illegible] | [illegible] | [illegible] | [illegible] | [illegible] | [illegible] | [illegible] | [illegible] | [illegible] | [illegible] | [illegible] | [illegible] | [illegible] | [illegible] | [illegible] | [illegible] | [illegible] | [illegible] | [illegible] | [illegible] | [illegible] | [illegible] | [illegible] | [illegible] | [illegible] | [illegible] | [illegible] | [illegible] | [illegible] | [illegible] | [illegible] | [illegible] | [illegible] | [illegible] |

| snp_B3577 | snp_B3553 | snp_B3540 | snp_B3475 | snp_B3392 | snp_B3385 | snp_B3352 | snp_B3295 | snp_B3291 | snp_B3278 | snp_B3255 | snp_B3245 | snp_B3228 | snp_B3130 | snp_B3120 | snp_B3117 | snp_B3106 | snp_B3069 | snp_B3038 | snp_B3021 | snp_B2946 | snp_B2945 | snp_B2944 | snp_B2892 | snp_B2859 | snp_B2856 | snp_B2821 | snp_B2799 | snp_B2795 | snp_B2783 | snp_B2779 | snp_B2774 | snp_B2750 | snp_B2628 | snp_B2581 | snp_B2568 | snp_B2481 | snp_B2451 | snp_B2448 | snp_B2439 | snp_B2378 | snp_B2360 | snp_B2344 | snp_B2309 | snp_B2268 | snp_B2198 | snp_B2191 | snp_B2170 | snp_B2157 | snp_B2129 | snp_B2127 | snp_B2126 | snp_B2122 | snp_B2088 | snp_B2064 | snp_B2051 | snp_B2012 | snp_B1978 | snp_B1976 | snp_B1876 | snp_B1851 | snp_B1825 | snp_B1805 | snp_B1792 | snp_B1783 | snp_B1757 | snp_B1734 | snp_B1708 | snp_B1663 | snp_B1661 | snp_B1649 | snp_B1644 | snp_B1631 | snp_B1585 | snp_B1575 | snp_B1565 | snp_B1389 |
|---|---|---|---|---|---|---|---|---|---|---|---|---|---|---|---|---|---|---|---|---|---|---|---|---|---|---|---|---|---|---|---|---|---|---|---|---|---|---|---|---|---|---|---|---|---|---|---|---|---|---|---|---|---|---|---|---|---|---|---|---|---|---|---|---|---|---|---|---|---|---|---|---|---|---|---|---|
| 12 | 12 | 12 | 12 | 12 | 12 | 12 | 12 | 12 | 12 | 12 | 12 | 12 | 12 | 12 | 12 | 12 | 12 | 12 | 12 | 12 | 12 | 12 | 12 | 12 | 12 | 12 | 12 | 12 | 12 | 12 | 12 | 12 | 12 | 12 | 12 | 12 | 12 | 12 | 12 | 12 | 12 | 12 | 12 | 12 | 12 | 12 | 12 | 12 | 12 | 12 | 12 | 12 | 12 | 12 | 12 | 12 | 12 | 12 | 12 | 12 | 12 | 12 | 12 | 12 | 12 | 12 | 12 | 12 | 12 | 12 | 12 | 12 | 12 | 12 | 12 | 12 |
| -- | LG7 | -- | LG7 | -- | -- | LG7 | -- | -- | -- | LG7 | -- | -- | -- | -- | -- | -- | LG7 | -- | -- | -- | -- | -- | -- | -- | LG7 | -- | -- | -- | -- | -- | -- | -- | -- | -- | -- | -- | -- | -- | -- | -- | -- | -- | -- | -- | -- | -- | -- | -- | -- | -- | -- | -- | -- | -- | -- | -- | -- | -- | -- | -- | -- | -- | -- | -- | -- | -- | -- | -- | -- | -- | -- | -- | LG7 | -- | LG7 | -- |

| snp_B1350 | snp_B1348 | snp_B1303 | snp_B1261 | snp_B1237 | snp_B1126 | snp_B1124 | snp_B1100 | snp_B1041 | snp_B1014 | snp_B1011 | snp_B0996 | snp_B0957 | snp_B0950 | snp_B0948 | snp_B0943 | snp_B0942 | snp_B0877 | snp_B0868 | snp_B0811 | snp_B0794 | snp_B0778 | snp_B0711 | snp_B0689 | snp_B0638 |
|---|---|---|---|---|---|---|---|---|---|---|---|---|---|---|---|---|---|---|---|---|---|---|---|---|
| 12 | 12 | 12 | 12 | 12 | 12 | 12 | 12 | 12 | 12 | 12 | 12 | 12 | 12 | 12 | 12 | 12 | 12 | 12 | 12 | 12 | 12 | 12 | 12 | 12 |
| -- | LG7 | -- | -- | -- | -- | -- | -- | -- | -- | LG7 | -- | -- | -- | -- | -- | -- | -- | -- | -- | LG7 | -- | -- | -- | -- |

| snp_B0538 | snp_B0434 | snp_B0419 | snp_B0364 | snp_B0354 | snp_B0340 | snp_B0327 | snp_B0325 | snp_B0317 | snp_B0291 | snp_B0281 | snp_B0271 | snp_B0239 | snp_B0225 | snp_B0212 | snp_B0199 | snp_B0176 | snp_B0127 | snp_B0113 | snp_B0098 | snp_B0078 | snp_B0075 |
|---|---|---|---|---|---|---|---|---|---|---|---|---|---|---|---|---|---|---|---|---|---|
| 12 | 12 | 12 | 12 | 12 | 12 | 12 | 12 | 12 | 12 | 12 | 12 | 12 | 12 | 12 | 12 | 12 | 12 | 12 | 12 | 12 | 12 |
| -- | -- | -- | -- | -- | -- | -- | -- | -- | -- | -- | -- | -- | -- | -- | LG7 | -- | -- | -- | -- | -- | -- |
